# Genome Profiling of SARS-CoV-2 in Indonesia, ASEAN, and the Neighbouring East Asian Countries: Features, Challenges, and Achievements

**DOI:** 10.1101/2021.07.06.451270

**Authors:** Inswasti Cahyani, Eko W. Putro, Asep M. Ridwanuloh, Satrio H.B. Wibowo, Hariyatun Hariyatun, Gita Syahputra, Gilang Akbariani, Ahmad R. Utomo, Mohammad Ilyas, Matthew W. Loose, Wien Kusharyoto

**Affiliations:** School of Life Sciences, University of Nottingham, Nottingham, United Kingdom; Research Centre for Biotechnology, Indonesian Institute of Sciences (Lembaga Ilmu Pengetahuan Indonesia/LIPI), Bogor, Indonesia; PT. PathGen Diagnostik Teknologi, Cibinong, Bogor, Indonesia; Biomedical Postgraduate Program, Universitas Yarsi, Jakarta, Indonesia; Molecular Pathology Research Group, School of Medicine, University of Nottingham, United Kingdom; National Institute of Health Research and Development, Ministry of Health, Republic of Indonesia; Department of Pharmacology and Clinical Pharmacy, Faculty of Pharmacy, Universitas Muhammadiyah, Purwokerto, Indonesia; National Institute of Health Research and Development, Ministry of Health (NIHRD); Eijkman Institute for Molecular Biology, Jakarta 10430, Indonesia (EIJK); Diagnostic and Research Center of Infectious Diseases, Medical Faculty, Andalas University (FKUA); Institute of Tropical Disease, Universitas Airlangga (RSDS-SCVTD-UNAIR); Tanjungpura University Hospital (TUH); Faculty of Medicine, Universitas Indonesia (FKUI); West Java Health Laboratory-ITB-UNPAD (WJHL-ITB); Universitas Sebelas Maret (UNS), Rumah sakit Universitas Sebelas Maret (RS-UNS) Surakarta, National Institute of Health Research and Development, Indonesian Ministry of Health; Medical Research Center, Faculty of Medicine, Syarif Hidayatullah State Islamic University Jakarta (SHSIU); Genomik Solidaritas Indonesia Laboratorium (GSI); Genetics Working Group (Pokja Genetik) Faculty of Medicine, Public Health and Nursing Universitas Gadjah Mada (UGM); Stem Cell Research and Development Center; Faculty of Medicine, Universitas Sumatera Utara (USU); Mochtar Riady Institute for Nanotechnology, Universitas Pelita Harapan (MRIN-UPH); Clinical Microbiology Lab RSPTN Universitas Hasanuddin; Indonesian Research Center for Veterinary Science/BBalitvet; Biosafety Level-3 Laboratory, Indonesian Institute of Sciences (LIPI); Stem Cell Lab, Universitas Pembangunan Nasional Veteran Jakarta (FKUPNVJ)

**Keywords:** ASEAN, COVID-19, genomic surveillance, GISAID, nanopore, NICCRAT, SARS-CoV-2, variant of concern, whole-genome sequencing

## Abstract

A year after the World Health Organisation (WHO) declared COVID-19 as a pandemic, much has been learned regarding SARS-CoV-2 epidemiology, vaccine production, and disease treatment. Whole-genome sequencing (WGS) has played a significant role in contributing to our understanding of the epidemiology and biology of this virus. In this paper, we investigate the use of SARS-CoV-2 WGS in Southeast and East Asia and the impact of technological development, access to resources, and demography of individual countries on its uptake. Using Oxford Nanopore Technology (ONT), Nottingham-Indonesia Collaboration for Clinical Research and Training (NICCRAT) initiative has facilitated collaboration between the University of Nottingham and a team in Research Centre for Biotechnology, Indonesian Institute of Sciences (*Lembaga Ilmu Pengetahuan Indonesia/LIPI*) to carry out a small number of SARS-CoV-2 WGS in Indonesia. The ONT offers sequencing advantages that fit within the Indonesian context. Analyses of SARS-CoV-2 genomes deposited on GISAID from Southeast and East Asian countries reveal the importance of collecting clinical and demographic metadata and the importance of open access and data sharing. Lineage and phylogenetic analyses per 1 June 2021 found that: 1) B.1.466.2 variants were the most predominant in Indonesia, with mutations in the spike protein including D614G at 100%, N439K at 99.1%, and P681R at 69.7% frequency, 2) The variants of concern (VoCs) B.1.1.7 (Alpha), B.1.351 (Beta), and B.1.617.2 (Delta) were first detected in Indonesia in January 2021, 3) B.1.470 was first detected in Indonesia and spread to the neighbouring regions, and 4) The highest rate of virus transmissions between Indonesia and the rest of the world appears to be through interactions with Singapore and Japan, two neighbouring countries with a high degree of access and travels to and from Indonesia.

## Introduction

Coronaviruses belong to a large family of RNA viruses that usually cause respiratory illnesses. Most of the time, diseases caused by coronaviruses, such as HCoV-229E, OC43, NL63, and HKU1, are mild^1^. Nonetheless, in the past two decades, more deadly forms of coronaviruses have emerged, including Severe Acute Respiratory Syndrome Coronavirus (SARS-CoV)^2^ and the Middle East Respiratory Syndrome Coronavirus (MERS)^3^. Most recently, COVID-19 emerged and was declared as a global pandemic by the World Health Organization (WHO) on 11^th^ March 2020^4^. This disease is caused by a novel previously unreported coronavirus strain subsequently named Severe Acute Respiratory Syndrome Coronavirus-2 (SARS-CoV-2)^5^.

COVID-19 was first reported in Wuhan in early December 2019^6^. In early January 2020, the Chinese government released the first genome sequence of SARS-CoV-2 (*i*.*e*. labelled as WH-Human_1)^7,8^. By the end of February 2020, the infection had been reported in 51 countries with almost 84,000 cases^9^. Despite being one of the world’s most populated countries, with more than 260 million people, and no travel restrictions in place at the time, Indonesia only diagnosed and announced its first case of infection on 2^nd^ March 2020^10^. However, neighbouring countries, such as Singapore and Malaysia, had reported their first cases of infection on 23^rd^ and 25^th^ January 2020, respectively. Indonesia’s later detection and action might have been influenced to a large degree by some presumed confidence in the society that the infection would not easily spread in the tropics^11^. There was a widely-held belief that the hot tropical climate and high intensity of UV rays in Indonesia would be sufficient to hinder SARS-CoV-2 survival and spread in the air, much like the first SARS epidemic where cases were low compared to its originating place in China^2,11,12^. Therefore, before March 2020, Indonesia only carried out early testing and screening of individuals with symptoms and travel links^13,14^.

In hindsight, the indifference toward COVID-19 when it was still an epidemic was not exclusive to Indonesia, as many governments around the globe shared a similar lack of anticipation and mitigation to prevent the COVID-19 epidemic from turning into a pandemic^15^. The Association of Southeast Asian Nations (ASEAN) accommodates free cross-border travels for its members, with Indonesia having the largest area and population size. Indonesia also has bilateral travel agreements that make travel more accessible to other East Asian countries, such as Japan and China. Global travel has been a significant contributing factor to the rapid spread of COVID-19 and establishing the pandemic^16^. Therefore, we sought to compare the genomic epidemiology and metadata of SARS-CoV-2 between ASEAN member countries and others in the region.

As of today, more than a year after Indonesia’s first COVID-19 case was announced (*i*.*e*., early-March 2020), it is thought that the number of cases in Indonesia is still significantly underestimated. Besides the utility of screening and tracing for containing and reducing transmission, there is also a need to pinpoint and follow variants of SARS-CoV-2 that might worsen the pandemic. World Health Organization defines a variant to be a Variant of Interest (VoI) if it has changed phenotypically from the reference genome (or shown phenotypic implications) and has been identified to cause community transmissions or detected in multiple countries^17^. A VoI can become a Variant of Concern (VoC) if it has been demonstrated to 1) increase transmissibility or detrimental changes in COVID-19 epidemiology; 2) an increase in virulence or change in clinical disease presentation; or 3) decrease in the effectiveness of public health and social measures or available diagnostics, vaccines, and therapeutics^17^. The existence and potential emergence of these variants highlight the need for reliable, easy-to-set-up, rapid, and cost-effective genome sequencing technology in Indonesia. We showed on a small number of samples (*n*=12) how the Oxford Nanopore Technology^®^ (ONT) platform could be used for whole-genome sequencing (WGS) of SARS-CoV-2 in the Indonesian context. Moreover, this report discusses the types and transmission patterns of SARS-CoV-2 variants in Indonesia in relation to ASEAN and other Asian countries until 1 June 2021. We highlighted how the knowledge obtained from the WGS of SARS-CoV-2 is pivotal in the understanding and tackling of COVID-19 spread in Indonesia and its surrounding regions.

## Results

### A. The rates of WGS in ASEAN and its neighbouring countries correlate to differences in technology development, access to resources, and demography

Whole-genome sequencing of viruses that play a role in an outbreak, epidemic and/or pandemic is emerging as a crucial tool in controlling and mitigating the spread of disease^18–21^. Whole-genome sequencing can inform epidemiological scientists on the nature, behaviour, and transmission patterns of viruses^22^. This information can support decision-makers in taking appropriate actions. In the era of next-generation sequencing (NGS), WGS for epidemiological monitoring should be straightforward, in principle, with the availability of many tools and relatively low cost of sequencing compared to one or two decades ago. Some examples of WGS that applied these NGS principles in (or after) an outbreak were in the case of the Zika virus in Brazil^18,19^, Ebola in West Africa^20^, and more recently, a mumps outbreak in Canada ^21^. Whole-genome sequencing is also at the heart of global research for vaccines and treatment discoveries of the COVID-19 pandemic and for surveying its VoCs^22^.

Unfortunately, WGS, which uses cutting-edge sequencing technology, typically means that only developed countries with access and resources to technology can readily use it in a pandemic. Figure 1A shows that despite being the largest continent in the world, Asia has submitted only about 6% of the total SARS-CoV-2 genomes deposited on GISAID per 1 June 2021, with Japan submitting the most at just over 44% (Figure 1B-1C). Indonesia contributed to 1.7% of genomes submitted from Asia, relatively behind its close neighbour, the Philippines, at 4% (Figure 1B-1C). Indonesia also experienced the highest total number of positive cases in the Southeast and East Asian region (Figure 2A). However, the rate of SARS-CoV-2 WGS did not follow the case rate as the number of genomes submitted to GISAID and Indonesia’s sequencing rate were relatively low (Figure 1C and 2B, respectively). Some of ASEAN’s neighbours, such as Hong Kong, Japan, and South Korea, tended to have significantly higher sequencing rates (Figure 1C and 2B), most likely because of better access to technology and resources. Better availability of resources also seems to help these countries to push their COVID-19 test rate to parallel their population size (Figure S1). Meanwhile, other countries with similar development status to Indonesia, such as Thailand, might fare better because of relatively smaller numbers of cases and population size (Figure 2A). In contrast to Indonesia, the Philippines relatively tripled its sequencing rate despite having the second-highest number of positive cases in the region (Figure 2).

**Figure 1.**
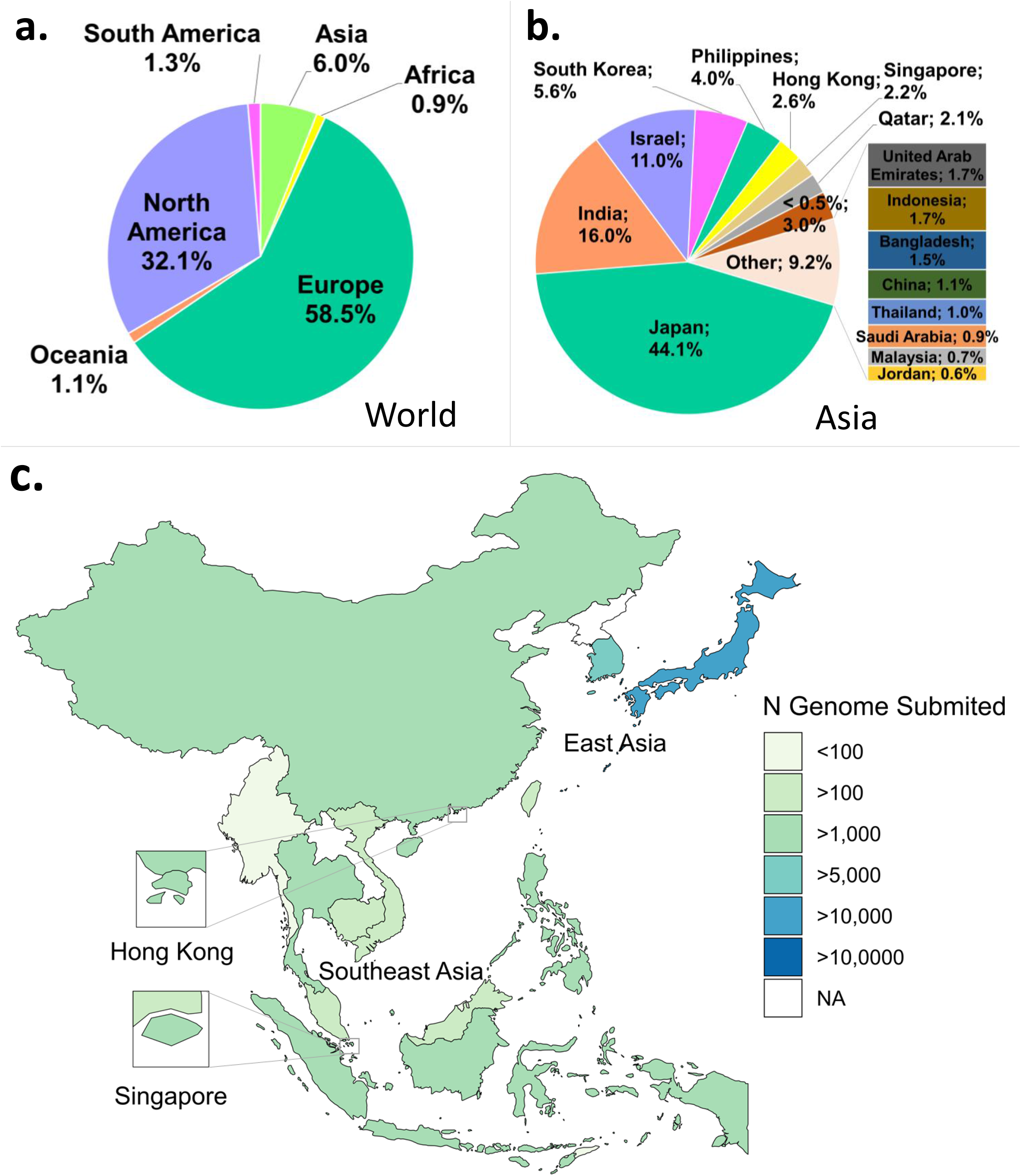
Profiles of SARS-CoV-2 WGS in Indonesia, its neighbouring countries in Asia, and the world. Metadata was downloaded from GISAID^50^ per 1 June 2021. A) Asia only accounts for about 6% of the total global number of genomes submitted in the world. B) In Asia, Japan is the country with the highest number of submitted genomes. C) Distribution map of the number of genomes submitted to GISAID in ASEAN and East Asia region.

**Figure 2.**
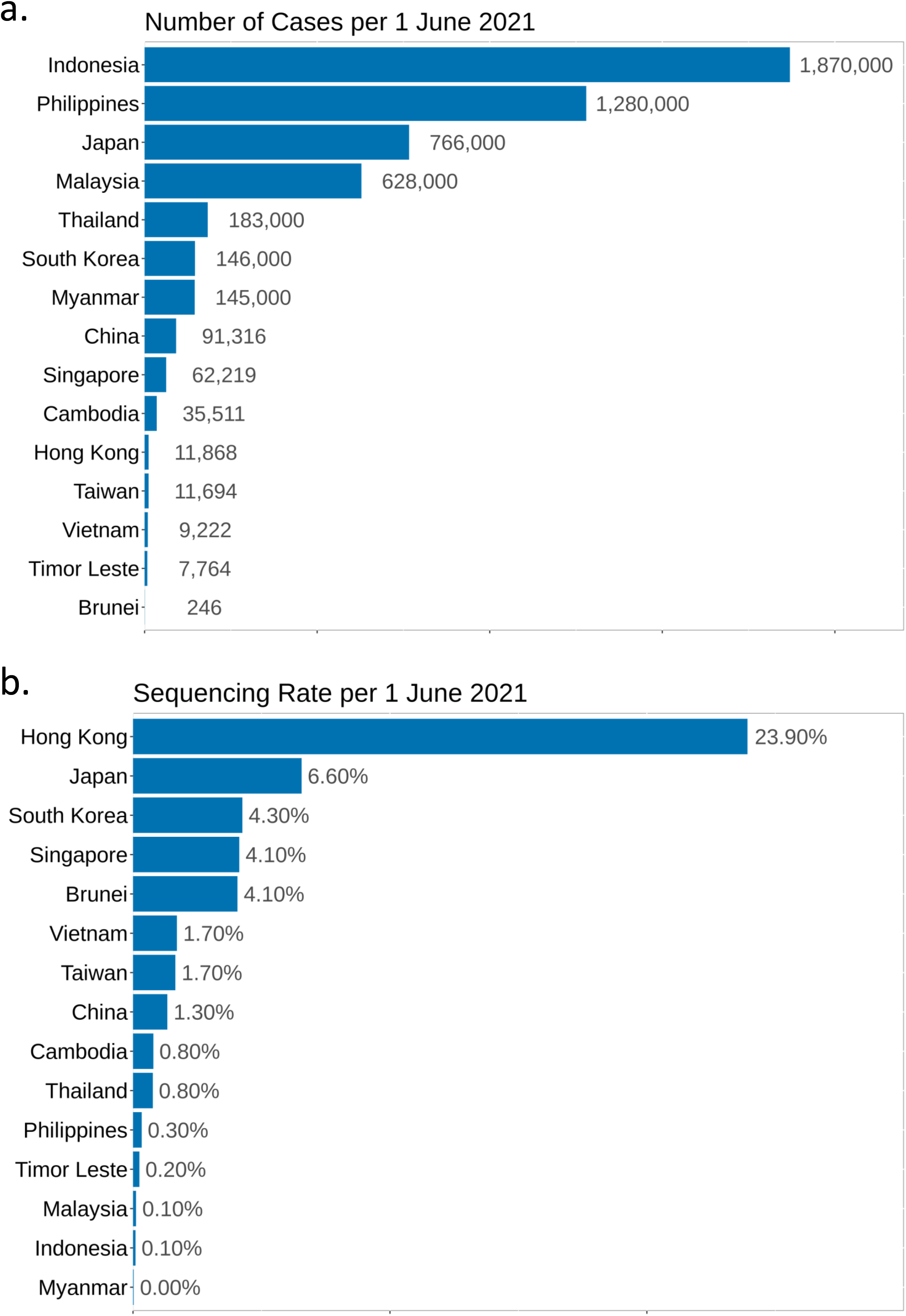
COVID-19 cases and SARS-CoV-2 sequencing rate. A) Confirmed positive number of COVID-19 cases in ASEAN and East Asia region, where Indonesia currently holds the highest number of cases, followed by the Philippines. B) The rates of WGS of countries in ASEAN and East Asia region, compared to the number of positive cases in Figure 1A; Hong Kong sequenced almost 24% of its positive cases and led the number in the region.

There has been a lot of debate on the ideal sequencing rate and whether it should be based on the population size positive cases or both^23^. As of 1 June 2021, Indonesia had sequenced 0.1% of its positive cases (Figure 2B). However, this sequencing rate was calculated on a known case number which might have been underestimated due to suboptimal testing rates (Figure S1).

### B. Establishment of WGS effort in Indonesia

In January 2021, the Indonesian Ministry of Health, working together with the Ministry of Research and Technology/National Agency for Research and Innovation, started a nationwide network for SARS-CoV-2 WGS for pandemic surveillance, bringing together laboratories and research, and academic institutions^24^ (Table 1). The network was expected to accelerate and scale-up SARS-CoV-2 WGS, informing the government on the genomic epidemiology of the pandemic in a timely manner^24^. The creation of nationwide consortia/network for SARS-CoV-2 WGS has been shown to work in the United Kingdom with the COG-UK consortium^25^. The UK is currently the second biggest contributor of SARS-CoV-2 genomes on GISAID after the United States of America^26^, sequencing about 10% of its positive cases^25^. Therefore, emulating some aspects of this consortium in Indonesia is feasible while considering the local context, such as demography and logistics.

**Table 1.** List of institutions in Indonesia and the numbers of submitted genomes to GISAID (Per 1 June 2021)

| Institution | Sequencing Platform |  |  |  |  |
| --- | --- | --- | --- | --- | --- |
|  | Illumina only | Illumina + Sanger | ONT | Sanger only | Sub Total |
| Andalas University | 108 | - | - | - | 108 |
| Eijkman Institute for Molecular Biology (including with the National Institute of Health Research & Development and Warmadewa University) | 651 | 38 | - | - | 689 |
| Genomik Solidaritas Indonesia Laboratorium | - | - | 1 | 1 | 2 |
| Indonesian Research Center for Veterinary Science/BBalitvet | - | - | 2 | - | 2 |
| Institute of Tropical Disease, Universitas Airlangga (including with Dr. Soetomo Hospital) | 126 | - | 10 | - | 136 |
| LIPI | - | - | 82 | - | 82 |
| NICCRAT-LIPI | - | - | 12 | - | 12 |
| Mochtar Riady Institute for Nanotechnology | - | - | - | 17 | 17 |
| National Institute of Health Research & Development | 473 | 7 | - | 15 | 495 |
| Syarif Hidayatullah State Islamic University | - | - | 14 | - | 14 |
| Tanjungpura University Hospital | 5 | - | 11 | - | 16 |
| Universitas Gadjah Mada | 66 | - | - | - | 66 |
| Universitas Indonesia | - | - | 15 | - | 15 |
| Universitas Pembangunan Nasional Veteran Jakarta | 9 | - | - | - | 9 |
| Universitas Sebelas Maret | 10 | - | - | - | 10 |
| Universitas Sumatera Utara | 23 | - | - | - | 23 |
| West Java Health Laboratory-ITB-UNPAD | 114 | - | 2 | - | 116 |
| **Total** | **1585** | **45** | **149** | **33** | **1812** |

Increasing the WGS of SARS-CoV-2 in Indonesia has been challenging. Up-to-standard laboratories are primarily found in the dense west-to-middle parts of Indonesia, such as Java, Sumatera, and Bali. The high-cost and state-of-the-art devices required for NGS are also a limiting factor as well as the expertise and infrastructure required. Oxford Nanopore Technology® offers a solution to these issues by providing a portable, scalable, and lower cost of sequencing platform compared to other NGS technologies^27^. As a result, ONT can potentially be the workhorse for WGS in Indonesia. In addition, the ARTIC network^28,29^ provided protocols, primer designs, and analysis pipelines and tools capable of running on basic computing infrastructure.

The Nottingham-Indonesia Collaboration in Clinical Research and Training (NICCRAT) initiative, established in 2019^30,31^ aims to foster partnership between the University of Nottingham and several Indonesian research and academic institutions in the field of health and clinical science^30,31^. As COVID-19 became a global pandemic in early 2020, the initiative contributed some expertise in SARS-CoV-2 WGS in Indonesia^31^. Collaboration between the University of Nottingham and some researchers in LIPI (NICCRAT-LIPI team) utilised the MinION sequencing platform to sequence 12 SARS-CoV-2 genomes that have been published to GISAID. The MinION sequencing platform offers speed, portability, relatively low cost, and has well-established protocols. These features are particularly advantageous within Indonesia, where specialized molecular biology laboratories that can handle WGS are not widely available. This project also benefits through the involvement of the University of Nottingham in the COG-UK consortium^25^. An online skill transfer workshop of SARS-CoV-2 WGS using the ONT platform was carried out once by the University of Nottingham team to some researchers in LIPI on 17 April 2020. Meanwhile, as 1 June 2021, a total of 94 whole genomes have been deposited by the LIPI-wide SARS-CoV-2 WGS team. Until 1 June 2021, the number of SARS-CoV-2 genomes from Indonesia submitted to GISAID is 1812 (Table 1).

### C. Comparison of clinical and demographic metadata of deposited genomic sequence in GISAID between Indonesia and its neighbouring countries

As part of SARS-CoV-2 WGS, recording sample metadata is a vital aspect to build a clearer picture of genomic epidemiology^32^. However, this is not always easy nor practical to achieve, especially when the institution collecting samples differs to the one sequencing them, such as a hospital and a sequencing research facility, respectively. Moreover, countries have different regulations concerning privacy, data access, and sharing. In this report, we categorize the metadata accompanying genomes on GISAID as clinical (*e*.*g*. disease severity, mortality, hospitalization rate, and sample type) or demographic (*e*.*g*. age, gender, and residency) metadata. Indonesia and its neighbours recorded and dealt with sample metadata in different methods, which influence the interpretation of WGS output.

Indonesia performs well in terms of metadata completeness, both in clinical (Table 2, Figure 3A) and demographic metadata (Figure 3B-3C) when compared to other ASEAN and neighbouring countries. Over 70% of the Indonesian genomes have the corresponding clinical metadata (Table 2). In addition, Indonesia also recorded the sources of the majority of samples (*i*.*e*., nasal and oropharyngeal swab) used in WGS, which is lacking in many other large datasets (*e*.*g*., COG-UK) (Figure 3A). On the other hand, many countries, including Japan as the top contributor of SARS-CoV-2 genomes shown in Figure 1B, do not have accompanying clinical and/or demographic metadata submitted to GISAID (Figure 3B-3C). Most likely, these metadata are stored in a local/national repository but not reported to GISAID to comply with respective data-sharing regulations.

**Table 2.** Clinical metadata of genomes in Indonesia and Southeast and East Asia (based on data downloaded from GISAID per 1 June 2021)

| Country | Live | Deceased | Not Hospitalized/<br>Asymptomatic | Hospitalized/<br>Symptomatic | Total No.<br>of<br>Genomes | %<br>Genomes<br>with<br>Metadata |
| --- | --- | --- | --- | --- | --- | --- |
| Brunei | 0 | 0 | 0 | 0 | 10 | 0.0% |
| Cambodia | 278 | 0 | 0 | 0 | 281 | 98.9% |
| China | 239 | 1 | 2 | 153 | 1,220 | 32.4% |
| Hong Kong | 601 | 26 | 0 | 323 | 2,837 | 33.5% |
| Indonesia | 811 | 25 | 26 | 477 | 1,806 | 74.1% |
| Japan | 114 | 16 | 62 | 9,114 | 50,241 | 18.5% |
| Malaysia | 476 | 7 | 3 | 64 | 745 | 73.8% |
| Myanmar | 2 | 0 | 0 | 0 | 41 | 4.9% |
| Philippines | 3,593 | 73 | 0 | 11 | 4,295 | 85.6% |
| Singapore | 0 | 0 | 0 | 553 | 2,576 | 21.5% |
| South Korea | 25 | 9 | 0 | 0 | 6,216 | 0.5% |
| Taiwan | 25 | 0 | 0 | 74 | 193 | 51.3% |
| Thailand | 8 | 0 | 0 | 3 | 1,404 | 0.8% |
| Timor-Leste | 0 | 0 | 0 | 0 | 19 | 0.0% |
| Vietnam | 79 | 0 | 0 | 3 | 158 | 51.9% |

**Figure 3.**
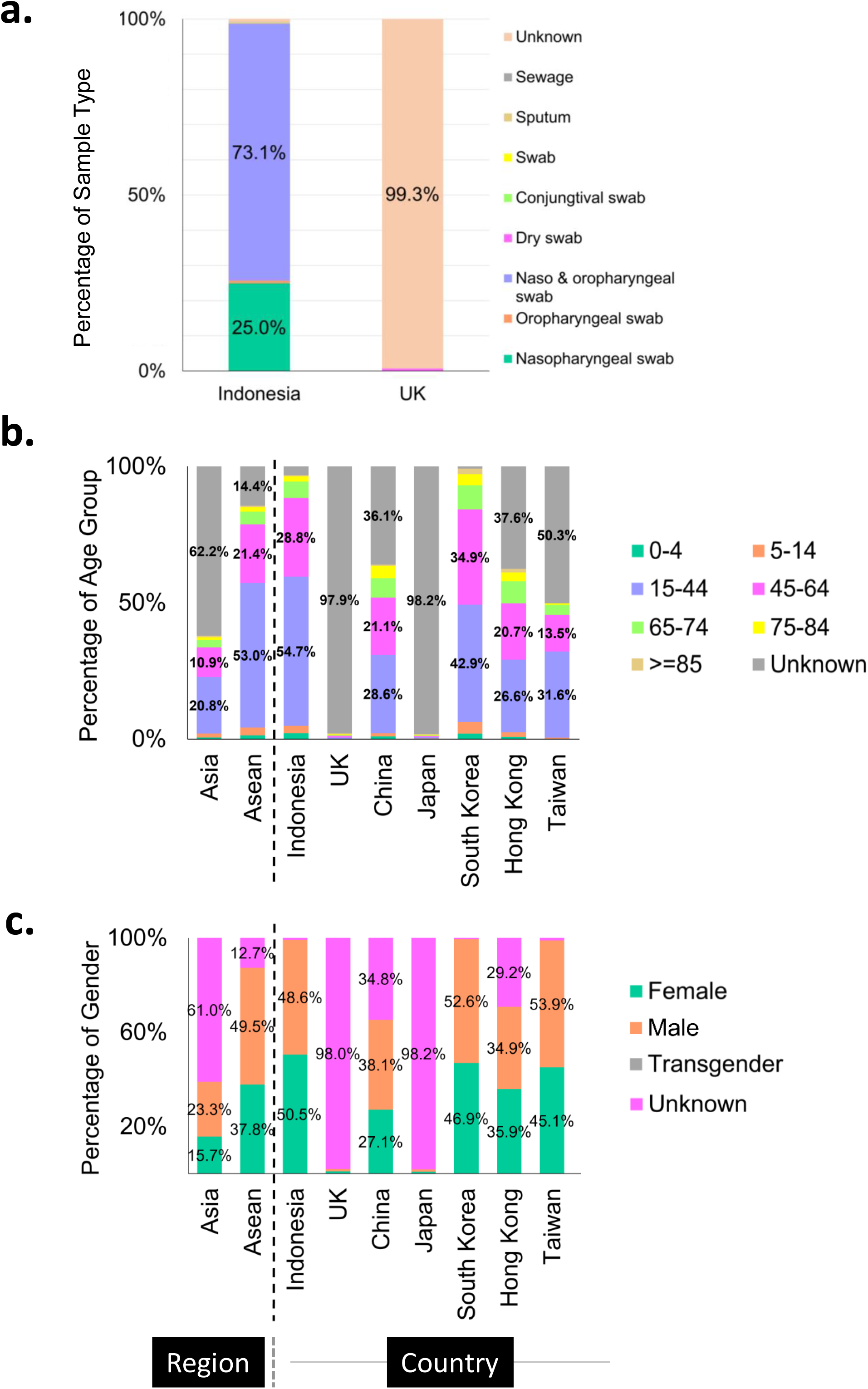
Metadata profiles between ASEAN, Asian countries, and the UK; Indonesia is relatively thorough in collecting and registering its genomic metadata, comparable to those of South Korea. A) Proportion of sample sources used for sequencing; Indonesia obtained most of its sequencing samples from swab while the UK did not provide this information to GISAID. B) Distribution of the age groups of patient samples. Indonesia submitted almost complete metadata of this category to GISAID, as well as South Korea. C) Distribution of the gender groups of patient samples. Indonesia submitted almost complete metadata of this category to GISAID, as well as South Korea and Taiwan.

The majority of the Indonesian samples came from hospitalized patients (Table 2). More than half of these sequenced cases in Indonesia belong to the productive age group of 15– 44-year-olds and the second-largest group (29%) being 45-64-year-olds (Figure 3B). This data can merely represent the age distribution of the Indonesian population (Table S2), as it is also reflected in the population age distribution of South Korea (Figure 3B, Table S2). However, it is hard to say for countries, such as Hong Kong and China, as their unknown categories of this metadata are relatively large (Figure 3B). The same assumption could also be applied to the metadata distribution of gender in Indonesia, where 50.5% of the infection came from female patients (Figure 3C), in correlation with the 50:50 gender distribution in the Indonesian population (Table S2). However, a study shows that the male sex of higher age with comorbidity may have more prevalence in requiring intensive care treatment and mortality^33,34^.

In conclusion, Indonesia has successfully recorded and shared its genomic metadata on GISAID, despite the sporadic origins of the samples. However, it is notable that with the different national and international rules governing personal information and data sharing, metadata information should best be made available and accessible to selected relevant parties (*e*.*g*., health ministry, medical professionals, and epidemiological researchers). This way, the metadata will put the SARS-CoV-2 WGS output into context and draw as correct conclusions as possible to deal with the COVID-19 pandemic.

### D. The distribution of variants by PANGO lineages in Indonesia compared to its neighbours and the rest of the world

Lineage annotation by Phylogenetic Assignment of Named Global Outbreak Lineages (PANGO Lineages; PANGOLIN) presents a dynamic nomenclature of the SARS-CoV-2^35^. Because it labels the variants that are circulating and active, it can track the transmission of SARS-CoV-2 based on sequence differences while also considering new virus diversity^35^. Therefore, we use the PANGOLIN annotations analysed by GISAID and included in the metadata of the respective genomes (Figure 4 and S2). In the following sections, we also use the WHO’s nomenclature of Greek alphabets^36^, when generally discussing lineages that have become VoCs.

**Figure 4.**
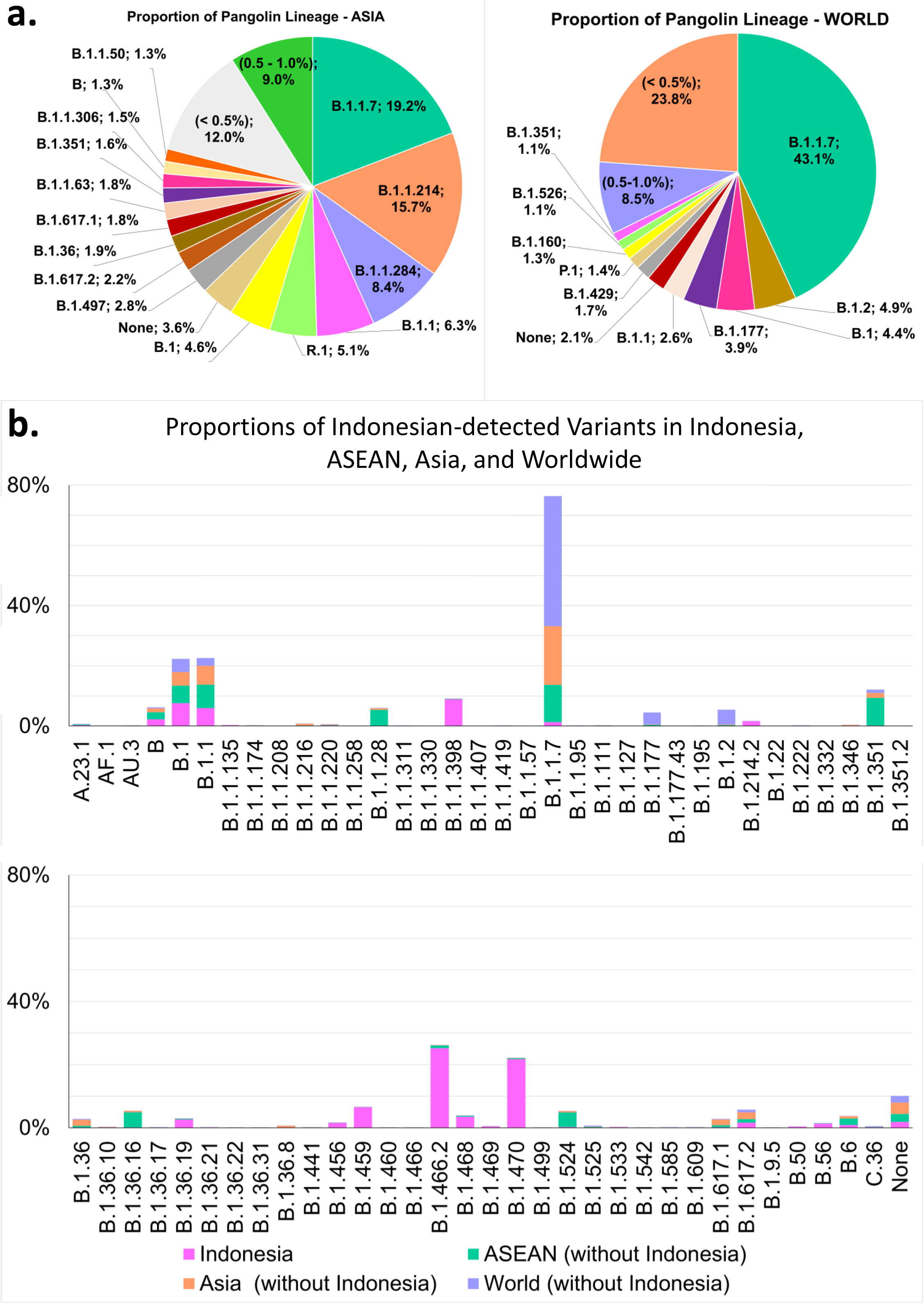
The distribution of SARS-CoV-2 variants based on PANGOLIN^69^ per data downloaded from GISAID on 1 June 2021. A) In Asia, B.1.1.7 and B.1.1.214 are the most common variants found with a total of more than 35% between them (left panel). B.1.1.7 is also the dominant variant of the submitted genomes in the world (right panel). B) Distributions of Indonesia-detected variants as proportions in Indonesia, ASEAN, Asia, and the rest of the world. Variants B.1.466.2 and B.1.470 (lower panel) were the two most prevalent variants in Indonesia.

Rambaut *et*.*al*. hypothesized that lineage A was the founding lineage^35^. However, currently, the B lineage, in particular B.1.1.7 is the most prevalent variant both in Asia (*i*.*e*., 19.2%) and the world (*i*.*e*., 43.1%) (Figure 4A). Lineage B.1.1.7 (the Alpha variant^36^) was first detected in Kent in the United Kingdom as early as September 2020^37^ and has now become a VoC^36^. Previous data downloaded in early February 2021 shows that the Alpha variant had not been significantly detected in Asia and only accounted for 8.3% of the sequenced genomes in the world at that time (Figure S2). The most prevalent variants in early February 2021 in Asia and the world were B.1.1.284 and B.1.177, respectively (Figure S2). However, Indonesia had detected the Alpha variant along with two other VoCs, Beta and Delta, in January 2021 (Table 3).

**Table 3.** The first detection dates of VoCs in Indonesia per 1 June 2021.

| VoC | Number of<br>Genomes | First Detected Case |  |
| --- | --- | --- | --- |
|  |  | Accession ID | Date |
| B.1.1.7 (Alpha) | 23 | hCoV-19/Indonesia/SS-NIHRD-WGS00427/2021 | 5/01/2021 |
| B.1.351 (Beta) | 4 | hCoV-19/Indonesia/BA-NIHRD-WGS00725/2021 | 25/01/2021 |
| B.1.617.2 (Delta) | 29 | hCoV-19/Indonesia/JK-NIHRD-WGS00007/2021 | 7/01/2021 |

Figure 4B shows the frequency distribution of Indonesia-detected variants as a proportion of the total number of cases in Indonesia (in pink), ASEAN area (in green), Asia continent (in orange), and the world (in purple). Ten of the B.1.1.7 variant had been detected as of 24 April 2021^38^ but it has not significantly influenced variant distribution in Indonesia, in contrast to the world’s proportion (Figure 4B and S3, upper panels). The two most predominant variants in Indonesia per 1 June 2021 were B.1.466.2 and B.1.470 at about 25% (456 genomes) and 21% (392 genomes), respectively (Figure 4B, lower panel). Moreover, B.1.470 variant was first detected in Indonesia on 9 April 2020 and then spread to near- and far-neighbouring countries, such as Malaysia, South Korea, and Japan^39^.

All B.1.466.2 variants in Indonesia carried the D614G spike protein mutation (Table 4). The G614 virus was shown to have an increased capacity to enter ACE2-expressing cells^40^ and to enhance its replication in human respiratory cells^41^ compared to the original D614 virus. These G614 mutation characteristics were hypothesized to confer a moderate level of infectivity and transmissibility and could be the reasons it maintained its presence in the globally circulating variants^42,43^. The B.1.466.2 variant also frequently carried two other mutations: 1) N439K at 99.1% frequency and 2) P681R at 69.7% frequency (Table 4). Studies on viruses with these two mutations also showed correlation to increased infectivity, by way of ACE2 receptor binding, and viral transmissibility^43–46^. Moreover, the P681R mutation was frequently detected in the more-infectious B.1.617.2 (Delta) variant^46^. Therefore, the high prevalence of all three mutations in the Indonesian B.1.466.2 variant might have given it a cumulative fitness advantage over the other variants. Furthermore, the B.1.617.2 (Delta) variant carried more mutations in its spike protein, including the two mutations found in the B.1.466.2 variant (Table 4). All the Delta variants had mutations in D614G, P681R, T19R, T478K, and L452R (Table 4).

**Table 4.** Most frequent mutations of SARS-CoV-2 spike protein in B.1.466.2, B.1.617.2 (Delta) and all other variants in Indonesia.

| Amino Acid Change | B.1.466.2 | B.1.617.2 (Delta) | All Variants |
| --- | --- | --- | --- |
| D614G | 100.0% | 100.0% | 94.2% |
| N439K | 99.1% | 0% | 26.5% |
| P681R | 69.7% | 100.0% | 21.0% |
| T19R | 0% | 100.0% | 0% |
| T478K | 0% | 100.0% | 0% |
| L452R | 0% | 100.0% | 0% |

There was another variant, B.50, with only five genomes recorded in Indonesia per 1 June 2021, designated as an Indonesian lineage on the PANGOLIN website (https://cov-lineages.org/)^47^. B.50 variant was also detected in Japan and Germany, both at around 11% of the world’s total^47^. Hence, this variant is not unique to Indonesia. There is so far no variant detected only in Indonesia and not elsewhere in the world (Figure S3). This might mean that the lineages found in Indonesia: 1) accumulated mutations during local transmissions from imported parental variants, or 2) were exported to other countries, particularly its neighbours. It might also be possible that both scenarios occurred intermittently, as border closure for foreigners and quarantine measures for returning citizens were only put into effect from February 2021^48^.

### E. Phylogenetic relationship based on PANGOLIN exists between different Indonesian lineages and those exported to the neighbouring countries

In Indonesia, the earliest dated sample for sequencing was collected by *Universitas Airlangga* (Airlangga University) in Surabaya on 12 March 2020 (Figure 5A, Table S3). We used this sequence as the root in the phylogenetic tree analysis (Figure 5B). Nonetheless, the first Indonesian cases (*i*.*e*. a cluster of two) were detected in Jakarta and officially announced on 2 March 2020^10,13,14^. As these patients immediately underwent quarantine and hospitalization, it was less likely that the first sequencing sample was the result of direct transmission from this cluster, only ten days apart. This implies that COVID-19 might have been transmitted and circulated locally and presumably undetected (as modelled by De Salazar *et*.*al*.^49^), prior to these first official cases.

**Figure 5.**
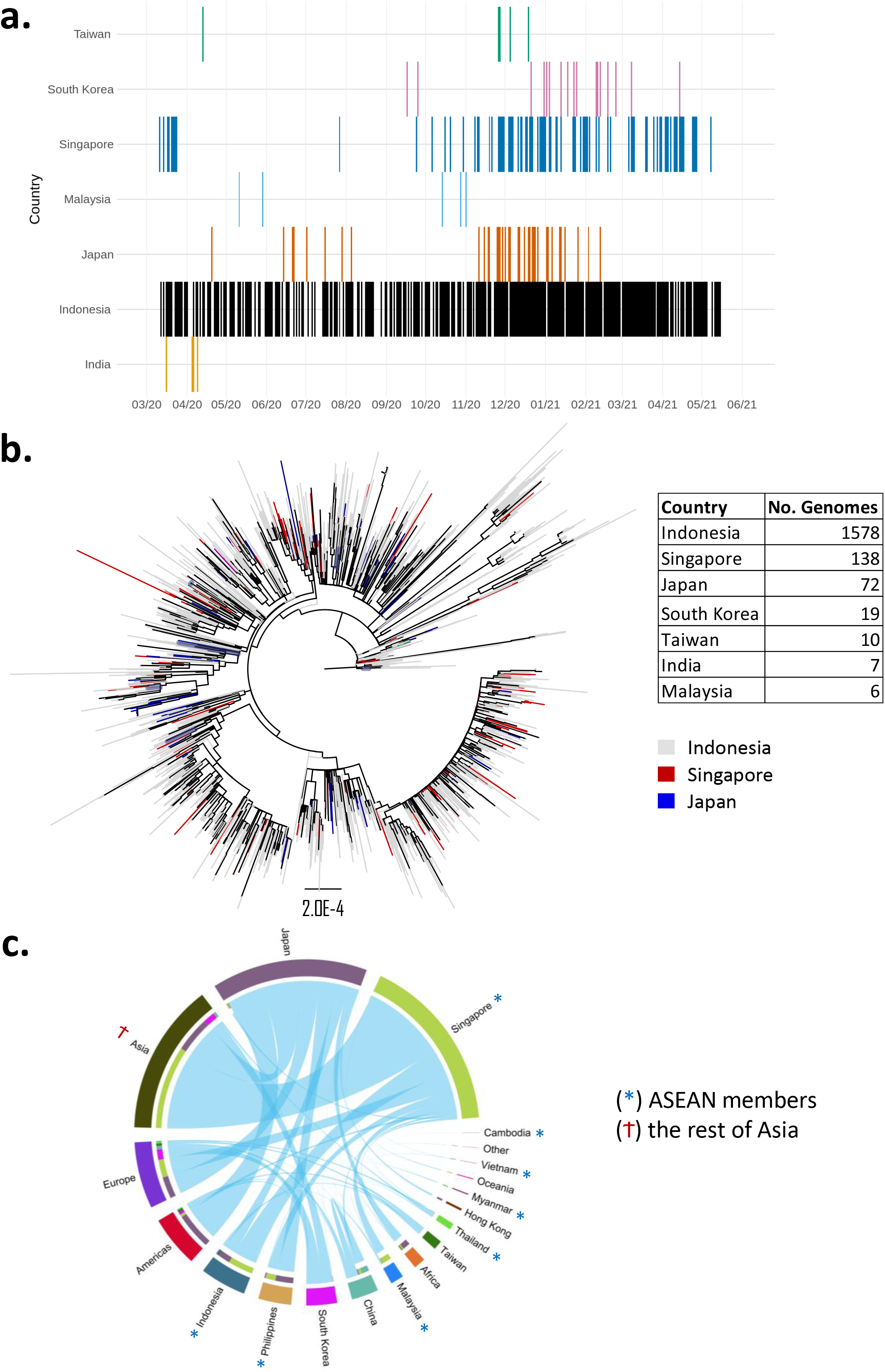
Phylogenetic analyses of the locally transmitted and exported Indonesian genomes (*n* = 1830) based on data downloaded from GISAID on 1 June 2021. A) Timeline of Indonesian-associated variants by country; exported transmissions were primarily within Southeast and East Asia. B) Phylogenetic tree rooted to the earliest Indonesian genome shows the Indonesian lineages in grey, spread across the tree. Tree was ordered by nodes, with Singapore and Japan represented by red and blue, respectively. Scale of branch length is given. C) Circular plot shows associations of variants (*i*.*e*., exported/imported transmissions) between different countries in the world; transmissions from Indonesia were the highest to Singapore and Japan, both in the number of events and variant types.

In terms of Indonesian variants being associated with those in other countries (*i*.*e*., country of exposure in the metadata was stated to be Indonesia and/or other mentions of a link to Indonesia), a few near- and far-neighbouring countries are on the list: Singapore, Malaysia, South Korea, Taiwan, Japan, and a bit further away in the region, India (Figure 5A, Figure S4). We term these Indonesian-associated variants as exported variants that occurred in an intermittent manner related to travel and/or contact history with Indonesia (Figure 5A, Figure S4). Since the beginning of March 2020, Indonesian-exported variants could be found in Singapore in parallel to the local transmission of these variants within Indonesia (Figure 5A). Japan also shows a similar pattern to a lesser degree (Figure 5A). This could also suggest that an ongoing, multi-way transmission patterns have existed between these countries, facilitated by relatively easy access and travel between them, and considering the high level of economic and industrial collaborations. The fact that these associated variants still exist up to recently in these countries highlights the importance of genomic surveillance to inform border control policies during a pandemic. Meanwhile, based on the downloaded data per 1 June 2021, there was only one imported Indonesian case from Saudi Arabia (*i*.*e*. EPI_ISL_576387)^50^. However, this seemingly rare, imported case was likely due to a low rate of contact tracing in Indonesia.

All the sequenced genomes of SARS-CoV-2 in Indonesia plus the exported variants were analysed to measure the phylogenetic relationship between them (*n* = 1830). From the total number of exported variants, Singapore holds the highest number of associations at ∼55%, followed by Japan at ∼29% (table in Figure 5B). A phylogenetic tree rooted in the earliest Indonesian genome is presented in Figure 5B, with variants from Singapore and Japan annotated in red and blue, respectively, and Indonesia in grey (Figure 5B). The tree shows a relatively even distribution of Singapore- and Japan-exported variants, branching out from the Indonesian lineages (Figure 5B).

In a more general context, associations between imported/exported lineages in different countries are visualized in Figure 5C (data per 1 June 2021) and Figure S5 (data per 1 February 2021). We filtered 3850 genomes that had different country annotations compared to their countries of exposure, or other location information (Figure 5C)^50^. Worldwide associations show that Singapore was the country having the most links to variants in other countries, both in a total number of genomes and a total number of linked countries, followed by Japan (Figure 5C). These numbers are not surprising considering Singapore is a busy travel hub in the Asian region as well as the world ^51^. Meanwhile, the diverse foreign transmission links in Japan might have been catalysed by the outbreak on-board the Diamond Princess cruise ship in February 2020.

Interestingly, the GISAID data that we downloaded in early February 2021, did not have much information on Singapore variants in terms of their associations to other countries (Figure S5). This contrasts with the newest data (Figure 5C). Thus, it is possible that new genomic metadata was added within the four months prior to our last data download (*i*.*e*., 1 June 2021) and/or backtracked, or both.

## Discussion

The importance of WGS to tracing and understanding pathogens has been demonstrated repeatedly during the ongoing COVID-19 pandemic^27,52,53^. However, establishing distributed networks of a cost-effective variable through sequencing in low- to middle-income countries is challenging. For this purpose, ONT is a robust sequencing platform to do WGS and fits the Indonesian context well. It has a range of outputs that can be suited to the need and budget of each sequencing facility and, crucially, samples can be processed in small or large batches.

One of the difficulties in WGS within a pandemic context is sampling, *i*.*e*., how to obtain obtaining an optimum number of samples representing cases in the population. It is clearly better if sequenced samples come from a wide range of subjects rather than only one group or a subset of individuals^54^. In Indonesia, this has been challenging as samples were mostly non-randomly taken from symptomatic and/or hospitalized patients (Table 2). This will likely affect how the pandemic is observed and may not typify the events occurring in the broader community. More recently, the Indonesian government has been trying to distribute sample collection more evenly by gathering specimens from different provinces across the vast Indonesian archipelago^55^. This is good progress that needs to be maintained and even improved.

In the case of VoCs, the three major variants of Alpha (B.1.1.7), Beta (B.1.351),), and Delta (B.1.617.2) have recently been detected in Indonesia, with the Delta variant quickly rising in the days following our data analyses per 1 June 2021^56^. This pattern is also occurring in the UK, where the Delta variant is gaining number against the Alpha variant^57,58^. The emergence of new variants that may become VoIs and VoCs highlights the importance of genomic surveillance of the SARS-CoV-2 to control the COVID-19 pandemic. Furthermore, functional studies of variants with high prevalence in Indonesia, such as B.1.466.2, B.1.470, and B.50, can be pursued to understand their biological significance. However, such effort needs to be approached with caution as the prevalence of these variants will change with time as the pandemic progresses.

Based on our finding of the Indonesian exported cases to ASEAN countries and beyond, governments in the region could use the existing diplomatic relationships to set more synchronous efforts in border control, trace-and-track, and vaccination scheme, for example. In addition, reducing cross-border spread would conceivably reduce mutation opportunities and thus the emergence of new variants.

Lastly, Indonesia must increase its sequencing rate through the newly established COVID-19 WGS network, in parallel with testing (Figure S1). It is also important to allocate resources for asymptomatic case research to study transmission patterns by asymptomatic individuals, which could influence case numbers^59–61^. These aspects of epidemiological research (*i*.*e*., genomics and diagnostics) will help Indonesia be better prepared and build toward sustainable mitigation of outbreaks. In parallel, the good quality of clinical and demographic metadata recording, despite the difficult terrain faced by many sequencing facilities, should be maintained or even improved to maximize research output.

## Methods

### NICCRAT-LIPI Team Sample collection and WGS

Samples (*n*=12) were collected from nasopharyngeal and oropharyngeal swabs of patients in viral transport medium^62–65^ or DNA/RNA Shield™ (Zymo Research, USA) abides by the Center for Disease Control’s (CDC) guidelines for pathogen inactivation^66^. RNA extraction was done using Viral Nucleic Acid Extraction Kit II (Geneaid Biotech Ltd, Taiwan) according to the manufacturers’ instructions. All samples were tested for the presence of SARS-CoV-2 with an RT–qPCR assay^62–65,67^ using Real-Q 2019-nCoV Detection Kit (Rev.2 (2020.03.25); BioSewoom, Korea) according to the manufacturers’ instructions in a CFX96 Touch Real-Time PCR Detection System (BioRad, USA) machine. Samples with a Ct value of 11-30 were chosen for sequencing^29^.

Sequencing libraries were prepared based on the ARTIC nCoV-2019 sequencing protocol v2 (GunIt) V.2^29^, with minor modifications, using reverse-transcribed cDNA. Primers are obtained from IDT (Version 3)^29^. Libraries were loaded on a MinION device and flow cell. ONT’s MinKNOW software was used to run sequencing. The Rampart software was applied to monitor coverage of each barcoded sample in real-time by running fast base-calling.

### NICCRAT-LIPI Team Bioinformatics processes

A validated pipeline was followed to conduct high accuracy base-calling, demultiplexing, trimming, alignment, and consensus establishment^28^. Medaka workflow was used in generating consensus genome sequences due to the speed and GPU compatibility. A minimum of 20× coverage was set in the pipeline^68^. A concatenated consensus sequences of all barcoded samples which also contained the number of ambiguous nucleotides (Ns) were then further analysed. Assignment of clade membership was done by PANGOLIN (version 27-05-2021)^69^. FASTA files of assembled sequences along with metadata in comma-separated values (.csv) files were prepared for submission to GISAID. The metadata included collection date, location, the origin of samples, passage history, and sequencing technology. Further analysis for possible missing metadata, FASTA sequences, a certain information, and frameshift mutations was undertaken following the result of GISAID quality control checking after submission^70^. In total, 12 genomes were submitted to GISAID^50^ from NICCRAT-LIPI Team.

### Lineage and phylogenetic analyses

Full-genome of SARS-CoV-2 sequences from ASEAN member countries, China, Hong Kong, Japan, South Korea, Taiwan, and the United Kingdom along with their metadata, were downloaded from the GISAID database^50^ on 4 February and again on 1 June 2021. The metadata already contains annotations of genomes based on PANGOLIN^35^. These lineages were used to sub-divide the genomes in further analyses.

All Indonesian sequences and those transmitted from or historically linked to Indonesia (*i*.*e*., Indonesia as the country of exposure) were selected for phylogenetic analysis; only complete genomes with high coverage were chosen for phylogenetic analysis. Up to 1 June 2021, there were 1830 sequences that were aligned and the 5’ and 3’ ends trimmed using MAFFT v.7.475 with automatic flavour selection^71^. Maximum-likelihood phylogenetic trees were produced from the trimmed sequences using IQ-TREE2^72^, employing the GTR+F+R3 model of nucleotide substitution as suggested by the software’s model finder^73^, with 1000 SH-like approximate likelihood ratio test (SH-aLRT) Ultra-Fast Bootstrap^74^. The output tree was rooted using the earliest Indonesian genome, *i*.*e*., Indonesia/JI-ITD-136N/2020|EPI_ISL_529961|2020-03-12. The tree was then visualized and annotated with FigTree, where the nodes were ordered and rooted^75^.

### Data Availability

The datasets analysed during the current study are available for registered users in the GISAID repository, https://www.gisaid.org/.

## Supporting information

Acknowledgement Table

## Acknowledgements and Funding

We would like to express our gratitude to the labs and institutions involved in WGS of SARS-CoV-2 and deposited their data on GISAID. Specifically, we thank those in Indonesia, especially the Indonesian SARS-CoV-2 Genomics Surveillance Network, and the countries whose data are used for this paper. A list of contributing labs whose data were used in phylogenetic analyses is presented in Supplementary Table S3. We also convey our appreciation to those who had given their consents to their samples being used for SARS-CoV-2 research, and Josh Quick for developing and his guidance of the ARTIC Network’s protocol for the ONT platform. We thank Puspita Lisdiyanti, Director of the Research Centre for Biotechnology, LIPI, for facilitating the collaboration. We are grateful for the help of Realtime Surveillance Genome SARS-CoV-2 (VenomCoV) team in LIPI for sample collection, sample preparation, library preparation, sequencing, and bioinformatics analysis. We also thank Safarina G. Malik in Eijkman Institute for Molecular Biology and Vivi Setiawati, Director of National Institute for Health Research and Development of the Ministry of Health Republic of Indonesia, for valuable input to the manuscript.

This work was funded by the Government of Indonesia through the Indonesian Endowment Fund (LPDP-RISPRO) grant scheme, on which WK is the Principal Investigator. IC was supported by a Wellcome Trust UK grant. SS was funded by Bowel Cancer Research UK.

## Author Contributions

IC, WK and SS conceptualised and designed the study. IC, MWL, and SS performed data, analysis, and wrote the main manuscript text. ARU, MI, and WK revised the manuscript. IC prepared Figure 2, 5A, 5B, Table 3, 4. EWP and GS prepared Figure 1A, 1B, 3, 4, Table 1. SHBW prepared Figure 1C, 5C, Table 2. AMR, SHBW, HH, and GA involved in data acquisition. ISGSN submitted the Indonesian SARS-CoV-2 genomes to GISAID. All authors reviewed and approved the manuscript.

