## Supplementary Materials for "Genome Profiling of SARS-CoV-2 in Indonesia, ASEAN, and the Neighbouring East Asian Countries: Features, Challenges, and Achievements"

**Supplementary Table S1 Genomes submitted by NICCRAT-LIPI team**

| **No.** | **Virus Name** | **Accession ID** | **Submission Date** | **Collection Date** | **Patient Status** |
| --- | --- | --- | --- | --- | --- |
| 1. | hCoV-19/Indonesia/BGR-LIPI046/2020 | EPI_ISL_1020188 | 18/02/2021 | 13/01/2021 | symptomatic |
| 2. | hCoV-19/Indonesia/BGR-LIPI035/2020 | EPI_ISL_1020192 | 18/02/2021 | 13/10/2020 | asymptomatic |
| 3. | hCoV-19/Indonesia/BGR-LIPI043/2020 | EPI_ISL_1020197 | 18/02/2021 | 1/12/2020 | asymptomatic |
| 4. | hCoV-19/Indonesia/JK-LIPI032/2020 | EPI_ISL_1020203 | 18/02/2021 | 18/09/2020 | symptomatic |
| 5. | hCoV-19/Indonesia/BGR-LIPI041/2020 | EPI_ISL_1117451 | 2/03/2021 | 9/11/2020 | unknown |
| 6. | hCoV-19/Indonesia/BGR-LIPI031/2020 | EPI_ISL_1117456 | 2/03/2021 | 7/09/2020 | unknown |
| 7. | hCoV-19/Indonesia/BGR-LIPI033/2020 | EPI_ISL_1117457 | 2/03/2021 | 7/09/2020 | asymptomatic |
| 8. | hCoV-19/Indonesia/BGR-LIPI070/2020 | EPI_ISL_1117590 | 2/03/2021 | 30/11/2020 | symptomatic |
| 9. | hCoV-19/Indonesia/BGR-LIPI071/2020 | EPI_ISL_1117591 | 2/03/2021 | 1/12/2020 | symptomatic |
| 10. | hCoV-19/Indonesia/BGR-LIPI073/2020 | EPI_ISL_1117592 | 2/03/2021 | 7/12/2020 | hospitalized |
| 11. | hCoV-19/Indonesia/BGR-LIPI072/2020 | EPI_ISL_1118280 | 2/03/2021 | 6/12/2020 | symptomatic |
| 12. | hCoV-19/Indonesia/JB-BGR-LIPI075/2020 | EPI_ISL_1137601 | 4/03/2021 | 30/10/2020 | symptomatic |

**Supplementary Table S2 Age and Gender Distribution of Countries**

| **Country** | **% Ages of Total Population** | | | **% Gender of Total Population** | |
| --- | --- | --- | --- | --- | --- |
|  | **0-14 years** [1] | **15-64 years** [2] | **>65 years** [3] | **Female** [4] | **Male** [5] |
| China | 13 | 71 | 11 | 48.7 | 51 |
| Hong Kong | 12 | 70 | 17 | 54.0 | 46 |
| Indonesia | 26 | 68 | 6 | 49.6 | 50 |
| Japan | 13 | 59 | 28 | 51.2 | 49 |
| South Korea | 13 | 72 | 15 | 49.9 | 50 |
| United Kingdom | 18 | 64 | 19 | 50.6 | 49 |

**Supplementary Figure S1 and S2**

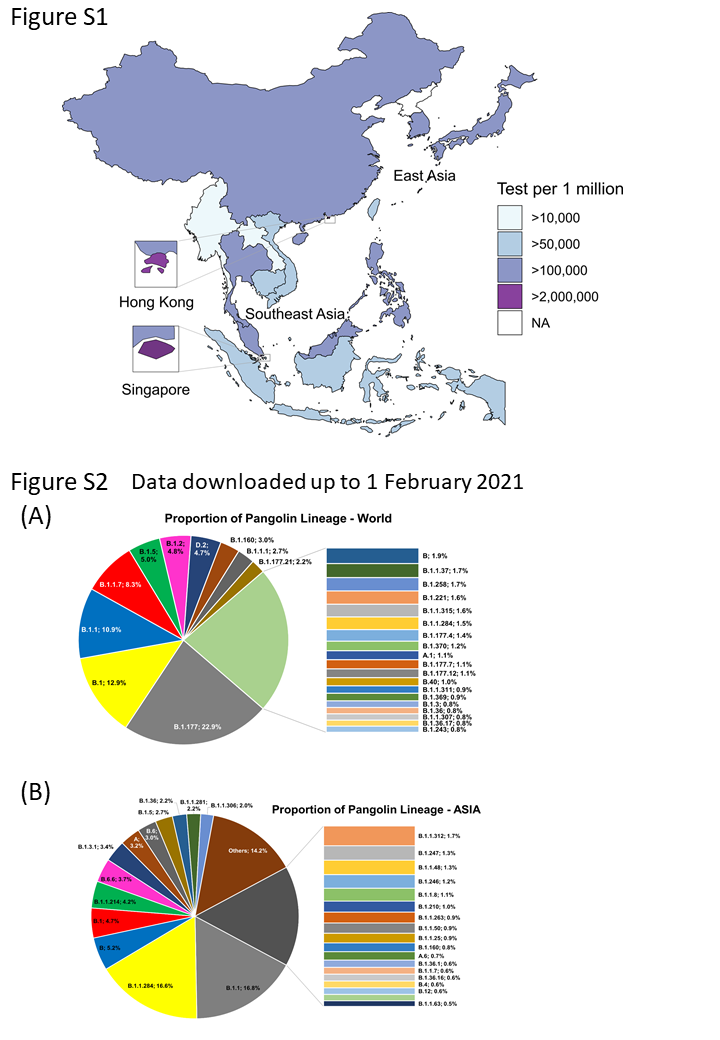

**Supplementary Figure S3**

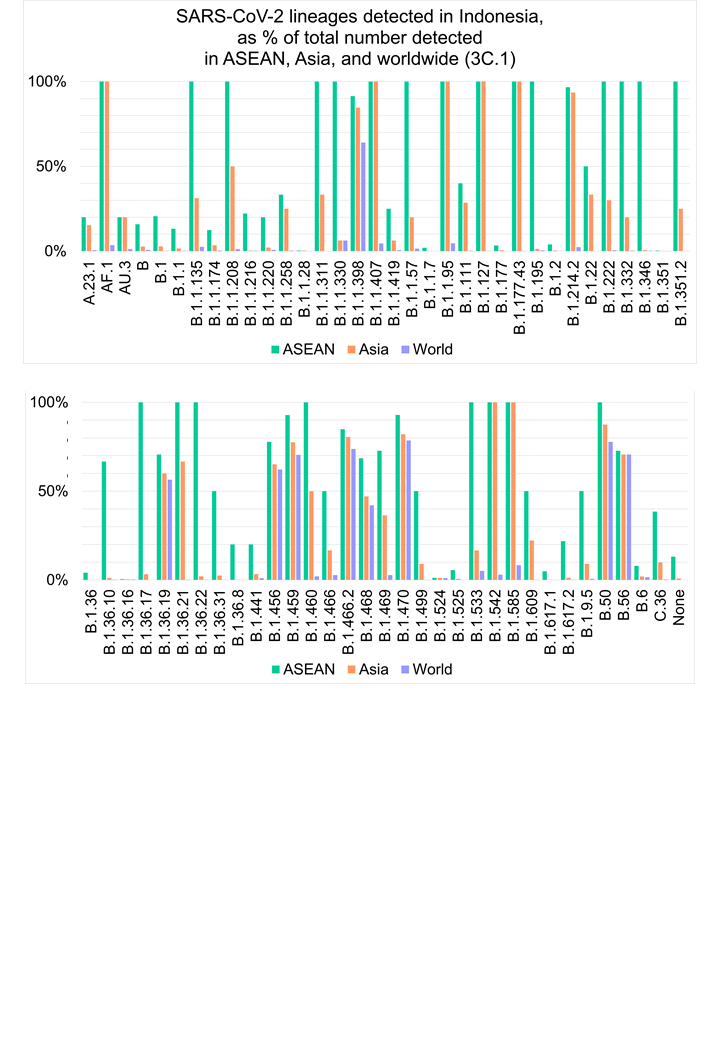

**Supplementary Figure S4**

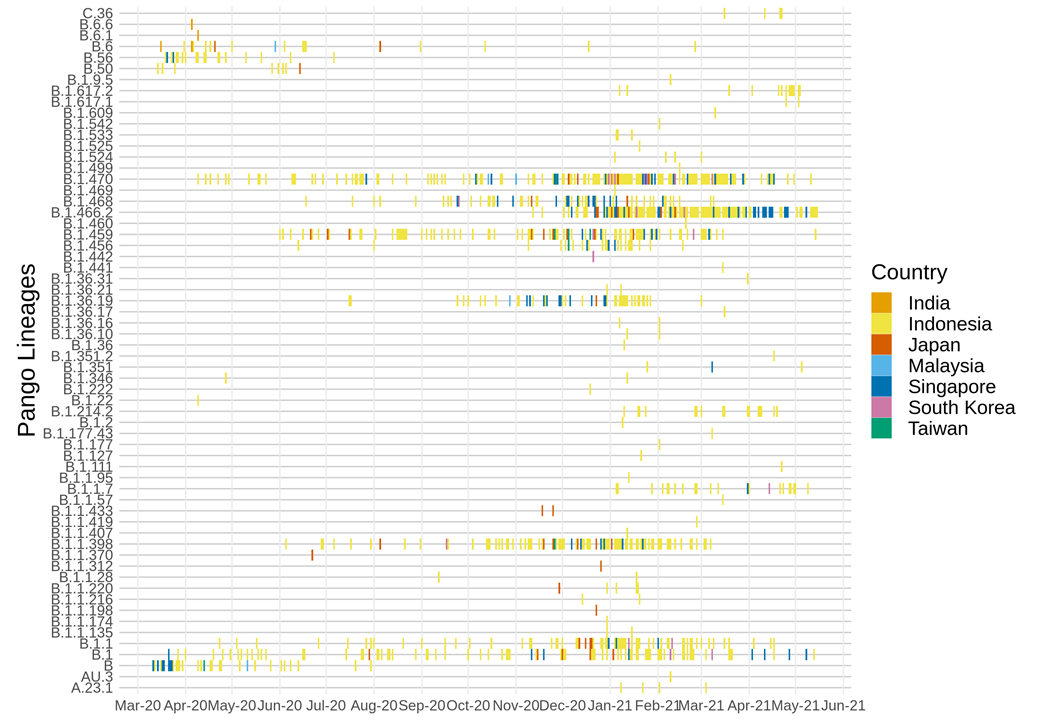

**Supplementary Figure S5**

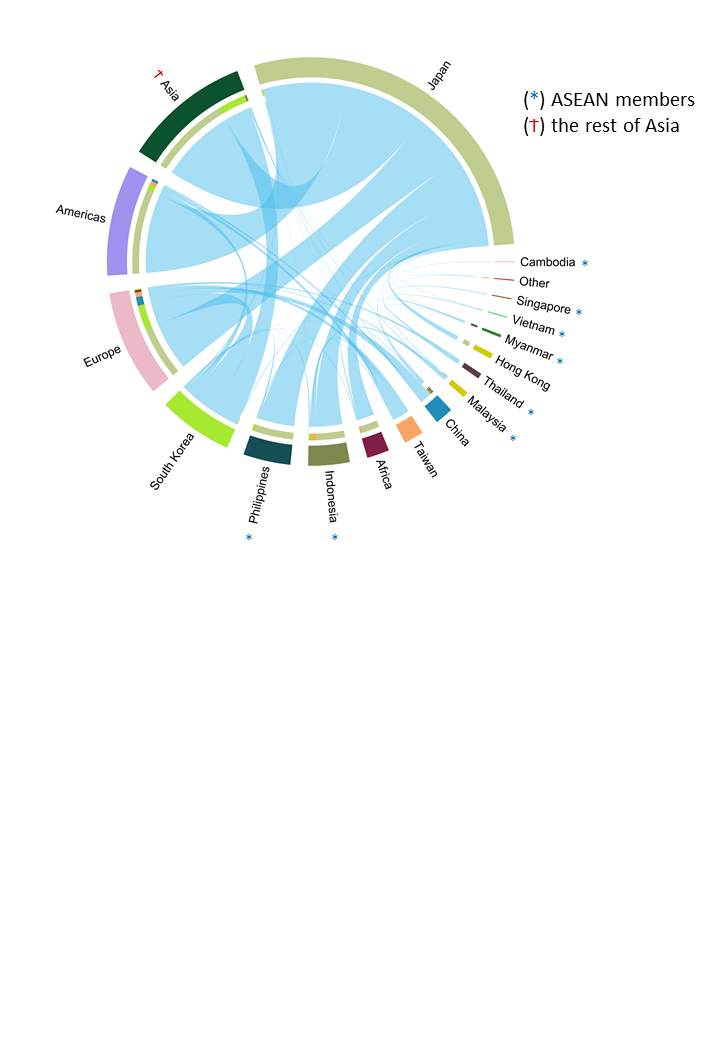

**Supplementary Figure Descriptions**

Figure S1. COVID-19 test rates of ASEAN and East Asian countries per 1 June 2021

Figure S2. Proportion of PANGO lineages based on the data downloaded from GISAID per 1 February 2021

A) Proportion of PANGO lineages in the world; the most common variant at the time was B.1.177.
B) Proportion of PANGO lineages in Asia; the two most common variants at the time were B.1.1 and B.1.1.284.

Figure S3. Distribution of variants in Indonesia as proportions to the neighbouring regions and the rest of the world; B.1.177.43 is an example of variant found only in Indonesia (upper panel).

Figure S4. Timeline of Indonesian-associated variants by PANGO lineage based on data downloaded from GISAID on 1 June 2021 and coloured by country; exported transmissions were primarily within Southeast and East Asia.

Figure S5 Circular plot shows associations of variants (*i.e.* exported/imported transmissions) between different countries in the world of data per 1 February 2021; transmission events were the highest to Japan at almost 74% of the total number of exported cases.
