## Supplementary material for "Genome Profiling of SARS-CoV-2 in Indonesia, ASEAN, and the Neighbouring East Asian Countries: Features, Challenges, and Achievements": Acknowledgement Table

We gratefully acknowledge the following Authors from the Originating laboratories responsible for obtaining the specimens, as well as the Submitting laboratories where the genome data were generated and shared via GISAID, on which this research is based.

All Submitters of data may be contacted directly via www.gisaid.org

Authors are sorted alphabetically.

| Accession ID | Originating Laboratory | Submitting Laboratory | Authors |
|---|---|---|---|
| EPI_ISL_1005695 | RSUD Dr. Soedirman | Genetics Working Group (Pokja Genetik) Faculty of Medicine, Public Health and Nursing Universitas Gadjah Mada (FK-KMK UGM); Disease Investigation Center Wates Ministry of Agriculture Indonesia; Department of Microbiology FK-KMK UGM; Laboratorium Diagnostik Yayasan Tahija World Mosquito Program (WMP) Yogyakarta Center for Tropical Medicine FK-KMK UGM; Integrated Research Center FK-KMK UGM; Department of Computer Science and Electronics FMIPA UGM; Balai Besar Teknik Kesehatan Lingkungan dan Pengendalian Penyakit (BBTKLPP) Yogyakarta | Gunadi, Hendra Wibawa, Marcellus, Mohamad S. Hakim, Edwin W. Daniwijaya, Ludhang P. Rizki, Endah Supriyati, Eggi Arguni, Titik Nuryastuti, Tri Wibawa, Dwi AA Nugrahaningsih, Afiahayati, Siswanto, Kristy Iskandar, Nungki Anggorowati, Irene, Indaryati, Havi[illegible] Sari, Audric Kenny Tedja, Maria Patricia Inggriani |
| EPI_ISL_1005696 | RSUD Dr. Tjitrowardojo | Genetics Working Group (Pokja Genetik) Faculty of Medicine, Public Health and Nursing Universitas Gadjah Mada (FK-KMK UGM); Disease Investigation Center Wates Ministry of Agriculture Indonesia; Department of Microbiology FK-KMK UGM; Laboratorium Diagnostik Yayasan Tahija World Mosquito Program (WMP) Yogyakarta Center for Tropical Medicine FK-KMK UGM; Integrated Research Center FK-KMK UGM; Department of Computer Science and Electronics FMIPA UGM; Balai Besar Teknik Kesehatan Lingkungan dan Pengendalian Penyakit (BBTKLPP) Yogyakarta | Gunadi, Hendra Wibawa, Marcellus, Mohamad S. Hakim, Edwin W. Daniwijaya, Ludhang P. Rizki, Endah Supriyati, Eggi Arguni, Titik Nuryastuti, Tri Wibawa, Dwi AA Nugrahaningsih, Afiahayati, Siswanto, Kristy Iskandar, Nungki Anggorowati, Irene, Indaryati, Havi[illegible] Sari, Susan Simanjaya, Alvin Santoso Kalim |
| EPI_ISL_1005697 | RSUD Dr. Tjitrowardojo | Genetics Working Group (Pokja Genetik) Faculty of Medicine, Public Health and Nursing Universitas Gadjah Mada (FK-KMK UGM); Disease Investigation Center Wates Ministry of Agriculture Indonesia; Department of Microbiology FK-KMK UGM; Laboratorium Diagnostik Yayasan Tahija World Mosquito Program (WMP) Yogyakarta Center for Tropical Medicine FK-KMK UGM; Integrated Research Center FK-KMK UGM; Department of Computer Science and Electronics FMIPA UGM; Balai Besar Teknik Kesehatan Lingkungan dan Pengendalian Penyakit (BBTKLPP) Yogyakarta | Gunadi, Hendra Wibawa, Marcellus, Mohamad S. Hakim, Edwin W. Daniwijaya, Ludhang P. Rizki, Endah Supriyati, Eggi Arguni, Titik Nuryastuti, Tri Wibawa, Dwi AA Nugrahaningsih, Afiahayati, Siswanto, Kristy Iskandar, Nungki Anggorowati, Irene, Indaryati, Havi[illegible] Sari, Kemala Athollah, Dyah Ayu Puspitarani |
| EPI_ISL_1005698 | RSUD Dr. | Genetics Working Group | Gunadi, Hendra Wibawa, Marcellus, Mohamad S. Hakim, Edwin W. Daniwijaya, Ludhang P. Rizki, Endah Supriyati, Eggi Arguni, Titik Nuryastuti, Tri Wibawa, Dwi AA Nugrahaningsih, Afiahayati, Siswanto, Kristy Iskandar, Nungki Anggorowati, Irene, Indaryati, Havi[illegible] |

| | | | |
|---|---|---|---|
| | Tjitrowardojo | (Pokja Genetik) Faculty of Medicine, Public Health and Nursing Universitas Gadjah Mada (FK-KMK UGM); Disease Investigation Center Wates Ministry of Agriculture Indonesia; Department of Microbiology FK-KMK UGM; Laboratorium Diagnostik Yayasan Tahija World Mosquito Program (WMP) Yogyakarta Center for Tropical Medicine FK-KMK UGM; Integrated Research Center FK-KMK UGM; Department of Computer Science and Electronics FMIPA UGM; Balai Besar Teknik Kesehatan Lingkungan dan Pengendalian Penyakit (BBTKLPP) Yogyakarta | Sari, Aditya Rifqi Fauzi, Fadil Fahri |
| EPI_ISL_1020175 | RS PMI, Bogor, West Java | Biosafety Level-3 Laboratory, Indonesian Institute of Sciences (LIPI) | Isa Nuryana, Syaiful Rizal, Indriawati, Ari S. Wulandari, Syam Budi Iryanto, Anik B. Dharmayanthi, Andri Wardiana, Sugiyono Saputra |
| EPI_ISL_1020176 | RS Trimitra, Bogor, West java | Biosafety Level-3 Laboratory, Indonesian Institute of Sciences (LIPI) | Isa Nuryana, Syaiful Rizal, Indriawati, Ari S. Wulandari, Syam Budi Iryanto, Anik B. Dharmayanthi, Andri Wardiana, Sugiyono Saputra |
| EPI_ISL_1020177 | RS Trimitra, Bogor, West java | Biosafety Level-3 Laboratory, Indonesian Institute of Sciences (LIPI) | Isa Nuryana, Syaiful Rizal, Indriawati, Ari S. Wulandari, Syam Budi Iryanto, Ratih Asmana Ningrum, Andri Wardiana, Sugiyono Saputra |
| EPI_ISL_1020178 | RS Trimitra, Bogor, West java | Biosafety Level-3 Laboratory, Indonesian Institute of Sciences (LIPI) | Isa Nuryana, Syaiful Rizal, Indriawati, Ari S. Wulandari, Syam Budi Iryanto, Asep M. Ridwanuloh, Andri Wardiana, Sugiyono Saputra |
| EPI_ISL_1020179 | Primera Clinica, Jakarta | Biosafety Level-3 Laboratory, Indonesian Institute of Sciences (LIPI) | Isa Nuryana, Syaiful Rizal, Indriawati, Ari S. Wulandari, Syam Budi Iryanto, Ratih Asmana Ningrum, Andri Wardiana, Sugiyono Saputra |
| EPI_ISL_1020180 | RS PMI, Bogor, West Java | Biosafety Level-3 Laboratory, Indonesian Institute of Sciences (LIPI) | Isa Nuryana, Syaiful Rizal, Indriawati, Ari S. Wulandari, Syam Budi Iryanto, Gita Syahputra, Andri Wardiana, Sugiyono Saputra |
| EPI_ISL_1020181 | Primera Clinica, Jakarta | Biosafety Level-3 Laboratory, Indonesian Institute of Sciences (LIPI) | Isa Nuryana, Syaiful Rizal, Indriawati, Ari S. Wulandari, Syam Budi Iryanto, Ratih Asmana Ningrum, Andri Wardiana, Sugiyono Saputra |
| EPI_ISL_1020182 | RS Trimitra, Bogor, West java | Biosafety Level-3 Laboratory, Indonesian Institute of Sciences (LIPI) | Isa Nuryana, Syaiful Rizal, Indriawati, Ari S. Wulandari, Syam Budi Iryanto, Sugiyono Saputra, Andri Wardiana |
| EPI_ISL_1020183 | RS Trimitra, Bogor, West java | Biosafety Level-3 Laboratory, Indonesian Institute of Sciences (LIPI) | Sri Swasthikawati, Linda Sukmarini, Idris, Masrukhin, Alfi T. Fathurahman, Ruby Setiawan, Ratih Asmana Ningrum, Sugiyono Saputra |
| EPI_ISL_1020184 | RS PMI, Bogor, West Java | Biosafety Level-3 Laboratory, Indonesian Institute of Sciences (LIPI) | Isa Nuryana, Syaiful Rizal, Indriawati, Ari S. Wulandari, Syam Budi Iryanto, Gita Syahputra, Andri Wardiana, Sugiyono Saputra |
| EPI_ISL_1020185 | RS Trimitra, Bogor, West java | Biosafety Level-3 Laboratory, Indonesian Institute of Sciences (LIPI) | Isa Nuryana, Syaiful Rizal, Indriawati, Ari S. Wulandari, Syam Budi Iryanto, Sugiyono Saputra, Andri Wardiana |
| EPI_ISL_1020186 | BSL3 Laboratory, Bogor, West Java | Biosafety Level-3 Laboratory, Indonesian Institute of Sciences (LIPI) | Anggia Prasetyoputri, Isa Nuryana, Sri Swasthikawati, Sugiyono Saputra, Syam Budi Iryanto, Gita Syahputra, Ratih Asmana Ningrum |
| EPI_ISL_1020187 | RS Trimitra, Bogor, West java | Biosafety Level-3 Laboratory, Indonesian Institute of Sciences (LIPI) | Anggia Prasetyoputri, Erwin Hasrianda, Herjuno A. Nugroho, Nova Dhilla Yanthi, Fahrurrozi, Syam Budi Iryanto, Ratih Asmana Ningrum |
| EPI_ISL_1020188 | RS Trimitra, Bogor, West java | Biosafety Level-3 Laboratory, Indonesian Institute of Sciences (LIPI) | Ade Andriani, Hariyatun, Asep M. Ridwanuloh, Yuliawati, Eko Wahyu Putro, Ruby Setiawan, Sugiyono Saputra, Susanti, Inswasti Cahyani, Matthew W. Loose, Mohammad Ilyas, Wien Kusharyoto |
| EPI_ISL_1020189 | RS Trimitra, Bogor, West java | Biosafety Level-3 Laboratory, Indonesian Institute of Sciences (LIPI) | Isa Nuryana, Syaiful Rizal, Indriawati, Ari S. Wulandari, Syam Budi Iryanto, Sugiyono Saputra, Andri Wardiana |
| EPI_ISL_1020190 | RS Trimitra, Bogor, West java | Biosafety Level-3 Laboratory, Indonesian Institute of Sciences (LIPI) | Isa Nuryana, Syaiful Rizal, Indriawati, Ari S. Wulandari, Syam Budi Iryanto, Asep M. Ridwanuloh, Andri Wardiana, Sugiyono Saputra |
| EPI_ISL_1020191 | Primera Clinica, Jakarta | Biosafety Level-3 Laboratory, Indonesian Institute of Sciences (LIPI) | Isa Nuryana, Syaiful Rizal, Indriawati, Ari S. Wulandari, Syam Budi Iryanto, Anik B. Dharmayanthi, Andri Wardiana, Sugiyono Saputra |
| EPI_ISL_1020192 | Melania Hospital, Bagor , west Java | Biosafety Level-3 Laboratory, Indonesian Institute of Sciences (LIPI) | Anik Budhi Dharmayanthi, Ade Andriani, Hariyatun, Asep M. Ridwanuloh, Syam Budi Iryanto, Eko Wahyu Putro, Gita Syahputra, Susanti, Inswasti Cahyani, Matthew W. Loose, Mohammad Ilyas, Wien Kushary[illegible] |
| EPI_ISL_1020193 | RS PMI, Bogor, | Biosafety Level-3 | Anggia Prasetyoputri, Isa Nuryana, Sri Swasthikawati, Sugiyono Saputra, Syam Budi Iryanto, Eko Wahyu Putro, Fahrurrozi, Ratih Asmana Ningrum |

| | | | |
|---|---|---|---|
| | West Java | Laboratory, Indonesian Institute of Sciences (LIPI) | |
| EPI_ISL_1020194 | UMMI Hospital, Bogor, West Java | Biosafety Level-3 Laboratory, Indonesian Institute of Sciences (LIPI) | Anggia Prasetyoputri, Isa Nuryana, Sri Swasthikawati, Sugiyono Saputra, Syam Budi Iryanto, Gita Syahputra, Ratih Asmana Ningrum |
| EPI_ISL_1020195 | Melania Hospital, Bogor , West Java | Biosafety Level-3 Laboratory, Indonesian Institute of Sciences (LIPI) | Anggia Prasetyoputri, Erwin Hasrianda, Herjuno A. Nugroho, Nova Dhilla Yanthi, Fahrurrozi, Syam Budi Iryanto, Gita Syahputra, Ratih Asmana Ningrum |
| EPI_ISL_1020196 | RS PMI, Bogor, West Java | Biosafety Level-3 Laboratory, Indonesian Institute of Sciences (LIPI) | Anggia Prasetyoputri, Isa Nuryana, Sri Swasthikawati, Sugiyono Saputra, Syam Budi Iryanto, Eko Wahyu Putro, Ratih Asmana Ningrum |
| EPI_ISL_1020197 | Dinkes, Bogor, West Java | Biosafety Level-3 Laboratory, Indonesian Institute of Sciences (LIPI) | Anik Budhi Dharmayanthi, Ade Andriani, Hariyatun, Asep M. Ridwanuloh, Syam Budi Iryanto, Eko Wahyu Putro, Gita Syahputra, Susanti, Inswasti Cahyani, Matthew W. Loose, Mohammad Ilyas, Wien Kushar[illegible] |
| EPI_ISL_1020198 | RS Trimitra, Bogor, West java | Biosafety Level-3 Laboratory, Indonesian Institute of Sciences (LIPI) | Sri Swasthikawati, Linda Sukmarini, Idris, Masrukhin, Alfi T. Fathurahman, Eko Wahyu Putro, Ratih Asmana Ningrum, Sugiyono Saputra |
| EPI_ISL_1020199 | Balai Besar Rehabilitasi (BNN) | Biosafety Level-3 Laboratory, Indonesian Institute of Sciences (LIPI) | Sri Swasthikawati, Linda Sukmarini, Idris, Masrukhin, Alfi T. Fathurahman, Ruby Setiawan, Ratih Asmana Ningrum, Sugiyono Saputra |
| EPI_ISL_1020200 | RS Trimitra, Bogor, West java | Biosafety Level-3 Laboratory, Indonesian Institute of Sciences (LIPI) | Sri Swasthikawati, Linda Sukmarini, Idris, Masrukhin, Alfi T. Fathurahman, Ruby Setiawan, Ratih Asmana Ningrum, Sugiyono Saputra |
| EPI_ISL_1020201 | RS Trimitra, Bogor, West java | Biosafety Level-3 Laboratory, Indonesian Institute of Sciences (LIPI) | Isa Nuryana, Syaiful Rizal, Indriawati, Ari S. Wulandari, Syam Budi Iryanto, Anik B. Dharmayanthi, Andri Wardiana, Sugiyono Saputra |
| EPI_ISL_1020202 | RS Trimitra, Bogor, West java | Biosafety Level-3 Laboratory, Indonesian Institute of Sciences (LIPI) | Isa Nuryana, Syaiful Rizal, Indriawati, Ari S. Wulandari, Syam Budi Iryanto, Ratih Asmana Ningrum, Andri Wardiana, Sugiyono Saputra |
| EPI_ISL_1020203 | Klinik Apotik Dien, Jakarta | Biosafety Level-3 Laboratory, Indonesian Institute of Sciences (LIPI) | Anik Budhi Dharmayanthi, Ade Andriani, Hariyatun, Asep M. Ridwanuloh, Syam Budi Iryanto, Eko Wahyu Putro, Gita Syahputra, Susanti, Inswasti Cahyani, Matthew W. Loose, Mohammad Ilyas, Wien Kushar[illegible] |
| EPI_ISL_1020204 | DD Klinik Ciputat, South Tangerang, Banten | Biosafety Level-3 Laboratory, Indonesian Institute of Sciences (LIPI) | Anggia Prasetyoputri, Isa Nuryana, Sri Swasthikawati, Sugiyono Saputra, Syam Budi Iryanto, Eko Wahyu Putro, Fahrurrozi, Ratih Asmana Ningrum |
| EPI_ISL_1020205 | DMC Dompet Dhuafa, South Tangerang, Banten | Biosafety Level-3 Laboratory, Indonesian Institute of Sciences (LIPI) | Anggia Prasetyoputri, Isa Nuryana, Sri Swasthikawati, Sugiyono Saputra, Syam Budi Iryanto, Anik B. Dharmayanthi, Nuruliawaty Utami, Ratih Asmana Ningrum |
| EPI_ISL_1020206 | Qadr Hospital, Tangerang, Banten | Biosafety Level-3 Laboratory, Indonesian Institute of Sciences (LIPI) | Anggia Prasetyoputri, Isa Nuryana, Sri Swasthikawati, Sugiyono Saputra, Syam Budi Iryanto, Eko Wahyu Putro, Gita Syahputra, Ari Sulistyo Wulandari, Ratih Asmana Ningrum |
| EPI_ISL_1041030 | Dr. Cipto Mangunkusumo (RSCM) Hospital | Faculty of Medicine, Universitas Indonesia | Rina La Distia Nora, Mei Riasanti, Rani Dwi Ningtias, Ari Fahrial Syam |
| EPI_ISL_1048372 | Farmalab Indoutama Jakarta | National Institute of Health Research and Development | Hana Apsari Pawestri, Subangkit, Kartika Dewi Puspa, Arie Ardiansyah Nugraha, Hartanti Dian Ikawati, Krisna Nur Andriana Pangesti, Nelly Puspandari, Vivi Setiawaty. |
| EPI_ISL_1048373 | Rumah Sakit Umum Pusat Dr. Kariadi Semarang | National Institute of Health Research and Development | Hana Apsari Pawestri, Subangkit, Kartika Dewi Puspa, Arie Ardiansyah Nugraha, Hartanti Dian Ikawati, Krisna Nur Andriana Pangesti, Nelly Puspandari, Iva Puspitasari, Vivi Setiawaty. |
| EPI_ISL_1048375, EPI_ISL_1048381 | Farmalab Indoutama Jakarta | National Institute of Health Research and Development | Hana Apsari Pawestri, Subangkit, Kartika Dewi Puspa, Arie Ardiansyah Nugraha, Hartanti Dian Ikawati, Krisna Nur Andriana Pangesti, Nelly Puspandari, Vivi Setiawaty. |
| EPI_ISL_1048406 | Balai Penelitian dan Pengembangan Biomedis Papua | National Institute of Health Research and Development | Hana Apsari Pawestri, Subangkit, Kartika Dewi Puspa, Arie Ardiansyah Nugraha, Hartanti Dian Ikawati, Krisna Nur Andriana Pangesti, Nelly Puspandari, Anton Oktavianus, Hotma Sitompul, Vivi Setiawaty. |
| EPI_ISL_1069153 | Binjai Health Office | Institute of Tropical Disease, Universitas Airlangga; Fakultas Kedokteran, Universitas Sumatra Utara | Krisnoadi Rahardjo, Aldise M Nastri, Jezzy R Dewantari, Rima R Prasetya, Inke N D Lubis, R Lia Kusumawati, Muhammad Ichwan, Franciscus Ginting, Meliani, Ramadhan Bestari, Mirzan Hasibuan, Irbah R Nainggolan, R Andika D Cahyadi, Yasuko M[illegible] |
| EPI_ISL_1072962 | Kota Nopan Primary Health Centre, Mandailing Natal District | Institute of Tropical Disease, Universitas Airlangga; Faculty of Medicine, Universitas Sumatera Utara | Aldise M Nastri, Jezzy R Dewantari, Rima R Prasetya, Krisnoadi Rahardjo, Inke N D Lubis, R Lia Kusumawati, Muhammad Ichwan, Franciscus Ginting, Meliani, Ramadhan Bestari, Mirzan Hasibuan, Irbah R Nainggolan, R Andika D Cahyadi, Yasuko M[illegible] |
| EPI_ISL_1072963 | North Sumatera Health Office | Institute of Tropical Disease, Universitas Airlangga; Faculty of Medicine, Universitas Sumatera Utara | Jezzy R Dewantari, Rima R Prasetya, Krisnoadi Rahardjo, Aldise M Nastri, Inke N D Lubis, R Lia Kusumawati, Muhammad Ichwan, Franciscus Ginting, Meliani, Ramadhan Bestari, Mirzan Hasibuan, Irbah R Nainggolan, R Andika D Cahyadi, Yasuko M[illegible] |
| EPI_ISL_1072964 | Bhayangkara Hospital Medan | Institute of Tropical Disease, Universitas Airlangga; Faculty of Medicine, Universitas Sumatera Utara | Rima R Prasetya, Krisnoadi Rahardjo, Aldise M Nastri, Jezzy R Dewantari, Inke N D Lubis, R Lia Kusumawati, Muhammad Ichwan, Franciscus Ginting, Meliani, Ramadhan Bestari, Mirzan Hasibuan, Irbah R Nainggolan, R Andika D Cahyadi, Yasuko M[illegible] |
| EPI_ISL_1072965 | Department of Microbiology, | Institute of Tropical Disease, Universitas | Kazufumi Shimizu, Krisnoadi Rahardjo, Aldise M Nastri, Jezzy R Dewantari, Rima R Prasetya, Inke N D Lubis, R Lia Kusumawati, Muhammad Ichwan, Franciscus Ginting, Meliani, Ramadhan Bestari, Mirzan Hasibuan, Irbah R Nainggolan, R Andika D [illegible] |

| | | | |
|---|---|---|---|
| | Faculty of Medicine, Universitas Sumatera Utara | Airlangga; Faculty of Medicine, Universitas Sumatera Utara | |
| EPI_ISL_1072986 | Department of Microbiology, Faculty of Medicine, Universitas Sumatera Utara | Institute of Tropical Disease, Universitas Airlangga; Faculty of Medicine, Universitas Sumatera Utara | Krisnoadi Rahardjo, Aldise M Nastri, Jezzy R Dewantari, Rima R Prasetya, Inke N D Lubis, R Lia Kusumawati, Muhammad Ichwan, Franciscus Ginting, Meliani, Ramadhan Bestari, Mirzan Hasibuan, Irbah R Nainggolan, R Andika D Cahyadi, Yasuko M |
| EPI_ISL_1117450 | Melania Hospital, Bagor , west Java | Biosafety Level-3 Laboratory, Indonesian Institute of Sciences (LIPI) | Anggia Prasetyoputri, Isa Nuryana, Sri Swasthikawati, Sugiyono Saputra, Syam Budi Iryanto, Eko Wahyu Putro, Gita Syahputra, Ari Sulistyo Wulandari, Ratih Asmana Ningrum |
| EPI_ISL_1117451 | RS PMI, Bogor, West Java | Biosafety Level-3 Laboratory, Indonesian Institute of Sciences (LIPI) | Anik Budhi Dharmayanthi, Ade Andriani, Hariyatun, Asep M. Ridwanuloh, Syam Budi Iryanto, Eko Wahyu Putro, Gita Syahputra, Susanti, Inswasti Cahyani, Matthew W. Loose, Mohammad Ilyas, Wien Kushar |
| EPI_ISL_1117452 | RS Trimitra, Bogor, West java | Biosafety Level-3 Laboratory, Indonesian Institute of Sciences (LIPI) | Anggia Prasetyoputri, Erwin Hasrianda, Herjuno A. Nugroho, Nova Dhilla Yanthi, Fahrurrozi, Syam Budi Iryanto, Ratih Asmana Ningrum |
| EPI_ISL_1117453 | RS Trimitra, Bogor, West java | Biosafety Level-3 Laboratory, Indonesian Institute of Sciences (LIPI) | Sri Swasthikawati, Linda Sukmarini, Idris, Masrukhin, Alfi T. Fathurahman, Eko Wahyu Putro, Ratih Asmana Ningrum, Sugiyono Saputra |
| EPI_ISL_1117454 | RS Trimitra, Bogor, West java | Biosafety Level-3 Laboratory, Indonesian Institute of Sciences (LIPI) | Sri Swasthikawati, Linda Sukmarini, Idris, Masrukhin, Alfi T. Fathurahman, Ruby Setiawan, Ratih Asmana Ningrum, Sugiyono Saputra |
| EPI_ISL_1117455 | RS Trimitra, Bogor, West java | Biosafety Level-3 Laboratory, Indonesian Institute of Sciences (LIPI) | Isa Nuryana, Syaiful Rizal, Indriawati, Ari S. Wulandari, Syam Budi Iryanto, Gita Syahputra, Andri Wardiana, Sugiyono Saputra |
| EPI_ISL_1117456 | Murni Teguh Hospital, Jakarta | Biosafety Level-3 Laboratory, Indonesian Institute of Sciences (LIPI) | Anik Budhi Dharmayanthi, Ade Andriani, Hariyatun, Asep M. Ridwanuloh, Syam Budi Iryanto, Eko Wahyu Putro, Gita Syahputra, Susanti, Inswasti Cahyani, Matthew W. Loose, Mohammad Ilyas, Wien Kushar |
| EPI_ISL_1117457 | Klinik Apotik Dien, Jakarta | Biosafety Level-3 Laboratory, Indonesian Institute of Sciences (LIPI) | Anik Budhi Dharmayanthi, Ade Andriani, Hariyatun, Asep M. Ridwanuloh, Syam Budi Iryanto, Eko Wahyu Putro, Gita Syahputra, Susanti, Inswasti Cahyani, Matthew W. Loose, Mohammad Ilyas, Wien Kushar |
| EPI_ISL_1117589 | RS PMI, Bogor, West Java | Biosafety Level-3 Laboratory, Indonesian Institute of Sciences (LIPI) | Anggia Prasetyoputri, Isa Nuryana, Sri Swasthikawati, Sugiyono Saputra, Syam Budi Iryanto, Gita Syahputra, Ratih Asmana Ningrum |
| EPI_ISL_1117590 | RS Trimitra, Bogor, West java | Biosafety Level-3 Laboratory, Indonesian Institute of Sciences (LIPI) | Anik Budhi Dharmayanthi, Ade Andriani, Hariyatun, Asep M. Ridwanuloh, Syam Budi Iryanto, Eko Wahyu Putro, Gita Syahputra, Susanti, Inswasti Cahyani, Matthew W. Loose, Mohammad Ilyas, Wien Kushar |
| EPI_ISL_1117591 | Dinkes, Bogor, West Java | Biosafety Level-3 Laboratory, Indonesian Institute of Sciences (LIPI) | Anik Budhi Dharmayanthi, Ade Andriani, Hariyatun, Asep M. Ridwanuloh, Syam Budi Iryanto, Eko Wahyu Putro, Gita Syahputra, Susanti, Inswasti Cahyani, Matthew W. Loose, Mohammad Ilyas, Wien Kushar |
| EPI_ISL_1117592 | Melania Hospital, Bagor , West Java | Biosafety Level-3 Laboratory, Indonesian Institute of Sciences (LIPI) | Anik Budhi Dharmayanthi, Ade Andriani, Hariyatun, Asep M. Ridwanuloh, Syam Budi Iryanto, Eko Wahyu Putro, Gita Syahputra, Susanti, Inswasti Cahyani, Matthew W. Loose, Mohammad Ilyas, Wien Kushar |
| EPI_ISL_1118206, EPI_ISL_1118207, EPI_ISL_1118208, EPI_ISL_1118209, EPI_ISL_1118210, EPI_ISL_1118211, EPI_ISL_1118212, EPI_ISL_1118213, EPI_ISL_1118214, EPI_ISL_1118215, EPI_ISL_1118216 | | | |
| see above | National Institute of Health Research and Development | National Institute of Health Research and Development | Hana Apsari Pawestri, Subangkit, Kartika Dewi Puspa, Arie Ardiansyah Nugraha, Hartanti Dian Ikawati, Krisna Nur Andriana Pangesti, Yuni Rukminiati, Ririn Ramadhany, Agustiningsih, Kindi Adam, Holy Arif Wibowo, Triyani Soekarso, Ni Ketut Susilarini, Nurika Ha Tati Febriyanti, Natalie Laurencia Kipuw, Fauzul Muna, Irene Lorinda Indalao, Nelly Puspandari, Vivi Setiawaty. |
| EPI_ISL_1118280 | Merial Health Hospital, Jakarta | Biosafety Level-3 Laboratory, Indonesian Institute of Sciences (LIPI) | Anik Budhi Dharmayanthi, Ade Andriani, Hariyatun, Asep M. Ridwanuloh, Syam Budi Iryanto, Eko Wahyu Putro, Gita Syahputra, Susanti, Inswasti Cahyani, Matthew W. Loose, Mohammad Ilyas, Wien Kushar |
| EPI_ISL_1118931, EPI_ISL_1118933 | National Institute of Health Research and Development | National Institute of Health Research and Development | Hana Apsari Pawestri, Subangkit, Kartika Dewi Puspa, Arie Ardiansyah Nugraha, Hartanti Dian Ikawati, Krisna Nur Andriana Pangesti, Yuni Rukminiati, Ririn Ramadhany, Agustiningsih, Kindi Adam, Holy Arif Wibowo, Triyani Soekarso, Ni Ketut Susilarini, Nurika Ha Tati Febriyanti, Natalie Laurencia Kipuw, Fauzul Muna, Irene Lorinda Indalao, Nelly Puspandari, Vivi Setiawaty. |
| EPI_ISL_1122422, EPI_ISL_1122423, EPI_ISL_1122424 | Clinical Microbiology Laboratory, Faculty of Medicine, Universitas Indonesia | Clinical Microbiology Laboratory, Faculty of Medicine, Universitas Indonesia | Fera Ibrahim, Augustine Natasha, Andi Yasmon, Fithriyah , Yulia Rosa Saharman, Erni J. Nelwan, Anis Karuniawati, Pratiwi Sudarmono |
| EPI_ISL_1137600 | BSL3 Laboratory, Bogor, West Java | Biosafety Level-3 Laboratory, Indonesian Institute of Sciences (LIPI) | Anggia Prasetyoputri, Isa Nuryana, Sri Swasthikawati, Sugiyono Saputra, Syam Budi Iryanto, Eko Wahyu Putro, Ratih Asmana Ningrum |
| EPI_ISL_1137601 | RS PMI, Bogor, West Java | Biosafety Level-3 Laboratory, Indonesian Institute of Sciences (LIPI) | Anik Budhi Dharmayanthi, Ade Andriani, Hariyatun, Asep M. Ridwanuloh, Syam Budi Iryanto, Eko Wahyu Putro, Gita Syahputra, Susanti, Inswasti Cahyani, Matthew W. Loose, Mohammad Ilyas, Wien Kushar |
| EPI_ISL_1137602 | RS Trimitra, Bogor, West java | Biosafety Level-3 Laboratory, Indonesian Institute of Sciences (LIPI) | Ade Andriani, Hariyatun, Asep M. Ridwanulloh, Yuliawati, Eko Wahyu Putro, Ruby Setiawan, Sugiyono Saputra |
| EPI_ISL_1137603 | RS Trimitra, Bogor, West java | Biosafety Level-3 Laboratory, Indonesian Institute of Sciences (LIPI) | Anggia Prasetyoputri, Erwin Hasrianda, Herjuno A. Nugroho, Nova Dhilla Yanthi, Fahrurrozi, Syam Budi Iryanto, Ratih Asmana Ningrum |
| EPI_ISL_1159381 | Surabaya Public Health Office | Institute of Tropical Disease, Universitas Airlangga | Aldise M Nastri, Jezzy R Dewantari, Rima R Prasetya, Krisnoadi Rahardjo, Febria Rachmanita, Gatot Soegiarto, Laksmi Wulandari, Resti Yudhawati, Yasuko Mori, Soetjipto, Kazufumi Shimizu, Maria I Lusid |
| EPI_ISL_1159382 | Adi Husada | Institute of Tropical | Maria I Lusida, Krisnoadi Rahardjo, Aldise M Nastri, Jezzy R Dewantari, Rima R Prasetya, Hermanto Wijaya, Gatot Soegiarto, Laksmi Wulandari, Resti Yudhawati, Soetjipto, Yasuko Mori, Kazufumi Shimiz |

| | | | |
|---|---|---|---|
| | Kapasari Hospital | Disease, Universitas Airlangga | |
| EPI_ISL_1159383 | Siti Khodijah Hospital | Institute of Tropical Disease, Universitas Airlangga | Aldise M Nastri, Jezzy R Dewantari, Rima R Prasetya, Krisnoadi Rahardjo, Muhammad Hamdan, Gatot Soegiarto, Laksmi Wulandari, Resti Yudhawati, Soetjipto, Yasuko Mori, Maria I Lusida, Kazufumi Shimi |
| EPI_ISL_1159384 | Institute of Tropical Disease | Institute of Tropical Disease, Universitas Airlangga | Jezzy R Dewantari, Rima R Prasetya, Krisnoadi Rahardjo, Aldise M Nastri, Gatot Soegiarto, Laksmi Wulandari, Resti Yudhawati, Soetjipto, Yasuko Mori, Maria I Lusida, Kazufumi Shimizu |
| EPI_ISL_1159385 | Graha Medika Maternity Hospital | Institute of Tropical Disease, Universitas Airlangga | Rima R Prasetya, Krisnoadi Rahardjo, Aldise M Nastri, Jezzy R Dewantari, Diah R Kusumawati, Gatot Soegiarto, Laksmi Wulandari, Resti Yudhawati, Soetjipto, Yasuko Mori, Maria I Lusida, Kazufumi Shimiz |
| EPI_ISL_1159386 | Siti Khodijah Hospital | Institute of Tropical Disease, Universitas Airlangga | Krisnoadi Rahardjo, Aldise M Nastri, Jezzy R Dewantari, Rima R Prasetya, Muhammad Hamdan, Gatot Soegiarto, Laksmi Wulandari, Resti Yudhawati, Yasuko Mori, Soetjipto, Kazufumi Shimizu, Maria I Lusi |
| EPI_ISL_1159387 | Adi Husada Kapasari Hospital | Institute of Tropical Disease, Universitas Airlangga | Maria I Lusida, Krisnoadi Rahardjo, Aldise M Nastri, Jezzy R Dewantari, Rima R Prasetya, Hermanto Wijaya, Gatot Soegiarto, Laksmi Wulandari, Resti Yudhawati, Soetjipto, Yasuko Mori, Kazufumi Shimizu |
| EPI_ISL_1159388 | Paru Hospital | Institute of Tropical Disease, Universitas Airlangga | Aldise M Nastri, Jezzy R Dewantari, Rima R Prasetya, Krisnoadi Rahardjo, Dyah Retno, Gatot Soegiarto, Laksmi Wulandari, Resti Yudhawati, Soetjipto, Yasuko Mori, Maria I Lusida, Kazufumi Shimizu |
| EPI_ISL_1159389 | Darmo Hospital | Institute of Tropical Disease, Universitas Airlangga | Jezzy R Dewantari, Rima R Prasetya, Krisnoadi Rahardjo, Aldise M Nastri, Sulung Budianto, Gatot Soegiarto, Laksmi Wulandari, Resti Yudhawati, Soetjipto, Yasuko Mori, Maria I Lusida, Kazufumi Shimizu |
| EPI_ISL_1159390 | Semen Gresik Hospital | Institute of Tropical Disease, Universitas Airlangga | Rima R Prasetya, Krisnoadi Rahardjo, Aldise M Nastri, Jezzy R Dewantari, Muchdor, Gatot Soegiarto, Laksmi Wulandari, Resti Yudhawati, Soetjipto, Yasuko Mori, Maria I Lusida, Kazufumi Shimizu |
| EPI_ISL_1159391 | Royal Hospital | Institute of Tropical Disease, Universitas Airlangga | Krisnoadi Rahardjo, Aldise M Nastri, Jezzy R Dewantari, Rima R Prasetya, Henny P Margastuti, Gatot Soegiarto, Laksmi Wulandari, Resti Yudhawati, Soetjipto, Yasuko Mori, Maria I Lusida, Kazufumi Shimiz |
| EPI_ISL_1165067 | Siti Khodijah Hospital | Institute of Tropical Disease, Universitas Airlangga | Aldise M Nastri, Jezzy R Dewantari, Rima R Prasetya, Krisnoadi Rahardjo, Muhammad Hamdan, Gatot Soegiarto, Laksmi Wulandari, Resti Yudhawati, Yasuko Mori, Soetjipto, Kazufumi Shimizu, Maria I Lusi |
| EPI_ISL_1165068 | Anwar Medika General Hospital | Institute of Tropical Disease, Universitas Airlangga | Jezzy R Dewantari, Rima R Prasetya, Krisnoadi Rahardjo, Aldise M Nastri, Nungky Taniasari ,Gatot Soegiarto, Laksmi Wulandari, Resti Yudhawati, Yasuko Mori, Soetjipto, Kazufumi Shimizu, Maria I Lusida |
| EPI_ISL_1165069, EPI_ISL_1165070 | Siti Khodijah Hospital | Institute of Tropical Disease, Universitas Airlangga | Rima R Prasetya, Krisnoadi Rahardjo, Aldise M Nastri, Jezzy R Dewantari, Muhammad Hamdan, Gatot Soegiarto, Laksmi Wulandari, Resti Yudhawati, Yasuko Mori, Soetjipto, Kazufumi Shimizu, Maria I Lusi |
| EPI_ISL_1165071 | Anwar Medika General Hospital | Institute of Tropical Disease, Universitas Airlangga | Kazufumi Shimizu, Krisnoadi Rahardjo, Aldise M Nastri, Jezzy R Dewantari, Rima R Prasetya, Nungky Taniasari, Gatot Soegiarto, Laksmi Wulandari, Resti Yudhawati, Yasuko Mori, Soetjipto, Maria I Lusida |
| EPI_ISL_1165072 | Institute of Tropical Disease | Institute of Tropical Disease, Universitas Airlangga | Kazufumi Shimizu, Krisnoadi Rahardjo, Aldise M Nastri, Jezzy R Dewantari, Rima R Prasetya, Gatot Soegiarto, Laksmi Wulandari, Resti Yudhawati, Yasuko Mori, Soetjipto, Maria I Lusida |
| EPI_ISL_1165073 | Darus Syifa' Islamic Hospital | Institute of Tropical Disease, Universitas Airlangga | Kazufumi Shimizu, Krisnoadi Rahardjo, Aldise M Nastri, Jezzy R Dewantari, Rima R Prasetya, HM Faiz, Gatot Soegiarto, Laksmi Wulandari, Resti Yudhawati, Yasuko Mori, Soetjipto, Maria I Lusida |
| EPI_ISL_1165074 | Institute of Tropical Disease | Institute of Tropical Disease, Universitas Airlangga | Krisnoadi Rahardjo, Aldise M Nastri, Jezzy R Dewantari, Rima R Prasetya, Gatot Soegiarto, Laksmi Wulandari, Resti Yudhawati, Yasuko Mori, Soetjipto, Kazufumi Shimizu, Maria I Lusida |
| EPI_ISL_1165075 | Paru Hospital | Institute of Tropical Disease, Universitas Airlangga | Jezzy R Dewantari, Rima R Prasetya, Krisnoadi Rahardjo, Aldise M Nastri, Dyah Retno, Gatot Soegiarto, Laksmi Wulandari, Resti Yudhawati, Yasuko Mori, Soetjipto, Kazufumi Shimizu, Maria I Lusida |
| EPI_ISL_1165076 | Rahman Rahim Hospital | Institute of Tropical Disease, Universitas Airlangga | Jezzy R Dewantari, Rima R Prasetya, Krisnoadi Rahardjo, Aldise M Nastri, Rendra E Febrianto, Gatot Soegiarto, Laksmi Wulandari, Resti Yudhawati, Yasuko Mori, Soetjipto, Kazufumi Shimizu, Maria I Lusid |
| EPI_ISL_1169046 | National Institute of Health Research and Development | National Institute of Health Research and Development | Hana Apsari Pawestri, Subangkit, Kartika Dewi Puspa, Arie Ardiansyah Nugraha, Hartanti Dian Ikawati, Krisna Nur Andriana Pangesti, Yuni Rukminiati, Ririn Ramadhany, Agustiningsih, Kindi Adam, Holy Arif Wibowo, Triyani Soekarso, Ni Ketut Susilarini, Nurika Har Tati Febriyanti, Natalie Laurencia Kipuw, Fauzul Muna, Irene Lorinda Indalao, Nelly Puspandari, Vivi Setiawaty. |
| EPI_ISL_1169047 | RSUP Dr. Mohammad Hoesin Palembang Sumatera Selatan | National Institute of Health Research and Development | Vivi Setiawaty, Hana Apsari Pawestri, Subangkit, Kartika Dewi Puspa, Arie Ardiansyah Nugraha, Hartanti Dian Ikawati, Nelly Puspandari |
| EPI_ISL_1169048 | BBTKLPP Banjarbaru, Kalimantan Selatan | National Institute of Health Research and Development | Vivi Setiawaty, Hana Apsari Pawestri, Subangkit, Kartika Dewi Puspa, Arie Ardiansyah Nugraha, Hartanti Dian Ikawati, Nelly Puspandari |
| EPI_ISL_1169049 | RSUP Adam Malik Medan Sumatera Utara | National Institute of Health Research and Development | Vivi Setiawaty, Hana Apsari Pawestri, Subangkit, Kartika Dewi Puspa, Arie Ardiansyah Nugraha, Hartanti Dian Ikawati, Nelly Puspandari |
| EPI_ISL_1169050 | National Institute of Health Research and Development | National Institute of Health Research and Development | Hartanti Dian Ikawati, Hana Apsari Pawestri, Subangkit, Kartika Dewi Puspa, Arie Ardiansyah Nugraha, risna Nur Andriana Pangesti, Yuni Rukminiati, Ririn Ramadhany, Agustiningsih, Kindi Adam, Holy Arif Wibowo, Triyani Soekarso, Ni Ketut Susilarini, Nurika Hari Tati Febriyanti, Natalie Laurencia Kipuw, Fauzul Muna, Irene Lorinda Indalao, Nelly Puspandari, Vivi Setiawaty. |
| EPI_ISL_1169051 | National Institute of Health Research and Development | National Institute of Health Research and Development | Kartika Dewi Puspa, Hana Apsari Pawestri, Subangkit, Arie Ardiansyah Nugraha, Hartanti Dian Ikawati, Krisna Nur Andriana Pangesti, Yuni Rukminiati, Ririn Ramadhany, Agustiningsih, Kindi Adam, Holy Arif Wibowo, Triyani Soekarso, Ni Ketut Susilarini, Nurika Har Tati Febriyanti, Natalie Laurencia Kipuw, Fauzul Muna, Irene Lorinda Indalao, Nelly Puspandari, Vivi Setiawaty. |

| | | | |
|---|---|---|---|
| EPI_ISL_1169052, EPI_ISL_1169053, EPI_ISL_1169054, EPI_ISL_1169055 | National Institute of Health Research and Development | National Institute of Health Research and Development | Subangkit, Hana Apsari Pawestri, Kartika Dewi Puspa, Arie Ardiansyah Nugraha, Hartanti Dian Ikawati, Krisna Nur Andriana Pangesti, Yuni Rukminiati, Ririn Ramadhany, Agustiningsih, Kindi Adam, Holy Arif Wibowo, Triyani Soekarso, Ni Ketut Susilarini, Nurika Ha Tati Febriyanti, Natalie Laurencia Kipuw, Fauzul Muna, Irene Lorinda Indalao, Nelly Puspandari, Vivi Setiawaty. |
| EPI_ISL_1169056 | National Institute of Health Research and Development | National Institute of Health Research and Development | Kartika Dewi Puspa, Hana Apsari Pawestri, Subangkit, Arie Ardiansyah Nugraha, Hartanti Dian Ikawati, Krisna Nur Andriana Pangesti, Yuni Rukminiati, Ririn Ramadhany, Agustiningsih, Kindi Adam, Holy Arif Wibowo, Triyani Soekarso, Ni Ketut Susilarini, Nurika Ha Tati Febriyanti, Natalie Laurencia Kipuw, Fauzul Muna, Irene Lorinda Indalao, Nelly Puspandari, Vivi Setiawaty. |
| EPI_ISL_1169057, EPI_ISL_1169058 | National Institute of Health Research and Development | National Institute of Health Research and Development | Arie Ardiansyah Nugraha, Kartika Dewi Puspa, Hana Apsari Pawestri, Subangkit, , Hartanti Dian Ikawati, Krisna Nur Andriana Pangesti, Yuni Rukminiati, Ririn Ramadhany, Agustiningsih, Kindi Adam, Holy Arif Wibowo, Triyani Soekarso, Ni Ketut Susilarini, Nurika Herna, Tati Febriyanti, Natalie Laurencia Kipuw, Fauzul Muna, Irene Lorinda Indalao, Nelly Puspandari, Vivi Setiawaty. |
| EPI_ISL_1169059, EPI_ISL_1169060 | National Institute of Health Research and Development | National Institute of Health Research and Development | Hartanti Dian Ikawati, Hana Apsari Pawestri, Subangkit, Kartika Dewi Puspa, Arie Ardiansyah Nugraha, risna Nur Andriana Pangesti, Yuni Rukminiati, Ririn Ramadhany, Agustiningsih, Kindi Adam, Holy Arif Wibowo, Triyani Soekarso, Ni Ketut Susilarini, Nurika Hari Tati Febriyanti, Natalie Laurencia Kipuw, Fauzul Muna, Irene Lorinda Indalao, Nelly Puspandari, Vivi Setiawaty. |
| EPI_ISL_1170948 | National Institute of Health Research and Development | National Institute of Health Research and Development | Ririn Ramadhany, Yuni Rukminiati, Agustiningsih, Kindi Adam, Holy Arif Wibowo, Hana Apsari Pawestri, Subangkit; Kartika Dewi Puspa, Arie Ardiansyah Nugraha, Hartanti Dian Ikawati, Krisna Nur Andriana Pangesti, Ni Ketut Susilarini, Nur Ika Hariastuti, Uly Alfi Febriyanti, Natalie Laurencia Kipuw, Fauzul Muna, Aulia Rizki, Nelly Puspandari, Vivi Setiawaty |
| EPI_ISL_1170955 | National Institute of Health Research and Development | National Institute of Health Research and Development | Yuni Rukminiati, Agustiningsih, Kindi Adam, Holy Arif Wibowo, Ririn Ramadhany, Hana Apsari Pawestri, Subangkit, Kartika Dewi Puspa, Arie Ardiansyah Nugraha, Hartanti Dian Ikawati, Krisna Nur Andriana Pangesti, Ni Ketut Susilarini, Nur Ika Hariastuti, Uly A Susanti, Herna, Tati Febriyanti, Natalie Laurencia Kipuw, Fauzul Muna, Irene Lorinda Indalao, Aulia Rizki, Nelly Puspandari, Vivi Setiawaty |
| EPI_ISL_1196279, EPI_ISL_1196280, EPI_ISL_1196281, EPI_ISL_1196303, EPI_ISL_1196304 | Universitas Indonesia Hospital | Faculty of Medicine, Universitas Indonesia | Fadilah, Rafika Indah Paramita, Linda Erlina, Khaerunissa Anbar Istiadi, Ardiana Kusumaningrum, Dimas Seto, Astuti Giantini, Doni Usman, Putra Mahanaim Tampubolon, Aulia Reski Widyaningrum, Ferdi Anda Sitepu, Irandi Putra Pratomo, Dwiana Ocviyanti, Budi |
| EPI_ISL_1196774, EPI_ISL_1196792, EPI_ISL_1196793, EPI_ISL_1196795, EPI_ISL_1196798 | Clinical Microbiology Laboratory, Faculty of Medicine, Universitas Indonesia | Faculty of Medicine, Universitas Indonesia | Fadilah, Khaerunissa Anbar Istiadi, Rafika Indah Paramita, Linda Erlina, Fera Ibrahim, Anis Karuniawati, Pratiwi Sudarmono, Beti Ernawati dewi, Andi Yasmon, Dwiana Ocviyanti, Budi Wiweko, Ari Fahrial Syam, Bad |
| EPI_ISL_1239143 | National Institute of Health Research and Development | National Institute of Health Research and Development | Agustiningsih, Kindi Adam, Holy Arif Wibowo, Ririn Ramadhany, Yuni Rukminiati, Hana A Pawestri, Subangkit, Kartika Dewi Puspa, Arie Ardiansyah Nugraha, Hartanti Dian Ikawati, Krisna Nur Andriana Pangesti, Triyani Soekarso, Ni Ketut Susilarini, Nur Ika Hariast Nike Susanti, Herna, Tati Febriyanti, Natalie Laurencia Kipuw, Fauzul Muna, Aulia Rizki, Nelly Puspandari, Vivi Setiawaty |
| EPI_ISL_1239144 | National Institute of Health Research and Development | National Institute of Health Research and Development | Kindi Adam, Holy Arif Wibowo, Ririn Ramadhany, Yuni Rukminiati, Agustiningsih, Hana A Pawestri, Subangkit, Kartika Dewi Puspa, Arie Ardiansyah Nugraha, Hartanti Dian Ikawati, Krisna Nur Andriana Pangesti, Triyani Soekarso, Ni Ketut Susilarini, Nur Ika Hariast Nike Susanti, Herna, Tati Febriyanti, Natalie Laurencia Kipuw, Fauzul Muna, Aulia Rizki, Nelly Puspandari, Vivi Setiawaty |
| EPI_ISL_1239145, EPI_ISL_1239216, EPI_ISL_1239217, EPI_ISL_1239218, EPI_ISL_1239219 | National Institute of Health Research and Development | National Institute of Health Research and Development | Hana Apsari Pawestri, Subangkit, Kartika Dewi Puspa, Arie Ardiansyah Nugraha, Hartanti Dian Ikawati, Krisna Nur Andriana Pangesti, Yuni Rukminiati, Ririn Ramadhany, Agustiningsih, Kindi Adam, Holy Arif Wibowo, Triyani Soekarso, Ni Ketut Susilarini, Nurika Ha Tati Febriyanti, Natalie Laurencia Kipuw, Fauzul Muna, Irene Lorinda Indalao, Nelly Puspandari, Vivi Setiawaty. |
| EPI_ISL_1239220 | RSUPN Dr. Cipto Mangunkusumo | National Institute of Health Research and Development | Hana Apsari Pawestri, Subangkit, Kartika Dewi Puspa, Arie Ardiansyah Nugraha, Hartanti Dian Ikawati, Krisna Nur Andriana Pangesti, Yuni Rukminiati, Ririn Ramadhany, Agustiningsih, Kindi Adam, Holy Arif Wibowo, Triyani Soekarso, Ni Ketut Susilarini, Nurika Ha Tati Febriyanti, Natalie Laurencia Kipuw, Fauzul Muna, Irene Lorinda Indalao, Nelly Puspandari, Vivi Setiawaty. |
| EPI_ISL_1239221, EPI_ISL_1239222, EPI_ISL_1239223 | National Institute of Health Research and Development | National Institute of Health Research and Development | Hana Apsari Pawestri, Subangkit, Kartika Dewi Puspa, Arie Ardiansyah Nugraha, Hartanti Dian Ikawati, Krisna Nur Andriana Pangesti, Yuni Rukminiati, Ririn Ramadhany, Agustiningsih, Kindi Adam, Holy Arif Wibowo, Triyani Soekarso, Ni Ketut Susilarini, Nurika Ha Tati Febriyanti, Natalie Laurencia Kipuw, Fauzul Muna, Irene Lorinda Indalao, Nelly Puspandari, Vivi Setiawaty. |
| EPI_ISL_1247019 | Clinical microbiology lab, RSPTN Universitas Hasanuddin | Clinical microbiology lab, RSPTN Universitas Hasanuddin | Muhammad Nasrum Massi, Rizalinda Sjahril, Gita Vita Soraya, Triwibawa, Handayani, Muhammad Yusuf Usman, Rufika Shari |
| EPI_ISL_1248934 | Clinical Microbiology Lab RSPTN Universitas Hasanuddin | Clinical Microbiology Lab RSPTN Universitas Hasanuddin | Muhammad Nasrum Massi, Rizalinda Sjahril, Gita Vita Soraya, Triwibawa, Handayani, Muhammad Yusuf Usman, Rufika Shari |
| EPI_ISL_1249610, EPI_ISL_1250222 | Clinical Microbiology Lab RSPTN Universitas Hasanuddin | Clinical Microbiology Lab RSPTN Universitas Hasanuddin | Muhammad Nasrum Massi, Rizalinda Sjahril, Gita Vita Soraya, Triwibawa, Handayani Halik, Muhammad Yusuf Usman, Rufika Shari |
| EPI_ISL_1250844 | Tanjungpura University Hospital | Tanjungpura University Hospital | Andriani; Mahyarudin; Virhan Novianry; Delima Fajar Liana; Sofi Siti Shofiyah; Puji Astuti, Muhammad Ibnu Kahtan; Eka Ardiani Putri; Helmi Sastriawan; Rini Andriani; Ambar Rialita; Hana Apsari Pawestri; Arie Ardiansyah Nugraha; Kartika Dewi P |
| EPI_ISL_1250845 | Tanjungpura University Hospital | Tanjungpura University Hospital | Mahyarudin; Andriani; Virhan Novianry; Delima Fajar Liana; Sofi Siti Shofiyah; Puji Astuti, Muhammad Ibnu Kahtan; Eka Ardiani Putri; Helmi Sastriawan; Rini Andriani; Ambar Rialita; Hana Apsari Pawestri; Arie Ardiansyah Nugraha; Kartika Dewi P |
| EPI_ISL_1250846 | Tanjungpura University Hospital | Tanjungpura University Hospital | Delima Fajar Liana; Mahyarudin; Andriani; Virhan Novianry; Sofi Siti Shofiyah; Puji Astuti, Muhammad Ibnu Kahtan; Eka Ardiani Putri; Helmi Sastriawan; Rini Andriani; Ambar Rialita; Hana Apsari Pawestri; Arie Ardiansyah Nugraha; Kartika Dewi P |
| EPI_ISL_1250851 | Tanjungpura University Hospital | Tanjungpura University Hospital | Sofi Siti Shofiyah; Delima Fajar Liana; Mahyarudin; Andriani; Virhan Novianry; Puji Astuti, Muhammad Ibnu Kahtan; Eka Ardiani Putri; Helmi Sastriawan; Rini Andriani; Ambar Rialita; Hana Apsari Pawestri; Arie Ardiansyah Nugraha; Kartika Dewi P |
| EPI_ISL_1250859 | Tanjungpura University Hospital | Tanjungpura University Hospital | Puji Astuti, Sofi Siti Shofiyah; Delima Fajar Liana; Mahyarudin; Andriani; Virhan Novianry; Muhammad Ibnu Kahtan; Eka Ardiani Putri; Helmi Sastriawan; Rini Andriani; Ambar Rialita; Hana Apsari Pawestri; Arie Ardiansyah Nugraha; Kartika Dewi P |
| EPI_ISL_1257816, EPI_ISL_1257817 | National Institute of Health Research and Development | National Institute of Health Research and Development | Subangkit, Hana Apsari Pawestri, Kartika Dewi Puspa, Arie Ardiansyah Nugraha, Hartanti Dian Ikawati, Krisna Nur Andriana Pangesti, Yuni Rukminiati, Ririn Ramadhany, Agustiningsih, Kindi Adam, Holy Arif Wibowo, Triyani Soekarso, Ni Ketut Susilarini, Nurika Ha Tati Febriyanti, Natalie Laurencia Kipuw, Fauzul Muna, Irene Lorinda Indalao, Nelly Puspandari, Vivi Setiawaty. |
| EPI_ISL_1257818, EPI_ISL_1257819, EPI_ISL_1257820, EPI_ISL_1257821, EPI_ISL_1257822 | Balai Litbangkes Papua | National Institute of Health Research and Development | Hana Apsari Pawestri, Subangkit, Kartika Dewi Puspa, Arie Ardiansyah Nugraha, Hartanti Dian Ikawati, Krisna Pangesti, Nelly Puspandari, Hotma Sitompul, Antonius Octavian, Vivi Setiawaty |
| EPI_ISL_1257823, | RSUP DR | National Institute of Health | Hana Apsari Pawestri, Subangkit, Kartika Dewi Puspa, Arie Ardiansyah Nugraha, Hartanti Dian Ikawati, Krisna Pangesti, Nelly Puspandari, Eny Rahmawati, Rogaya, Vivi Setiawaty |

| | | | |
|---|---|---|---|
| EPI_ISL_1257824, EPI_ISL_1257825, EPI_ISL_1257826 | Mohammad Hoesin | Research and Development | |
| EPI_ISL_1257827, EPI_ISL_1257828, EPI_ISL_1257829 | RSUP Dr. Sardjito | National Institute of Health Research and Development | Vivi Setiawaty, Hana Apsari Pawestri, Subangkit, Kartika Dewi Puspa, Arie Ardiansyah Nugraha, Hartanti Dian Ikawati, Nelly Puspandari, Krisna Pangesti |
| EPI_ISL_1257830, EPI_ISL_1257831 | Balai Litbangkes Tanah Bumbu | National Institute of Health Research and Development | Vivi Setiawaty, Hana Apsari Pawestri, Subangkit, Kartika Dewi Puspa, Arie Ardiansyah Nugraha, Hartanti Dian Ikawati, Nelly Puspandari, Juhairiyah, Syarif Hidayat, Krisna Pangesti |
| EPI_ISL_1257832 | BTKL-PP Ambon | National Institute of Health Research and Development | Vivi Setiawaty, Hana Apsari Pawestri, Subangkit, Kartika Dewi Puspa, Arie Ardiansyah Nugraha, Hartanti Dian Ikawati, Nelly Puspandari, Krisna Pangesti |
| EPI_ISL_1257833, EPI_ISL_1257834, EPI_ISL_1257835, EPI_ISL_1257836, EPI_ISL_1257837, EPI_ISL_1257838, EPI_ISL_1257839, EPI_ISL_1257840, EPI_ISL_1257841 | RSUP Sanglah | National Institute of Health Research and Development | Arie Ardiansyah Nugraha, Hana Apsari Pawestri, Subangkit, Kartika Dewi Puspa, Hartanti Dian Ikawati, Nelly Puspandari, Krisna Pangesti, I Nengah Tony Rustawan, I Gede Ketut Sajinadiyasa, Vivi Setiawa[illegible] |
| EPI_ISL_1257842, EPI_ISL_1257843, EPI_ISL_1257844, EPI_ISL_1257845, EPI_ISL_1257846 | RSUP Fatmawati | National Institute of Health Research and Development | Kartika Dewi Puspa, Hana Apsari Pawestri, Subangkit, Arie Ardiansyah Nugraha, Hartanti Dian Ikawati, Nelly Puspandari, Krisna Pangesti, Lidya Utami, Luci Liana, Lusiya Ningsih, Vivi Setiawaty |
| EPI_ISL_1257847, EPI_ISL_1257848, EPI_ISL_1257849, EPI_ISL_1257850 | RSPI Prof.Dr.Sulianti Saroso | National Institute of Health Research and Development | Hartanti Dian Ikawati, Hana Apsari Pawestri, Subangkit, Kartika Dewi Puspa, Arie Ardiansyah Nugraha, Krisna Pangesti, Nelly Puspandari, Vivi Setiawaty |
| EPI_ISL_1257851, EPI_ISL_1257852, EPI_ISL_1257853, EPI_ISL_1257854 | BBTKLPP Banjarbaru | National Institute of Health Research and Development | Hartanti Dian Ikawati, Hana Apsari Pawestri, Subangkit, Kartika Dewi Puspa, Arie Ardiansyah Nugraha, Krisna Pangesti, Nelly Puspandari, Dewi Hermawati, Mutia Raudah, Vivi Setiawaty |
| EPI_ISL_1257855 | BPOM Mamuju | National Institute of Health Research and Development | Hartanti Dian Ikawati, Hana Apsari Pawestri, Subangkit, Kartika Dewi Puspa, Arie Ardiansyah Nugraha, Krisna Pangesti, Nelly Puspandari, Vivi Setiawaty |
| EPI_ISL_1257856, EPI_ISL_1257857, EPI_ISL_1257858 | BBTKLPP Yogyakarta | National Institute of Health Research and Development | Kartika Dewi Puspa, Hana Apsari Pawestri, Subangkit, Arie Ardiansyah Nugraha, Hartanti Dian Ikawati, Nelly Puspandari, Krisna Pangesti, Irene, Havid Setyawan, Indaryati, Vivi Setiawaty |
| EPI_ISL_1257859 | KKP Soekarno Hatta | National Institute of Health Research and Development | Hana Apsari Pawestri, Subangkit, Kartika Dewi Puspa, Arie Ardiansyah Nugraha, Hartanti Dian Ikawati, Nelly Puspandari, Krisna Pangesti, Vivi Setiawaty |
| EPI_ISL_1257860, EPI_ISL_1257861 | Farmalab Indoutama | National Institute of Health Research and Development | Hana Apsari Pawestri, Subangkit, Kartika Dewi Puspa, Arie Ardiansyah Nugraha, Hartanti Dian Ikawati, Nelly Puspandari, Krisna Pangesti, Vivi Setiawaty |
| EPI_ISL_1257862 | Farmalab Indoutama | National Institute of Health Research and Development | Vivi Setiawaty, Hana Apsari Pawestri, Subangkit, Kartika Dewi Puspa, Arie Ardiansyah Nugraha, Hartanti Dian Ikawati, Nelly Puspandari, Krisna Pangesti |
| EPI_ISL_1257863 | RSUP Dr. Sardjito | National Institute of Health Research and Development | Kartika Dewi Puspa, Hana Apsari Pawestri, Subangkit, Arie Ardiansyah Nugraha, Hartanti Dian Ikawati, Nelly Puspandari, Krisna Pangesti, Vivi Setiawaty |
| EPI_ISL_1257864 | BTKL-PP Ambon | National Institute of Health Research and Development | Kartika Dewi Puspa, Hana Apsari Pawestri, Subangkit, Arie Ardiansyah Nugraha, Hartanti Dian Ikawati, Nelly Puspandari, Krisna Pangesti, Vivi Setiawaty |
| EPI_ISL_1257865 | RSUP Adam Malik | National Institute of Health Research and Development | Kartika Dewi Puspa, Hana Apsari Pawestri, Subangkit, Arie Ardiansyah Nugraha, Hartanti Dian Ikawati, Nelly Puspandari, Krisna Pangesti, Vivi Setiawaty |
| EPI_ISL_1257866 | RSUP Adam Malik | National Institute of Health Research and Development | Vivi Setiawaty, Hana Apsari Pawestri, Subangkit, Kartika Dewi Puspa, Arie Ardiansyah Nugraha, Hartanti Dian Ikawati, Nelly Puspandari, Maliana, Ade Rahmaini, Nelly Elfrida Samosir, Rina Yunita, Krisna Pan[illegible] |
| EPI_ISL_1258423, EPI_ISL_1258424 | Clinical Microbiology Lab RSPTN Universitas Hasanuddin | Clinical Microbiology Lab RSPTN Universitas Hasanuddin | Muhammad Nasrum Massi, Rizalinda Sjahril, Gita Vita Soraya, Triwibawa, Handayani, Muhammad Yusuf Usman, Rufika Shari |
| EPI_ISL_1258463, EPI_ISL_1258464, EPI_ISL_1258465, EPI_ISL_1263319, EPI_ISL_1263320, EPI_ISL_1263321, EPI_ISL_1263322, EPI_ISL_1263323 | Clinical Microbiology Lab RSPTN Universitas Hasanuddin | Clinical Microbiology Lab RSPTN Universitas Hasanuddin | Muhammad Nasrum Massi, Rizalinda Sjahril, Gita Vita Soraya, Triwibawa, Handayani Halik, Muhammad Yusuf Usman, Rufika Shari |
| EPI_ISL_1263459 | BTKLPP Ambon | National Institute of Health Research and Development | Vivi Setiawaty, Hana Apsari Pawestri, Subangkit, Kartika Dewi Puspa, Arie Ardiansyah Nugraha, Hartanti Dian Ikawati, Nelly Puspandari, Krisna Pangesti |
| EPI_ISL_1265168 | Rumah Sakit Umum Pusat Sanglah | National Institute of Health Research and Development | Arie Ardiansyah Nugraha, Hana Apsari Pawestri, Subangkit, Kartika Dewi Puspa, Hartanti Dian Ikawati, Nelly Puspandari, Krisna Pangesti, I Nengah Tony Rustawan, I Gede Ketut Sajinadiyasa, Vivi Setiawa[illegible] |
| EPI_ISL_1265171 | Rumah Sakit Umum Pusat Fatmawati | National Institute of Health Research and Development | Kartika Dewi Puspa, Hana Apsari Pawestri, Subangkit, Arie Ardiansyah Nugraha, Hartanti Dian Ikawati, Nelly Puspandari, Krisna Pangesti, Lidya Utami, Luci Liana, Lusiya Ningsih, Vivi Setiawaty |
| EPI_ISL_1265173 | Siloam Hospitals Yogyakarta | National Institute of Health Research and Development | Hartanti Dian Ikawati, Hana Apsari Pawestri, Subangkit, Kartika Dewi Puspa, Arie Ardiansyah Nugraha, Krisna Pangesti, Nelly Puspandari, Vivi Setiawaty |
| EPI_ISL_1265273 | Rumah Sakit Umum Pusat H. Adam Malik | National Institute of Health Research and Development | Vivi Setiawaty, Hana Apsari Pawestri, Subangkit, Kartika Dewi Puspa, Arie Ardiansyah Nugraha, Hartanti Dian Ikawati, Nelly Puspandari, Maliana, Ade Rahmaini, Nelly Elfrida Samosir, Rina Yunita, Krisna Pan[illegible] |
| EPI_ISL_1265441, | BTKLPP Kelas I | Eijkman Institute for | Edison Johar, Hidayat Trimarsanto, Frilasita A Yudhaputri, Iskandar Adnan, Willy Agustine, Sukma Oktavianthi, Lydia V. Panggalo, Slamet, Vivi Setiawaty, Hana Apsari Pawestri, Safarina G Malik, Khin Saw Myint, Ami[illegible] |

| | | | |
|---|---|---|---|
| EPI_ISL_1265442, EPI_ISL_1265443 | Makassar | Molecular Biology, Ministry of Research and Technology/National Agency for Research and Innovation; National Institute of Health Research and Development | |
| EPI_ISL_1265444, EPI_ISL_1265445, EPI_ISL_1265446 | BTKLPP Kelas I Makassar | Eijkman Institute for Molecular Biology, Ministry of Research and Technology/National Agency for Research and Innovation; National Institute of Health Research and Development | Hidayat Trimarsanto, Frilasita A Yudhaputri, Iskandar Adnan, Willy Agustine, Sukma Oktavianthi, Lydia V. Panggalo, Edison Johar, Slamet, Vivi Setiawaty, Hana Apsari Pawestri, Safarina G Malik, Khin Saw Myint, Amir |
| EPI_ISL_1265447, EPI_ISL_1265448, EPI_ISL_1265449 | BTKLPP Kelas I Makassar | Eijkman Institute for Molecular Biology, Ministry of Research and Technology/National Agency for Research and Innovation; National Institute of Health Research and Development | Frilasita A Yudhaputri, Iskandar Adnan, Willy Agustine, Sukma Oktavianthi, Lydia V. Panggalo, Edison Johar, Hidayat Trimarsanto, Slamet, Vivi Setiawaty, Hana Apsari Pawestri, Safarina G Malik, Khin Saw Myint, Amir |
| EPI_ISL_1265450, EPI_ISL_1265451, EPI_ISL_1265452 | BTKLPP Kelas I Makassar | Eijkman Institute for Molecular Biology, Ministry of Research and Technology/National Agency for Research and Innovation; National Institute of Health Research and Development | Iskandar Adnan, Willy Agustine, Sukma Oktavianthi, Lydia V. Panggalo, Edison Johar, Hidayat Trimarsanto, Frilasita A Yudhaputri, Slamet, Vivi Setiawaty, Hana Apsari Pawestri, Safarina G Malik, Khin Saw Myint, Amir |
| EPI_ISL_1265453, EPI_ISL_1265454, EPI_ISL_1265455 | BTKLPP Kelas I Makassar | Eijkman Institute for Molecular Biology, Ministry of Research and Technology/National Agency for Research and Innovation; National Institute of Health Research and Development | Willy Agustine, Sukma Oktavianthi, Lydia V. Panggalo, Edison Johar, Hidayat Trimarsanto, Frilasita A Yudhaputri, Iskandar Adnan, Slamet, Vivi Setiawaty, Hana Apsari Pawestri, Safarina G Malik, Khin Saw Myint, Amir |
| EPI_ISL_1265456, EPI_ISL_1265457, EPI_ISL_1265458, EPI_ISL_1265459 | BTKLPP Kelas I Makassar | Eijkman Institute for Molecular Biology, Ministry of Research and Technology/National Agency for Research and Innovation; National Institute of Health Research and Development | Sukma Oktavianthi, Lydia V. Panggalo, Edison Johar, Hidayat Trimarsanto, Frilasita A Yudhaputri, Iskandar Adnan, Willy Agustine, Slamet, Vivi Setiawaty, Hana Apsari Pawestri, Safarina G Malik, Khin Saw Myint, Amir |
| EPI_ISL_1265460, EPI_ISL_1265461, EPI_ISL_1265462 | BTKLPP Kelas I Makassar | Eijkman Institute for Molecular Biology, Ministry of Research and Technology/National Agency for Research and Innovation; National Institute of Health Research and Development | Lydia V. Panggalo, Edison Johar, Hidayat Trimarsanto, Frilasita A Yudhaputri, Iskandar Adnan, Willy Agustine, Sukma Oktavianthi, Slamet, Vivi Setiawaty, Hana Apsari Pawestri, Safarina G Malik, Khin Saw Myint, Amir |
| EPI_ISL_1283923, EPI_ISL_1283924 | RSIA Sayyidah | Eijkman Institute for Molecular Biology, Ministry of Research and Technology/National Agency for Research and Innovation | Edison Johar, Frilasita A Yudhaputri, Hidayat Trimarsanto, Iskandar Adnan, Lydia V. Panggalo, Sukma Oktavianthi, Willy Agustine, Safarina G Malik, Khin Saw Myint, Amin Soebandrio |
| EPI_ISL_1283927 | RSU Harapan Bunda | Eijkman Institute for Molecular Biology, Ministry of Research and Technology/National Agency for Research and Innovation | Edison Johar, Frilasita A Yudhaputri, Hidayat Trimarsanto, Iskandar Adnan, Lydia V. Panggalo, Sukma Oktavianthi, Willy Agustine, Safarina G Malik, Khin Saw Myint, Amin Soebandrio |
| EPI_ISL_1283929 | RS Mitra Keluarga Kelapa Gading | Eijkman Institute for Molecular Biology, Ministry of Research and Technology/National Agency for Research and Innovation | Edison Johar, Frilasita A Yudhaputri, Hidayat Trimarsanto, Iskandar Adnan, Lydia V. Panggalo, Sukma Oktavianthi, Willy Agustine, Safarina G Malik, Khin Saw Myint, Amin Soebandrio |
| EPI_ISL_1283932 | RS Mulya Tangerang | Eijkman Institute for Molecular Biology, Ministry of Research and Technology/National Agency for Research and Innovation | Edison Johar, Frilasita A Yudhaputri, Hidayat Trimarsanto, Iskandar Adnan, Lydia V. Panggalo, Sukma Oktavianthi, Willy Agustine, Safarina G Malik, Khin Saw Myint, Amin Soebandrio |
| EPI_ISL_1283933 | RS Mitra Keluarga Gading Serpong | Eijkman Institute for Molecular Biology, Ministry of Research and | Edison Johar, Frilasita A Yudhaputri, Hidayat Trimarsanto, Iskandar Adnan, Lydia V. Panggalo, Sukma Oktavianthi, Willy Agustine, Safarina G Malik, Khin Saw Myint, Amin Soebandrio |

| | | | |
|---|---|---|---|
| | | Technology/National Agency for Research and Innovation | |
| EPI_ISL_1283936 | RS Sentra Medika Cikarang | Eijkman Institute for Molecular Biology, Ministry of Research and Technology/National Agency for Research and Innovation | Edison Johar, Frilasita A Yudhaputri, Hidayat Trimarsanto, Iskandar Adnan, Lydia V. Panggalo, Sukma Oktavianthi, Willy Agustine, Safarina G Malik, Khin Saw Myint, Amin Soebandrio |
| EPI_ISL_1283938 | RSUD Cileungsi | Eijkman Institute for Molecular Biology, Ministry of Research and Technology/National Agency for Research and Innovation | Frilasita A Yudhaputri, Hidayat Trimarsanto, Iskandar Adnan, Lydia V. Panggalo, Sukma Oktavianthi, Willy Agustine, Edison Johar, Safarina G Malik, Khin Saw Myint, Amin Soebandrio |
| EPI_ISL_1283941, EPI_ISL_1283945, EPI_ISL_1283949, EPI_ISL_1283953 | RS Mitra Keluarga Gading Serpong | Eijkman Institute for Molecular Biology, Ministry of Research and Technology/National Agency for Research and Innovation | Frilasita A Yudhaputri, Hidayat Trimarsanto, Iskandar Adnan, Lydia V. Panggalo, Sukma Oktavianthi, Willy Agustine, Edison Johar, Safarina G Malik, Khin Saw Myint, Amin Soebandrio |
| EPI_ISL_1283957 | RSIA Selasih Medika | Eijkman Institute for Molecular Biology, Ministry of Research and Technology/National Agency for Research and Innovation | Frilasita A Yudhaputri, Hidayat Trimarsanto, Iskandar Adnan, Lydia V. Panggalo, Sukma Oktavianthi, Willy Agustine, Edison Johar, Safarina G Malik, Khin Saw Myint, Amin Soebandrio |
| EPI_ISL_1283960 | RS Asy-Syifaa Leuwiliang | Eijkman Institute for Molecular Biology, Ministry of Research and Technology/National Agency for Research and Innovation | Hidayat Trimarsanto, Iskandar Adnan, Lydia V. Panggalo, Sukma Oktavianthi, Willy Agustine, Edison Johar, Frilasita A Yudhaputri, Safarina G Malik, Khin Saw Myint, Amin Soebandrio |
| EPI_ISL_1283964 | RS Mitra Keluarga Gading Serpong | Eijkman Institute for Molecular Biology, Ministry of Research and Technology/National Agency for Research and Innovation | Hidayat Trimarsanto, Iskandar Adnan, Lydia V. Panggalo, Sukma Oktavianthi, Willy Agustine, Edison Johar, Frilasita A Yudhaputri, Safarina G Malik, Khin Saw Myint, Amin Soebandrio |
| EPI_ISL_1283967 | RS Bina Husada Bogor | Eijkman Institute for Molecular Biology, Ministry of Research and Technology/National Agency for Research and Innovation | Hidayat Trimarsanto, Iskandar Adnan, Lydia V. Panggalo, Sukma Oktavianthi, Willy Agustine, Edison Johar, Frilasita A Yudhaputri, Safarina G Malik, Khin Saw Myint, Amin Soebandrio |
| EPI_ISL_1283970 | RSIA Selasih Medika | Eijkman Institute for Molecular Biology, Ministry of Research and Technology/National Agency for Research and Innovation | Hidayat Trimarsanto, Iskandar Adnan, Lydia V. Panggalo, Sukma Oktavianthi, Willy Agustine, Edison Johar, Frilasita A Yudhaputri, Safarina G Malik, Khin Saw Myint, Amin Soebandrio |
| EPI_ISL_1283973 | RSU Bunda Mulia Cikarang | Eijkman Institute for Molecular Biology, Ministry of Research and Technology/National Agency for Research and Innovation | Hidayat Trimarsanto, Iskandar Adnan, Lydia V. Panggalo, Sukma Oktavianthi, Willy Agustine, Edison Johar, Frilasita A Yudhaputri, Safarina G Malik, Khin Saw Myint, Amin Soebandrio |
| EPI_ISL_1283979 | RS Cendana | Eijkman Institute for Molecular Biology, Ministry of Research and Technology/National Agency for Research and Innovation | Iskandar Adnan, Lydia V. Panggalo, Sukma Oktavianthi, Willy Agustine, Edison Johar, Hidayat Trimarsanto, Frilasita A Yudhaputri, Safarina G Malik, Khin Saw Myint, Amin Soebandrio |
| EPI_ISL_1283982, EPI_ISL_1283984 | RS Mitra Keluarga Gading Serpong | Eijkman Institute for Molecular Biology, Ministry of Research and Technology/National Agency for Research and Innovation | Iskandar Adnan, Lydia V. Panggalo, Sukma Oktavianthi, Willy Agustine, Edison Johar, Hidayat Trimarsanto, Frilasita A Yudhaputri, Safarina G Malik, Khin Saw Myint, Amin Soebandrio |
| EPI_ISL_1283987 | RS Islam Sari Asih Ar-Rahmah | Eijkman Institute for Molecular Biology, Ministry of Research and Technology/National Agency for Research and Innovation | Iskandar Adnan, Lydia V. Panggalo, Sukma Oktavianthi, Willy Agustine, Edison Johar, Hidayat Trimarsanto, Frilasita A Yudhaputri, Safarina G Malik, Khin Saw Myint, Amin Soebandrio |
| EPI_ISL_1283988 | RSUD Cileungsi | Eijkman Institute for Molecular Biology, Ministry of Research and Technology/National Agency for Research and Innovation | Iskandar Adnan, Lydia V. Panggalo, Sukma Oktavianthi, Willy Agustine, Edison Johar, Hidayat Trimarsanto, Frilasita A Yudhaputri, Safarina G Malik, Khin Saw Myint, Amin Soebandrio |

| | | | |
|---|---|---|---|
| EPI_ISL_1283991 | RS Bhakti Mulia | Eijkman Institute for Molecular Biology, Ministry of Research and Technology/National Agency for Research and Innovation | Iskandar Adnan, Lydia V. Panggalo, Sukma Oktavianthi, Willy Agustine, Edison Johar, Hidayat Trimarsanto, Frilasita A Yudhaputri, Safarina G Malik, Khin Saw Myint, Amin Soebandrio |
| EPI_ISL_1283994 | RS Mitra Keluarga Kelapa Gading | Eijkman Institute for Molecular Biology, Ministry of Research and Technology/National Agency for Research and Innovation | Lydia V. Panggalo, Sukma Oktavianthi, Willy Agustine, Edison Johar, Hidayat Trimarsanto, Iskandar Adnan, Frilasita A Yudhaputri, Safarina G Malik, Khin Saw Myint, Amin Soebandrio |
| EPI_ISL_1283996 | RSIA Bunda Sejahtera | Eijkman Institute for Molecular Biology, Ministry of Research and Technology/National Agency for Research and Innovation | Lydia V. Panggalo, Sukma Oktavianthi, Willy Agustine, Edison Johar, Hidayat Trimarsanto, Iskandar Adnan, Frilasita A Yudhaputri, Safarina G Malik, Khin Saw Myint, Amin Soebandrio |
| EPI_ISL_1283998 | RS Mitra Keluarga Kalideres | Eijkman Institute for Molecular Biology, Ministry of Research and Technology/National Agency for Research and Innovation | Lydia V. Panggalo, Sukma Oktavianthi, Willy Agustine, Edison Johar, Hidayat Trimarsanto, Iskandar Adnan, Frilasita A Yudhaputri, Safarina G Malik, Khin Saw Myint, Amin Soebandrio |
| EPI_ISL_1284000 | RSU HGA Depok | Eijkman Institute for Molecular Biology, Ministry of Research and Technology/National Agency for Research and Innovation | Lydia V. Panggalo, Sukma Oktavianthi, Willy Agustine, Edison Johar, Hidayat Trimarsanto, Iskandar Adnan, Frilasita A Yudhaputri, Safarina G Malik, Khin Saw Myint, Amin Soebandrio |
| EPI_ISL_1284002 | RS Omni Pulomas | Eijkman Institute for Molecular Biology, Ministry of Research and Technology/National Agency for Research and Innovation | Lydia V. Panggalo, Sukma Oktavianthi, Willy Agustine, Edison Johar, Hidayat Trimarsanto, Iskandar Adnan, Frilasita A Yudhaputri, Safarina G Malik, Khin Saw Myint, Amin Soebandrio |
| EPI_ISL_1284004 | RSU Sumber Waras | Eijkman Institute for Molecular Biology, Ministry of Research and Technology/National Agency for Research and Innovation | Lydia V. Panggalo, Sukma Oktavianthi, Willy Agustine, Edison Johar, Hidayat Trimarsanto, Iskandar Adnan, Frilasita A Yudhaputri, Safarina G Malik, Khin Saw Myint, Amin Soebandrio |
| EPI_ISL_1284007, EPI_ISL_1284008 | RS Mitra Keluarga Gading Serpong | Eijkman Institute for Molecular Biology, Ministry of Research and Technology/National Agency for Research and Innovation | Sukma Oktavianthi, Willy Agustine, Edison Johar, Hidayat Trimarsanto, Iskandar Adnan, Lydia V. Panggalo, Frilasita A Yudhaputri, Safarina G Malik, Khin Saw Myint, Amin Soebandrio |
| EPI_ISL_1284014 | RSUD Cileungsi | Eijkman Institute for Molecular Biology, Ministry of Research and Technology/National Agency for Research and Innovation | Sukma Oktavianthi, Willy Agustine, Edison Johar, Hidayat Trimarsanto, Iskandar Adnan, Lydia V. Panggalo, Frilasita A Yudhaputri, Safarina G Malik, Khin Saw Myint, Amin Soebandrio |
| EPI_ISL_1284017 | RS Mitra Keluarga Kalideres | Eijkman Institute for Molecular Biology, Ministry of Research and Technology/National Agency for Research and Innovation | Sukma Oktavianthi, Willy Agustine, Edison Johar, Hidayat Trimarsanto, Iskandar Adnan, Lydia V. Panggalo, Frilasita A Yudhaputri, Safarina G Malik, Khin Saw Myint, Amin Soebandrio |
| EPI_ISL_1284021, EPI_ISL_1284025 | RS Mitra Keluarga Gading Serpong | Eijkman Institute for Molecular Biology, Ministry of Research and Technology/National Agency for Research and Innovation | Sukma Oktavianthi, Willy Agustine, Edison Johar, Hidayat Trimarsanto, Iskandar Adnan, Lydia V. Panggalo, Frilasita A Yudhaputri, Safarina G Malik, Khin Saw Myint, Amin Soebandrio |
| EPI_ISL_1284027, EPI_ISL_1284029 | RS Sentra Medika Cikarang | Eijkman Institute for Molecular Biology, Ministry of Research and Technology/National Agency for Research and Innovation | Willy Agustine, Edison Johar, Hidayat Trimarsanto, Iskandar Adnan, Lydia V. Panggalo, Sukma Oktavianthi, Frilasita A Yudhaputri, Safarina G Malik, Khin Saw Myint, Amin Soebandrio |
| EPI_ISL_1284031 | RS OMNI Alam Sutera | Eijkman Institute for Molecular Biology, Ministry of Research and Technology/National Agency for Research and Innovation | Willy Agustine, Edison Johar, Hidayat Trimarsanto, Iskandar Adnan, Lydia V. Panggalo, Sukma Oktavianthi, Frilasita A Yudhaputri, Safarina G Malik, Khin Saw Myint, Amin Soebandrio |
| EPI_ISL_1284034, EPI_ISL_1284035 | RS Taman Harapan Baru | Eijkman Institute for Molecular Biology, Ministry of Research and | Willy Agustine, Edison Johar, Hidayat Trimarsanto, Iskandar Adnan, Lydia V. Panggalo, Sukma Oktavianthi, Frilasita A Yudhaputri, Safarina G Malik, Khin Saw Myint, Amin Soebandrio |

| | | | |
|---|---|---|---|
| | | Technology/National Agency for Research and Innovation | |
| EPI_ISL_1284040 | RSU Azra | Eijkman Institute for Molecular Biology, Ministry of Research and Technology/National Agency for Research and Innovation | Willy Agustine, Edison Johar, Hidayat Trimarsanto, Iskandar Adnan, Lydia V. Panggalo, Sukma Oktavianthi, Frilasita A Yudhaputri, Safarina G Malik, Khin Saw Myint, Amin Soebandrio |
| EPI_ISL_1284044, EPI_ISL_1284048 | RS Bhakti Husada Cikarang | Eijkman Institute for Molecular Biology, Ministry of Research and Technology/National Agency for Research and Innovation | Edison Johar, Frilasita A Yudhaputri, Hidayat Trimarsanto, Iskandar Adnan, Lydia V. Panggalo, Sukma Oktavianthi, Willy Agustine, Safarina G Malik, Khin Saw Myint, Amin Soebandrio |
| EPI_ISL_1284049 | RSU Harapan Bunda | Eijkman Institute for Molecular Biology, Ministry of Research and Technology/National Agency for Research and Innovation | Edison Johar, Frilasita A Yudhaputri, Hidayat Trimarsanto, Iskandar Adnan, Lydia V. Panggalo, Sukma Oktavianthi, Willy Agustine, Safarina G Malik, Khin Saw Myint, Amin Soebandrio |
| EPI_ISL_1284052 | RS Rumah Indonesia Sehat | Eijkman Institute for Molecular Biology, Ministry of Research and Technology/National Agency for Research and Innovation | Edison Johar, Frilasita A Yudhaputri, Hidayat Trimarsanto, Iskandar Adnan, Lydia V. Panggalo, Sukma Oktavianthi, Willy Agustine, Safarina G Malik, Khin Saw Myint, Amin Soebandrio |
| EPI_ISL_1284054 | Metro Hospitals Cikarang | Eijkman Institute for Molecular Biology, Ministry of Research and Technology/National Agency for Research and Innovation | Edison Johar, Frilasita A Yudhaputri, Hidayat Trimarsanto, Iskandar Adnan, Lydia V. Panggalo, Sukma Oktavianthi, Willy Agustine, Safarina G Malik, Khin Saw Myint, Amin Soebandrio |
| EPI_ISL_1284057 | RS Primaya Tangerang | Eijkman Institute for Molecular Biology, Ministry of Research and Technology/National Agency for Research and Innovation | Edison Johar, Frilasita A Yudhaputri, Hidayat Trimarsanto, Iskandar Adnan, Lydia V. Panggalo, Sukma Oktavianthi, Willy Agustine, Safarina G Malik, Khin Saw Myint, Amin Soebandrio |
| EPI_ISL_1284058 | RS Cendana | Eijkman Institute for Molecular Biology, Ministry of Research and Technology/National Agency for Research and Innovation | Edison Johar, Frilasita A Yudhaputri, Hidayat Trimarsanto, Iskandar Adnan, Lydia V. Panggalo, Sukma Oktavianthi, Willy Agustine, Safarina G Malik, Khin Saw Myint, Amin Soebandrio |
| EPI_ISL_1284061 | RSUD Cileungsi | Eijkman Institute for Molecular Biology, Ministry of Research and Technology/National Agency for Research and Innovation | Frilasita A Yudhaputri, Hidayat Trimarsanto, Iskandar Adnan, Lydia V. Panggalo, Sukma Oktavianthi, Willy Agustine, Edison Johar, Safarina G Malik, Khin Saw Myint, Amin Soebandrio |
| EPI_ISL_1284063 | RS Mitra Keluarga Gading Serpong | Eijkman Institute for Molecular Biology, Ministry of Research and Technology/National Agency for Research and Innovation | Frilasita A Yudhaputri, Hidayat Trimarsanto, Iskandar Adnan, Lydia V. Panggalo, Sukma Oktavianthi, Willy Agustine, Edison Johar, Safarina G Malik, Khin Saw Myint, Amin Soebandrio |
| EPI_ISL_1284064 | RSU HGA Depok | Eijkman Institute for Molecular Biology, Ministry of Research and Technology/National Agency for Research and Innovation | Frilasita A Yudhaputri, Hidayat Trimarsanto, Iskandar Adnan, Lydia V. Panggalo, Sukma Oktavianthi, Willy Agustine, Edison Johar, Safarina G Malik, Khin Saw Myint, Amin Soebandrio |
| EPI_ISL_1284067 | RS Mitra Keluarga Pratama Jatiasih | Eijkman Institute for Molecular Biology, Ministry of Research and Technology/National Agency for Research and Innovation | Frilasita A Yudhaputri, Hidayat Trimarsanto, Iskandar Adnan, Lydia V. Panggalo, Sukma Oktavianthi, Willy Agustine, Edison Johar, Safarina G Malik, Khin Saw Myint, Amin Soebandrio |
| EPI_ISL_1284069, EPI_ISL_1284072 | RS Bina Sehat Mandiri | Eijkman Institute for Molecular Biology, Ministry of Research and Technology/National Agency for Research and Innovation | Frilasita A Yudhaputri, Hidayat Trimarsanto, Iskandar Adnan, Lydia V. Panggalo, Sukma Oktavianthi, Willy Agustine, Edison Johar, Safarina G Malik, Khin Saw Myint, Amin Soebandrio |
| EPI_ISL_1284074, EPI_ISL_1284076 | RSIA PKU Muhammadiyah | Eijkman Institute for Molecular Biology, Ministry of Research and Technology/National Agency for Research and Innovation | Hidayat Trimarsanto, Iskandar Adnan, Lydia V. Panggalo, Sukma Oktavianthi, Willy Agustine, Edison Johar, Frilasita A Yudhaputri, Safarina G Malik, Khin Saw Myint, Amin Soebandrio |

| | | | |
|---|---|---|---|
| EPI_ISL_1284078 | RS Rumah Indonesia Sehat | Eijkman Institute for Molecular Biology, Ministry of Research and Technology/National Agency for Research and Innovation | Hidayat Trimarsanto, Iskandar Adnan, Lydia V. Panggalo, Sukma Oktavianthi, Willy Agustine, Edison Johar, Frilasita A Yudhaputri, Safarina G Malik, Khin Saw Myint, Amin Soebandrio |
| EPI_ISL_1284083 | RS Umum Harapan Bunda | Eijkman Institute for Molecular Biology, Ministry of Research and Technology/National Agency for Research and Innovation | Hidayat Trimarsanto, Iskandar Adnan, Lydia V. Panggalo, Sukma Oktavianthi, Willy Agustine, Edison Johar, Frilasita A Yudhaputri, Safarina G Malik, Khin Saw Myint, Amin Soebandrio |
| EPI_ISL_1284087 | RSJ Dr. Soeharto Heerdjan | Eijkman Institute for Molecular Biology, Ministry of Research and Technology/National Agency for Research and Innovation | Hidayat Trimarsanto, Iskandar Adnan, Lydia V. Panggalo, Sukma Oktavianthi, Willy Agustine, Edison Johar, Frilasita A Yudhaputri, Safarina G Malik, Khin Saw Myint, Amin Soebandrio |
| EPI_ISL_1284090 | RSU Sumber Waras | Eijkman Institute for Molecular Biology, Ministry of Research and Technology/National Agency for Research and Innovation | Hidayat Trimarsanto, Iskandar Adnan, Lydia V. Panggalo, Sukma Oktavianthi, Willy Agustine, Edison Johar, Frilasita A Yudhaputri, Safarina G Malik, Khin Saw Myint, Amin Soebandrio |
| EPI_ISL_1284093, EPI_ISL_1284096 | RSU Sumber Waras | Eijkman Institute for Molecular Biology, Ministry of Research and Technology/National Agency for Research and Innovation | Iskandar Adnan, Lydia V. Panggalo, Sukma Oktavianthi, Willy Agustine, Edison Johar, Hidayat Trimarsanto, Frilasita A Yudhaputri, Safarina G Malik, Khin Saw Myint, Amin Soebandrio |
| EPI_ISL_1284101 | RS Mitra Keluarga Kalideres | Eijkman Institute for Molecular Biology, Ministry of Research and Technology/National Agency for Research and Innovation | Iskandar Adnan, Lydia V. Panggalo, Sukma Oktavianthi, Willy Agustine, Edison Johar, Hidayat Trimarsanto, Frilasita A Yudhaputri, Safarina G Malik, Khin Saw Myint, Amin Soebandrio |
| EPI_ISL_1284105 | RS Mitra Husada Tangerang | Eijkman Institute for Molecular Biology, Ministry of Research and Technology/National Agency for Research and Innovation | Iskandar Adnan, Lydia V. Panggalo, Sukma Oktavianthi, Willy Agustine, Edison Johar, Hidayat Trimarsanto, Frilasita A Yudhaputri, Safarina G Malik, Khin Saw Myint, Amin Soebandrio |
| EPI_ISL_1284109, EPI_ISL_1284114 | RSU HGA Depok | Eijkman Institute for Molecular Biology, Ministry of Research and Technology/National Agency for Research and Innovation | Iskandar Adnan, Lydia V. Panggalo, Sukma Oktavianthi, Willy Agustine, Edison Johar, Hidayat Trimarsanto, Frilasita A Yudhaputri, Safarina G Malik, Khin Saw Myint, Amin Soebandrio |
| EPI_ISL_1284118 | RSU HGA Depok | Eijkman Institute for Molecular Biology, Ministry of Research and Technology/National Agency for Research and Innovation | Lydia V. Panggalo, Sukma Oktavianthi, Willy Agustine, Edison Johar, Hidayat Trimarsanto, Iskandar Adnan, Frilasita A Yudhaputri, Safarina G Malik, Khin Saw Myint, Amin Soebandrio |
| EPI_ISL_1284121, EPI_ISL_1284125 | RS Bhakti Mulia | Eijkman Institute for Molecular Biology, Ministry of Research and Technology/National Agency for Research and Innovation | Lydia V. Panggalo, Sukma Oktavianthi, Willy Agustine, Edison Johar, Hidayat Trimarsanto, Iskandar Adnan, Frilasita A Yudhaputri, Safarina G Malik, Khin Saw Myint, Amin Soebandrio |
| EPI_ISL_1284129 | RS Medika Permata Hijau | Eijkman Institute for Molecular Biology, Ministry of Research and Technology/National Agency for Research and Innovation | Lydia V. Panggalo, Sukma Oktavianthi, Willy Agustine, Edison Johar, Hidayat Trimarsanto, Iskandar Adnan, Frilasita A Yudhaputri, Safarina G Malik, Khin Saw Myint, Amin Soebandrio |
| EPI_ISL_1284133, EPI_ISL_1284136 | RS Mitra Keluarga Gading Serpong | Eijkman Institute for Molecular Biology, Ministry of Research and Technology/National Agency for Research and Innovation | Lydia V. Panggalo, Sukma Oktavianthi, Willy Agustine, Edison Johar, Hidayat Trimarsanto, Iskandar Adnan, Frilasita A Yudhaputri, Safarina G Malik, Khin Saw Myint, Amin Soebandrio |
| EPI_ISL_1284140, EPI_ISL_1284145 | RSAB Harapan Kita | Eijkman Institute for Molecular Biology, Ministry of Research and Technology/National Agency for Research and Innovation | Sukma Oktavianthi, Willy Agustine, Edison Johar, Hidayat Trimarsanto, Iskandar Adnan, Lydia V. Panggalo, Frilasita A Yudhaputri, Safarina G Malik, Khin Saw Myint, Amin Soebandrio |
| EPI_ISL_1284149, EPI_ISL_1284152 | RSIA Bunda Sejahtera | Eijkman Institute for Molecular Biology, Ministry of Research and | Sukma Oktavianthi, Willy Agustine, Edison Johar, Hidayat Trimarsanto, Iskandar Adnan, Lydia V. Panggalo, Frilasita A Yudhaputri, Safarina G Malik, Khin Saw Myint, Amin Soebandrio |

| | | | |
|---|---|---|---|
| | | Technology/National Agency for Research and Innovation | |
| EPI_ISL_1284156 | RS Sentra Medika Cikarang | Eijkman Institute for Molecular Biology, Ministry of Research and Technology/National Agency for Research and Innovation | Sukma Oktavianthi, Willy Agustine, Edison Johar, Hidayat Trimarsanto, Iskandar Adnan, Lydia V. Panggalo, Frilasita A Yudhaputri, Safarina G Malik, Khin Saw Myint, Amin Soebandrio |
| EPI_ISL_1284160 | RS Sentra Medika Cikarang | Eijkman Institute for Molecular Biology, Ministry of Research and Technology/National Agency for Research and Innovation | Willy Agustine, Edison Johar, Hidayat Trimarsanto, Iskandar Adnan, Lydia V. Panggalo, Sukma Oktavianthi, Frilasita A Yudhaputri, Safarina G Malik, Khin Saw Myint, Amin Soebandrio |
| EPI_ISL_1284163, EPI_ISL_1284165 | RS Mitra Keluarga Kalideres | Eijkman Institute for Molecular Biology, Ministry of Research and Technology/National Agency for Research and Innovation | Willy Agustine, Edison Johar, Hidayat Trimarsanto, Iskandar Adnan, Lydia V. Panggalo, Sukma Oktavianthi, Frilasita A Yudhaputri, Safarina G Malik, Khin Saw Myint, Amin Soebandrio |
| EPI_ISL_1284168 | RSU Harapan Bunda | Eijkman Institute for Molecular Biology, Ministry of Research and Technology/National Agency for Research and Innovation | Willy Agustine, Edison Johar, Hidayat Trimarsanto, Iskandar Adnan, Lydia V. Panggalo, Sukma Oktavianthi, Frilasita A Yudhaputri, Safarina G Malik, Khin Saw Myint, Amin Soebandrio |
| EPI_ISL_1284169 | RSU Azra | Eijkman Institute for Molecular Biology, Ministry of Research and Technology/National Agency for Research and Innovation | Willy Agustine, Edison Johar, Hidayat Trimarsanto, Iskandar Adnan, Lydia V. Panggalo, Sukma Oktavianthi, Frilasita A Yudhaputri, Safarina G Malik, Khin Saw Myint, Amin Soebandrio |
| EPI_ISL_1284172 | RSU Azra | Eijkman Institute for Molecular Biology, Ministry of Research and Technology/National Agency for Research and Innovation | Edison Johar, Frilasita A Yudhaputri, Hidayat Trimarsanto, Iskandar Adnan, Lydia V. Panggalo, Sukma Oktavianthi, Willy Agustine, Safarina G Malik, Khin Saw Myint, Amin Soebandrio |
| EPI_ISL_1284174 | RSU Sumber Waras | Eijkman Institute for Molecular Biology, Ministry of Research and Technology/National Agency for Research and Innovation | Edison Johar, Frilasita A Yudhaputri, Hidayat Trimarsanto, Iskandar Adnan, Lydia V. Panggalo, Sukma Oktavianthi, Willy Agustine, Safarina G Malik, Khin Saw Myint, Amin Soebandrio |
| EPI_ISL_1284176 | RS Harapan Mulia Cibarusah | Eijkman Institute for Molecular Biology, Ministry of Research and Technology/National Agency for Research and Innovation | Edison Johar, Frilasita A Yudhaputri, Hidayat Trimarsanto, Iskandar Adnan, Lydia V. Panggalo, Sukma Oktavianthi, Willy Agustine, Safarina G Malik, Khin Saw Myint, Amin Soebandrio |
| EPI_ISL_1284178 | RSIA Selasih Medika | Eijkman Institute for Molecular Biology, Ministry of Research and Technology/National Agency for Research and Innovation | Edison Johar, Frilasita A Yudhaputri, Hidayat Trimarsanto, Iskandar Adnan, Lydia V. Panggalo, Sukma Oktavianthi, Willy Agustine, Safarina G Malik, Khin Saw Myint, Amin Soebandrio |
| EPI_ISL_1284182 | RSJ Dr. Soeharto Heerdjan | Eijkman Institute for Molecular Biology, Ministry of Research and Technology/National Agency for Research and Innovation | Edison Johar, Frilasita A Yudhaputri, Hidayat Trimarsanto, Iskandar Adnan, Lydia V. Panggalo, Sukma Oktavianthi, Willy Agustine, Safarina G Malik, Khin Saw Myint, Amin Soebandrio |
| EPI_ISL_1284186 | RS Cendana | Eijkman Institute for Molecular Biology, Ministry of Research and Technology/National Agency for Research and Innovation | Edison Johar, Frilasita A Yudhaputri, Hidayat Trimarsanto, Iskandar Adnan, Lydia V. Panggalo, Sukma Oktavianthi, Willy Agustine, Safarina G Malik, Khin Saw Myint, Amin Soebandrio |
| EPI_ISL_1284190 | RS Permata Cibubur | Eijkman Institute for Molecular Biology, Ministry of Research and Technology/National Agency for Research and Innovation | Edison Johar, Frilasita A Yudhaputri, Hidayat Trimarsanto, Iskandar Adnan, Lydia V. Panggalo, Sukma Oktavianthi, Willy Agustine, Safarina G Malik, Khin Saw Myint, Amin Soebandrio |
| EPI_ISL_1284193 | RS Permata Cibubur | Eijkman Institute for Molecular Biology, Ministry of Research and Technology/National Agency for Research and Innovation | Frilasita A Yudhaputri, Hidayat Trimarsanto, Iskandar Adnan, Lydia V. Panggalo, Sukma Oktavianthi, Willy Agustine, Edison Johar, Safarina G Malik, Khin Saw Myint, Amin Soebandrio |

| | | | |
|---|---|---|---|
| EPI_ISL_1284197 | RS Mitra Keluarga Pratama Jatiasih | Eijkman Institute for Molecular Biology, Ministry of Research and Technology/National Agency for Research and Innovation | Frilasita A Yudhaputri, Hidayat Trimarsanto, Iskandar Adnan, Lydia V. Panggalo, Sukma Oktavianthi, Willy Agustine, Edison Johar, Safarina G Malik, Khin Saw Myint, Amin Soebandrio |
| EPI_ISL_1284201 | RS Bhakti Husada Cikarang | Eijkman Institute for Molecular Biology, Ministry of Research and Technology/National Agency for Research and Innovation | Frilasita A Yudhaputri, Hidayat Trimarsanto, Iskandar Adnan, Lydia V. Panggalo, Sukma Oktavianthi, Willy Agustine, Edison Johar, Safarina G Malik, Khin Saw Myint, Amin Soebandrio |
| EPI_ISL_1284204 | Klinik Ciputra Medical Center | Eijkman Institute for Molecular Biology, Ministry of Research and Technology/National Agency for Research and Innovation | Frilasita A Yudhaputri, Hidayat Trimarsanto, Iskandar Adnan, Lydia V. Panggalo, Sukma Oktavianthi, Willy Agustine, Edison Johar, Safarina G Malik, Khin Saw Myint, Amin Soebandrio |
| EPI_ISL_1284207 | RS Mitra Keluarga Cibubur | Eijkman Institute for Molecular Biology, Ministry of Research and Technology/National Agency for Research and Innovation | Frilasita A Yudhaputri, Hidayat Trimarsanto, Iskandar Adnan, Lydia V. Panggalo, Sukma Oktavianthi, Willy Agustine, Edison Johar, Safarina G Malik, Khin Saw Myint, Amin Soebandrio |
| EPI_ISL_1284210 | RS Mitra Husada Pringsewu | Eijkman Institute for Molecular Biology, Ministry of Research and Technology/National Agency for Research and Innovation | Hidayat Trimarsanto, Iskandar Adnan, Lydia V. Panggalo, Sukma Oktavianthi, Willy Agustine, Edison Johar, Frilasita A Yudhaputri, Safarina G Malik, Khin Saw Myint, Amin Soebandrio |
| EPI_ISL_1284212 | RSU Medirossa | Eijkman Institute for Molecular Biology, Ministry of Research and Technology/National Agency for Research and Innovation | Hidayat Trimarsanto, Iskandar Adnan, Lydia V. Panggalo, Sukma Oktavianthi, Willy Agustine, Edison Johar, Frilasita A Yudhaputri, Safarina G Malik, Khin Saw Myint, Amin Soebandrio |
| EPI_ISL_1284215 | RS Medirossa Cikarang | Eijkman Institute for Molecular Biology, Ministry of Research and Technology/National Agency for Research and Innovation | Iskandar Adnan, Lydia V. Panggalo, Sukma Oktavianthi, Willy Agustine, Edison Johar, Hidayat Trimarsanto, Frilasita A Yudhaputri, Safarina G Malik, Khin Saw Myint, Amin Soebandrio |
| EPI_ISL_1284219, EPI_ISL_1284223 | RS Mitra Keluarga Kalideres | Eijkman Institute for Molecular Biology, Ministry of Research and Technology/National Agency for Research and Innovation | Iskandar Adnan, Lydia V. Panggalo, Sukma Oktavianthi, Willy Agustine, Edison Johar, Hidayat Trimarsanto, Frilasita A Yudhaputri, Safarina G Malik, Khin Saw Myint, Amin Soebandrio |
| EPI_ISL_1284226, EPI_ISL_1284230, EPI_ISL_1284234 | RSU Harapan Bunda | Eijkman Institute for Molecular Biology, Ministry of Research and Technology/National Agency for Research and Innovation | Iskandar Adnan, Lydia V. Panggalo, Sukma Oktavianthi, Willy Agustine, Edison Johar, Hidayat Trimarsanto, Frilasita A Yudhaputri, Safarina G Malik, Khin Saw Myint, Amin Soebandrio |
| EPI_ISL_1284237 | RSU Harapan Bunda | Eijkman Institute for Molecular Biology, Ministry of Research and Technology/National Agency for Research and Innovation | Lydia V. Panggalo, Sukma Oktavianthi, Willy Agustine, Edison Johar, Hidayat Trimarsanto, Iskandar Adnan, Frilasita A Yudhaputri, Safarina G Malik, Khin Saw Myint, Amin Soebandrio |
| EPI_ISL_1284240 | RS Harapan Mulia Cibarusah | Eijkman Institute for Molecular Biology, Ministry of Research and Technology/National Agency for Research and Innovation | Lydia V. Panggalo, Sukma Oktavianthi, Willy Agustine, Edison Johar, Hidayat Trimarsanto, Iskandar Adnan, Frilasita A Yudhaputri, Safarina G Malik, Khin Saw Myint, Amin Soebandrio |
| EPI_ISL_1284244, EPI_ISL_1284247 | RS Masmitra | Eijkman Institute for Molecular Biology, Ministry of Research and Technology/National Agency for Research and Innovation | Lydia V. Panggalo, Sukma Oktavianthi, Willy Agustine, Edison Johar, Hidayat Trimarsanto, Iskandar Adnan, Frilasita A Yudhaputri, Safarina G Malik, Khin Saw Myint, Amin Soebandrio |
| EPI_ISL_1284251 | RS Mitra Keluarga Gading Serpong | Eijkman Institute for Molecular Biology, Ministry of Research and Technology/National Agency for Research and Innovation | Lydia V. Panggalo, Sukma Oktavianthi, Willy Agustine, Edison Johar, Hidayat Trimarsanto, Iskandar Adnan, Frilasita A Yudhaputri, Safarina G Malik, Khin Saw Myint, Amin Soebandrio |
| EPI_ISL_1284254 | RS Mitra Keluarga Gading Serpong | Eijkman Institute for Molecular Biology, Ministry of Research and | Sukma Oktavianthi, Willy Agustine, Edison Johar, Hidayat Trimarsanto, Iskandar Adnan, Lydia V. Panggalo, Frilasita A Yudhaputri, Safarina G Malik, Khin Saw Myint, Amin Soebandrio |

| | | | |
|---|---|---|---|
| | | Technology/National Agency for Research and Innovation | |
| EPI_ISL_1284258 | RS Mulia Pajajaran | Eijkman Institute for Molecular Biology, Ministry of Research and Technology/National Agency for Research and Innovation | Sukma Oktavianthi, Willy Agustine, Edison Johar, Hidayat Trimarsanto, Iskandar Adnan, Lydia V. Panggalo, Frilasita A Yudhaputri, Safarina G Malik, Khin Saw Myint, Amin Soebandrio |
| EPI_ISL_1284262 | RS OMNI Alam Sutera | Eijkman Institute for Molecular Biology, Ministry of Research and Technology/National Agency for Research and Innovation | Sukma Oktavianthi, Willy Agustine, Edison Johar, Hidayat Trimarsanto, Iskandar Adnan, Lydia V. Panggalo, Frilasita A Yudhaputri, Safarina G Malik, Khin Saw Myint, Amin Soebandrio |
| EPI_ISL_1284266 | RS Mulya Tangerang | Eijkman Institute for Molecular Biology, Ministry of Research and Technology/National Agency for Research and Innovation | Sukma Oktavianthi, Willy Agustine, Edison Johar, Hidayat Trimarsanto, Iskandar Adnan, Lydia V. Panggalo, Frilasita A Yudhaputri, Safarina G Malik, Khin Saw Myint, Amin Soebandrio |
| EPI_ISL_1284269 | RS Mitra Keluarga Bekasi Barat | Eijkman Institute for Molecular Biology, Ministry of Research and Technology/National Agency for Research and Innovation | Willy Agustine, Edison Johar, Hidayat Trimarsanto, Iskandar Adnan, Lydia V. Panggalo, Sukma Oktavianthi, Frilasita A Yudhaputri, Safarina G Malik, Khin Saw Myint, Amin Soebandrio |
| EPI_ISL_1284273, EPI_ISL_1284277 | RS Medirossa Cikarang | Eijkman Institute for Molecular Biology, Ministry of Research and Technology/National Agency for Research and Innovation | Willy Agustine, Edison Johar, Hidayat Trimarsanto, Iskandar Adnan, Lydia V. Panggalo, Sukma Oktavianthi, Frilasita A Yudhaputri, Safarina G Malik, Khin Saw Myint, Amin Soebandrio |
| EPI_ISL_1284281 | RS Royal Taruma | Eijkman Institute for Molecular Biology, Ministry of Research and Technology/National Agency for Research and Innovation | Willy Agustine, Edison Johar, Hidayat Trimarsanto, Iskandar Adnan, Lydia V. Panggalo, Sukma Oktavianthi, Frilasita A Yudhaputri, Safarina G Malik, Khin Saw Myint, Amin Soebandrio |
| EPI_ISL_1284284, EPI_ISL_1284288 | RS Mitra Keluarga Kalideres | Eijkman Institute for Molecular Biology, Ministry of Research and Technology/National Agency for Research and Innovation | Willy Agustine, Edison Johar, Hidayat Trimarsanto, Iskandar Adnan, Lydia V. Panggalo, Sukma Oktavianthi, Frilasita A Yudhaputri, Safarina G Malik, Khin Saw Myint, Amin Soebandrio |
| EPI_ISL_1284292 | RS Mitra Keluarga Kalideres | Eijkman Institute for Molecular Biology, Ministry of Research and Technology/National Agency for Research and Innovation | Edison Johar, Frilasita A Yudhaputri, Hidayat Trimarsanto, Iskandar Adnan, Lydia V. Panggalo, Sukma Oktavianthi, Willy Agustine, Safarina G Malik, Khin Saw Myint, Amin Soebandrio |
| EPI_ISL_1284296 | RSJ Dr. Soeharto Heerdjan | Eijkman Institute for Molecular Biology, Ministry of Research and Technology/National Agency for Research and Innovation | Edison Johar, Frilasita A Yudhaputri, Hidayat Trimarsanto, Iskandar Adnan, Lydia V. Panggalo, Sukma Oktavianthi, Willy Agustine, Safarina G Malik, Khin Saw Myint, Amin Soebandrio |
| EPI_ISL_1284299 | RSU HGA Depok | Eijkman Institute for Molecular Biology, Ministry of Research and Technology/National Agency for Research and Innovation | Edison Johar, Frilasita A Yudhaputri, Hidayat Trimarsanto, Iskandar Adnan, Lydia V. Panggalo, Sukma Oktavianthi, Willy Agustine, Safarina G Malik, Khin Saw Myint, Amin Soebandrio |
| EPI_ISL_1284303 | RS Grha Kedoya | Eijkman Institute for Molecular Biology, Ministry of Research and Technology/National Agency for Research and Innovation | Edison Johar, Frilasita A Yudhaputri, Hidayat Trimarsanto, Iskandar Adnan, Lydia V. Panggalo, Sukma Oktavianthi, Willy Agustine, Safarina G Malik, Khin Saw Myint, Amin Soebandrio |
| EPI_ISL_1284307 | RS Mitra Keluarga Gading Serpong | Eijkman Institute for Molecular Biology, Ministry of Research and Technology/National Agency for Research and Innovation | Edison Johar, Frilasita A Yudhaputri, Hidayat Trimarsanto, Iskandar Adnan, Lydia V. Panggalo, Sukma Oktavianthi, Willy Agustine, Safarina G Malik, Khin Saw Myint, Amin Soebandrio |
| EPI_ISL_1284310 | RS Bhakti Husada Cikarang | Eijkman Institute for Molecular Biology, Ministry of Research and Technology/National Agency for Research and Innovation | Edison Johar, Frilasita A Yudhaputri, Hidayat Trimarsanto, Iskandar Adnan, Lydia V. Panggalo, Sukma Oktavianthi, Willy Agustine, Safarina G Malik, Khin Saw Myint, Amin Soebandrio |

| | | | |
|---|---|---|---|
| EPI_ISL_1284314 | RS Mitra Keluarga Kalideres | Eijkman Institute for Molecular Biology, Ministry of Research and Technology/National Agency for Research and Innovation | Edison Johar, Frilasita A Yudhaputri, Hidayat Trimarsanto, Iskandar Adnan, Lydia V. Panggalo, Sukma Oktavianthi, Willy Agustine, Safarina G Malik, Khin Saw Myint, Amin Soebandrio |
| EPI_ISL_1284318 | RS Sentra Medika Cikarang | Eijkman Institute for Molecular Biology, Ministry of Research and Technology/National Agency for Research and Innovation | Edison Johar, Frilasita A Yudhaputri, Hidayat Trimarsanto, Iskandar Adnan, Lydia V. Panggalo, Sukma Oktavianthi, Willy Agustine, Safarina G Malik, Khin Saw Myint, Amin Soebandrio |
| EPI_ISL_1284322 | RS Islam Sari Asih Ar-Rahmah | Eijkman Institute for Molecular Biology, Ministry of Research and Technology/National Agency for Research and Innovation | Frilasita A Yudhaputri, Hidayat Trimarsanto, Iskandar Adnan, Lydia V. Panggalo, Sukma Oktavianthi, Willy Agustine, Edison Johar, Safarina G Malik, Khin Saw Myint, Amin Soebandrio |
| EPI_ISL_1284327, EPI_ISL_1284331, EPI_ISL_1284335 | RS Mitra Keluarga Kalideres | Eijkman Institute for Molecular Biology, Ministry of Research and Technology/National Agency for Research and Innovation | Frilasita A Yudhaputri, Hidayat Trimarsanto, Iskandar Adnan, Lydia V. Panggalo, Sukma Oktavianthi, Willy Agustine, Edison Johar, Safarina G Malik, Khin Saw Myint, Amin Soebandrio |
| EPI_ISL_1284340 | RS Sentra Medika Cikarang | Eijkman Institute for Molecular Biology, Ministry of Research and Technology/National Agency for Research and Innovation | Hidayat Trimarsanto, Iskandar Adnan, Lydia V. Panggalo, Sukma Oktavianthi, Willy Agustine, Edison Johar, Frilasita A Yudhaputri, Safarina G Malik, Khin Saw Myint, Amin Soebandrio |
| EPI_ISL_1284344 | RSU Sumber Waras | Eijkman Institute for Molecular Biology, Ministry of Research and Technology/National Agency for Research and Innovation | Hidayat Trimarsanto, Iskandar Adnan, Lydia V. Panggalo, Sukma Oktavianthi, Willy Agustine, Edison Johar, Frilasita A Yudhaputri, Safarina G Malik, Khin Saw Myint, Amin Soebandrio |
| EPI_ISL_1297194 | RSUD Pasarminggu | Stem Cell Lab. Universitas Pembangunan Nasional Veteran Jakarta (UPNVJ) | Andri Pramesyanti Pramono, Yudhi Nugraha, Ria Maria Theresa, Yudi Amiarno, Megadianty Mokoginta |
| EPI_ISL_1300496 | RSUD Pasarminggu | Stem Cell Lab, Universitas Pembangunan Nasional Veteran Jakarta (UPNVJ) | Andri Pramesyanti Pramono, Yudhi Nugraha, Ria Maria Theresa, Yudi Amiarno, Megadianty Mokoginta |
| EPI_ISL_1300521 | RSUD Pasarminggu | Stem Cell Lab, Universitas Pembangunan Nasional Veteran Jakarta | Andri Pramesyanti Pramono, Yudhi Nugraha, Ria Maria Theresa, Yudi Amiarno, Megadianty Mokoginta |
| EPI_ISL_1300522, EPI_ISL_1300535, EPI_ISL_1300540, EPI_ISL_1300644, EPI_ISL_1300660, EPI_ISL_1300661 | RSUD Pasarminggu | Stem Cell Lab, Universitas Pembangunan Nasional Veteran Jakarta (UPNVJ) | Andri Pramesyanti Pramono, Yudhi Nugraha, Ria Maria Theresa, Yudi Amiarno, Megadianty Mokoginta |
| EPI_ISL_1335883 | North Sumatera Health Office | Faculty of Medicine, Universitas Sumatera Utara; Institute of Tropical Disease, Universitas Airlangga | Aldise M Nastri, Jezzy R Dewantari, Rima R Prasetya, Krisnoadi Rahardjo, Yasuko Mori, Soetjipto, Kazufumi Shimizu, Maria I Lusida, Meliani, R Andika D Cahyadi, Mirzan Hasibuan, Irbah R Nainggolan, Ramadhan Bestari, Franciscus Ginting, [illegible] |
| EPI_ISL_1335884 | Department of Microbiology, Faculty of Medicine, Universitas Sumatera Utara | Faculty of Medicine, Universitas Sumatera Utara; Institute of Tropical Disease, Universitas Airlangga | Jezzy R Dewantari, Rima R Prasetya, Krisnoadi Rahardjo, Aldise M Nastri, Yasuko Mori, Soetjipto, Kazufumi Shimizu, Maria I Lusida, Meliani, R Andika D Cahyadi, Mirzan Hasibuan, Irbah R Nainggolan, Ramadhan Bestari, Franciscus Ginting, [illegible] |
| EPI_ISL_1335885 | North Sumatera Health Office | Faculty of Medicine, Universitas Sumatera Utara; Institute of Tropical Disease, Universitas Airlangga | Rima R Prasetya, Krisnoadi Rahardjo, Aldise M Nastri, Jezzy R Dewantari, Yasuko Mori, Soetjipto, Kazufumi Shimizu, Maria I Lusida, Meliani, R Andika D Cahyadi, Mirzan Hasibuan, Irbah R Nainggolan, Ramadhan Bestari, Franciscus Ginting, Muhamr |
| EPI_ISL_1335886 | Sundari Hospital Medan | Faculty of Medicine, Universitas Sumatera Utara; Institute of Tropical Disease, Universitas Airlangga | Kazufumi Shimizu, Krisnoadi Rahardjo, Aldise M Nastri, Jezzy R Dewantari, Rima R Prasetya, Yasuko Mori, Soetjipto, Maria I Lusida, Meliani, R Andika D Cahyadi, Mirzan Hasibuan, Irbah R Nainggolan, Ramadhan Bestari, Franciscus Ginting, Muhamr |
| EPI_ISL_1335887 | North Sumatera Health Office | Faculty of Medicine, Universitas Sumatera Utara; Institute of Tropical Disease, Universitas Airlangga | Maria I Lusida, Krisnoadi Rahardjo, Aldise M Nastri, Jezzy R Dewantari, Rima R Prasetya, Yasuko Mori, Soetjipto, Kazufumi Shimizu, Meliani, R Andika D Cahyadi, Mirzan Hasibuan, Irbah R Nainggolan, Ramadhan Bestari, Franciscus Ginting, Muhamr |
| EPI_ISL_1335888 | Martha Friska Multatuli Hospital Medan | Faculty of Medicine, Universitas Sumatera | Krisnoadi Rahardjo, Aldise M Nastri, Jezzy R Dewantari, Rima R Prasetya, Yasuko Mori, Soetjipto, Kazufumi Shimizu, Maria I Lusida, Meliani, R Andika D Cahyadi, Mirzan Hasibuan, Irbah R Nainggolan, Ramadhan Bestari, Franciscus Ginting, Muhamr |

| | | | |
|---|---|---|---|
| | | Utara; Institute of Tropical Disease, Universitas Airlangga | |
| EPI_ISL_1335889 | Advent Hospital Medan | Faculty of Medicine, Universitas Sumatera Utara; Institute of Tropical Disease, Universitas Airlangga | Krisnoadi Rahardjo, Aldise M Nastri, Jezzy R Dewantari, Rima R Prasetya, Yasuko Mori, Soetjipto, Kazufumi Shimizu, Maria I Lusida, Meliani, R Andika D Cahyadi, Mirzan Hasibuan, Irbah R Nainggolan, Ramadhan Bestari, Franciscus Ginting, Muhamr |
| EPI_ISL_1335890 | Elisabeth Hospital Medan | Faculty of Medicine, Universitas Sumatera Utara; Institute of Tropical Disease, Universitas Airlangga | Aldise M Nastri, Jezzy R Dewantari, Rima R Prasetya, Krisnoadi Rahardjo, Yasuko Mori, Soetjipto, Kazufumi Shimizu, Maria I Lusida, Meliani, R Andika D Cahyadi, Mirzan Hasibuan, Irbah R Nainggolan, Ramadhan Bestari, Franciscus Ginting, Muhamr |
| EPI_ISL_1335891 | Advent Hospital Medan | Faculty of Medicine, Universitas Sumatera Utara; Institute of Tropical Disease, Universitas Airlangga | Jezzy R Dewantari, Rima R Prasetya, Krisnoadi Rahardjo, Aldise M Nastri, Yasuko Mori, Soetjipto, Kazufumi Shimizu, Maria I Lusida, Meliani, R Andika D Cahyadi, Mirzan Hasibuan, Irbah R Nainggolan, Ramadhan Bestari, Franciscus Ginting, Muhamr |
| EPI_ISL_1335892 | Department of Microbiology, Faculty of Medicine, Universitas Sumatera Utara | Faculty of Medicine, Universitas Sumatera Utara; Institute of Tropical Disease, Universitas Airlangga | Rima R Prasetya, Krisnoadi Rahardjo, Aldise M Nastri, Jezzy R Dewantari, Yasuko Mori, Soetjipto, Kazufumi Shimizu, Maria I Lusida, Meliani, R Andika D Cahyadi, Mirzan Hasibuan, Irbah R Nainggolan, Ramadhan Bestari, Franciscus Ginting, Muhamr |
| EPI_ISL_1335893 | Department of Microbiology, Faculty of Medicine, Universitas Sumatera Utara | Faculty of Medicine, Universitas Sumatera Utara; Institute of Tropical Disease, Universitas Airlangga | Kazufumi Shimizu, Krisnoadi Rahardjo, Aldise M Nastri, Jezzy R Dewantari, Rima R Prasetya, Yasuko Mori, Soetjipto, Maria I Lusida, Meliani, R Andika D Cahyadi, Mirzan Hasibuan, Irbah R Nainggolan, Ramadhan Bestari, Franciscus Ginting, Muhamr |
| EPI_ISL_1335894 | Department of Microbiology, Faculty of Medicine, Universitas Sumatera Utara | Faculty of Medicine, Universitas Sumatera Utara; Institute of Tropical Disease, Universitas Airlangga | Maria I Lusida, Krisnoadi Rahardjo, Aldise M Nastri, Jezzy R Dewantari, Rima R Prasetya, Yasuko Mori, Soetjipto, Kazufumi Shimizu, Meliani, R Andika D Cahyadi, Mirzan Hasibuan, Irbah R Nainggolan, Ramadhan Bestari, Franciscus Ginting, Muhamr |
| EPI_ISL_1335895 | Martha Friska Multatuli Hospital Medan | Faculty of Medicine, Universitas Sumatera Utara; Institute of Tropical Disease, Universitas Airlangga | Krisnoadi Rahardjo, Aldise M Nastri, Jezzy R Dewantari, Rima R Prasetya, Yasuko Mori, Soetjipto, Kazufumi Shimizu, Maria I Lusida, Meliani, R Andika D Cahyadi, Mirzan Hasibuan, Irbah R Nainggolan, Ramadhan Bestari, Franciscus Ginting, Muhamr |
| EPI_ISL_1335896 | Columbia Asia Hospital Medan | Faculty of Medicine, Universitas Sumatera Utara; Institute of Tropical Disease, Universitas Airlangga | Aldise M Nastri, Jezzy R Dewantari, Rima R Prasetya, Krisnoadi Rahardjo, Yasuko Mori, Soetjipto, Kazufumi Shimizu, Maria I Lusida, Meliani, R Andika D Cahyadi, Mirzan Hasibuan, Irbah R Nainggolan, Ramadhan Bestari, Franciscus Ginting, Muhamr |
| EPI_ISL_1335897 | North Sumatera Health Office | Faculty of Medicine, Universitas Sumatera Utara; Institute of Tropical Disease, Universitas Airlangga | Jezzy R Dewantari, Rima R Prasetya, Krisnoadi Rahardjo, Aldise M Nastri, Yasuko Mori, Soetjipto, Kazufumi Shimizu, Maria I Lusida, Meliani, R Andika D Cahyadi, Mirzan Hasibuan, Irbah R Nainggolan, Ramadhan Bestari, Franciscus Ginting, Muhamr |
| EPI_ISL_1335898 | Department of Microbiology, Faculty of Medicine, Universitas Sumatera Utara | Faculty of Medicine, Universitas Sumatera Utara; Institute of Tropical Disease, Universitas Airlangga | Rima R Prasetya, Krisnoadi Rahardjo, Aldise M Nastri, Jezzy R Dewantari, Yasuko Mori, Soetjipto, Kazufumi Shimizu, Maria I Lusida, Meliani, R Andika D Cahyadi, Mirzan Hasibuan, Irbah R Nainggolan, Ramadhan Bestari, Franciscus Ginting, Muhamr |
| EPI_ISL_1335899 | Department of Microbiology, Faculty of Medicine, Universitas Sumatera Utara | Faculty of Medicine, Universitas Sumatera Utara; Institute of Tropical Disease, Universitas Airlangga | Kazufumi Shimizu, Krisnoadi Rahardjo, Aldise M Nastri, Jezzy R Dewantari, Rima R Prasetya, Yasuko Mori, Soetjipto, Maria I Lusida, Meliani, R Andika D Cahyadi, Mirzan Hasibuan, Irbah R Nainggolan, Ramadhan Bestari, Franciscus Ginting, Muhamr |
| EPI_ISL_1340862, EPI_ISL_1340863, EPI_ISL_1340864, EPI_ISL_1340865, EPI_ISL_1340866, EPI_ISL_1340867, EPI_ISL_1341792, EPI_ISL_1341793, EPI_ISL_1341794, EPI_ISL_1341795, EPI_ISL_1341796, EPI_ISL_1341797 | | | |
| see above | West Bandung Public Health | West Java Health Laboratory; School of Life Sciences and Technology, Institut Teknologi Bandung | Azzania Fibriani, Ema Rahmawati, Ryan Bayusantika Ristandi, Rifky Waluyajati Rachman, Cut Nur Cinthia Alamanda, Isak Solihin, Rini Robiani, Miftahul Faridl, Karimatu Khoirunnisa |
| EPI_ISL_1341798 | Depok Public Health | West Java Health Laboratory; School of Life Sciences and Technology, Institut Teknologi Bandung | Azzania Fibriani, Ema Rahmawati, Ryan Bayusantika Ristandi, Rifky Waluyajati Rachman, Cut Nur Cinthia Alamanda, Isak Solihin, Rini Robiani, Miftahul Faridl, Karimatu Khoirunnisa |
| EPI_ISL_1341799, EPI_ISL_1341800 | Tasikmalaya Public Health | West Java Health Laboratory; School of Life Sciences and Technology, Institut Teknologi Bandung | Azzania Fibriani, Ema Rahmawati, Ryan Bayusantika Ristandi, Rifky Waluyajati Rachman, Cut Nur Cinthia Alamanda, Isak Solihin, Rini Robiani, Miftahul Faridl, Karimatu Khoirunnisa |
| EPI_ISL_1341801 | Sukabumi Health Laboratory | West Java Health Laboratory; School of Life Sciences and Technology, Institut Teknologi Bandung | Azzania Fibriani, Ema Rahmawati, Ryan Bayusantika Ristandi, Rifky Waluyajati Rachman, Cut Nur Cinthia Alamanda, Isak Solihin, Rini Robiani, Miftahul Faridl, Karimatu Khoirunnisa |
| EPI_ISL_1341802, EPI_ISL_1341907 | Subang Public Health | West Java Health Laboratory; School of Life Sciences and Technology, Institut Teknologi Bandung | Azzania Fibriani, Ema Rahmawati, Ryan Bayusantika Ristandi, Rifky Waluyajati Rachman, Cut Nur Cinthia Alamanda, Isak Solihin, Rini Robiani, Miftahul Faridl, Karimatu Khoirunnisa |

| | | | |
|---|---|---|---|
| EPI_ISL_1343218 | West Java Health Laboratory | West Java Health Laboratory; School of Life Sciences and Technology, Institut Teknologi Bandung | Azzania Fibriani, Ema Rahmawati, Ryan Bayusantika Ristandi, Rifky Waluyajati Rachman, Cut Nur Cinthia Alamanda, Isak Solihin, Rini Robiani, Miftahul Faridl, Karimatu Khoirunnisa |
| EPI_ISL_1343818 | Subang Public Health | West Java Health Laboratory; School of Life Sciences and Technology, Institut Teknologi Bandung | Azzania Fibriani, Ema Rahmawati, Ryan Bayusantika Ristandi, Rifky Waluyajati Rachman, Cut Nur Cinthia Alamanda, Isak Solihin, Rini Robiani, Miftahul Faridl, Karimatu Khoirunnisa |
| EPI_ISL_1344587 | West Java Health Laboratory | West Java Health Laboratory; School of Life Sciences and Technology, Institut Teknologi Bandung | Azzania Fibriani, Ema Rahmawati, Ryan Bayusantika Ristandi, Rifky Waluyajati Rachman, Cut Nur Cinthia Alamanda, Isak Solihin, Rini Robiani, Miftahul Faridl, Karimatu Khoirunnisa |
| EPI_ISL_1345232 | West Bandung Public Health | West Java Health Laboratory; School of Life Sciences and Technology, Institut Teknologi Bandung | Azzania Fibriani, Ema Rahmawati, Ryan Bayusantika Ristandi, Rifky Waluyajati Rachman, Cut Nur Cinthia Alamanda, Isak Solihin, Rini Robiani, Miftahul Faridl, Karimatu Khoirunnisa |
| EPI_ISL_1346047 | West Java Health Laboratory | West Java Health Laboratory; School of Life Sciences and Technology, Institut Teknologi Bandung | Azzania Fibriani, Ema Rahmawati, Ryan Bayusantika Ristandi, Rifky Waluyajati Rachman, Cut Nur Cinthia Alamanda, Isak Solihin, Rini Robiani, Miftahul Faridl, Karimatu Khoirunnisa |
| EPI_ISL_1346484 | West Bandung Public Health | West Java Health Laboratory; School of Life Sciences and Technology, Institut Teknologi Bandung | Azzania Fibriani, Ema Rahmawati, Ryan Bayusantika Ristandi, Rifky Waluyajati Rachman, Cut Nur Cinthia Alamanda, Isak Solihin, Rini Robiani, Miftahul Faridl, Karimatu Khoirunnisa |
| EPI_ISL_1346485 | Cianjur Public Health | West Java Health Laboratory; School of Life Sciences and Technology, Institut Teknologi Bandung | Azzania Fibriani, Ema Rahmawati, Ryan Bayusantika Ristandi, Rifky Waluyajati Rachman, Cut Nur Cinthia Alamanda, Isak Solihin, Rini Robiani, Miftahul Faridl, Karimatu Khoirunnisa |
| EPI_ISL_1346625, EPI_ISL_1346627 | Hasan Sadikin Hospital | West Java Health Laboratory; School of Life Sciences and Technology, Institut Teknologi Bandung | Azzania Fibriani, Ema Rahmawati, Ryan Bayusantika Ristandi, Rifky Waluyajati Rachman, Cut Nur Cinthia Alamanda, Isak Solihin, Rini Robiani, Miftahul Faridl, Karimatu Khoirunnisa, Nina Susana Dewi, Anggraini Alam, Basti Andriyoko, |
| EPI_ISL_1364466, EPI_ISL_1364467, EPI_ISL_1364468, EPI_ISL_1366083, EPI_ISL_1366238, EPI_ISL_1366269, EPI_ISL_1366271 | Dr. Soetomo Teaching Hospital | Research Center for Vaccine Technology and Development, Institute of Tropical Disease, Airlangga University | Fedik Abdul Rantam, Cita Rosita Sigit Prakoeswa, Damayanti Tinduh, Jusak Nugraha, Helen Susilowati, Andi Yasmin Wijaya, Ni Nyoman Tri Puspaningsih, Dwiyanti Puspitasari, Dominicus Husada, Neneng Dewi Kur |
| EPI_ISL_1366503, EPI_ISL_1366505, EPI_ISL_1366509 | Indrapura Field Hospital (part of Dr. Soetomo Teaching Hospital) | Research Center for Vaccine Technology and Development, Institute of Tropical Disease, Airlangga University | Fedik Abdul Rantam, Christijogo Soemartono Waloejo, Cita Rosita Sigit Prakoeswa, Damayanti Tinduh, Jusak Nugraha, Helen Susilowati, Andi Yasmin Wijaya, Ni Nyoman Tri Puspaningsih, Dwiyanti Puspitasari, Dominicus Husada, N |
| EPI_ISL_1391009, EPI_ISL_1391010, EPI_ISL_1391011, EPI_ISL_1391012, EPI_ISL_1391013, EPI_ISL_1391014, EPI_ISL_1391015, EPI_ISL_1391016, EPI_ISL_1391017, EPI_ISL_1391018 | National Institute of Health Research and Development | National Institute of Health Research and Development | Subangkit, Hana Apsari Pawestri, Kartika Dewi Puspa, Arie Ardiansyah Nugraha, Hartanti Dian Ikawati, Krisna Nur Andriana Pangesti, Yuni Rukminiati, Ririn Ramadhany, Agustiningsih, Kindi Adam, Holy Arif Wibowo, Triyani Soekarso, Ni Ketut Susilarini, Nurika Ha Tati Febriyanti, Natalie Laurencia Kipuw, Fauzul Muna, Irene Lorinda Indalao, Nelly Puspandari, Vivi Setiawaty. |
| EPI_ISL_1398267 | Subang Public Health | West Java Health Laboratory; School of Life Sciences and Technology, Institut Teknologi Bandung | Azzania Fibriani, Ema Rahmawati, Ryan Bayusantika Ristandi, Rifky Waluyajati Rachman, Cut Nur Cinthia Alamanda, Isak Solihin, Rini Robiani, Miftahul Faridl, Karimatu Khoirunnisa, Kamila Tania |
| EPI_ISL_1398353 | Kuningan Public Health | West Java Health Laboratory; School of Life Sciences and Technology, Institut Teknologi Bandung | Azzania Fibriani, Ema Rahmawati, Ryan Bayusantika Ristandi, Rifky Waluyajati Rachman, Cut Nur Cinthia Alamanda, Isak Solihin, Rini Robiani, Miftahul Faridl, Karimatu Khoirunnisa, Kamila Tania |
| EPI_ISL_1398355 | Subang Public Health | West Java Health Laboratory; School of Life Sciences and Technology, Institut Teknologi Bandung | Azzania Fibriani, Ema Rahmawati, Ryan Bayusantika Ristandi, Rifky Waluyajati Rachman, Cut Nur Cinthia Alamanda, Isak Solihin, Rini Robiani, Miftahul Faridl, Karimatu Khoirunnisa, Kamila Tania |
| EPI_ISL_1398357 | Kasih Bunda Hospital | West Java Health Laboratory; School of Life Sciences and Technology, Institut Teknologi Bandung | Azzania Fibriani, Ema Rahmawati, Ryan Bayusantika Ristandi, Rifky Waluyajati Rachman, Cut Nur Cinthia Alamanda, Isak Solihin, Rini Robiani, Miftahul Faridl, Karimatu Khoirunnisa, Kamila Tania |
| EPI_ISL_1398359 | Subang Public Health | West Java Health Laboratory; School of Life Sciences and Technology, Institut Teknologi Bandung | Azzania Fibriani, Ema Rahmawati, Ryan Bayusantika Ristandi, Rifky Waluyajati Rachman, Cut Nur Cinthia Alamanda, Isak Solihin, Rini Robiani, Miftahul Faridl, Karimatu Khoirunnisa, Kamila Tania |
| EPI_ISL_1398361 | West Bandung Public Health | West Java Health Laboratory; School of Life Sciences and Technology, Institut Teknologi Bandung | Azzania Fibriani, Ema Rahmawati, Ryan Bayusantika Ristandi, Rifky Waluyajati Rachman, Cut Nur Cinthia Alamanda, Isak Solihin, Rini Robiani, Miftahul Faridl, Karimatu Khoirunnisa, Kamila Tania |
| EPI_ISL_1398363 | West Java Health | West Java Health | Azzania Fibriani, Ema Rahmawati, Ryan Bayusantika Ristandi, Rifky Waluyajati Rachman, Cut Nur Cinthia Alamanda, Isak Solihin, Rini Robiani, Miftahul Faridl, Karimatu Khoirunnisa, Kamila Tania |

| | | | |
|---|---|---|---|
| | Laboratory | Laboratory; School of Life Sciences and Technology, Institut Teknologi Bandung | |
| EPI_ISL_1398365 | Sumedang Public Health | West Java Health Laboratory; School of Life Sciences and Technology, Institut Teknologi Bandung | Azzania Fibriani, Ema Rahmawati, Ryan Bayusantika Ristandi, Rifky Waluyajati Rachman, Cut Nur Cinthia Alamanda, Isak Solihin, Rini Robiani, Miftahul Faridl, Karimatu Khoirunnisa, Kamila Tania |
| EPI_ISL_1398367 | Cianjur Public Health | West Java Health Laboratory; School of Life Sciences and Technology, Institut Teknologi Bandung | Azzania Fibriani, Ema Rahmawati, Ryan Bayusantika Ristandi, Rifky Waluyajati Rachman, Cut Nur Cinthia Alamanda, Isak Solihin, Rini Robiani, Miftahul Faridl, Karimatu Khoirunnisa, Kamila Tania |
| EPI_ISL_1398369 | Kuningan Public Health | West Java Health Laboratory; School of Life Sciences and Technology, Institut Teknologi Bandung | Azzania Fibriani, Ema Rahmawati, Ryan Bayusantika Ristandi, Rifky Waluyajati Rachman, Cut Nur Cinthia Alamanda, Isak Solihin, Rini Robiani, Miftahul Faridl, Karimatu Khoirunnisa, Kamila Tania |
| EPI_ISL_1398371 | West Java Health Laboratory | West Java Health Laboratory; School of Life Sciences and Technology, Institut Teknologi Bandung | Azzania Fibriani, Ema Rahmawati, Ryan Bayusantika Ristandi, Rifky Waluyajati Rachman, Cut Nur Cinthia Alamanda, Isak Solihin, Rini Robiani, Miftahul Faridl, Karimatu Khoirunnisa, Kamila Tania |
| EPI_ISL_1398374 | Sukabumi Public Health | West Java Health Laboratory; School of Life Sciences and Technology, Institut Teknologi Bandung | Azzania Fibriani, Ema Rahmawati, Ryan Bayusantika Ristandi, Rifky Waluyajati Rachman, Cut Nur Cinthia Alamanda, Isak Solihin, Rini Robiani, Miftahul Faridl, Karimatu Khoirunnisa, Kamila Tania |
| EPI_ISL_1398375, EPI_ISL_1398561 | Subang Public Health | West Java Health Laboratory; School of Life Sciences and Technology, Institut Teknologi Bandung | Azzania Fibriani, Ema Rahmawati, Ryan Bayusantika Ristandi, Rifky Waluyajati Rachman, Cut Nur Cinthia Alamanda, Isak Solihin, Rini Robiani, Miftahul Faridl, Karimatu Khoirunnisa, Kamila Tania |
| EPI_ISL_1398562 | West Java Health Laboratory | West Java Health Laboratory; School of Life Sciences and Technology, Institut Teknologi Bandung | Azzania Fibriani, Ema Rahmawati, Ryan Bayusantika Ristandi, Rifky Waluyajati Rachman, Cut Nur Cinthia Alamanda, Isak Solihin, Rini Robiani, Miftahul Faridl, Karimatu Khoirunnisa, Kamila Tania |
| EPI_ISL_1398563 | West Bandung Public Health | West Java Health Laboratory; School of Life Sciences and Technology, Institut Teknologi Bandung | Azzania Fibriani, Ema Rahmawati, Ryan Bayusantika Ristandi, Rifky Waluyajati Rachman, Cut Nur Cinthia Alamanda, Isak Solihin, Rini Robiani, Miftahul Faridl, Karimatu Khoirunnisa, Kamila Tania |
| EPI_ISL_1398565 | Cianjur Public Health | West Java Health Laboratory; School of Life Sciences and Technology, Institut Teknologi Bandung | Azzania Fibriani, Ema Rahmawati, Ryan Bayusantika Ristandi, Rifky Waluyajati Rachman, Cut Nur Cinthia Alamanda, Isak Solihin, Rini Robiani, Miftahul Faridl, Karimatu Khoirunnisa, Kamila Tania |
| EPI_ISL_1398567 | West Java Health Laboratory | West Java Health Laboratory; School of Life Sciences and Technology, Institut Teknologi Bandung | Azzania Fibriani, Ema Rahmawati, Ryan Bayusantika Ristandi, Rifky Waluyajati Rachman, Cut Nur Cinthia Alamanda, Isak Solihin, Rini Robiani, Miftahul Faridl, Karimatu Khoirunnisa, Kamila Tania |
| EPI_ISL_1398568 | West Bandung Public Health | West Java Health Laboratory; School of Life Sciences and Technology, Institut Teknologi Bandung | Azzania Fibriani, Ema Rahmawati, Ryan Bayusantika Ristandi, Rifky Waluyajati Rachman, Cut Nur Cinthia Alamanda, Isak Solihin, Rini Robiani, Miftahul Faridl, Karimatu Khoirunnisa, Kamila Tania |
| EPI_ISL_1398924, EPI_ISL_1398927 | Subang Public Health | West Java Health Laboratory; School of Life Sciences and Technology, Institut Teknologi Bandung | Azzania Fibriani, Ema Rahmawati, Ryan Bayusantika Ristandi, Rifky Waluyajati Rachman, Cut Nur Cinthia Alamanda, Isak Solihin, Rini Robiani, Miftahul Faridl, Karimatu Khoirunnisa, Kamila Tania |
| EPI_ISL_1398929 | West Bandung Public Health | West Java Health Laboratory; School of Life Sciences and Technology, Institut Teknologi Bandung | Azzania Fibriani, Ema Rahmawati, Ryan Bayusantika Ristandi, Rifky Waluyajati Rachman, Cut Nur Cinthia Alamanda, Isak Solihin, Rini Robiani, Miftahul Faridl, Karimatu Khoirunnisa, Kamila Tania |
| EPI_ISL_1398930 | Subang Public Health | West Java Health Laboratory; School of Life Sciences and Technology, Institut Teknologi Bandung | Azzania Fibriani, Ema Rahmawati, Ryan Bayusantika Ristandi, Rifky Waluyajati Rachman, Cut Nur Cinthia Alamanda, Isak Solihin, Rini Robiani, Miftahul Faridl, Karimatu Khoirunnisa, Kamila Tania |
| EPI_ISL_1398932 | Cianjur Public Health | West Java Health Laboratory; School of Life Sciences and Technology, Institut Teknologi Bandung | Azzania Fibriani, Ema Rahmawati, Ryan Bayusantika Ristandi, Rifky Waluyajati Rachman, Cut Nur Cinthia Alamanda, Isak Solihin, Rini Robiani, Miftahul Faridl, Karimatu Khoirunnisa, Kamila Tania |
| EPI_ISL_1398934 | West Bandung Public Health | West Java Health Laboratory | Azzania Fibriani, Ema Rahmawati, Ryan Bayusantika Ristandi, Rifky Waluyajati Rachman, Cut Nur Cinthia Alamanda, Isak Solihin, Rini Robiani, Miftahul Faridl, Karimatu Khoirunnisa, Kamila Tania |
| EPI_ISL_1398936 | Purwakarta Public Health | West Java Health Laboratory | Azzania Fibriani, Ema Rahmawati, Ryan Bayusantika Ristandi, Rifky Waluyajati Rachman, Cut Nur Cinthia Alamanda, Isak Solihin, Rini Robiani, Miftahul Faridl, Karimatu Khoirunnisa, Kamila Tania |
| EPI_ISL_1398939, EPI_ISL_1398942, EPI_ISL_1398955 | Cianjur Public Health | West Java Health Laboratory; School of Life Sciences and Technology, Institut Teknologi Bandung | Azzania Fibriani, Ema Rahmawati, Ryan Bayusantika Ristandi, Rifky Waluyajati Rachman, Cut Nur Cinthia Alamanda, Isak Solihin, Rini Robiani, Miftahul Faridl, Karimatu Khoirunnisa, Kamila Tania |
| EPI_ISL_1398980 | Sumedang Public Health | West Java Health Laboratory; School of Life Sciences and Technology, Institut Teknologi Bandung | Azzania Fibriani, Ema Rahmawati, Ryan Bayusantika Ristandi, Rifky Waluyajati Rachman, Cut Nur Cinthia Alamanda, Isak Solihin, Rini Robiani, Miftahul Faridl, Karimatu Khoirunnisa, Kamila Tania |
| EPI_ISL_1398981 | Subang Public Health | West Java Health Laboratory; School of Life | Azzania Fibriani, Ema Rahmawati, Ryan Bayusantika Ristandi, Rifky Waluyajati Rachman, Cut Nur Cinthia Alamanda, Isak Solihin, Rini Robiani, Miftahul Faridl, Karimatu Khoirunnisa, Kamila Tania |

| | | | |
|---|---|---|---|
| | | Sciences and Technology, Institut Teknologi Bandung | |
| EPI_ISL_1398983 | Cianjur Public Health | West Java Health Laboratory; School of Life Sciences and Technology, Institut Teknologi Bandung | Azzania Fibriani, Ema Rahmawati, Ryan Bayusantika Ristandi, Rifky Waluyajati Rachman, Cut Nur Cinthia Alamanda, Isak Solihin, Rini Robiani, Miftahul Faridl, Karimatu Khoirunnisa, Kamila Tania |
| EPI_ISL_1398984 | West Bandung Public Health | West Java Health Laboratory; School of Life Sciences and Technology, Institut Teknologi Bandung | Azzania Fibriani, Ema Rahmawati, Ryan Bayusantika Ristandi, Rifky Waluyajati Rachman, Cut Nur Cinthia Alamanda, Isak Solihin, Rini Robiani, Miftahul Faridl, Karimatu Khoirunnisa, Kamila Tania |
| EPI_ISL_1411159, EPI_ISL_1413427, EPI_ISL_1415427 | National Institute of Health Research and Development | National Institute of Health Research and Development | Subangkit, Hana Apsari Pawestri, Kartika Dewi Puspa, Arie Ardiansyah Nugraha, Hartanti Dian Ikawati, Krisna Nur Andriana Pangesti, Yuni Rukminiati, Ririn Ramadhany, Agustiningsih, Kindi Adam, Holy Arif Wibowo, Triyani Soekarso, Ni Ketut Susilarini, Nurika Ha Tati Febriyanti, Natalie Laurencia Kipuw, Fauzul Muna, Irene Lorinda Indalao, Nelly Puspandari, Vivi Setiawaty. |
| EPI_ISL_1415898 | Laboratorium Intibios Karawang | National Institute of Health Research and Development | Vivi Setiawaty, Hana Apsari Pawestri, Subangkit, Kartika Dewi Puspa, Arie Ardiansyah Nugraha, Hartanti Dian Ikawati, Nelly Puspandari, Krisna Nur Andriana Pangesti |
| EPI_ISL_1416191 | Fakultas Kedokteran Universitas Sumatera Utara | National Institute of Health Research and Development | Vivi Setiawaty, Hana Apsari Pawestri, Subangkit, Kartika Dewi Puspa, Arie Ardiansyah Nugraha, Hartanti Dian Ikawati, Nelly Puspandari, Krisna Nur Andriana Pangesti |
| EPI_ISL_1447329, EPI_ISL_1447330, EPI_ISL_1447331, EPI_ISL_1447332, EPI_ISL_1447333, EPI_ISL_1447334, EPI_ISL_1447335, EPI_ISL_1447336, EPI_ISL_1447337, EPI_ISL_1447338, EPI_ISL_1447339, EPI_ISL_1447340, EPI_ISL_1447341, EPI_ISL_1447342, EPI_ISL_1447343, EPI_ISL_1447344, EPI_ISL_1447345, EP EPI_ISL_1447349, EPI_ISL_1447350, EPI_ISL_1447351, EPI_ISL_1447352, EPI_ISL_1447353, EPI_ISL_1447354, EPI_ISL_1447355, EPI_ISL_1447356, EPI_ISL_1447357, EPI_ISL_1447358, EPI_ISL_1447359, EPI_ISL_1447360, EPI_ISL_1447361, EPI_ISL_1447362, EPI_ISL_1469232, EPI_ISL_1469233, EPI_ISL_1469234, EP | | | |
| see above | Diagnostic and Research Center of Infectious Diseases, Medical Faculty, Andalas University | Diagnostic and Research Center of Infectious Diseases, Medical Faculty, Andalas University | Andani Eka Putra, Syafrizayanti, Ikhwan R. Sudji, Linosefa, Dede Rahman Agustian, Ayu Novita Trisnawati, Siskalil Fahma, Nia Ayuni Putri, Sekar Asri Tresnaningtyas, SM Rezvi, Fauzul Azhim, Dessy Arisanty, Gestina Aliska, Desmawati, Nita Afriani ,Syandrez F Lailani |
| EPI_ISL_1469238 | National Institute Health Research and Development | National Institute of Health Research and Development | Reni Herman, Krisna Pangesti, Hana Apsari Pawestri, Subangkit, Kartika Dewi Puspa, Arie Ardiansyah Nugraha, Hartanti Dian Ikawati, Nelly Puspandari, Vivi Setiawaty |
| EPI_ISL_1469239 | BBTKL PP Jakarta | National Institute of Health Research and Development | Hartanti Dian Ikawati, Hana Apsari Pawestri, Subangkit, Kartika Dewi Puspa, Arie Ardiansyah Nugraha, Krisna Pangesti, Nelly Puspandari, Vivi Setiawaty |
| EPI_ISL_1469240 | Dinas Kesehatan Kabupaten Ponorogo | National Institute of Health Research and Development | Hartanti Dian Ikawati, Hana Apsari Pawestri, Subangkit, Kartika Dewi Puspa, Arie Ardiansyah Nugraha, Krisna Pangesti, Nelly Puspandari, Vivi Setiawaty |
| EPI_ISL_1469241, EPI_ISL_1469242 | National Institute Health Research and Development | National Institute of Health Research and Development | Subangkit, Hana Apsari Pawestri, Kartika Dewi Puspa, Arie Ardiansyah Nugraha, Hartanti Dian Ikawati, Krisna Nur Andriana Pangesti, Yuni Rukminiati, Ririn Ramadhany, Agustiningsih, Kindi Adam, Holy Arif Wibowo, Triyani Soekarso, Ni Ketut Susilarini, Nurika Ha Tati Febriyanti, Natalie Laurencia Kipuw, Fauzul Muna, Irene Lorinda Indalao, Nelly Puspandari, Vivi Setiawaty. |
| EPI_ISL_1469243, EPI_ISL_1469244 | National Institute Health Research and Development | National Institute of Health Research and Development | Reni Herman, Krisna Pangesti, Hana Apsari Pawestri, Subangkit, Kartika Dewi Puspa, Arie Ardiansyah Nugraha, Hartanti Dian Ikawati, Nelly Puspandari, Vivi Setiawaty |
| EPI_ISL_1469245 | National Institute Health Research and Development | National Institute of Health Research and Development | Subangkit, Hana Apsari Pawestri, Kartika Dewi Puspa, Arie Ardiansyah Nugraha, Hartanti Dian Ikawati, Krisna Nur Andriana Pangesti, Yuni Rukminiati, Ririn Ramadhany, Agustiningsih, Kindi Adam, Holy Arif Wibowo, Triyani Soekarso, Ni Ketut Susilarini, Nurika Ha Tati Febriyanti, Natalie Laurencia Kipuw, Fauzul Muna, Irene Lorinda Indalao, Nelly Puspandari, Vivi Setiawaty. |
| EPI_ISL_1469246 | Dinas Kesehatan Kabupaten Ponorogo | National Institute of Health Research and Development | Arie Ardiansyah Nugraha, Hana Apsari Pawestri, Subangkit, Kartika Dewi Puspa, Hartanti Dian Ikawati, Nelly Puspandari, Krisna Pangesti, Vivi Setiawaty |
| EPI_ISL_1469247 | RS Khusus Paru Karawang | National Institute of Health Research and Development | Hartanti Dian Ikawati, Hana Apsari Pawestri, Subangkit, Kartika Dewi Puspa, Arie Ardiansyah Nugraha, Krisna Pangesti, Nelly Puspandari, Vivi Setiawaty |
| EPI_ISL_1469248 | RS Mandaya Karawang | National Institute of Health Research and Development | Vivi Setiawaty, Hana Apsari Pawestri, Subangkit, Kartika Dewi Puspa, Arie Ardiansyah Nugraha, Hartanti Dian Ikawati, Nelly Puspandari, Krisna Pangesti |
| EPI_ISL_1469249 | Lab Intibios Karawang | National Institute of Health Research and Development | Hana Apsari Pawestri, Subangkit, Kartika Dewi Puspa, Arie Ardiansyah Nugraha, Hartanti Dian Ikawati, Nelly Puspandari, Krisna Pangesti, Vivi Setiawaty |
| EPI_ISL_1469250 | Dinas Kesehatan Blitar | National Institute of Health Research and Development | Hana Apsari Pawestri, Subangkit, Kartika Dewi Puspa, Arie Ardiansyah Nugraha, Hartanti Dian Ikawati, Nelly Puspandari, Krisna Pangesti, Vivi Setiawaty |
| EPI_ISL_1469251 | Lab Intibios Karawang | National Institute of Health Research and Development | Hana Apsari Pawestri, Subangkit, Kartika Dewi Puspa, Arie Ardiansyah Nugraha, Hartanti Dian Ikawati, Nelly Puspandari, Krisna Pangesti, Vivi Setiawaty |
| EPI_ISL_1469252 | National Institute Health Research and Development | National Institute of Health Research and Development | Subangkit, Hana Apsari Pawestri, Kartika Dewi Puspa, Arie Ardiansyah Nugraha, Hartanti Dian Ikawati, Krisna Nur Andriana Pangesti, Yuni Rukminiati, Ririn Ramadhany, Agustiningsih, Kindi Adam, Holy Arif Wibowo, Triyani Soekarso, Ni Ketut Susilarini, Nurika Ha Tati Febriyanti, Natalie Laurencia Kipuw, Fauzul Muna, Irene Lorinda Indalao, Nelly Puspandari, Vivi Setiawaty. |
| EPI_ISL_1469253 | RS Mandaya Karawang | National Institute of Health Research and Development | Vivi Setiawaty, Hana Apsari Pawestri, Subangkit, Kartika Dewi Puspa, Arie Ardiansyah Nugraha, Hartanti Dian Ikawati, Nelly Puspandari, Krisna Pangesti |
| EPI_ISL_1469254 | RS Lira Medika Karawang | National Institute of Health Research and Development | Vivi Setiawaty, Hana Apsari Pawestri, Subangkit, Kartika Dewi Puspa, Arie Ardiansyah Nugraha, Hartanti Dian Ikawati, Nelly Puspandari, Krisna Pangesti |
| EPI_ISL_1469255 | National Institute Health Research and Development | National Institute of Health Research and Development | Subangkit, Hana Apsari Pawestri, Kartika Dewi Puspa, Arie Ardiansyah Nugraha, Hartanti Dian Ikawati, Krisna Nur Andriana Pangesti, Yuni Rukminiati, Ririn Ramadhany, Agustiningsih, Kindi Adam, Holy Arif Wibowo, Triyani Soekarso, Ni Ketut Susilarini, Nurika Ha Tati Febriyanti, Natalie Laurencia Kipuw, Fauzul Muna, Irene Lorinda Indalao, Nelly Puspandari, Vivi Setiawaty. |
| EPI_ISL_1469256, EPI_ISL_1469257 | RS Khusus Paru Karawang | National Institute of Health Research and Development | Hartanti Dian Ikawati, Hana Apsari Pawestri, Subangkit, Kartika Dewi Puspa, Arie Ardiansyah Nugraha, Krisna Pangesti, Nelly Puspandari, Vivi Setiawaty |
| EPI_ISL_1469258 | PKM TEMPURAN KARAWANG | National Institute of Health Research and Development | Subangkit, Hana Apsari Pawestri, Kartika Dewi Puspa, Arie Ardiansyah Nugraha, Hartanti Dian Ikawati, Krisna Nur Andriana Pangesti, Yuni Rukminiati, Ririn Ramadhany, Agustiningsih, Kindi Adam, Holy Arif Wibowo, Triyani Soekarso, Ni Ketut Susilarini, Nurika Ha Tati Febriyanti, Natalie Laurencia Kipuw, Fauzul Muna, Irene Lorinda Indalao, Nelly Puspandari, Vivi Setiawaty. |
| EPI_ISL_1469259, EPI_ISL_1469260, EPI_ISL_1469261, EPI_ISL_1469262, EPI_ISL_1469263, EPI_ISL_1469264, EPI_ISL_1469265, EPI_ISL_1469266, EPI_ISL_1469267, EPI_ISL_1469268, EPI_ISL_1469269, EPI_ISL_1469270 | | | |
| see above | National Institute Health Research and Development | National Institute of Health Research and Development | Subangkit, Hana Apsari Pawestri, Kartika Dewi Puspa, Arie Ardiansyah Nugraha, Hartanti Dian Ikawati, Krisna Nur Andriana Pangesti, Yuni Rukminiati, Ririn Ramadhany, Agustiningsih, Kindi Adam, Holy Arif Wibowo, Triyani Soekarso, Ni Ketut Susilarini, Nurika Ha Tati Febriyanti, Natalie Laurencia Kipuw, Fauzul Muna, Irene Lorinda Indalao, Nelly Puspandari, Vivi Setiawaty. |

| | | | |
|---|---|---|---|
| EPI_ISL_1469271, EPI_ISL_1469272, EPI_ISL_1469273, EPI_ISL_1469274, EPI_ISL_1469275 | Dinas Kesehatan Kabupaten Ponorogo | National Institute of Health Research and Development | Arie Ardiansyah Nugraha, Hana Apsari Pawestri, Subangkit, Kartika Dewi Puspa, Hartanti Dian Ikawati, Nelly Puspandari, Krisna Pangesti, Vivi Setiawaty |
| EPI_ISL_1469276, EPI_ISL_1469277, EPI_ISL_1469278 | National Institute Health Research and Development | National Institute of Health Research and Development | Subangkit, Hana Apsari Pawestri, Kartika Dewi Puspa, Arie Ardiansyah Nugraha, Hartanti Dian Ikawati, Krisna Nur Andriana Pangesti, Yuni Rukminiati, Ririn Ramadhany, Agustiningsih, Kindi Adam, Holy Arif Wibowo, Triyani Soekarso, Ni Ketut Susilarini, Nurika Ha[illegible] Tati Febriyanti, Natalie Laurencia Kipuw, Fauzul Muna, Irene Lorinda Indalao, Nelly Puspandari, Vivi Setiawaty. |
| EPI_ISL_1469279, EPI_ISL_1469280, EPI_ISL_1469281, EPI_ISL_1469282, EPI_ISL_1469283 | RSUD Surakarta | National Institute of Health Research and Development | Kartika Dewi Puspa, Hana Apsari Pawestri, Subangkit, Arie Ardiansyah Nugraha, Hartanti Dian Ikawati, Nelly Puspandari, Krisna Pangesti, Vivi Setiawaty |
| EPI_ISL_1469284 | Dinas Kesehatan Kabupaten Ponorogo | National Institute of Health Research and Development | Kartika Dewi Puspa, Hana Apsari Pawestri, Subangkit, Arie Ardiansyah Nugraha, Hartanti Dian Ikawati, Nelly Puspandari, Krisna Pangesti, Vivi Setiawaty |
| EPI_ISL_1469285 | RS Khusus Paru Karawang | National Institute of Health Research and Development | Hartanti Dian Ikawati, Hana Apsari Pawestri, Subangkit, Kartika Dewi Puspa, Arie Ardiansyah Nugraha, Krisna Pangesti, Nelly Puspandari, Vivi Setiawaty |
| EPI_ISL_1469286 | RS Khusus Paru Karawang | National Institute of Health Research and Development | Vivi Setiawaty, Hana Apsari Pawestri, Subangkit, Kartika Dewi Puspa, Arie Ardiansyah Nugraha, Hartanti Dian Ikawati, Nelly Puspandari, Krisna Pangesti |
| EPI_ISL_1469287 | RS Mandaya Karawang | National Institute of Health Research and Development | Vivi Setiawaty, Hana Apsari Pawestri, Subangkit, Kartika Dewi Puspa, Arie Ardiansyah Nugraha, Hartanti Dian Ikawati, Nelly Puspandari, Krisna Pangesti |
| EPI_ISL_1469288, EPI_ISL_1469289 | Labkesda Karawang | National Institute of Health Research and Development | Hana Apsari Pawestri, Subangkit, Kartika Dewi Puspa, Arie Ardiansyah Nugraha, Hartanti Dian Ikawati, Nelly Puspandari, Krisna Pangesti, Vivi Setiawaty |
| EPI_ISL_1469290, EPI_ISL_1469291 | National Institute Health Research and Development | National Institute of Health Research and Development | Nur Ika Hariastuti, Nike Susanti, Hana Apsari Pawestri, Subangkit, Kartika Dewi Puspa, Arie Ardiansyah Nugraha, Hartanti Dian Ikawati, Nelly Puspandari, Vivi Setiawaty |
| EPI_ISL_1523640, EPI_ISL_1523641 | Medical Research Center, Faculty of Medicine, Syarif Hidayatullah State Islamic University Jakarta | Medical Research Center, Faculty of Medicine, Syarif Hidayatullah State Islamic University Jakarta | Erike A Suwarsono, Chris Adhiyanto, Laifa Hendarmin, Zeti Harriyati, Endah Wulandari, Flori R Sari, Fika Ekayanti, Hari Hendarto |
| EPI_ISL_1523642 | Medical Research Center, Faculty of Medicine, Syarif Hidayatullah State Islamic University Jakarta | Medical Research Center, Faculty of Medicine, Syarif Hidayatullah State Islamic University Jakarta | Erike A Suwarsono, Chris Adhiyanto, Laifa Hendarmin, Zeti Harriyati, Flori R Sari, Endah Wulandari, Fika Ekayanti, Hari Hendarto |
| EPI_ISL_1533034, EPI_ISL_1533035, EPI_ISL_1533036, EPI_ISL_1533037, EPI_ISL_1533038, EPI_ISL_1533039, EPI_ISL_1533040, EPI_ISL_1533041, EPI_ISL_1533042, EPI_ISL_1533043, EPI_ISL_1533044, EPI_ISL_1533045, EPI_ISL_1533046, EPI_ISL_1533047, EPI_ISL_1533048, EPI_ISL_1533049, EPI_ISL_1533050, EP[illegible] EPI_ISL_1533054, EPI_ISL_1533055, EPI_ISL_1533056, EPI_ISL_1533057, EPI_ISL_1533058, EPI_ISL_1533059, EPI_ISL_1533060, EPI_ISL_1533061, EPI_ISL_1533062, EPI_ISL_1533063, EPI_ISL_1533064, EPI_ISL_1533065, EPI_ISL_1533066, EPI_ISL_1533067, EPI_ISL_1533068, EPI_ISL_1533069, EPI_ISL_1533070, EP[illegible] EPI_ISL_1533074, EPI_ISL_1533075, EPI_ISL_1533076, EPI_ISL_1533077, EPI_ISL_1533078, EPI_ISL_1533079, EPI_ISL_1533080, EPI_ISL_1533081, EPI_ISL_1533082, EPI_ISL_1533083, EPI_ISL_1533084, EPI_ISL_1533085, EPI_ISL_1533086, EPI_ISL_1533087, EPI_ISL_1533088, EPI_ISL_1533089, EPI_ISL_1533090, EP[illegible] EPI_ISL_1533094, EPI_ISL_1533095, EPI_ISL_1533096, EPI_ISL_1533097, EPI_ISL_1533098, EPI_ISL_1533099, EPI_ISL_1533100, EPI_ISL_1533101 | | | |
| see above | Diagnostic and Research Center of Infectious Diseases, Medical Faculty, Andalas University | Diagnostic and Research Center of Infectious Diseases, Medical Faculty, Andalas University | Andani Eka Putra, Syafrizayanti, Ikhwan R. Sudji, Linosefa, Dede Rahman Agustian, Ayu Novita Trisnawati, Siskalil Fahma, Nia Ayuni Putri, Sekar Asri Tresnaningtyas, SM Rezvi, Fauzul Azhim, Dessy Arisanty, Gestina Aliska, Desmawati, Nita Afriani ,Syandrez [illegible] Lailani |
| EPI_ISL_1534690, EPI_ISL_1534691, EPI_ISL_1534692, EPI_ISL_1534693, EPI_ISL_1534694, EPI_ISL_1534695, EPI_ISL_1534696 | BTKLPP kelas 1 Batam | National Institute of Health Research and Development | Hana Apsari Pawestri, Subangkit, Kartika Dewi Puspa, Arie Ardiansyah Nugraha, Hartanti Dian Ikawati, Nelly Puspandari, Krisna Pangesti, Budi Santosa, Ismail, Ida Neni Haryanti, Vivi Setiawaty |
| EPI_ISL_1534697, EPI_ISL_1534698, EPI_ISL_1534699 | Ciputra Hospital Citra Garden City | National Institute of Health Research and Development | Vivi Setiawaty, Hana Apsari Pawestri, Subangkit, Kartika Dewi Puspa, Arie Ardiansyah Nugraha, Hartanti Dian Ikawati, Nelly Puspandari, Krisna Pangesti |
| EPI_ISL_1534700 | Ciputra Hospital Citra Garden City | National Institute of Health Research and Development | Hartanti Dian Ikawati, Hana Apsari Pawestri, Subangkit, Kartika Dewi Puspa, Arie Ardiansyah Nugraha, Krisna Pangesti, Nelly Puspandari, Vivi Setiawaty |
| EPI_ISL_1534701, EPI_ISL_1534702 | Fakultas kedokteran Universitas Sumatra Utara | National Institute of Health Research and Development | Hartanti Dian Ikawati, Hana Apsari Pawestri, Subangkit, Kartika Dewi Puspa, Arie Ardiansyah Nugraha, Krisna Pangesti, Nelly Puspandari, Vivi Setiawaty |
| EPI_ISL_1534703 | Fakultas kedokteran Universitas Sumatra Utara | National Institute of Health Research and Development | Kartika Dewi Puspa, Hana Apsari Pawestri, Subangkit, Arie Ardiansyah Nugraha, Hartanti Dian Ikawati, Nelly Puspandari, Krisna Pangesti, Vivi Setiawaty |
| EPI_ISL_1534704, EPI_ISL_1534705 | BBLK Surabaya | National Institute of Health Research and Development | Kartika Dewi Puspa, Hana Apsari Pawestri, Subangkit, Arie Ardiansyah Nugraha, Hartanti Dian Ikawati, Nelly Puspandari, Krisna Pangesti, Vivi Setiawaty |
| EPI_ISL_1534706, EPI_ISL_1534707 | National Institute Health Research and Development | National Institute of Health Research and Development | Subangkit, Hana Apsari Pawestri, Kartika Dewi Puspa, Arie Ardiansyah Nugraha, Hartanti Dian Ikawati, Krisna Nur Andriana Pangesti, Yuni Rukminiati, Ririn Ramadhany, Agustiningsih, Kindi Adam, Holy Arif Wibowo, Triyani Soekarso, Ni Ketut Susilarini, Nurika Ha[illegible] Tati Febriyanti, Natalie Laurencia Kipuw, Fauzul Muna, Irene Lorinda Indalao, Nelly Puspandari, Vivi Setiawaty. |
| EPI_ISL_1534708 | Labkesda Karawang | National Institute of Health Research and Development | Arie Ardiansyah Nugraha, Hana Apsari Pawestri, Subangkit, Kartika Dewi Puspa, Hartanti Dian Ikawati, Nelly Puspandari, Krisna Pangesti, Vivi Setiawaty |
| EPI_ISL_1615586 | Santosa Kopo Hospital | West Java Health Laboratory; School of Life Sciences and Technology, Institut Teknologi Bandung | Azzania Fibriani, Ema Rahmawati, Ryan Bayusantika Ristandi, Rifky Waluyajati Rachman, Cut Nur Cinthia Alamanda, Rini Robiani, Miftahul Faridl, Karimatu Khoirunnisa, Aulia Saraswati Wicaksono |

| | | | |
|---|---|---|---|
| EPI_ISL_1615587 | Bandung Public Health | West Java Health Laboratory; School of Life Sciences and Technology, Institut Teknologi Bandung | Azzania Fibriani, Ema Rahmawati, Ryan Bayusantika Ristandi, Rifky Waluyajati Rachman, Cut Nur Cinthia Alamanda, Rini Robiani, Miftahul Faridl, Karimatu Khoirunnisa, Aulia Saraswati Wicaksono |
| EPI_ISL_1615588 | Mitra Kasih Hospital | West Java Health Laboratory; School of Life Sciences and Technology, Institut Teknologi Bandung | Azzania Fibriani, Ema Rahmawati, Ryan Bayusantika Ristandi, Rifky Waluyajati Rachman, Cut Nur Cinthia Alamanda, Rini Robiani, Miftahul Faridl, Karimatu Khoirunnisa, Aulia Saraswati Wicaksono |
| EPI_ISL_1615589 | Sumedang Public Health | West Java Health Laboratory; School of Life Sciences and Technology, Institut Teknologi Bandung | Azzania Fibriani, Ema Rahmawati, Ryan Bayusantika Ristandi, Rifky Waluyajati Rachman, Cut Nur Cinthia Alamanda, Rini Robiani, Miftahul Faridl, Karimatu Khoirunnisa, Aulia Saraswati Wicaksono |
| EPI_ISL_1615590 | Subang Public Health | West Java Health Laboratory; School of Life Sciences and Technology, Institut Teknologi Bandung | Azzania Fibriani, Ema Rahmawati, Ryan Bayusantika Ristandi, Rifky Waluyajati Rachman, Cut Nur Cinthia Alamanda, Rini Robiani, Miftahul Faridl, Karimatu Khoirunnisa, Aulia Saraswati Wicaksono |
| EPI_ISL_1615600 | West Java Health Laboratory | West Java Health Laboratory; School of Life Sciences and Technology, Institut Teknologi Bandung | Azzania Fibriani, Ema Rahmawati, Ryan Bayusantika Ristandi, Rifky Waluyajati Rachman, Cut Nur Cinthia Alamanda, Rini Robiani, Miftahul Faridl, Karimatu Khoirunnisa, Aulia Saraswati Wicaksono |
| EPI_ISL_1615617 | Kuningan Public Health | West Java Health Laboratory; School of Life Sciences and Technology, Institut Teknologi Bandung | Azzania Fibriani, Ema Rahmawati, Ryan Bayusantika Ristandi, Rifky Waluyajati Rachman, Cut Nur Cinthia Alamanda, Rini Robiani, Miftahul Faridl, Karimatu Khoirunnisa, Aulia Saraswati Wicaksono |
| EPI_ISL_1615620 | Subang Public Health | West Java Health Laboratory; School of Life Sciences and Technology, Institut Teknologi Bandung | Azzania Fibriani, Ema Rahmawati, Ryan Bayusantika Ristandi, Rifky Waluyajati Rachman, Cut Nur Cinthia Alamanda, Rini Robiani, Miftahul Faridl, Karimatu Khoirunnisa, Aulia Saraswati Wicaksono |
| EPI_ISL_1615621 | Sumedang Public Health | West Java Health Laboratory; School of Life Sciences and Technology, Institut Teknologi Bandung | Azzania Fibriani, Ema Rahmawati, Ryan Bayusantika Ristandi, Rifky Waluyajati Rachman, Cut Nur Cinthia Alamanda, Rini Robiani, Miftahul Faridl, Karimatu Khoirunnisa, Aulia Saraswati Wicaksono |
| EPI_ISL_1615656 | Bandung Public Health | West Java Health Laboratory; School of Life Sciences and Technology, Institut Teknologi Bandung | Azzania Fibriani, Ema Rahmawati, Ryan Bayusantika Ristandi, Rifky Waluyajati Rachman, Cut Nur Cinthia Alamanda, Rini Robiani, Miftahul Faridl, Karimatu Khoirunnisa, Aulia Saraswati Wicaksono |
| EPI_ISL_1615658 | Santo Borromeus Hospital | West Java Health Laboratory; School of Life Sciences and Technology, Institut Teknologi Bandung | Azzania Fibriani, Ema Rahmawati, Ryan Bayusantika Ristandi, Rifky Waluyajati Rachman, Cut Nur Cinthia Alamanda, Rini Robiani, Miftahul Faridl, Karimatu Khoirunnisa, Aulia Saraswati Wicaksono |
| EPI_ISL_1615660 | West Java Health Laboratory | West Java Health Laboratory; School of Life Sciences and Technology, Institut Teknologi Bandung | Azzania Fibriani, Ema Rahmawati, Ryan Bayusantika Ristandi, Rifky Waluyajati Rachman, Cut Nur Cinthia Alamanda, Rini Robiani, Miftahul Faridl, Karimatu Khoirunnisa, Aulia Saraswati Wicaksono |
| EPI_ISL_1622360, EPI_ISL_1622361 | RS Mitra Keluarga Gading Serpong | Eijkman Institute for Molecular Biology, Ministry of Research and Technology/National Agency for Research and Innovation | Willy Agustine, Hidayat Trimarsanto, Lydia V. Panggalo, Iskandar Adnan, Sukma Oktavianthi, Edison Johar, Frilasita A Yudhaputri, Muhammad Rezki Rasyak, Safarina G Malik, Khin Saw Myint, Amin Soeban |
| EPI_ISL_1622362 | RS Sentra Medika Cikarang | Eijkman Institute for Molecular Biology, Ministry of Research and Technology/National Agency for Research and Innovation | Willy Agustine, Hidayat Trimarsanto, Lydia V. Panggalo, Iskandar Adnan, Sukma Oktavianthi, Edison Johar, Frilasita A Yudhaputri, Muhammad Rezki Rasyak, Safarina G Malik, Khin Saw Myint, Amin Soeban |
| EPI_ISL_1622363, EPI_ISL_1622364 | RS Sari Asih Ciledug | Eijkman Institute for Molecular Biology, Ministry of Research and Technology/National Agency for Research and Innovation | Willy Agustine, Hidayat Trimarsanto, Lydia V. Panggalo, Iskandar Adnan, Sukma Oktavianthi, Edison Johar, Frilasita A Yudhaputri, Muhammad Rezki Rasyak, Safarina G Malik, Khin Saw Myint, Amin Soeban |
| EPI_ISL_1622365 | RS Bhakti Husada Cikarang | Eijkman Institute for Molecular Biology, Ministry of Research and Technology/National Agency for Research and Innovation | Willy Agustine, Hidayat Trimarsanto, Lydia V. Panggalo, Iskandar Adnan, Sukma Oktavianthi, Edison Johar, Frilasita A Yudhaputri, Muhammad Rezki Rasyak, Safarina G Malik, Khin Saw Myint, Amin Soeban |
| EPI_ISL_1622366 | RSUD Cileungsi | Eijkman Institute for Molecular Biology, Ministry of Research and Technology/National Agency for Research and Innovation | Willy Agustine, Hidayat Trimarsanto, Lydia V. Panggalo, Iskandar Adnan, Sukma Oktavianthi, Edison Johar, Frilasita A Yudhaputri, Muhammad Rezki Rasyak, Safarina G Malik, Khin Saw Myint, Amin Soeban |
| EPI_ISL_1622367 | RS Harapan Bunda | Eijkman Institute for Molecular Biology, Ministry of Research and | Willy Agustine, Hidayat Trimarsanto, Lydia V. Panggalo, Iskandar Adnan, Sukma Oktavianthi, Edison Johar, Frilasita A Yudhaputri, Muhammad Rezki Rasyak, Safarina G Malik, Khin Saw Myint, Amin Soeban |

| | | | |
|---|---|---|---|
| | | Technology/National Agency for Research and Innovation | |
| EPI_ISL_1622368 | RSJ Dr. Soeharto Heerdjan | Eijkman Institute for Molecular Biology, Ministry of Research and Technology/National Agency for Research and Innovation | Willy Agustine, Hidayat Trimarsanto, Lydia V. Panggalo, Iskandar Adnan, Sukma Oktavianthi, Edison Johar, Frilasita A Yudhaputri, Muhammad Rezki Rasyak, Safarina G Malik, Khin Saw Myint, Amin Soeban[illegible] |
| EPI_ISL_1622369 | RS Sumber Waras | Eijkman Institute for Molecular Biology, Ministry of Research and Technology/National Agency for Research and Innovation | Willy Agustine, Hidayat Trimarsanto, Lydia V. Panggalo, Iskandar Adnan, Sukma Oktavianthi, Edison Johar, Frilasita A Yudhaputri, Muhammad Rezki Rasyak, Safarina G Malik, Khin Saw Myint, Amin Soeban[illegible] |
| EPI_ISL_1622370 | RSIA Selasih Medika | Eijkman Institute for Molecular Biology, Ministry of Research and Technology/National Agency for Research and Innovation | Willy Agustine, Hidayat Trimarsanto, Lydia V. Panggalo, Iskandar Adnan, Sukma Oktavianthi, Edison Johar, Frilasita A Yudhaputri, Muhammad Rezki Rasyak, Safarina G Malik, Khin Saw Myint, Amin Soeban[illegible] |
| EPI_ISL_1622371 | RS Mitra Keluarga Gading Serpong | Eijkman Institute for Molecular Biology, Ministry of Research and Technology/National Agency for Research and Innovation | Willy Agustine, Hidayat Trimarsanto, Lydia V. Panggalo, Iskandar Adnan, Sukma Oktavianthi, Edison Johar, Frilasita A Yudhaputri, Muhammad Rezki Rasyak, Safarina G Malik, Khin Saw Myint, Amin Soeban[illegible] |
| EPI_ISL_1622372 | RS Mitra Keluarga Gading Serpong | Eijkman Institute for Molecular Biology, Ministry of Research and Technology/National Agency for Research and Innovation | Hidayat Trimarsanto, Lydia V. Panggalo, Iskandar Adnan, Sukma Oktavianthi, Edison Johar, Frilasita A Yudhaputri, Muhammad Rezki Rasyak, Willy Agustine, Safarina G Malik, Khin Saw Myint, Amin Soeban[illegible] |
| EPI_ISL_1622373 | RS Mary Cileungsi Hijau | Eijkman Institute for Molecular Biology, Ministry of Research and Technology/National Agency for Research and Innovation | Hidayat Trimarsanto, Lydia V. Panggalo, Iskandar Adnan, Sukma Oktavianthi, Edison Johar, Frilasita A Yudhaputri, Muhammad Rezki Rasyak, Willy Agustine, Safarina G Malik, Khin Saw Myint, Amin Soeban[illegible] |
| EPI_ISL_1622374 | RS Sentra Medika Cikarang | Eijkman Institute for Molecular Biology, Ministry of Research and Technology/National Agency for Research and Innovation | Hidayat Trimarsanto, Lydia V. Panggalo, Iskandar Adnan, Sukma Oktavianthi, Edison Johar, Frilasita A Yudhaputri, Muhammad Rezki Rasyak, Willy Agustine, Safarina G Malik, Khin Saw Myint, Amin Soeban[illegible] |
| EPI_ISL_1622375 | RS Taman Harapan Baru | Eijkman Institute for Molecular Biology, Ministry of Research and Technology/National Agency for Research and Innovation | Hidayat Trimarsanto, Lydia V. Panggalo, Iskandar Adnan, Sukma Oktavianthi, Edison Johar, Frilasita A Yudhaputri, Muhammad Rezki Rasyak, Willy Agustine, Safarina G Malik, Khin Saw Myint, Amin Soeban[illegible] |
| EPI_ISL_1622376, EPI_ISL_1622377 | RSAB Harapan Kita | Eijkman Institute for Molecular Biology, Ministry of Research and Technology/National Agency for Research and Innovation | Hidayat Trimarsanto, Lydia V. Panggalo, Iskandar Adnan, Sukma Oktavianthi, Edison Johar, Frilasita A Yudhaputri, Muhammad Rezki Rasyak, Willy Agustine, Safarina G Malik, Khin Saw Myint, Amin Soeban[illegible] |
| EPI_ISL_1622378 | RSIA Bun | Eijkman Institute for Molecular Biology, Ministry of Research and Technology/National Agency for Research and Innovation | Hidayat Trimarsanto, Lydia V. Panggalo, Iskandar Adnan, Sukma Oktavianthi, Edison Johar, Frilasita A Yudhaputri, Muhammad Rezki Rasyak, Willy Agustine, Safarina G Malik, Khin Saw Myint, Amin Soeban[illegible] |
| EPI_ISL_1622379, EPI_ISL_1622380 | RS Mitra Keluarga Gading Serpong | Eijkman Institute for Molecular Biology, Ministry of Research and Technology/National Agency for Research and Innovation | Hidayat Trimarsanto, Lydia V. Panggalo, Iskandar Adnan, Sukma Oktavianthi, Edison Johar, Frilasita A Yudhaputri, Muhammad Rezki Rasyak, Willy Agustine, Safarina G Malik, Khin Saw Myint, Amin Soeban[illegible] |
| EPI_ISL_1622381 | RS Bhakti Husada Cikarang | Eijkman Institute for Molecular Biology, Ministry of Research and Technology/National Agency for Research and Innovation | Hidayat Trimarsanto, Lydia V. Panggalo, Iskandar Adnan, Sukma Oktavianthi, Edison Johar, Frilasita A Yudhaputri, Muhammad Rezki Rasyak, Willy Agustine, Safarina G Malik, Khin Saw Myint, Amin Soeban[illegible] |
| EPI_ISL_1622382 | RS Medika Dramaga | Eijkman Institute for Molecular Biology, Ministry of Research and Technology/National Agency for Research and Innovation | Hidayat Trimarsanto, Lydia V. Panggalo, Iskandar Adnan, Sukma Oktavianthi, Edison Johar, Frilasita A Yudhaputri, Muhammad Rezki Rasyak, Willy Agustine, Safarina G Malik, Khin Saw Myint, Amin Soeban[illegible] |

| | | | |
|---|---|---|---|
| EPI_ISL_1622383 | RS Bina Sehat Mandiri | Eijkman Institute for Molecular Biology, Ministry of Research and Technology/National Agency for Research and Innovation | Hidayat Trimarsanto, Lydia V. Panggalo, Iskandar Adnan, Sukma Oktavianthi, Edison Johar, Frilasita A Yudhaputri, Muhammad Rezki Rasyak, Willy Agustine, Safarina G Malik, Khin Saw Myint, Amin Soeband |
| EPI_ISL_1622384, EPI_ISL_1622385 | RS Harapan Bunda | Eijkman Institute for Molecular Biology, Ministry of Research and Technology/National Agency for Research and Innovation | Lydia V. Panggalo, Iskandar Adnan, Sukma Oktavianthi, Edison Johar, Frilasita A Yudhaputri, Muhammad Rezki Rasyak, Willy Agustine, Hidayat Trimarsanto, Safarina G Malik, Khin Saw Myint, Amin Soeband |
| EPI_ISL_1622386 | RSJ Dr. Soeharto Heerdjan | Eijkman Institute for Molecular Biology, Ministry of Research and Technology/National Agency for Research and Innovation | Lydia V. Panggalo, Iskandar Adnan, Sukma Oktavianthi, Edison Johar, Frilasita A Yudhaputri, Muhammad Rezki Rasyak, Willy Agustine, Hidayat Trimarsanto, Safarina G Malik, Khin Saw Myint, Amin Soeband |
| EPI_ISL_1622387 | RSU HGA Depok | Eijkman Institute for Molecular Biology, Ministry of Research and Technology/National Agency for Research and Innovation | Lydia V. Panggalo, Iskandar Adnan, Sukma Oktavianthi, Edison Johar, Frilasita A Yudhaputri, Muhammad Rezki Rasyak, Willy Agustine, Hidayat Trimarsanto, Safarina G Malik, Khin Saw Myint, Amin Soeband |
| EPI_ISL_1622388 | RS Mitra Keluarga Gading Serpong | Eijkman Institute for Molecular Biology, Ministry of Research and Technology/National Agency for Research and Innovation | Lydia V. Panggalo, Iskandar Adnan, Sukma Oktavianthi, Edison Johar, Frilasita A Yudhaputri, Muhammad Rezki Rasyak, Willy Agustine, Hidayat Trimarsanto, Safarina G Malik, Khin Saw Myint, Amin Soeband |
| EPI_ISL_1622389 | RS Masmitra | Eijkman Institute for Molecular Biology, Ministry of Research and Technology/National Agency for Research and Innovation | Lydia V. Panggalo, Iskandar Adnan, Sukma Oktavianthi, Edison Johar, Frilasita A Yudhaputri, Muhammad Rezki Rasyak, Willy Agustine, Hidayat Trimarsanto, Safarina G Malik, Khin Saw Myint, Amin Soeband |
| EPI_ISL_1622390 | RS Sentra Medika Cikarang | Eijkman Institute for Molecular Biology, Ministry of Research and Technology/National Agency for Research and Innovation | Lydia V. Panggalo, Iskandar Adnan, Sukma Oktavianthi, Edison Johar, Frilasita A Yudhaputri, Muhammad Rezki Rasyak, Willy Agustine, Hidayat Trimarsanto, Safarina G Malik, Khin Saw Myint, Amin Soeband |
| EPI_ISL_1622391 | RS Medirossa Cikarang | Eijkman Institute for Molecular Biology, Ministry of Research and Technology/National Agency for Research and Innovation | Lydia V. Panggalo, Iskandar Adnan, Sukma Oktavianthi, Edison Johar, Frilasita A Yudhaputri, Muhammad Rezki Rasyak, Willy Agustine, Hidayat Trimarsanto, Safarina G Malik, Khin Saw Myint, Amin Soeband |
| EPI_ISL_1622392 | RSIA Kenari Graha Medika | Eijkman Institute for Molecular Biology, Ministry of Research and Technology/National Agency for Research and Innovation | Iskandar Adnan, Sukma Oktavianthi, Edison Johar, Frilasita A Yudhaputri, Muhammad Rezki Rasyak, Willy Agustine, Hidayat Trimarsanto, Lydia V. Panggalo, Safarina G Malik, Khin Saw Myint, Amin Soeband |
| EPI_ISL_1622393 | RSJ Dr. Soeharto Heerdjan | Eijkman Institute for Molecular Biology, Ministry of Research and Technology/National Agency for Research and Innovation | Iskandar Adnan, Sukma Oktavianthi, Edison Johar, Frilasita A Yudhaputri, Muhammad Rezki Rasyak, Willy Agustine, Hidayat Trimarsanto, Lydia V. Panggalo, Safarina G Malik, Khin Saw Myint, Amin Soeband |
| EPI_ISL_1622394 | RSU Proklamasi | Eijkman Institute for Molecular Biology, Ministry of Research and Technology/National Agency for Research and Innovation | Iskandar Adnan, Sukma Oktavianthi, Edison Johar, Frilasita A Yudhaputri, Muhammad Rezki Rasyak, Willy Agustine, Hidayat Trimarsanto, Lydia V. Panggalo, Safarina G Malik, Khin Saw Myint, Amin Soeband |
| EPI_ISL_1622395, EPI_ISL_1622396 | RSUD Cileungsi | Eijkman Institute for Molecular Biology, Ministry of Research and Technology/National Agency for Research and Innovation | Iskandar Adnan, Sukma Oktavianthi, Edison Johar, Frilasita A Yudhaputri, Muhammad Rezki Rasyak, Willy Agustine, Hidayat Trimarsanto, Lydia V. Panggalo, Safarina G Malik, Khin Saw Myint, Amin Soeband |
| EPI_ISL_1622397, EPI_ISL_1622398, EPI_ISL_1622399 | RSAB Harapan Kita | Eijkman Institute for Molecular Biology, Ministry of Research and Technology/National Agency for Research and Innovation | Iskandar Adnan, Sukma Oktavianthi, Edison Johar, Frilasita A Yudhaputri, Muhammad Rezki Rasyak, Willy Agustine, Hidayat Trimarsanto, Lydia V. Panggalo, Safarina G Malik, Khin Saw Myint, Amin Soeband |
| EPI_ISL_1622400 | RSU HGA Depok | Eijkman Institute for Molecular Biology, Ministry of Research and | Iskandar Adnan, Sukma Oktavianthi, Edison Johar, Frilasita A Yudhaputri, Muhammad Rezki Rasyak, Willy Agustine, Hidayat Trimarsanto, Lydia V. Panggalo, Safarina G Malik, Khin Saw Myint, Amin Soeband |

| | | | |
|---|---|---|---|
| | | Technology/National Agency for Research and Innovation | |
| EPI_ISL_1622401 | RSU Proklamasi | Eijkman Institute for Molecular Biology, Ministry of Research and Technology/National Agency for Research and Innovation | Iskandar Adnan, Sukma Oktavianthi, Edison Johar, Frilasita A Yudhaputri, Muhammad Rezki Rasyak, Willy Agustine, Hidayat Trimarsanto, Lydia V. Panggalo, Safarina G Malik, Khin Saw Myint, Amin Soeban[illegible] |
| EPI_ISL_1622402 | RSU Proklamasi | Eijkman Institute for Molecular Biology, Ministry of Research and Technology/National Agency for Research and Innovation | Sukma Oktavianthi, Edison Johar, Frilasita A Yudhaputri, Muhammad Rezki Rasyak, Willy Agustine, Hidayat Trimarsanto, Lydia V. Panggalo, Iskandar Adnan, Safarina G Malik, Khin Saw Myint, Amin Soeban[illegible] |
| EPI_ISL_1622403, EPI_ISL_1622404 | RS Bhakti Husada Cikarang | Eijkman Institute for Molecular Biology, Ministry of Research and Technology/National Agency for Research and Innovation | Sukma Oktavianthi, Edison Johar, Frilasita A Yudhaputri, Muhammad Rezki Rasyak, Willy Agustine, Hidayat Trimarsanto, Lydia V. Panggalo, Iskandar Adnan, Safarina G Malik, Khin Saw Myint, Amin Soeban[illegible] |
| EPI_ISL_1622405, EPI_ISL_1622406, EPI_ISL_1622407 | RSUD Cileungsi | Eijkman Institute for Molecular Biology, Ministry of Research and Technology/National Agency for Research and Innovation | Sukma Oktavianthi, Edison Johar, Frilasita A Yudhaputri, Muhammad Rezki Rasyak, Willy Agustine, Hidayat Trimarsanto, Lydia V. Panggalo, Iskandar Adnan, Safarina G Malik, Khin Saw Myint, Amin Soeban[illegible] |
| EPI_ISL_1622408 | RS Medirossa Cikarang | Eijkman Institute for Molecular Biology, Ministry of Research and Technology/National Agency for Research and Innovation | Edison Johar, Frilasita A Yudhaputri, Muhammad Rezki Rasyak, Willy Agustine, Hidayat Trimarsanto, Lydia V. Panggalo, Iskandar Adnan, Sukma Oktavianthi, Safarina G Malik, Khin Saw Myint, Amin Soeban[illegible] |
| EPI_ISL_1622409 | RS Mitra Keluarga Kalideres | Eijkman Institute for Molecular Biology, Ministry of Research and Technology/National Agency for Research and Innovation | Edison Johar, Frilasita A Yudhaputri, Muhammad Rezki Rasyak, Willy Agustine, Hidayat Trimarsanto, Lydia V. Panggalo, Iskandar Adnan, Sukma Oktavianthi, Safarina G Malik, Khin Saw Myint, Amin Soeban[illegible] |
| EPI_ISL_1622410 | RS Cendana | Eijkman Institute for Molecular Biology, Ministry of Research and Technology/National Agency for Research and Innovation | Edison Johar, Frilasita A Yudhaputri, Muhammad Rezki Rasyak, Willy Agustine, Hidayat Trimarsanto, Lydia V. Panggalo, Iskandar Adnan, Sukma Oktavianthi, Safarina G Malik, Khin Saw Myint, Amin Soeban[illegible] |
| EPI_ISL_1622411 | EKA Hospital BSD | Eijkman Institute for Molecular Biology, Ministry of Research and Technology/National Agency for Research and Innovation | Edison Johar, Frilasita A Yudhaputri, Muhammad Rezki Rasyak, Willy Agustine, Hidayat Trimarsanto, Lydia V. Panggalo, Iskandar Adnan, Sukma Oktavianthi, Safarina G Malik, Khin Saw Myint, Amin Soeban[illegible] |
| EPI_ISL_1622412 | RS Medirossa Cikarang | Eijkman Institute for Molecular Biology, Ministry of Research and Technology/National Agency for Research and Innovation | Edison Johar, Frilasita A Yudhaputri, Muhammad Rezki Rasyak, Willy Agustine, Hidayat Trimarsanto, Lydia V. Panggalo, Iskandar Adnan, Sukma Oktavianthi, Safarina G Malik, Khin Saw Myint, Amin Soeban[illegible] |
| EPI_ISL_1622413 | RS Mitra Keluarga Gading Serpong | Eijkman Institute for Molecular Biology, Ministry of Research and Technology/National Agency for Research and Innovation | Edison Johar, Frilasita A Yudhaputri, Muhammad Rezki Rasyak, Willy Agustine, Hidayat Trimarsanto, Lydia V. Panggalo, Iskandar Adnan, Sukma Oktavianthi, Safarina G Malik, Khin Saw Myint, Amin Soeban[illegible] |
| EPI_ISL_1622414 | RS Bhakti Mulia | Eijkman Institute for Molecular Biology, Ministry of Research and Technology/National Agency for Research and Innovation | Edison Johar, Frilasita A Yudhaputri, Muhammad Rezki Rasyak, Willy Agustine, Hidayat Trimarsanto, Lydia V. Panggalo, Iskandar Adnan, Sukma Oktavianthi, Safarina G Malik, Khin Saw Myint, Amin Soeban[illegible] |
| EPI_ISL_1622415 | RSU HGA Depok | Eijkman Institute for Molecular Biology, Ministry of Research and Technology/National Agency for Research and Innovation | Edison Johar, Frilasita A Yudhaputri, Muhammad Rezki Rasyak, Willy Agustine, Hidayat Trimarsanto, Lydia V. Panggalo, Iskandar Adnan, Sukma Oktavianthi, Safarina G Malik, Khin Saw Myint, Amin Soeban[illegible] |
| EPI_ISL_1622416, EPI_ISL_1622417 | RS Permata Cibubur | Eijkman Institute for Molecular Biology, Ministry of Research and Technology/National Agency for Research and Innovation | Frilasita A Yudhaputri, Muhammad Rezki Rasyak, Willy Agustine, Hidayat Trimarsanto, Lydia V. Panggalo, Iskandar Adnan, Sukma Oktavianthi, Edison Johar, Safarina G Malik, Khin Saw Myint, Amin Soeban[illegible] |

| | | | |
|---|---|---|---|
| EPI_ISL_1622418 | RS Qadr | Eijkman Institute for Molecular Biology, Ministry of Research and Technology/National Agency for Research and Innovation | Frilasita A Yudhaputri, Muhammad Rezki Rasyak, Willy Agustine, Hidayat Trimarsanto, Lydia V. Panggalo, Iskandar Adnan, Sukma Oktavianthi, Edison Johar, Safarina G Malik, Khin Saw Myint, Amin Soeban |
| EPI_ISL_1622419, EPI_ISL_1622420 | RS Sentra Medika Cikarang | Eijkman Institute for Molecular Biology, Ministry of Research and Technology/National Agency for Research and Innovation | Frilasita A Yudhaputri, Muhammad Rezki Rasyak, Willy Agustine, Hidayat Trimarsanto, Lydia V. Panggalo, Iskandar Adnan, Sukma Oktavianthi, Edison Johar, Safarina G Malik, Khin Saw Myint, Amin Soeban |
| EPI_ISL_1622421, EPI_ISL_1622422 | RS Medirossa Cikarang | Eijkman Institute for Molecular Biology, Ministry of Research and Technology/National Agency for Research and Innovation | Frilasita A Yudhaputri, Muhammad Rezki Rasyak, Willy Agustine, Hidayat Trimarsanto, Lydia V. Panggalo, Iskandar Adnan, Sukma Oktavianthi, Edison Johar, Safarina G Malik, Khin Saw Myint, Amin Soeban |
| EPI_ISL_1622423 | RSJ Dr. Soeharto Heerdjan | Eijkman Institute for Molecular Biology, Ministry of Research and Technology/National Agency for Research and Innovation | Frilasita A Yudhaputri, Muhammad Rezki Rasyak, Willy Agustine, Hidayat Trimarsanto, Lydia V. Panggalo, Iskandar Adnan, Sukma Oktavianthi, Edison Johar, Safarina G Malik, Khin Saw Myint, Amin Soeban |
| EPI_ISL_1622424 | RS Sumber Waras | Eijkman Institute for Molecular Biology, Ministry of Research and Technology/National Agency for Research and Innovation | Muhammad Rezki Rasyak, Willy Agustine, Hidayat Trimarsanto, Lydia V. Panggalo, Iskandar Adnan, Sukma Oktavianthi, Edison Johar, Frilasita A Yudhaputri, Safarina G Malik, Khin Saw Myint, Amin Soeban |
| EPI_ISL_1622425 | RS Mitra Keluarga Kelapa Gading | Eijkman Institute for Molecular Biology, Ministry of Research and Technology/National Agency for Research and Innovation | Muhammad Rezki Rasyak, Willy Agustine, Hidayat Trimarsanto, Lydia V. Panggalo, Iskandar Adnan, Sukma Oktavianthi, Edison Johar, Frilasita A Yudhaputri, Safarina G Malik, Khin Saw Myint, Amin Soeban |
| EPI_ISL_1622426 | RS Mary Cileungsi Hijau | Eijkman Institute for Molecular Biology, Ministry of Research and Technology/National Agency for Research and Innovation | Muhammad Rezki Rasyak, Willy Agustine, Hidayat Trimarsanto, Lydia V. Panggalo, Iskandar Adnan, Sukma Oktavianthi, Edison Johar, Frilasita A Yudhaputri, Safarina G Malik, Khin Saw Myint, Amin Soeban |
| EPI_ISL_1622427 | RS OMNI Pulomas | Eijkman Institute for Molecular Biology, Ministry of Research and Technology/National Agency for Research and Innovation | Muhammad Rezki Rasyak, Willy Agustine, Hidayat Trimarsanto, Lydia V. Panggalo, Iskandar Adnan, Sukma Oktavianthi, Edison Johar, Frilasita A Yudhaputri, Safarina G Malik, Khin Saw Myint, Amin Soeban |
| EPI_ISL_1622428, EPI_ISL_1622429, EPI_ISL_1622430 | RS Mitra Keluarga Gading Serpong | Eijkman Institute for Molecular Biology, Ministry of Research and Technology/National Agency for Research and Innovation | Muhammad Rezki Rasyak, Willy Agustine, Hidayat Trimarsanto, Lydia V. Panggalo, Iskandar Adnan, Sukma Oktavianthi, Edison Johar, Frilasita A Yudhaputri, Safarina G Malik, Khin Saw Myint, Amin Soeban |
| EPI_ISL_1622431, EPI_ISL_1622432 | RS Sentra Medika Cikarang | Eijkman Institute for Molecular Biology, Ministry of Research and Technology/National Agency for Research and Innovation | Muhammad Rezki Rasyak, Willy Agustine, Hidayat Trimarsanto, Lydia V. Panggalo, Iskandar Adnan, Sukma Oktavianthi, Edison Johar, Frilasita A Yudhaputri, Safarina G Malik, Khin Saw Myint, Amin Soeban |
| EPI_ISL_1622433 | RSJ Dr. Soeharto Heerdjan | Eijkman Institute for Molecular Biology, Ministry of Research and Technology/National Agency for Research and Innovation | Muhammad Rezki Rasyak, Willy Agustine, Hidayat Trimarsanto, Lydia V. Panggalo, Iskandar Adnan, Sukma Oktavianthi, Edison Johar, Frilasita A Yudhaputri, Safarina G Malik, Khin Saw Myint, Amin Soeban |
| EPI_ISL_1622434 | RS Permata Cibubur | Eijkman Institute for Molecular Biology, Ministry of Research and Technology/National Agency for Research and Innovation | Iskandar Adnan, Sukma Oktavianthi, Edison Johar, Frilasita A Yudhaputri, Muhammad Rezki Rasyak, Willy Agustine, Hidayat Trimarsanto, Lydia V. Panggalo, Safarina G Malik, Khin Saw Myint, Amin Soeban |
| EPI_ISL_1622435 | RSIA Selasih Medika | Eijkman Institute for Molecular Biology, Ministry of Research and Technology/National Agency for Research and Innovation | Lydia V. Panggalo, Iskandar Adnan, Sukma Oktavianthi, Edison Johar, Frilasita A Yudhaputri, Muhammad Rezki Rasyak, Willy Agustine, Hidayat Trimarsanto, Safarina G Malik, Khin Saw Myint, Amin Soeban |
| EPI_ISL_1630278, EPI_ISL_1630279, EPI_ISL_1630280 | Clinical Microbiology Lab RSPTN Universitas | Clinical Microbiology Lab RSPTN Universitas Hasanuddin | Muhammad Nasrum Massi, Rizalinda Sjahril, Gita Vita Soraya, Triwibawa, Handayani Halik, Muhammad Yusuf Usman, Rufika Shari |

| | | | |
|---|---|---|---|
| | Hasanuddin | | |
| EPI_ISL_1633339 | Bandung Public Health | West Java Health Laboratory; School of Life Sciences and Technology, Institut Teknologi Bandung | Azzania Fibriani, Ema Rahmawati, Ryan Bayusantika Ristandi, Rifky Waluyajati Rachman, Cut Nur Cinthia Alamanda, Rini Robiani, Miftahul Faridl, Karimatu Khoirunnisa, Aulia Saraswati Wicaksono |
| EPI_ISL_1634442 | Mitra Kasih Hospital | West Java Health Laboratory; School of Life Sciences and Technology, Institut Teknologi Bandung | Azzania Fibriani, Ema Rahmawati, Ryan Bayusantika Ristandi, Rifky Waluyajati Rachman, Cut Nur Cinthia Alamanda, Rini Robiani, Miftahul Faridl, Karimatu Khoirunnisa, Aulia Saraswati Wicaksono |
| EPI_ISL_1634465 | Bandung Public Health | West Java Health Laboratory; School of Life Sciences and Technology, Institut Teknologi Bandung | Azzania Fibriani, Ema Rahmawati, Ryan Bayusantika Ristandi, Rifky Waluyajati Rachman, Cut Nur Cinthia Alamanda, Rini Robiani, Miftahul Faridl, Karimatu Khoirunnisa, Aulia Saraswati Wicaksono |
| EPI_ISL_1634467 | Purwakarta Public Health | West Java Health Laboratory; School of Life Sciences and Technology, Institut Teknologi Bandung | Azzania Fibriani, Ema Rahmawati, Ryan Bayusantika Ristandi, Rifky Waluyajati Rachman, Cut Nur Cinthia Alamanda, Rini Robiani, Miftahul Faridl, Karimatu Khoirunnisa, Aulia Saraswati Wicaksono |
| EPI_ISL_1634470 | Tasikmalaya Public Health | West Java Health Laboratory; School of Life Sciences and Technology, Institut Teknologi Bandung | Azzania Fibriani, Ema Rahmawati, Ryan Bayusantika Ristandi, Rifky Waluyajati Rachman, Cut Nur Cinthia Alamanda, Rini Robiani, Miftahul Faridl, Karimatu Khoirunnisa, Aulia Saraswati Wicaksono |
| EPI_ISL_1634471 | Santosa Kopo Hospital | West Java Health Laboratory; School of Life Sciences and Technology, Institut Teknologi Bandung | Azzania Fibriani, Ema Rahmawati, Ryan Bayusantika Ristandi, Rifky Waluyajati Rachman, Cut Nur Cinthia Alamanda, Rini Robiani, Miftahul Faridl, Karimatu Khoirunnisa, Aulia Saraswati Wicaksono |
| EPI_ISL_1634474 | Cimahi Public Health | West Java Health Laboratory; School of Life Sciences and Technology, Institut Teknologi Bandung | Azzania Fibriani, Ema Rahmawati, Ryan Bayusantika Ristandi, Rifky Waluyajati Rachman, Cut Nur Cinthia Alamanda, Rini Robiani, Miftahul Faridl, Karimatu Khoirunnisa, Aulia Saraswati Wicaksono |
| EPI_ISL_1634475 | Santosa Kopo Hospital | West Java Health Laboratory; School of Life Sciences and Technology, Institut Teknologi Bandung | Azzania Fibriani, Ema Rahmawati, Ryan Bayusantika Ristandi, Rifky Waluyajati Rachman, Cut Nur Cinthia Alamanda, Rini Robiani, Miftahul Faridl, Karimatu Khoirunnisa, Aulia Saraswati Wicaksono |
| EPI_ISL_1634477 | Subang Public Health | West Java Health Laboratory; School of Life Sciences and Technology, Institut Teknologi Bandung | Azzania Fibriani, Ema Rahmawati, Ryan Bayusantika Ristandi, Rifky Waluyajati Rachman, Cut Nur Cinthia Alamanda, Rini Robiani, Miftahul Faridl, Karimatu Khoirunnisa, Aulia Saraswati Wicaksono |
| EPI_ISL_1660412 | AMC Cileunyi Hospital | West Java Health Laboratory; School of Life Sciences and Technology, Institut Teknologi Bandung | Azzania Fibriani, Ema Rahmawati, Ryan Bayusantika Ristandi, Rifky Waluyajati Rachman, Cut Nur Cinthia Alamanda, Rini Robiani, Miftahul Faridl, Karimatu Khoirunnisa, Aulia Saraswati Wicaksono |
| EPI_ISL_1660413 | West Java Health Laboratory | West Java Health Laboratory; School of Life Sciences and Technology, Institut Teknologi Bandung | Azzania Fibriani, Ema Rahmawati, Ryan Bayusantika Ristandi, Rifky Waluyajati Rachman, Cut Nur Cinthia Alamanda, Rini Robiani, Miftahul Faridl, Karimatu Khoirunnisa, Aulia Saraswati Wicaksono |
| EPI_ISL_1660414 | Mitra Kasih Hospital | West Java Health Laboratory; School of Life Sciences and Technology, Institut Teknologi Bandung | Azzania Fibriani, Ema Rahmawati, Ryan Bayusantika Ristandi, Rifky Waluyajati Rachman, Cut Nur Cinthia Alamanda, Rini Robiani, Miftahul Faridl, Karimatu Khoirunnisa, Aulia Saraswati Wicaksono |
| EPI_ISL_1660415 | West Bandung Public Health | West Java Health Laboratory; School of Life Sciences and Technology, Institut Teknologi Bandung | Azzania Fibriani, Ema Rahmawati, Ryan Bayusantika Ristandi, Rifky Waluyajati Rachman, Cut Nur Cinthia Alamanda, Rini Robiani, Miftahul Faridl, Karimatu Khoirunnisa, Aulia Saraswati Wicaksono |
| EPI_ISL_1660416 | West Java Health Laboratory | West Java Health Laboratory; School of Life Sciences and Technology, Institut Teknologi Bandung | Azzania Fibriani, Ema Rahmawati, Ryan Bayusantika Ristandi, Rifky Waluyajati Rachman, Cut Nur Cinthia Alamanda, Rini Robiani, Miftahul Faridl, Karimatu Khoirunnisa, Aulia Saraswati Wicaksono |
| EPI_ISL_1660417 | Subang Public Health | West Java Health Laboratory; School of Life Sciences and Technology, Institut Teknologi Bandung | Azzania Fibriani, Ema Rahmawati, Ryan Bayusantika Ristandi, Rifky Waluyajati Rachman, Cut Nur Cinthia Alamanda, Rini Robiani, Miftahul Faridl, Karimatu Khoirunnisa, Aulia Saraswati Wicaksono |
| EPI_ISL_1660418 | AMC Cileunyi Hospital | West Java Health Laboratory; School of Life Sciences and Technology, Institut Teknologi Bandung | Azzania Fibriani, Ema Rahmawati, Ryan Bayusantika Ristandi, Rifky Waluyajati Rachman, Cut Nur Cinthia Alamanda, Rini Robiani, Miftahul Faridl, Karimatu Khoirunnisa, Aulia Saraswati Wicaksono |
| EPI_ISL_1660419 | Santosa Kopo Hospital | West Java Health Laboratory; School of Life Sciences and Technology, Institut Teknologi Bandung | Azzania Fibriani, Ema Rahmawati, Ryan Bayusantika Ristandi, Rifky Waluyajati Rachman, Cut Nur Cinthia Alamanda, Rini Robiani, Miftahul Faridl, Karimatu Khoirunnisa, Aulia Saraswati Wicaksono |
| EPI_ISL_1660420 | Subang Public Health | West Java Health Laboratory; School of Life Sciences and Technology, Institut Teknologi Bandung | Azzania Fibriani, Ema Rahmawati, Ryan Bayusantika Ristandi, Rifky Waluyajati Rachman, Cut Nur Cinthia Alamanda, Rini Robiani, Miftahul Faridl, Karimatu Khoirunnisa, Aulia Saraswati Wicaksono |
| EPI_ISL_1660472 | Bandung Public Health | West Java Health Laboratory; School of Life Sciences and Technology, Institut Teknologi Bandung | Azzania Fibriani, Ema Rahmawati, Ryan Bayusantika Ristandi, Rifky Waluyajati Rachman, Cut Nur Cinthia Alamanda, Rini Robiani, Miftahul Faridl, Karimatu Khoirunnisa, Aulia Saraswati Wicaksono |

| | | | |
|---|---|---|---|
| EPI_ISL_1660473 | West Bandung Public Health | West Java Health Laboratory; School of Life Sciences and Technology, Institut Teknologi Bandung | Azzania Fibriani, Ema Rahmawati, Ryan Bayusantika Ristandi, Rifky Waluyajati Rachman, Cut Nur Cinthia Alamanda, Rini Robiani, Miftahul Faridl, Karimatu Khoirunnisa, Aulia Saraswati Wicaksono |
| EPI_ISL_1660474 | Immanuel Hospital | West Java Health Laboratory; School of Life Sciences and Technology, Institut Teknologi Bandung | Azzania Fibriani, Ema Rahmawati, Ryan Bayusantika Ristandi, Rifky Waluyajati Rachman, Cut Nur Cinthia Alamanda, Rini Robiani, Miftahul Faridl, Karimatu Khoirunnisa, Aulia Saraswati Wicaksono |
| EPI_ISL_1789112 | Klinik Tanjung Laboratory | Institute of Tropical Disease, Universitas Airlangga | Krisnoadi Rahardjo, Aldise M Nastri, Jezzy R Dewantari, Rima R Prasetya, Gatot Soegiarto, Laksmi Wulandari, Resti Yudhawati, Soetjipto, Yasuko Mori, Maria I Lusida, Kazufumi Shimizu |
| EPI_ISL_1789113 | Siti Khodijah Hospital | Institute of Tropical Disease, Universitas Airlangga | Aldise M Nastri, Jezzy R Dewantari, Rima R Prasetya, Krisnoadi Rahardjo, Muhammad Hamdan, Gatot Soegiarto, Laksmi Wulandari, Resti Yudhawati, Soetjipto, Yasuko Mori, Maria I Lusida, Kazufumi Shimi |
| EPI_ISL_1789114 | Siti Khodijah Hospital | Institute of Tropical Disease, Universitas Airlangga | Jezzy R Dewantari, Rima R Prasetya, Krisnoadi Rahardjo, Aldise M Nastri, Muhammad Hamdan, Gatot Soegiarto, Laksmi Wulandari, Resti Yudhawati, Soetjipto, Yasuko Mori, Maria I Lusida, Kazufumi Shimi |
| EPI_ISL_1789115 | Siti Khodijah Hospital | Institute of Tropical Disease, Universitas Airlangga | Rima R Prasetya, Krisnoadi Rahardjo, Aldise M Nastri, Jezzy R Dewantari, Muhammad Hamdan, Gatot Soegiarto, Laksmi Wulandari, Resti Yudhawati, Soetjipto, Yasuko Mori, Maria I Lusida, Kazufumi Shimi |
| EPI_ISL_1789116 | Anwar Medika General Hospital | Institute of Tropical Disease, Universitas Airlangga | Maria I Lusida, Krisnoadi Rahardjo, Aldise M Nastri, Jezzy R Dewantari, Rima R Prasetya, Nungky Taniasari, Gatot Soegiarto, Laksmi Wulandari, Resti Yudhawati, Soetjipto, Yasuko Mori, Kazufumi Shimizu |
| EPI_ISL_1789117 | Anwar Medika General Hospital | Institute of Tropical Disease, Universitas Airlangga | Krisnoadi Rahardjo, Aldise M Nastri, Jezzy R Dewantari, Rima R Prasetya, Muhammad Hamdan, Gatot Soegiarto, Laksmi Wulandari, Resti Yudhawati, Yasuko Mori, Soetjipto, Kazufumi Shimizu, Maria I Lusi |
| EPI_ISL_1789118 | Institute of Tropical Disease, Universitas Airlangga | Institute of Tropical Disease, Universitas Airlangga | Aldise M Nastri, Jezzy R Dewantari, Rima R Prasetya, Krisnoadi Rahardjo, Gatot Soegiarto, Laksmi Wulandari, Resti Yudhawati, Soetjipto, Yasuko Mori, Maria I Lusida, Kazufumi Shimizu |
| EPI_ISL_1789119 | Institute of Tropical Disease, Universitas Airlangga | Institute of Tropical Disease, Universitas Airlangga | Jezzy R Dewantari, Rima R Prasetya, Krisnoadi Rahardjo, Aldise M Nastri, Gatot Soegiarto, Laksmi Wulandari, Resti Yudhawati, Soetjipto, Yasuko Mori, Maria I Lusida, Kazufumi Shimizu |
| EPI_ISL_1789172 | Institute of Tropical Disease, Universitas Airlangga | Institute of Tropical Disease, Universitas Airlangga | Rima R Prasetya, Krisnoadi Rahardjo, Aldise M Nastri, Jezzy R Dewantari, Gatot Soegiarto, Laksmi Wulandari, Resti Yudhawati, Soetjipto, Yasuko Mori, Maria I Lusida, Kazufumi Shimizu |
| EPI_ISL_1789202 | Paru Hospital | Institute of Tropical Disease, Universitas Airlangga | Maria I Lusida, Krisnoadi Rahardjo, Aldise M Nastri, Jezzy R Dewantari, Rima R Prasetya, Dyah Retno, Gatot Soegiarto, Laksmi Wulandari, Resti Yudhawati, Soetjipto, Yasuko Mori, Kazufumi Shimizu |
| EPI_ISL_1795053 | Anwar Medika General Hospital | Institute of Tropical Disease, Universitas Airlangga | Aldise M Nastri, Jezzy R Dewantari, Rima R Prasetya, Krisnoadi Rahardjo, Nungky Taniasari, Gatot Soegiarto, Laksmi Wulandari, Resti Yudhawati, Yasuko Mori, Soetjipto, Kazufumi Shimizu, Maria I Lusida |
| EPI_ISL_1795054 | Darus Syifa' Islamic Hospital | Institute of Tropical Disease, Universitas Airlangga | Krisnoadi Rahardjo, Aldise M Nastri, Jezzy R Dewantari, Rima R Prasetya, HM Faiz, Gatot Soegiarto, Laksmi Wulandari, Resti Yudhawati, Yasuko Mori, Soetjipto, Kazufumi Shimizu, Maria I Lusida |
| EPI_ISL_1795055 | Muhammadiyah Hospital | Institute of Tropical Disease, Universitas Airlangga | Rima R Prasetya, Krisnoadi Rahardjo, Aldise M Nastri, Jezzy R Dewantari, Imam Suyuthi, Gatot Soegiarto, Laksmi Wulandari, Resti Yudhawati, Yasuko Mori, Soetjipto, Kazufumi Shimizu, Maria I Lusida |
| EPI_ISL_1795056 | Siloam Hospital | Institute of Tropical Disease, Universitas Airlangga | Kazufumi Shimizu, Krisnoadi Rahardjo, Aldise M Nastri, Jezzy R Dewantari, Rima R Prasetya, Maria M Padmidewi, Gatot Soegiarto, Laksmi Wulandari, Retno A Setyoningrum, Resti Yudhawati, Yohko K Shimizu, Mitsuhiro Nishimura, Yasu |
| EPI_ISL_1795057 | Muhammadiyah Hospital | Institute of Tropical Disease, Universitas Airlangga | Jezzy R Dewantari, Rima R Prasetya, Krisnoadi Rahardjo, Aldise M Nastri, Imam Suyuthi, Gatot Soegiarto, Laksmi Wulandari, Resti Yudhawati, Yasuko Mori, Soetjipto, Kazufumi Shimizu, Maria I Lusida |
| EPI_ISL_1795058 | Institute of Tropical Disease, Universitas Airlangga | Institute of Tropical Disease, Universitas Airlangga | Krisnoadi Rahardjo, Aldise M Nastri, Jezzy R Dewantari, Rima R Prasetya, Gatot Soegiarto, Laksmi Wulandari, Resti Yudhawati, Yasuko Mori, Soetjipto, Kazufumi Shimizu, Maria I Lusida |
| EPI_ISL_1795059 | Citra Medika Hospital | Institute of Tropical Disease, Universitas Airlangga | Aldise M Nastri, Jezzy R Dewantari, Rima R Prasetya, Krisnoadi Rahardjo, Wahidah Rachmaniyah, Gatot Soegiarto, Laksmi Wulandari, Resti Yudhawati, Yasuko Mori, Soetjipto, Kazufumi Shimizu, Maria I Lus |
| EPI_ISL_1795060 | Institute of Tropical Disease, Universitas Airlangga | Institute of Tropical Disease, Universitas Airlangga | Kazufumi Shimizu, Krisnoadi Rahardjo, Aldise M Nastri, Jezzy R Dewantari, Rima R Prasetya, Gatot Soegiarto, Laksmi Wulandari, Retno A Setyoningrum, Resti Yudhawati, Yohko K Shimizu, Mitsuhiro Nishimura, Yasuko Mori, S |
| EPI_ISL_1824601, EPI_ISL_1824602, EPI_ISL_1824603 | National Institute of Health Research and Development | National Institute of Health Research and Development | Subangkit, Hana Apsari Pawestri, Kartika Dewi Puspa, Arie Ardiansyah Nugraha, Hartanti Dian Ikawati, Krisna Nur Andriana Pangesti, Yuni Rukminiati, Ririn Ramadhany, Agustiningsih, Kindi Adam, Holy Arif Wibowo, Triyani Soekarso, Ni Ketut Susilarini, Nurika Ha[illegible] Tati Febriyanti, Natalie Laurencia Kipuw, Fauzul Muna, Irene Lorinda Indalao, Nelly Puspandari, Vivi Setiawaty. |
| EPI_ISL_1824604 | PRVKP FK UI | National Institute of Health Research and Development | Vivi Setiawaty, Hana Apsari Pawestri, Subangkit, Kartika Dewi Puspa, Arie Ardiansyah Nugraha, Hartanti Dian Ikawati, Krisna Pangesti, Nelly Puspandari |
| EPI_ISL_1824605, EPI_ISL_1824606 | RSUP Sanglah | National Institute of Health Research and Development | Vivi Setiawaty, Hana Apsari Pawestri, Subangkit, Kartika Dewi Puspa, Arie Ardiansyah Nugraha, Hartanti Dian Ikawati, Krisna Pangesti, Nelly Puspandari, I Nengah Tony Rustawan, I Gede Ketut Sajinadiyas |
| EPI_ISL_1824607 | PKC Jagakarsa | National Institute of Health Research and Development | Vivi Setiawaty, Hana Apsari Pawestri, Subangkit, Kartika Dewi Puspa, Arie Ardiansyah Nugraha, Hartanti Dian Ikawati, Krisna Pangesti, Nelly Puspandari |
| EPI_ISL_1824608, EPI_ISL_1824609, EPI_ISL_1824610, EPI_ISL_1824611, | National Institute of Health Research and Development | National Institute of Health Research and Development | Subangkit, Hana Apsari Pawestri, Kartika Dewi Puspa, Arie Ardiansyah Nugraha, Hartanti Dian Ikawati, Krisna Nur Andriana Pangesti, Yuni Rukminiati, Ririn Ramadhany, Agustiningsih, Kindi Adam, Holy Arif Wibowo, Triyani Soekarso, Ni Ketut Susilarini, Nurika Ha[illegible] Tati Febriyanti, Natalie Laurencia Kipuw, Fauzul Muna, Irene Lorinda Indalao, Nelly Puspandari, Vivi Setiawaty. |

| | | | |
|---|---|---|---|
| EPI_ISL_1824612, EPI_ISL_1824613, EPI_ISL_1824614 | | | |
| EPI_ISL_1824615 | RSUD dr. Hasri Ainun Habibie Gorontalo | National Institute of Health Research and Development | Hana Apsari Pawestri, Subangkit, Kartika Dewi Puspa, Arie Ardiansyah Nugraha, Hartanti Dian Ikawati, Krisna Pangesti, Nelly Puspandari, Vivi Setiawaty |
| EPI_ISL_1824616, EPI_ISL_1824617, EPI_ISL_1824618 | RSUD Bahteramas Kendari | National Institute of Health Research and Development | Hana Apsari Pawestri, Subangkit, Kartika Dewi Puspa, Arie Ardiansyah Nugraha, Hartanti Dian Ikawati, Krisna Pangesti, Nelly Puspandari, Vivi Setiawaty |
| EPI_ISL_1824719 | National Institute of Health Research and Development | National Institute of Health Research and Development | Subangkit, Hana Apsari Pawestri, Kartika Dewi Puspa, Arie Ardiansyah Nugraha, Hartanti Dian Ikawati, Krisna Nur Andriana Pangesti, Yuni Rukminiati, Ririn Ramadhany, Agustiningsih, Kindi Adam, Holy Arif Wibowo, Triyani Soekarso, Ni Ketut Susilarini, Nurika Ha[illegible] Tati Febriyanti, Natalie Laurencia Kipuw, Fauzul Muna, Irene Lorinda Indalao, Nelly Puspandari, Vivi Setiawaty. |
| EPI_ISL_1855390 | Indonesian Research Center for Veterinary Science/BBalitvet | Indonesian Research Center for Veterinary Science/BBalitvet | Ni Luh Putu Indi Dharmayanti, Diana Nurjanah, Risa Indriani, Harimurti Nuradji, Ilham Chaidir, Sri Nowo Retno, Fadjry Djufry |
| EPI_ISL_1855939 | Dinas Kesehatan Kota Bogor | Indonesian Research Center for Veterinary Science / BBalitvet | Ni Luh Putu Indi Dharmayanti, Diana Nurjanah, Risa Indriani, Harimurti Nuradji, Ilham Chaidir, Sri Nowo Retno, Fadjry Djufry |
| EPI_ISL_1915511, EPI_ISL_1915512, EPI_ISL_1915513, EPI_ISL_1915514, EPI_ISL_1915515, EPI_ISL_1915516, EPI_ISL_1915517, EPI_ISL_1915518, EPI_ISL_1915519, EPI_ISL_1915520, EPI_ISL_1915521, EPI_ISL_1915522, EPI_ISL_1915523, EPI_ISL_1915524, EPI_ISL_1915525, EPI_ISL_1915526, EPI_ISL_1915527, EP[illegible] EPI_ISL_1915531, EPI_ISL_1915532, EPI_ISL_1915533, EPI_ISL_1915534, EPI_ISL_1915535, EPI_ISL_1915536, EPI_ISL_1915537, EPI_ISL_1915538, EPI_ISL_1915539, EPI_ISL_1915540, EPI_ISL_1915541, EPI_ISL_1915542, EPI_ISL_1915543, EPI_ISL_1915544, EPI_ISL_1915545, EPI_ISL_1915546, EPI_ISL_1915547, EP[illegible] | | | |
| see above | National Institute of Health Research and Development | National Institute of Health Research and Development | Subangkit, Hana Apsari Pawestri, Kartika Dewi Puspa, Arie Ardiansyah Nugraha, Hartanti Dian Ikawati, Krisna Nur Andriana Pangesti, Yuni Rukminiati, Ririn Ramadhany, Agustiningsih, Kindi Adam, Holy Arif Wibowo, Triyani Soekarso, Ni Ketut Susilarini, Nurika Ha[illegible] Tati Febriyanti, Natalie Laurencia Kipuw, Fauzul Muna, Irene Lorinda Indalao, Nelly Puspandari, Vivi Setiawaty. |
| EPI_ISL_1915550 | RSUP Sanglah | National Institute of Health Research and Development | Vivi Setiawaty, Hana Apsari Pawestri, Subangkit, Kartika Dewi Puspa, Arie Ardiansyah Nugraha, Hartanti Dian Ikawati, Krisna Pangesti, Triyani Soekarso, Nelly Puspandari, I Nengah Tony Rustawan, I Gede Ketut S[illegible] |
| EPI_ISL_1915551, EPI_ISL_1915552, EPI_ISL_1915553 | PRVKP FK UI | National Institute of Health Research and Development | Hartanti Dian Ikawati, Hana Apsari Pawestri, Subangkit, Kartika Dewi Puspa, Arie Ardiansyah Nugraha, Triyani Soekarso, Krisna Pangesti, Nelly Puspandari, Vivi Setiawaty |
| EPI_ISL_1915554, EPI_ISL_1915555, EPI_ISL_1915556, EPI_ISL_1915557, EPI_ISL_1915558, EPI_ISL_1915559 | RSUP Sanglah | National Institute of Health Research and Development | Vivi Setiawaty, Hana Apsari Pawestri, Subangkit, Kartika Dewi Puspa, Arie Ardiansyah Nugraha, Hartanti Dian Ikawati, Krisna Pangesti, Triyani Soekarso, Nelly Puspandari, I Nengah Tony Rustawan, I Gede Ketut S[illegible] |
| EPI_ISL_1915560, EPI_ISL_1915561 | RSPI Sulianti Saroso Jakarta Utara | National Institute of Health Research and Development | Hartanti Dian Ikawati, Hana Apsari Pawestri, Subangkit, Kartika Dewi Puspa, Arie Ardiansyah Nugraha, Triyani Soekarso, Krisna Pangesti, Nelly Puspandari, Vivi Setiawaty |
| EPI_ISL_1915562, EPI_ISL_1915563 | BBTKL PP Banjarbaru | National Institute of Health Research and Development | Hartanti Dian Ikawati, Hana Apsari Pawestri, Subangkit, Kartika Dewi Puspa, Arie Ardiansyah Nugraha, Triyani Soekarso, Krisna Pangesti, Nelly Puspandari, Vivi Setiawaty |
| EPI_ISL_1915564, EPI_ISL_1915565 | Kantor Pengelola Sumbawa Technopark | National Institute of Health Research and Development | Hana Apsari Pawestri, Subangkit, Kartika Dewi Puspa, Arie Ardiansyah Nugraha, Hartanti Dian Ikawati, Krisna Pangesti, Nelly Puspandari, Arief Budi Witarto, Rian Adha Ardinata, Lalu Hamzi Fikri, Nurhandini Eka Dewi, Laily In[illegible] |
| EPI_ISL_1915566, EPI_ISL_1915567 | RSPI Prof DR Sulianti Saroso Jakarta Utara | National Institute of Health Research and Development | Hartanti Dian Ikawati, Hana Apsari Pawestri, Subangkit, Kartika Dewi Puspa, Arie Ardiansyah Nugraha, Triyani Soekarso, Krisna Pangesti, Nelly Puspandari, Vivi Setiawaty |
| EPI_ISL_1915568 | RSPI Prof DR Sulianti Saroso Jakarta Utara | National Institute of Health Research and Development | Hana Apsari Pawestri, Subangkit, Kartika Dewi Puspa, Arie Ardiansyah Nugraha, Hartanti Dian Ikawati, Krisna Pangesti, Nelly Puspandari, Vivi Setiawaty |
| EPI_ISL_1915569, EPI_ISL_1915570 | RSUP Sanglah | National Institute of Health Research and Development | Hana Apsari Pawestri, Subangkit, Kartika Dewi Puspa, Arie Ardiansyah Nugraha, Hartanti Dian Ikawati, Krisna Pangesti, Nelly Puspandari, Vivi Setiawaty |
| EPI_ISL_1915571 | PKC Cilincing | National Institute of Health Research and Development | Hana Apsari Pawestri, Subangkit, Kartika Dewi Puspa, Arie Ardiansyah Nugraha, Hartanti Dian Ikawati, Krisna Pangesti, Nelly Puspandari, Vivi Setiawaty |
| EPI_ISL_1915572, EPI_ISL_1915573, EPI_ISL_1915574, EPI_ISL_1915575 | Balitbang Tanah Bumbu | National Institute of Health Research and Development | Hana Apsari Pawestri, Subangkit, Kartika Dewi Puspa, Arie Ardiansyah Nugraha, Hartanti Dian Ikawati, Krisna Pangesti, Nelly Puspandari, Vivi Setiawaty |
| EPI_ISL_1915576 | RSUD Siti Fatimah Prov Sumatera Selatan | National Institute of Health Research and Development | Hana Apsari Pawestri, Subangkit, Kartika Dewi Puspa, Arie Ardiansyah Nugraha, Hartanti Dian Ikawati, Krisna Pangesti, Nelly Puspandari, Vivi Setiawaty |
| EPI_ISL_1915577 | RS Bunda | National Institute of Health Research and Development | Hana Apsari Pawestri, Subangkit, Kartika Dewi Puspa, Arie Ardiansyah Nugraha, Hartanti Dian Ikawati, Krisna Pangesti, Nelly Puspandari, Vivi Setiawaty |
| EPI_ISL_1915578 | Labkesda Jakarta | National Institute of Health Research and Development | Hana Apsari Pawestri, Subangkit, Kartika Dewi Puspa, Arie Ardiansyah Nugraha, Hartanti Dian Ikawati, Krisna Pangesti, Nelly Puspandari, Vivi Setiawaty |
| EPI_ISL_1915579 | BPOM Gorontalo | National Institute of Health Research and Development | Hartanti Dian Ikawati, Hana Apsari Pawestri, Subangkit, Kartika Dewi Puspa, Arie Ardiansyah Nugraha, Triyani Soekarso, Krisna Pangesti, Nelly Puspandari, Vivi Setiawaty |
| EPI_ISL_1915580, EPI_ISL_1915581, EPI_ISL_1915582 | RSUD UNDATA Palu | National Institute of Health Research and Development | Hana Apsari Pawestri, Subangkit, Kartika Dewi Puspa, Arie Ardiansyah Nugraha, Hartanti Dian Ikawati, Krisna Pangesti, Nelly Puspandari, Rosa Dwi Wahyuni, Vivi Setiawaty |
| EPI_ISL_1969239, EPI_ISL_1969240 | Hotel Grand Sahid Jaya | National Institute of Health Research and Development | Vivi Setiawaty, Hana Apsari Pawestri, Subangkit, Kartika Dewi Puspa, Arie Ardiansyah Nugraha, Hartanti Dian Ikawati, Krisna Pangesti, Triyani Soekarso, Nelly Puspandari |
| EPI_ISL_1969241 | Hotel Grand Tjokro | National Institute of Health Research and Development | Vivi Setiawaty, Hana Apsari Pawestri, Subangkit, Kartika Dewi Puspa, Arie Ardiansyah Nugraha, Hartanti Dian Ikawati, Krisna Pangesti, Triyani Soekarso, Nelly Puspandari |
| EPI_ISL_1969242, | Swissbel Hotel | National Institute of Health | Vivi Setiawaty, Hana Apsari Pawestri, Subangkit, Kartika Dewi Puspa, Arie Ardiansyah Nugraha, Hartanti Dian Ikawati, Krisna Pangesti, Triyani Soekarso, Nelly Puspandari |

| | | | |
|---|---|---|---|
| EPI_ISL_1969243 | Airport | Research and Development | |
| EPI_ISL_1969244 | Lab RSUP DR Mohammad Hoesin Palembang | National Institute of Health Research and Development | Vivi Setiawaty, Hana Apsari Pawestri, Subangkit, Kartika Dewi Puspa, Arie Ardiansyah Nugraha, Hartanti Dian Ikawati, Krisna Pangesti, Triyani Soekarso, Nelly Puspandari |
| EPI_ISL_1969245 | Lab RSUP DR Mohammad Hoesin Palembang | National Institute of Health Research and Development | Subangkit, Hana Apsari Pawestri, Kartika Dewi Puspa, Arie Ardiansyah Nugraha, Hartanti Dian Ikawati, Krisna Pangesti, Triyani Soekarso, Nelly Puspandari, Vivi Setiawaty |
| EPI_ISL_1969246, EPI_ISL_1969247, EPI_ISL_1969248 | Rumah Sakit Umum Daerah Palangkaraya | National Institute of Health Research and Development | Subangkit, Hana Apsari Pawestri, Kartika Dewi Puspa, Arie Ardiansyah Nugraha, Hartanti Dian Ikawati, Krisna Pangesti, Triyani Soekarso, Nelly Puspandari, Vivi Setiawaty |
| EPI_ISL_1969249, EPI_ISL_1969250 | BBLK Palembang | National Institute of Health Research and Development | Subangkit, Hana Apsari Pawestri, Kartika Dewi Puspa, Arie Ardiansyah Nugraha, Hartanti Dian Ikawati, Krisna Pangesti, Triyani Soekarso, Nelly Puspandari, Vivi Setiawaty |
| EPI_ISL_2013043 | RS Sentra Medika Cikarang | Eijkman Institute for Molecular Biology, National Agency for Research and Innovation | Edison Johar, Frilasita A Yudhaputri, Muhammad Rezki Rasyak, Willy Agustine, Hidayat Trimarsanto, Lydia V. Panggalo, Iskandar Adnan, Sukma Oktavianthi, Safarina G Malik, Khin Saw Myint, Amin Soeban[illegible] |
| EPI_ISL_2013044 | RS Sentra Medika Cikarang | Eijkman Institute for Molecular Biology, National Agency for Research and Innovation | Willy Agustine, Hidayat Trimarsanto, Lydia V. Panggalo, Iskandar Adnan, Sukma Oktavianthi, Edison Johar, Frilasita A Yudhaputri, Muhammad Rezki Rasyak, Safarina G Malik, Khin Saw Myint, Amin Soeban[illegible] |
| EPI_ISL_2013045 | RS Mitra Keluarga Cikarang | Eijkman Institute for Molecular Biology, National Agency for Research and Innovation | Edison Johar, Frilasita A Yudhaputri, Muhammad Rezki Rasyak, Willy Agustine, Hidayat Trimarsanto, Lydia V. Panggalo, Iskandar Adnan, Sukma Oktavianthi, Safarina G Malik, Khin Saw Myint, Amin Soeban[illegible] |
| EPI_ISL_2013046 | RSUD Cileungsi | Eijkman Institute for Molecular Biology, National Agency for Research and Innovation | Lydia V. Panggalo, Iskandar Adnan, Sukma Oktavianthi, Edison Johar, Frilasita A Yudhaputri, Muhammad Rezki Rasyak, Willy Agustine, Hidayat Trimarsanto, Safarina G Malik, Khin Saw Myint, Amin Soeban[illegible] |
| EPI_ISL_2013047 | RS Mitra Keluarga Kelapa Gading | Eijkman Institute for Molecular Biology, National Agency for Research and Innovation | Hidayat Trimarsanto, Lydia V. Panggalo, Iskandar Adnan, Sukma Oktavianthi, Edison Johar, Frilasita A Yudhaputri, Muhammad Rezki Rasyak, Willy Agustine, Safarina G Malik, Khin Saw Myint, Amin Soeban[illegible] |
| EPI_ISL_2013509 | RS Harapan Bunda | Eijkman Institute for Molecular Biology, National Agency for Research and Innovation | Lydia V. Panggalo, Iskandar Adnan, Sukma Oktavianthi, Edison Johar, Frilasita A Yudhaputri, Muhammad Rezki Rasyak, Willy Agustine, Hidayat Trimarsanto, Safarina G Malik, Khin Saw Myint, Amin Soeban[illegible] |
| EPI_ISL_2047507, EPI_ISL_2047508, EPI_ISL_2047509, EPI_ISL_2047510, EPI_ISL_2047511, EPI_ISL_2047512 | Lab RSUP DR Mohammad Hoesin Palembang | National Institute of Health Research and Development | Arie Ardiansyah Nugraha, Kartika Dewi Puspa, Hana Apsari Pawestri, Hartanti Dian Ikawati, Krisna Pangesti, Triyani Soekarso, Nelly Puspandari, Subangkit, Eny Rahmawati, Rogaya, Vivi Setiawaty |
| EPI_ISL_2047513, EPI_ISL_2047514, EPI_ISL_2047515, EPI_ISL_2047516, EPI_ISL_2047517, EPI_ISL_2047518, EPI_ISL_2047519, EPI_ISL_2047520, EPI_ISL_2047521, EPI_ISL_2047522, EPI_ISL_2047523, EPI_ISL_2047524, EPI_ISL_2047525, EPI_ISL_2047526, EPI_ISL_2047527, EPI_ISL_2047528 | | | |
| see above | RSUP H Adam Malik Medan | National Institute of Health Research and Development | Kartika Dewi Puspa, Hana Apsari Pawestri, Arie Ardiansyah Nugraha, Hartanti Dian Ikawati, Krisna Pangesti, Triyani Soekarso, Nelly Puspandari, Subangkit, Maliana, Ade Rahmaini, Nelly Elfrida Samosir, Rina Yunita, [illegible] |
| EPI_ISL_2047529 | RSUD Kab Buleleng | National Institute of Health Research and Development | Hana Apsari Pawestri, Kartika Dewi Puspa, Arie Ardiansyah Nugraha, Hartanti Dian Ikawati, Krisna Pangesti, Triyani Soekarso, Nelly Puspandari, Subangkit, Vivi Setiawaty |
| EPI_ISL_2047530 | UPT LabKes Sulawesi Tengah | National Institute of Health Research and Development | Hana Apsari Pawestri, Kartika Dewi Puspa, Arie Ardiansyah Nugraha, Hartanti Dian Ikawati, Krisna Pangesti, Triyani Soekarso, Nelly Puspandari, Subangkit, Vivi Setiawaty |
| EPI_ISL_2047531 | Dinas Kesehatan Kota Denpasar | National Institute of Health Research and Development | Hana Apsari Pawestri, Kartika Dewi Puspa, Arie Ardiansyah Nugraha, Hartanti Dian Ikawati, Krisna Pangesti, Triyani Soekarso, Nelly Puspandari, Subangkit, Vivi Setiawaty |
| EPI_ISL_2047532 | RSUD Bali Mandara | National Institute of Health Research and Development | Hana Apsari Pawestri, Kartika Dewi Puspa, Arie Ardiansyah Nugraha, Hartanti Dian Ikawati, Krisna Pangesti, Triyani Soekarso, Nelly Puspandari, Subangkit, I Wayan Agus Gede Manik Saputra, Vivi Setiawa[illegible] |
| EPI_ISL_2047533, EPI_ISL_2047534, EPI_ISL_2047535, EPI_ISL_2047536, EPI_ISL_2047537, EPI_ISL_2047538, EPI_ISL_2047539, EPI_ISL_2047540, EPI_ISL_2047541, EPI_ISL_2047542, EPI_ISL_2047543, EPI_ISL_2047544, EPI_ISL_2047545, EPI_ISL_2047546 | | | |
| see above | Rumah Sakit Umum Daerah Palangkaraya | National Institute of Health Research and Development | Arie Ardiansyah Nugraha, Kartika Dewi Puspa, Hana Apsari Pawestri, Hartanti Dian Ikawati, Krisna Pangesti, Triyani Soekarso, Nelly Puspandari, Subangkit, Alia Yunita, Fransiska JLA, Mayawati Mewo, Vivi Set[illegible] |
| EPI_ISL_2047547, EPI_ISL_2047548, EPI_ISL_2047549, EPI_ISL_2047550 | Rumah Sakit Umum Daerah Palangkaraya | National Institute of Health Research and Development | Hana Apsari Pawestri, Arie Ardiansyah Nugraha, Kartika Dewi Puspa, Hartanti Dian Ikawati, Krisna Pangesti, Triyani Soekarso, Nelly Puspandari, Subangkit, Alia Yunita, Fransiska JLA, Mayawati Mewo, Vivi Set[illegible] |
| EPI_ISL_2047551, EPI_ISL_2047552, EPI_ISL_2047553, EPI_ISL_2047554 | BBLK Palembang | National Institute of Health Research and Development | Hana Apsari Pawestri, Kartika Dewi Puspa, Arie Ardiansyah Nugraha, Hartanti Dian Ikawati, Krisna Pangesti, Triyani Soekarso, Nelly Puspandari, Subangkit, Vivi Setiawaty |
| EPI_ISL_2047555, EPI_ISL_2047556, EPI_ISL_2047557, EPI_ISL_2047558, EPI_ISL_2047559 | Kantor Pengelola Sumbawa Technopark | National Institute of Health Research and Development | Subangkit, Hana Apsari Pawestri, Kartika Dewi Puspa, Arie Ardiansyah Nugraha, Hartanti Dian Ikawati, Krisna Pangesti, Triyani Soekarso, Nelly Puspandari, Arief Budi Witarto, Rian Adha Ardinata, Lalu Hamzi Fikri, Nurhandini Eka Dewi[illegible] |
| EPI_ISL_2047560 | RSUD Tarakan Jakarta | National Institute of Health Research and Development | Hana Apsari Pawestri, Kartika Dewi Puspa, Arie Ardiansyah Nugraha, Hartanti Dian Ikawati, Krisna Pangesti, Triyani Soekarso, Nelly Puspandari, Subangkit, Juhairiyah, Syarif Hidayat, Vivi Setiawaty |
| EPI_ISL_2047561, EPI_ISL_2047562, EPI_ISL_2047563, | Balai Litbangkes Tanah Bumbu | National Institute of Health Research and Development | Hana Apsari Pawestri, Kartika Dewi Puspa, Arie Ardiansyah Nugraha, Hartanti Dian Ikawati, Krisna Pangesti, Triyani Soekarso, Nelly Puspandari, Subangkit, Juhairiyah, Syarif Hidayat, Vivi Setiawaty |

| | | | |
|---|---|---|---|
| EPI_ISL_2047564, EPI_ISL_2047565, EPI_ISL_2047566, EPI_ISL_2047567, EPI_ISL_2047568, EPI_ISL_2047569 | | | |
| EPI_ISL_2047570, EPI_ISL_2047571 | Dinas Kesehatan Kab Blitar | National Institute of Health Research and Development | Hana Apsari Pawestri, Kartika Dewi Puspa, Arie Ardiansyah Nugraha, Hartanti Dian Ikawati, Krisna Pangesti, Triyani Soekarso, Nelly Puspandari, Subangkit, Vivi Setiawaty |
| EPI_ISL_2047572, EPI_ISL_2047573 | RSUD Siti Fatimah Prov Sumatera Selatan | National Institute of Health Research and Development | Hana Apsari Pawestri, Kartika Dewi Puspa, Arie Ardiansyah Nugraha, Hartanti Dian Ikawati, Krisna Pangesti, Triyani Soekarso, Nelly Puspandari, Subangkit, Vivi Setiawaty |
| EPI_ISL_2047574, EPI_ISL_2047575, EPI_ISL_2047576 | RSPI Prof. Dr. Sulianti Saroso | National Institute of Health Research and Development | Vivi Setiawaty, Hana Apsari Pawestri, Subangkit, Kartika Dewi Puspa, Arie Ardiansyah Nugraha, Hartanti Dian Ikawati, Krisna Pangesti, Triyani Soekarso, Nelly Puspandari |
| EPI_ISL_2101022 | Molecular Biology Laboratory, Faculty Medicine and Health Sciences, Warmadewa University | Eijkman Institute for Molecular Biology, National Agency for Research and Innovation; Molecular Biology Laboratory, Faculty Medicine and Health Sciences, Warmadewa University | Frilasita A Yudhaputri, Muhammad Rezki Rasyak, Willy Agustine, Hidayat Trimarsanto, Lydia V. Panggalo, Iskandar Adnan, Sukma Oktavianthi, Lidwina Priliani, Edison Johar, Sri Masyeni, Erly Sintya, Safarina G Malik, Khin Saw |
| EPI_ISL_2101023 | Molecular Biology Laboratory, Faculty Medicine and Health Sciences, Warmadewa University | Eijkman Institute for Molecular Biology, National Agency for Research and Innovation; Molecular Biology Laboratory, Faculty Medicine and Health Sciences, Warmadewa University | Muhammad Rezki Rasyak, Willy Agustine, Hidayat Trimarsanto, Lydia V. Panggalo, Iskandar Adnan, Sukma Oktavianthi, Lidwina Priliani, Edison Johar, Frilasita A Yudhaputri, Sri Masyeni, Erly Sintya, Safarina G Malik, Khin Saw |
| EPI_ISL_2101024 | Molecular Biology Laboratory, Faculty Medicine and Health Sciences, Warmadewa University | Eijkman Institute for Molecular Biology, National Agency for Research and Innovation; Molecular Biology Laboratory, Faculty Medicine and Health Sciences, Warmadewa University | Willy Agustine, Hidayat Trimarsanto, Lydia V. Panggalo, Iskandar Adnan, Sukma Oktavianthi, Lidwina Priliani, Edison Johar, Frilasita A Yudhaputri, Muhammad Rezki Rasyak, Sri Masyeni, Erly Sintya, Safarina G Malik, Khin Saw |
| EPI_ISL_2101025 | Molecular Biology Laboratory, Faculty Medicine and Health Sciences, Warmadewa University | Eijkman Institute for Molecular Biology, National Agency for Research and Innovation; Molecular Biology Laboratory, Faculty Medicine and Health Sciences, Warmadewa University | Hidayat Trimarsanto, Lydia V. Panggalo, Iskandar Adnan, Sukma Oktavianthi, Lidwina Priliani, Edison Johar, Frilasita A Yudhaputri, Muhammad Rezki Rasyak, Willy Agustine, Sri Masyeni, Erly Sintya, Safarina G Malik, Khin Saw |
| EPI_ISL_2101026 | Molecular Biology Laboratory, Faculty Medicine and Health Sciences, Warmadewa University | Eijkman Institute for Molecular Biology, National Agency for Research and Innovation; Molecular Biology Laboratory, Faculty Medicine and Health Sciences, Warmadewa University | Willy Agustine, Hidayat Trimarsanto, Lydia V. Panggalo, Iskandar Adnan, Sukma Oktavianthi, Lidwina Priliani, Edison Johar, Frilasita A Yudhaputri, Muhammad Rezki Rasyak, Sri Masyeni, Erly Sintya, Safarina G Malik, Khin Saw |
| EPI_ISL_2101027 | Molecular Biology Laboratory, Faculty Medicine and Health Sciences, Warmadewa University | Eijkman Institute for Molecular Biology, National Agency for Research and Innovation; Molecular Biology Laboratory, Faculty Medicine and Health Sciences, Warmadewa University | Iskandar Adnan, Sukma Oktavianthi, Lidwina Priliani, Edison Johar, Frilasita A Yudhaputri, Muhammad Rezki Rasyak, Willy Agustine, Hidayat Trimarsanto, Lydia V. Panggalo, Sri Masyeni, Erly Sintya, Safarina G Malik, Khin Saw |
| EPI_ISL_2101028 | Molecular Biology Laboratory, Faculty Medicine and Health Sciences, Warmadewa University | Eijkman Institute for Molecular Biology, National Agency for Research and Innovation; Molecular Biology Laboratory, Faculty Medicine and Health Sciences, Warmadewa University | Edison Johar, Frilasita A Yudhaputri, Muhammad Rezki Rasyak, Willy Agustine, Hidayat Trimarsanto, Lydia V. Panggalo, Iskandar Adnan, Sukma Oktavianthi, Lidwina Priliani, Sri Masyeni, Erly Sintya, Safarina G Malik, Khin Saw |
| EPI_ISL_2101029 | Molecular Biology Laboratory, Faculty Medicine and Health Sciences, Warmadewa University | Eijkman Institute for Molecular Biology, National Agency for Research and Innovation; Molecular Biology Laboratory, Faculty Medicine and Health Sciences, Warmadewa University | Hidayat Trimarsanto, Lydia V. Panggalo, Iskandar Adnan, Sukma Oktavianthi, Lidwina Priliani, Edison Johar, Frilasita A Yudhaputri, Muhammad Rezki Rasyak, Willy Agustine, Sri Masyeni, Erly Sintya, Safarina G Malik, Khin Saw |
| EPI_ISL_2101030 | Molecular Biology Laboratory, Faculty Medicine and | Eijkman Institute for Molecular Biology, National Agency for Research and | Sukma Oktavianthi, Lidwina Priliani, Edison Johar, Frilasita A Yudhaputri, Muhammad Rezki Rasyak, Willy Agustine, Hidayat Trimarsanto, Lydia V. Panggalo, Iskandar Adnan, Sri Masyeni, Erly Sintya, Safarina G Malik, Khin Saw |

| | | | |
|---|---|---|---|
| | Health Sciences, Warmadewa University | Innovation; Molecular Biology Laboratory, Faculty Medicine and Health Sciences, Warmadewa University | |
| EPI_ISL_2101031 | Molecular Biology Laboratory, Faculty Medicine and Health Sciences, Warmadewa University | Eijkman Institute for Molecular Biology, National Agency for Research and Innovation; Molecular Biology Laboratory, Faculty Medicine and Health Sciences, Warmadewa University | Frilasita A Yudhaputri, Muhammad Rezki Rasyak, Willy Agustine, Hidayat Trimarsanto, Lydia V. Panggalo, Iskandar Adnan, Sukma Oktavianthi, Lidwina Priliani, Edison Johar, Sri Masyeni, Erly Sintya, Safarina G Malik, Khin Saw |
| EPI_ISL_2101032 | Molecular Biology Laboratory, Faculty Medicine and Health Sciences, Warmadewa University | Eijkman Institute for Molecular Biology, National Agency for Research and Innovation; Molecular Biology Laboratory, Faculty Medicine and Health Sciences, Warmadewa University | Lydia V. Panggalo, Iskandar Adnan, Sukma Oktavianthi, Lidwina Priliani, Edison Johar, Frilasita A Yudhaputri, Muhammad Rezki Rasyak, Willy Agustine, Hidayat Trimarsanto, Sri Masyeni, Erly Sintya, Safarina G Malik, Khin Saw |
| EPI_ISL_2101034 | Molecular Biology Laboratory, Faculty Medicine and Health Sciences, Warmadewa University | Eijkman Institute for Molecular Biology, National Agency for Research and Innovation; Molecular Biology Laboratory, Faculty Medicine and Health Sciences, Warmadewa University | Willy Agustine, Hidayat Trimarsanto, Lydia V. Panggalo, Iskandar Adnan, Sukma Oktavianthi, Lidwina Priliani, Edison Johar, Frilasita A Yudhaputri, Muhammad Rezki Rasyak, Sri Masyeni, Erly Sintya, Safarina G Malik, Khin Saw |
| EPI_ISL_2101035, EPI_ISL_2101036, EPI_ISL_2101037 | Molecular Biology Laboratory, Faculty Medicine and Health Sciences, Warmadewa University | Eijkman Institute for Molecular Biology, National Agency for Research and Innovation; Molecular Biology Laboratory, Faculty Medicine and Health Sciences, Warmadewa University | Lydia V. Panggalo, Iskandar Adnan, Sukma Oktavianthi, Lidwina Priliani, Edison Johar, Frilasita A Yudhaputri, Muhammad Rezki Rasyak, Willy Agustine, Hidayat Trimarsanto, Sri Masyeni, Erly Sintya, Safarina G Malik, Khin Saw |
| EPI_ISL_2101038 | Molecular Biology Laboratory, Faculty Medicine and Health Sciences, Warmadewa University | Eijkman Institute for Molecular Biology, National Agency for Research and Innovation; Molecular Biology Laboratory, Faculty Medicine and Health Sciences, Warmadewa University | Iskandar Adnan, Sukma Oktavianthi, Lidwina Priliani, Edison Johar, Frilasita A Yudhaputri, Muhammad Rezki Rasyak, Willy Agustine, Hidayat Trimarsanto, Lydia V. Panggalo, Sri Masyeni, Erly Sintya, Safarina G Malik, Khin Saw |
| EPI_ISL_2101039 | Molecular Biology Laboratory, Faculty Medicine and Health Sciences, Warmadewa University | Eijkman Institute for Molecular Biology, National Agency for Research and Innovation; Molecular Biology Laboratory, Faculty Medicine and Health Sciences, Warmadewa University | Edison Johar, Frilasita A Yudhaputri, Muhammad Rezki Rasyak, Willy Agustine, Hidayat Trimarsanto, Lydia V. Panggalo, Iskandar Adnan, Sukma Oktavianthi, Lidwina Priliani, Sri Masyeni, Erly Sintya, Safarina G Malik, Khin Saw |
| EPI_ISL_2101040 | Molecular Biology Laboratory, Faculty Medicine and Health Sciences, Warmadewa University | Eijkman Institute for Molecular Biology, National Agency for Research and Innovation; Molecular Biology Laboratory, Faculty Medicine and Health Sciences, Warmadewa University | Hidayat Trimarsanto, Lydia V. Panggalo, Iskandar Adnan, Sukma Oktavianthi, Lidwina Priliani, Edison Johar, Frilasita A Yudhaputri, Muhammad Rezki Rasyak, Willy Agustine, Sri Masyeni, Erly Sintya, Safarina G Malik, Khin Saw |
| EPI_ISL_2101041 | Molecular Biology Laboratory, Faculty Medicine and Health Sciences, Warmadewa University | Eijkman Institute for Molecular Biology, National Agency for Research and Innovation; Molecular Biology Laboratory, Faculty Medicine and Health Sciences, Warmadewa University | Sukma Oktavianthi, Lidwina Priliani, Edison Johar, Frilasita A Yudhaputri, Muhammad Rezki Rasyak, Willy Agustine, Hidayat Trimarsanto, Lydia V. Panggalo, Iskandar Adnan, Sri Masyeni, Erly Sintya, Safarina G Malik, Khin Saw |
| EPI_ISL_2101042 | Molecular Biology Laboratory, Faculty Medicine and Health Sciences, Warmadewa University | Eijkman Institute for Molecular Biology, National Agency for Research and Innovation; Molecular Biology Laboratory, Faculty Medicine and Health Sciences, Warmadewa University | Willy Agustine, Hidayat Trimarsanto, Lydia V. Panggalo, Iskandar Adnan, Sukma Oktavianthi, Lidwina Priliani, Edison Johar, Frilasita A Yudhaputri, Muhammad Rezki Rasyak, Sri Masyeni, Erly Sintya, Safarina G Malik, Khin Saw |
| EPI_ISL_2101043 | Molecular Biology Laboratory, Faculty Medicine and | Eijkman Institute for Molecular Biology, National Agency for Research and Innovation; Molecular Biology Laboratory, Faculty | Hidayat Trimarsanto, Lydia V. Panggalo, Iskandar Adnan, Sukma Oktavianthi, Lidwina Priliani, Edison Johar, Frilasita A Yudhaputri, Muhammad Rezki Rasyak, Willy Agustine, Sri Masyeni, Erly Sintya, Safarina G Malik, Khin Saw |

| | | | |
|---|---|---|---|
| | Health Sciences, Warmadewa University | Medicine and Health Sciences, Warmadewa University | |
| EPI_ISL_2101044 | Molecular Biology Laboratory, Faculty Medicine and Health Sciences, Warmadewa University | Eijkman Institute for Molecular Biology, National Agency for Research and Innovation; Molecular Biology Laboratory, Faculty Medicine and Health Sciences, Warmadewa University | Lydia V. Panggalo, Iskandar Adnan, Sukma Oktavianthi, Lidwina Priliani, Edison Johar, Frilasita A Yudhaputri, Muhammad Rezki Rasyak, Willy Agustine, Hidayat Trimarsanto, Sri Masyeni, Erly Sintya, Safarina G Malik, Khin Saw |
| EPI_ISL_2101045 | Molecular Biology Laboratory, Faculty Medicine and Health Sciences, Warmadewa University | Eijkman Institute for Molecular Biology, National Agency for Research and Innovation; Molecular Biology Laboratory, Faculty Medicine and Health Sciences, Warmadewa University | Iskandar Adnan, Sukma Oktavianthi, Lidwina Priliani, Edison Johar, Frilasita A Yudhaputri, Muhammad Rezki Rasyak, Willy Agustine, Hidayat Trimarsanto, Lydia V. Panggalo, Sri Masyeni, Erly Sintya, Safarina G Malik, Khin Saw |
| EPI_ISL_2101046 | Molecular Biology Laboratory, Faculty Medicine and Health Sciences, Warmadewa University | Eijkman Institute for Molecular Biology, National Agency for Research and Innovation; Molecular Biology Laboratory, Faculty Medicine and Health Sciences, Warmadewa University | Sukma Oktavianthi, Lidwina Priliani, Edison Johar, Frilasita A Yudhaputri, Muhammad Rezki Rasyak, Willy Agustine, Hidayat Trimarsanto, Lydia V. Panggalo, Iskandar Adnan, Sri Masyeni, Erly Sintya, Safarina G Malik, Khin Saw |
| EPI_ISL_2101047 | Molecular Biology Laboratory, Faculty Medicine and Health Sciences, Warmadewa University | Eijkman Institute for Molecular Biology, National Agency for Research and Innovation; Molecular Biology Laboratory, Faculty Medicine and Health Sciences, Warmadewa University | Lidwina Priliani, Edison Johar, Frilasita A Yudhaputri, Muhammad Rezki Rasyak, Willy Agustine, Hidayat Trimarsanto, Lydia V. Panggalo, Iskandar Adnan, Sukma Oktavianthi, Sri Masyeni, Erly Sintya, Safarina G Malik, Khin Saw |
| EPI_ISL_2101048 | Molecular Biology Laboratory, Faculty Medicine and Health Sciences, Warmadewa University | Eijkman Institute for Molecular Biology, National Agency for Research and Innovation; Molecular Biology Laboratory, Faculty Medicine and Health Sciences, Warmadewa University | Edison Johar, Frilasita A Yudhaputri, Muhammad Rezki Rasyak, Willy Agustine, Hidayat Trimarsanto, Lydia V. Panggalo, Iskandar Adnan, Sukma Oktavianthi, Lidwina Priliani, Sri Masyeni, Erly Sintya, Safarina G Malik, Khin Saw |
| EPI_ISL_2101049 | Molecular Biology Laboratory, Faculty Medicine and Health Sciences, Warmadewa University | Eijkman Institute for Molecular Biology, National Agency for Research and Innovation; Molecular Biology Laboratory, Faculty Medicine and Health Sciences, Warmadewa University | Frilasita A Yudhaputri, Muhammad Rezki Rasyak, Willy Agustine, Hidayat Trimarsanto, Lydia V. Panggalo, Iskandar Adnan, Sukma Oktavianthi, Lidwina Priliani, Edison Johar, Sri Masyeni, Erly Sintya, Safarina G Malik, Khin Saw |
| EPI_ISL_2101050 | Molecular Biology Laboratory, Faculty Medicine and Health Sciences, Warmadewa University | Eijkman Institute for Molecular Biology, National Agency for Research and Innovation; Molecular Biology Laboratory, Faculty Medicine and Health Sciences, Warmadewa University | Willy Agustine, Hidayat Trimarsanto, Lydia V. Panggalo, Iskandar Adnan, Sukma Oktavianthi, Lidwina Priliani, Edison Johar, Frilasita A Yudhaputri, Muhammad Rezki Rasyak, Sri Masyeni, Erly Sintya, Safarina G Malik, Khin Saw |
| EPI_ISL_2101051 | Molecular Biology Laboratory, Faculty Medicine and Health Sciences, Warmadewa University | Eijkman Institute for Molecular Biology, National Agency for Research and Innovation; Molecular Biology Laboratory, Faculty Medicine and Health Sciences, Warmadewa University | Hidayat Trimarsanto, Lydia V. Panggalo, Iskandar Adnan, Sukma Oktavianthi, Lidwina Priliani, Edison Johar, Frilasita A Yudhaputri, Muhammad Rezki Rasyak, Willy Agustine, Sri Masyeni, Erly Sintya, Safarina G Malik, Khin Saw |
| EPI_ISL_2101052 | Molecular Biology Laboratory, Faculty Medicine and Health Sciences, Warmadewa University | Eijkman Institute for Molecular Biology, National Agency for Research and Innovation; Molecular Biology Laboratory, Faculty Medicine and Health Sciences, Warmadewa University | Lydia V. Panggalo, Iskandar Adnan, Sukma Oktavianthi, Lidwina Priliani, Edison Johar, Frilasita A Yudhaputri, Muhammad Rezki Rasyak, Willy Agustine, Hidayat Trimarsanto, Sri Masyeni, Erly Sintya, Safarina G Malik, Khin Saw |
| EPI_ISL_2101053 | Molecular Biology Laboratory, Faculty Medicine and Health Sciences, Warmadewa University | Eijkman Institute for Molecular Biology, National Agency for Research and Innovation; Molecular Biology Laboratory, Faculty | Iskandar Adnan, Sukma Oktavianthi, Lidwina Priliani, Edison Johar, Frilasita A Yudhaputri, Muhammad Rezki Rasyak, Willy Agustine, Hidayat Trimarsanto, Lydia V. Panggalo, Sri Masyeni, Erly Sintya, Safarina G Malik, Khin Saw |

| | | | |
|---|---|---|---|
| | | Medicine and Health Sciences, Warmadewa University | |
| EPI_ISL_2101054 | Molecular Biology Laboratory, Faculty Medicine and Health Sciences, Warmadewa University | Eijkman Institute for Molecular Biology, National Agency for Research and Innovation; Molecular Biology Laboratory, Faculty Medicine and Health Sciences, Warmadewa University | Sukma Oktavianthi, Lidwina Priliani, Edison Johar, Frilasita A Yudhaputri, Muhammad Rezki Rasyak, Willy Agustine, Hidayat Trimarsanto, Lydia V. Panggalo, Iskandar Adnan, Sri Masyeni, Erly Sintya, Safarina G Malik, Khin Saw |
| EPI_ISL_2101055 | Molecular Biology Laboratory, Faculty Medicine and Health Sciences, Warmadewa University | Eijkman Institute for Molecular Biology, National Agency for Research and Innovation; Molecular Biology Laboratory, Faculty Medicine and Health Sciences, Warmadewa University | Lidwina Priliani, Edison Johar, Frilasita A Yudhaputri, Muhammad Rezki Rasyak, Willy Agustine, Hidayat Trimarsanto, Lydia V. Panggalo, Iskandar Adnan, Sukma Oktavianthi, Sri Masyeni, Erly Sintya, Safarina G Malik, Khin Saw |
| EPI_ISL_2101056 | Molecular Biology Laboratory, Faculty Medicine and Health Sciences, Warmadewa University | Eijkman Institute for Molecular Biology, National Agency for Research and Innovation; Molecular Biology Laboratory, Faculty Medicine and Health Sciences, Warmadewa University | Edison Johar, Frilasita A Yudhaputri, Muhammad Rezki Rasyak, Willy Agustine, Hidayat Trimarsanto, Lydia V. Panggalo, Iskandar Adnan, Sukma Oktavianthi, Lidwina Priliani, Sri Masyeni, Erly Sintya, Safarina G Malik, Khin Saw |
| EPI_ISL_2101057 | Molecular Biology Laboratory, Faculty Medicine and Health Sciences, Warmadewa University | Eijkman Institute for Molecular Biology, National Agency for Research and Innovation; Molecular Biology Laboratory, Faculty Medicine and Health Sciences, Warmadewa University | Frilasita A Yudhaputri, Muhammad Rezki Rasyak, Willy Agustine, Hidayat Trimarsanto, Lydia V. Panggalo, Iskandar Adnan, Sukma Oktavianthi, Lidwina Priliani, Edison Johar, Sri Masyeni, Erly Sintya, Safarina G Malik, Khin Saw |
| EPI_ISL_2101058 | Molecular Biology Laboratory, Faculty Medicine and Health Sciences, Warmadewa University | Eijkman Institute for Molecular Biology, National Agency for Research and Innovation; Molecular Biology Laboratory, Faculty Medicine and Health Sciences, Warmadewa University | Willy Agustine, Hidayat Trimarsanto, Lydia V. Panggalo, Iskandar Adnan, Sukma Oktavianthi, Lidwina Priliani, Edison Johar, Frilasita A Yudhaputri, Muhammad Rezki Rasyak, Sri Masyeni, Erly Sintya, Safarina G Malik, Khin Saw |
| EPI_ISL_2101059 | Molecular Biology Laboratory, Faculty Medicine and Health Sciences, Warmadewa University | Eijkman Institute for Molecular Biology, National Agency for Research and Innovation; Molecular Biology Laboratory, Faculty Medicine and Health Sciences, Warmadewa University | Hidayat Trimarsanto, Lydia V. Panggalo, Iskandar Adnan, Sukma Oktavianthi, Lidwina Priliani, Edison Johar, Frilasita A Yudhaputri, Muhammad Rezki Rasyak, Willy Agustine, Sri Masyeni, Erly Sintya, Safarina G Malik, Khin Saw |
| EPI_ISL_2101060 | Molecular Biology Laboratory, Faculty Medicine and Health Sciences, Warmadewa University | Eijkman Institute for Molecular Biology, National Agency for Research and Innovation; Molecular Biology Laboratory, Faculty Medicine and Health Sciences, Warmadewa University | Lydia V. Panggalo, Iskandar Adnan, Sukma Oktavianthi, Lidwina Priliani, Edison Johar, Frilasita A Yudhaputri, Muhammad Rezki Rasyak, Willy Agustine, Hidayat Trimarsanto, Sri Masyeni, Erly Sintya, Safarina G Malik, Khin Saw |
| EPI_ISL_2101061 | Molecular Biology Laboratory, Faculty Medicine and Health Sciences, Warmadewa University | Eijkman Institute for Molecular Biology, National Agency for Research and Innovation; Molecular Biology Laboratory, Faculty Medicine and Health Sciences, Warmadewa University | Iskandar Adnan, Sukma Oktavianthi, Lidwina Priliani, Edison Johar, Frilasita A Yudhaputri, Muhammad Rezki Rasyak, Willy Agustine, Hidayat Trimarsanto, Lydia V. Panggalo, Sri Masyeni, Erly Sintya, Safarina G Malik, Khin Saw |
| EPI_ISL_2101062 | Molecular Biology Laboratory, Faculty Medicine and Health Sciences, Warmadewa University | Eijkman Institute for Molecular Biology, National Agency for Research and Innovation; Molecular Biology Laboratory, Faculty Medicine and Health Sciences, Warmadewa University | Sukma Oktavianthi, Lidwina Priliani, Edison Johar, Frilasita A Yudhaputri, Muhammad Rezki Rasyak, Willy Agustine, Hidayat Trimarsanto, Lydia V. Panggalo, Iskandar Adnan, Sri Masyeni, Erly Sintya, Safarina G Malik, Khin Saw |
| EPI_ISL_2101063 | Molecular Biology Laboratory, Faculty Medicine and Health Sciences, Warmadewa University | Eijkman Institute for Molecular Biology, National Agency for Research and Innovation; Molecular Biology Laboratory, Faculty | Lidwina Priliani, Edison Johar, Frilasita A Yudhaputri, Muhammad Rezki Rasyak, Willy Agustine, Hidayat Trimarsanto, Lydia V. Panggalo, Iskandar Adnan, Sukma Oktavianthi, Sri Masyeni, Erly Sintya, Safarina G Malik, Khin Saw |

| | | | |
|---|---|---|---|
| | | Medicine and Health Sciences, Warmadewa University | |
| EPI_ISL_2101064 | Molecular Biology Laboratory, Faculty Medicine and Health Sciences, Warmadewa University | Eijkman Institute for Molecular Biology, National Agency for Research and Innovation; Molecular Biology Laboratory, Faculty Medicine and Health Sciences, Warmadewa University | Edison Johar, Frilasita A Yudhaputri, Muhammad Rezki Rasyak, Willy Agustine, Hidayat Trimarsanto, Lydia V. Panggalo, Iskandar Adnan, Sukma Oktavianthi, Lidwina Priliani, Sri Masyeni, Erly Sintya, Safarina G Malik, Khin Saw |
| EPI_ISL_2101065 | Molecular Biology Laboratory, Faculty Medicine and Health Sciences, Warmadewa University | Eijkman Institute for Molecular Biology, National Agency for Research and Innovation; Molecular Biology Laboratory, Faculty Medicine and Health Sciences, Warmadewa University | Lydia V. Panggalo, Iskandar Adnan, Sukma Oktavianthi, Lidwina Priliani, Edison Johar, Frilasita A Yudhaputri, Muhammad Rezki Rasyak, Willy Agustine, Hidayat Trimarsanto, Sri Masyeni, Erly Sintya, Safarina G Malik, Khin Saw |
| EPI_ISL_2101066 | Molecular Biology Laboratory, Faculty Medicine and Health Sciences, Warmadewa University | Eijkman Institute for Molecular Biology, National Agency for Research and Innovation; Molecular Biology Laboratory, Faculty Medicine and Health Sciences, Warmadewa University | Iskandar Adnan, Sukma Oktavianthi, Lidwina Priliani, Edison Johar, Frilasita A Yudhaputri, Muhammad Rezki Rasyak, Willy Agustine, Hidayat Trimarsanto, Lydia V. Panggalo, Sri Masyeni, Erly Sintya, Safarina G Malik, Khin Saw |
| EPI_ISL_2101067 | Molecular Biology Laboratory, Faculty Medicine and Health Sciences, Warmadewa University | Eijkman Institute for Molecular Biology, National Agency for Research and Innovation; Molecular Biology Laboratory, Faculty Medicine and Health Sciences, Warmadewa University | Sukma Oktavianthi, Lidwina Priliani, Edison Johar, Frilasita A Yudhaputri, Muhammad Rezki Rasyak, Willy Agustine, Hidayat Trimarsanto, Lydia V. Panggalo, Iskandar Adnan, Sri Masyeni, Erly Sintya, Safarina G Malik, Khin Saw |
| EPI_ISL_2101068 | Molecular Biology Laboratory, Faculty Medicine and Health Sciences, Warmadewa University | Eijkman Institute for Molecular Biology, National Agency for Research and Innovation; Molecular Biology Laboratory, Faculty Medicine and Health Sciences, Warmadewa University | Lidwina Priliani, Edison Johar, Frilasita A Yudhaputri, Muhammad Rezki Rasyak, Willy Agustine, Hidayat Trimarsanto, Lydia V. Panggalo, Iskandar Adnan, Sukma Oktavianthi, Sri Masyeni, Erly Sintya, Safarina G Malik, Khin Saw |
| EPI_ISL_2101069 | Molecular Biology Laboratory, Faculty Medicine and Health Sciences, Warmadewa University | Eijkman Institute for Molecular Biology, National Agency for Research and Innovation; Molecular Biology Laboratory, Faculty Medicine and Health Sciences, Warmadewa University | Edison Johar, Frilasita A Yudhaputri, Muhammad Rezki Rasyak, Willy Agustine, Hidayat Trimarsanto, Lydia V. Panggalo, Iskandar Adnan, Sukma Oktavianthi, Lidwina Priliani, Sri Masyeni, Erly Sintya, Safarina G Malik, Khin Saw |
| EPI_ISL_2101070 | Molecular Biology Laboratory, Faculty Medicine and Health Sciences, Warmadewa University | Eijkman Institute for Molecular Biology, National Agency for Research and Innovation; Molecular Biology Laboratory, Faculty Medicine and Health Sciences, Warmadewa University | Frilasita A Yudhaputri, Muhammad Rezki Rasyak, Willy Agustine, Hidayat Trimarsanto, Lydia V. Panggalo, Iskandar Adnan, Sukma Oktavianthi, Lidwina Priliani, Edison Johar, Sri Masyeni, Erly Sintya, Safarina G Malik, Khin Saw |
| EPI_ISL_2101071 | Molecular Biology Laboratory, Faculty Medicine and Health Sciences, Warmadewa University | Eijkman Institute for Molecular Biology, National Agency for Research and Innovation; Molecular Biology Laboratory, Faculty Medicine and Health Sciences, Warmadewa University | Hidayat Trimarsanto, Lydia V. Panggalo, Iskandar Adnan, Sukma Oktavianthi, Lidwina Priliani, Edison Johar, Frilasita A Yudhaputri, Muhammad Rezki Rasyak, Willy Agustine, Sri Masyeni, Erly Sintya, Safarina G Malik, Khin Saw |
| EPI_ISL_2101072 | Molecular Biology Laboratory, Faculty Medicine and Health Sciences, Warmadewa University | Eijkman Institute for Molecular Biology, National Agency for Research and Innovation; Molecular Biology Laboratory, Faculty Medicine and Health Sciences, Warmadewa University | Lydia V. Panggalo, Iskandar Adnan, Sukma Oktavianthi, Lidwina Priliani, Edison Johar, Frilasita A Yudhaputri, Muhammad Rezki Rasyak, Willy Agustine, Hidayat Trimarsanto, Sri Masyeni, Erly Sintya, Safarina G Malik, Khin Saw |
| EPI_ISL_2101073 | Molecular Biology Laboratory, Faculty Medicine and Health Sciences, Warmadewa University | Eijkman Institute for Molecular Biology, National Agency for Research and Innovation; Molecular Biology Laboratory, Faculty | Iskandar Adnan, Sukma Oktavianthi, Lidwina Priliani, Edison Johar, Frilasita A Yudhaputri, Muhammad Rezki Rasyak, Willy Agustine, Hidayat Trimarsanto, Lydia V. Panggalo, Sri Masyeni, Erly Sintya, Safarina G Malik, Khin Saw |

| | | | |
|---|---|---|---|
| | | Medicine and Health Sciences, Warmadewa University | |
| EPI_ISL_2101074 | Molecular Biology Laboratory, Faculty Medicine and Health Sciences, Warmadewa University | Eijkman Institute for Molecular Biology, National Agency for Research and Innovation; Molecular Biology Laboratory, Faculty Medicine and Health Sciences, Warmadewa University | Sukma Oktavianthi, Lidwina Priliani, Edison Johar, Frilasita A Yudhaputri, Muhammad Rezki Rasyak, Willy Agustine, Hidayat Trimarsanto, Lydia V. Panggalo, Iskandar Adnan, Sri Masyeni, Erly Sintya, Safarina G Malik, Khin Saw |
| EPI_ISL_2101075 | Molecular Biology Laboratory, Faculty Medicine and Health Sciences, Warmadewa University | Eijkman Institute for Molecular Biology, National Agency for Research and Innovation; Molecular Biology Laboratory, Faculty Medicine and Health Sciences, Warmadewa University | Lidwina Priliani, Edison Johar, Frilasita A Yudhaputri, Muhammad Rezki Rasyak, Willy Agustine, Hidayat Trimarsanto, Lydia V. Panggalo, Iskandar Adnan, Sukma Oktavianthi, Sri Masyeni, Erly Sintya, Safarina G Malik, Khin Saw |
| EPI_ISL_2101076 | Molecular Biology Laboratory, Faculty Medicine and Health Sciences, Warmadewa University | Eijkman Institute for Molecular Biology, National Agency for Research and Innovation; Molecular Biology Laboratory, Faculty Medicine and Health Sciences, Warmadewa University | Edison Johar, Frilasita A Yudhaputri, Muhammad Rezki Rasyak, Willy Agustine, Hidayat Trimarsanto, Lydia V. Panggalo, Iskandar Adnan, Sukma Oktavianthi, Lidwina Priliani, Sri Masyeni, Erly Sintya, Safarina G Malik, Khin Saw |
| EPI_ISL_2101077 | Molecular Biology Laboratory, Faculty Medicine and Health Sciences, Warmadewa University | Eijkman Institute for Molecular Biology, National Agency for Research and Innovation; Molecular Biology Laboratory, Faculty Medicine and Health Sciences, Warmadewa University | Frilasita A Yudhaputri, Muhammad Rezki Rasyak, Willy Agustine, Hidayat Trimarsanto, Lydia V. Panggalo, Iskandar Adnan, Sukma Oktavianthi, Lidwina Priliani, Edison Johar, Sri Masyeni, Erly Sintya, Safarina G Malik, Khin Saw |
| EPI_ISL_2101078 | Molecular Biology Laboratory, Faculty Medicine and Health Sciences, Warmadewa University | Eijkman Institute for Molecular Biology, National Agency for Research and Innovation; Molecular Biology Laboratory, Faculty Medicine and Health Sciences, Warmadewa University | Willy Agustine, Hidayat Trimarsanto, Lydia V. Panggalo, Iskandar Adnan, Sukma Oktavianthi, Lidwina Priliani, Edison Johar, Frilasita A Yudhaputri, Muhammad Rezki Rasyak, Sri Masyeni, Erly Sintya, Safarina G Malik, Khin Saw |
| EPI_ISL_2101079 | Molecular Biology Laboratory, Faculty Medicine and Health Sciences, Warmadewa University | Eijkman Institute for Molecular Biology, National Agency for Research and Innovation; Molecular Biology Laboratory, Faculty Medicine and Health Sciences, Warmadewa University | Hidayat Trimarsanto, Lydia V. Panggalo, Iskandar Adnan, Sukma Oktavianthi, Lidwina Priliani, Edison Johar, Frilasita A Yudhaputri, Muhammad Rezki Rasyak, Willy Agustine, Sri Masyeni, Erly Sintya, Safarina G Malik, Khin Saw |
| EPI_ISL_2101080 | Molecular Biology Laboratory, Faculty Medicine and Health Sciences, Warmadewa University | Eijkman Institute for Molecular Biology, National Agency for Research and Innovation; Molecular Biology Laboratory, Faculty Medicine and Health Sciences, Warmadewa University | Lydia V. Panggalo, Iskandar Adnan, Sukma Oktavianthi, Lidwina Priliani, Edison Johar, Frilasita A Yudhaputri, Muhammad Rezki Rasyak, Willy Agustine, Hidayat Trimarsanto, Sri Masyeni, Erly Sintya, Safarina G Malik, Khin Saw |
| EPI_ISL_2101081 | Molecular Biology Laboratory, Faculty Medicine and Health Sciences, Warmadewa University | Eijkman Institute for Molecular Biology, National Agency for Research and Innovation; Molecular Biology Laboratory, Faculty Medicine and Health Sciences, Warmadewa University | Sukma Oktavianthi, Lidwina Priliani, Edison Johar, Frilasita A Yudhaputri, Muhammad Rezki Rasyak, Willy Agustine, Hidayat Trimarsanto, Lydia V. Panggalo, Iskandar Adnan, Sri Masyeni, Erly Sintya, Safarina G Malik, Khin Saw |
| EPI_ISL_2101082 | Molecular Biology Laboratory, Faculty Medicine and Health Sciences, Warmadewa University | Eijkman Institute for Molecular Biology, National Agency for Research and Innovation; Molecular Biology Laboratory, Faculty Medicine and Health Sciences, Warmadewa University | Lidwina Priliani, Edison Johar, Frilasita A Yudhaputri, Muhammad Rezki Rasyak, Willy Agustine, Hidayat Trimarsanto, Lydia V. Panggalo, Iskandar Adnan, Sukma Oktavianthi, Sri Masyeni, Erly Sintya, Safarina G Malik, Khin Saw |
| EPI_ISL_2101083 | Molecular Biology Laboratory, Faculty Medicine and Health Sciences, Warmadewa University | Eijkman Institute for Molecular Biology, National Agency for Research and Innovation; Molecular Biology Laboratory, Faculty | Edison Johar, Frilasita A Yudhaputri, Muhammad Rezki Rasyak, Willy Agustine, Hidayat Trimarsanto, Lydia V. Panggalo, Iskandar Adnan, Sukma Oktavianthi, Lidwina Priliani, Sri Masyeni, Erly Sintya, Safarina G Malik, Khin Saw |

| | | | |
|---|---|---|---|
| | | Medicine and Health Sciences, Warmadewa University | |
| EPI_ISL_2101084 | Molecular Biology Laboratory, Faculty Medicine and Health Sciences, Warmadewa University | Eijkman Institute for Molecular Biology, National Agency for Research and Innovation; Molecular Biology Laboratory, Faculty Medicine and Health Sciences, Warmadewa University | Frilasita A Yudhaputri, Muhammad Rezki Rasyak, Willy Agustine, Hidayat Trimarsanto, Lydia V. Panggalo, Iskandar Adnan, Sukma Oktavianthi, Lidwina Priliani, Edison Johar, Sri Masyeni, Erly Sintya, Safarina G Malik, Khin Saw |
| EPI_ISL_2101085 | Molecular Biology Laboratory, Faculty Medicine and Health Sciences, Warmadewa University | Eijkman Institute for Molecular Biology, National Agency for Research and Innovation; Molecular Biology Laboratory, Faculty Medicine and Health Sciences, Warmadewa University | Willy Agustine, Hidayat Trimarsanto, Lydia V. Panggalo, Iskandar Adnan, Sukma Oktavianthi, Lidwina Priliani, Edison Johar, Frilasita A Yudhaputri, Muhammad Rezki Rasyak, Sri Masyeni, Erly Sintya, Safarina G Malik, Khin Saw |
| EPI_ISL_2101086 | Molecular Biology Laboratory, Faculty Medicine and Health Sciences, Warmadewa University | Eijkman Institute for Molecular Biology, National Agency for Research and Innovation; Molecular Biology Laboratory, Faculty Medicine and Health Sciences, Warmadewa University | Hidayat Trimarsanto, Lydia V. Panggalo, Iskandar Adnan, Sukma Oktavianthi, Lidwina Priliani, Edison Johar, Frilasita A Yudhaputri, Muhammad Rezki Rasyak, Willy Agustine, Sri Masyeni, Erly Sintya, Safarina G Malik, Khin Saw |
| EPI_ISL_2101087 | Molecular Biology Laboratory, Faculty Medicine and Health Sciences, Warmadewa University | Eijkman Institute for Molecular Biology, National Agency for Research and Innovation; Molecular Biology Laboratory, Faculty Medicine and Health Sciences, Warmadewa University | Lydia V. Panggalo, Iskandar Adnan, Sukma Oktavianthi, Lidwina Priliani, Edison Johar, Frilasita A Yudhaputri, Muhammad Rezki Rasyak, Willy Agustine, Hidayat Trimarsanto, Sri Masyeni, Erly Sintya, Safarina G Malik, Khin Saw |
| EPI_ISL_2101088 | Molecular Biology Laboratory, Faculty Medicine and Health Sciences, Warmadewa University | Eijkman Institute for Molecular Biology, National Agency for Research and Innovation; Molecular Biology Laboratory, Faculty Medicine and Health Sciences, Warmadewa University | Iskandar Adnan, Sukma Oktavianthi, Lidwina Priliani, Edison Johar, Frilasita A Yudhaputri, Muhammad Rezki Rasyak, Willy Agustine, Hidayat Trimarsanto, Lydia V. Panggalo, Sri Masyeni, Erly Sintya, Safarina G Malik, Khin Saw |
| EPI_ISL_2101089 | Molecular Biology Laboratory, Faculty Medicine and Health Sciences, Warmadewa University | Eijkman Institute for Molecular Biology, National Agency for Research and Innovation; Molecular Biology Laboratory, Faculty Medicine and Health Sciences, Warmadewa University | Sukma Oktavianthi, Lidwina Priliani, Edison Johar, Frilasita A Yudhaputri, Muhammad Rezki Rasyak, Willy Agustine, Hidayat Trimarsanto, Lydia V. Panggalo, Iskandar Adnan, Sri Masyeni, Erly Sintya, Safarina G Malik, Khin Saw |
| EPI_ISL_2101090 | Molecular Biology Laboratory, Faculty Medicine and Health Sciences, Warmadewa University | Eijkman Institute for Molecular Biology, National Agency for Research and Innovation; Molecular Biology Laboratory, Faculty Medicine and Health Sciences, Warmadewa University | Lidwina Priliani, Edison Johar, Frilasita A Yudhaputri, Muhammad Rezki Rasyak, Willy Agustine, Hidayat Trimarsanto, Lydia V. Panggalo, Iskandar Adnan, Sukma Oktavianthi, Sri Masyeni, Erly Sintya, Safarina G Malik, Khin Saw |
| EPI_ISL_2101091 | Molecular Biology Laboratory, Faculty Medicine and Health Sciences, Warmadewa University | Eijkman Institute for Molecular Biology, National Agency for Research and Innovation; Molecular Biology Laboratory, Faculty Medicine and Health Sciences, Warmadewa University | Frilasita A Yudhaputri, Muhammad Rezki Rasyak, Willy Agustine, Hidayat Trimarsanto, Lydia V. Panggalo, Iskandar Adnan, Sukma Oktavianthi, Lidwina Priliani, Edison Johar, Sri Masyeni, Erly Sintya, Safarina G Malik, Khin Saw |
| EPI_ISL_2101092 | Molecular Biology Laboratory, Faculty Medicine and Health Sciences, Warmadewa University | Eijkman Institute for Molecular Biology, National Agency for Research and Innovation; Molecular Biology Laboratory, Faculty Medicine and Health Sciences, Warmadewa University | Willy Agustine, Hidayat Trimarsanto, Lydia V. Panggalo, Iskandar Adnan, Sukma Oktavianthi, Lidwina Priliani, Edison Johar, Frilasita A Yudhaputri, Muhammad Rezki Rasyak, Sri Masyeni, Erly Sintya, Safarina G Malik, Khin Saw |
| EPI_ISL_2101093 | Molecular Biology Laboratory, Faculty Medicine and Health Sciences, Warmadewa University | Eijkman Institute for Molecular Biology, National Agency for Research and Innovation; Molecular Biology Laboratory, Faculty | Hidayat Trimarsanto, Lydia V. Panggalo, Iskandar Adnan, Sukma Oktavianthi, Lidwina Priliani, Edison Johar, Frilasita A Yudhaputri, Muhammad Rezki Rasyak, Willy Agustine, Sri Masyeni, Erly Sintya, Safarina G Malik, Khin Saw |

| | | | |
|---|---|---|---|
| | | Medicine and Health Sciences, Warmadewa University | |
| EPI_ISL_2101094 | Molecular Biology Laboratory, Faculty Medicine and Health Sciences, Warmadewa University | Eijkman Institute for Molecular Biology, National Agency for Research and Innovation; Molecular Biology Laboratory, Faculty Medicine and Health Sciences, Warmadewa University | Lydia V. Panggalo, Iskandar Adnan, Sukma Oktavianthi, Lidwina Priliani, Edison Johar, Frilasita A Yudhaputri, Muhammad Rezki Rasyak, Willy Agustine, Hidayat Trimarsanto, Sri Masyeni, Erly Sintya, Safarina G Malik, Khin Saw |
| EPI_ISL_2101095 | Molecular Biology Laboratory, Faculty Medicine and Health Sciences, Warmadewa University | Eijkman Institute for Molecular Biology, National Agency for Research and Innovation; Molecular Biology Laboratory, Faculty Medicine and Health Sciences, Warmadewa University | Iskandar Adnan, Sukma Oktavianthi, Lidwina Priliani, Edison Johar, Frilasita A Yudhaputri, Muhammad Rezki Rasyak, Willy Agustine, Hidayat Trimarsanto, Lydia V. Panggalo, Sri Masyeni, Erly Sintya, Safarina G Malik, Khin Saw |
| EPI_ISL_2101096 | Molecular Biology Laboratory, Faculty Medicine and Health Sciences, Warmadewa University | Eijkman Institute for Molecular Biology, National Agency for Research and Innovation; Molecular Biology Laboratory, Faculty Medicine and Health Sciences, Warmadewa University | Lidwina Priliani, Edison Johar, Frilasita A Yudhaputri, Muhammad Rezki Rasyak, Willy Agustine, Hidayat Trimarsanto, Lydia V. Panggalo, Iskandar Adnan, Sukma Oktavianthi, Sri Masyeni, Erly Sintya, Safarina G Malik, Khin Saw |
| EPI_ISL_2101097 | Molecular Biology Laboratory, Faculty Medicine and Health Sciences, Warmadewa University | Eijkman Institute for Molecular Biology, National Agency for Research and Innovation; Molecular Biology Laboratory, Faculty Medicine and Health Sciences, Warmadewa University | Edison Johar, Frilasita A Yudhaputri, Muhammad Rezki Rasyak, Willy Agustine, Hidayat Trimarsanto, Lydia V. Panggalo, Iskandar Adnan, Sukma Oktavianthi, Lidwina Priliani, Sri Masyeni, Erly Sintya, Safarina G Malik, Khin Saw |
| EPI_ISL_2101098 | Molecular Biology Laboratory, Faculty Medicine and Health Sciences, Warmadewa University | Eijkman Institute for Molecular Biology, National Agency for Research and Innovation; Molecular Biology Laboratory, Faculty Medicine and Health Sciences, Warmadewa University | Frilasita A Yudhaputri, Muhammad Rezki Rasyak, Willy Agustine, Hidayat Trimarsanto, Lydia V. Panggalo, Iskandar Adnan, Sukma Oktavianthi, Lidwina Priliani, Edison Johar, Sri Masyeni, Erly Sintya, Safarina G Malik, Khin Saw |
| EPI_ISL_2101099 | Molecular Biology Laboratory, Faculty Medicine and Health Sciences, Warmadewa University | Eijkman Institute for Molecular Biology, National Agency for Research and Innovation; Molecular Biology Laboratory, Faculty Medicine and Health Sciences, Warmadewa University | Willy Agustine, Hidayat Trimarsanto, Lydia V. Panggalo, Iskandar Adnan, Sukma Oktavianthi, Lidwina Priliani, Edison Johar, Frilasita A Yudhaputri, Muhammad Rezki Rasyak, Sri Masyeni, Erly Sintya, Safarina G Malik, Khin Saw |
| EPI_ISL_2101100 | Molecular Biology Laboratory, Faculty Medicine and Health Sciences, Warmadewa University | Eijkman Institute for Molecular Biology, National Agency for Research and Innovation; Molecular Biology Laboratory, Faculty Medicine and Health Sciences, Warmadewa University | Lydia V. Panggalo, Iskandar Adnan, Sukma Oktavianthi, Lidwina Priliani, Edison Johar, Frilasita A Yudhaputri, Muhammad Rezki Rasyak, Willy Agustine, Hidayat Trimarsanto, Sri Masyeni, Erly Sintya, Safarina G Malik, Khin Saw |
| EPI_ISL_2101102 | Molecular Biology Laboratory, Faculty Medicine and Health Sciences, Warmadewa University | Eijkman Institute for Molecular Biology, National Agency for Research and Innovation; Molecular Biology Laboratory, Faculty Medicine and Health Sciences, Warmadewa University | Sukma Oktavianthi, Lidwina Priliani, Edison Johar, Frilasita A Yudhaputri, Muhammad Rezki Rasyak, Willy Agustine, Hidayat Trimarsanto, Lydia V. Panggalo, Iskandar Adnan, Sri Masyeni, Erly Sintya, Safarina G Malik, Khin Saw |
| EPI_ISL_2101103 | Molecular Biology Laboratory, Faculty Medicine and Health Sciences, Warmadewa University | Eijkman Institute for Molecular Biology, National Agency for Research and Innovation; Molecular Biology Laboratory, Faculty Medicine and Health Sciences, Warmadewa University | Lidwina Priliani, Edison Johar, Frilasita A Yudhaputri, Muhammad Rezki Rasyak, Willy Agustine, Hidayat Trimarsanto, Lydia V. Panggalo, Iskandar Adnan, Sukma Oktavianthi, Sri Masyeni, Erly Sintya, Safarina G Malik, Khin Saw |
| EPI_ISL_2101104 | Molecular Biology Laboratory, Faculty Medicine and Health Sciences, Warmadewa University | Eijkman Institute for Molecular Biology, National Agency for Research and Innovation; Molecular Biology Laboratory, Faculty | Edison Johar, Frilasita A Yudhaputri, Muhammad Rezki Rasyak, Willy Agustine, Hidayat Trimarsanto, Lydia V. Panggalo, Iskandar Adnan, Sukma Oktavianthi, Lidwina Priliani, Sri Masyeni, Erly Sintya, Safarina G Malik, Khin Saw |

| | | | |
|---|---|---|---|
| | | Medicine and Health Sciences, Warmadewa University | |
| EPI_ISL_2101105 | Molecular Biology Laboratory, Faculty Medicine and Health Sciences, Warmadewa University | Eijkman Institute for Molecular Biology, National Agency for Research and Innovation; Molecular Biology Laboratory, Faculty Medicine and Health Sciences, Warmadewa University | Frilasita A Yudhaputri, Muhammad Rezki Rasyak, Willy Agustine, Hidayat Trimarsanto, Lydia V. Panggalo, Iskandar Adnan, Sukma Oktavianthi, Lidwina Priliani, Edison Johar, Sri Masyeni, Erly Sintya, Safarina G Malik, Khin Saw |
| EPI_ISL_2101106 | Molecular Biology Laboratory, Faculty Medicine and Health Sciences, Warmadewa University | Eijkman Institute for Molecular Biology, National Agency for Research and Innovation; Molecular Biology Laboratory, Faculty Medicine and Health Sciences, Warmadewa University | Willy Agustine, Hidayat Trimarsanto, Lydia V. Panggalo, Iskandar Adnan, Sukma Oktavianthi, Lidwina Priliani, Edison Johar, Frilasita A Yudhaputri, Muhammad Rezki Rasyak, Sri Masyeni, Erly Sintya, Safarina G Malik, Khin Saw |
| EPI_ISL_2101107 | Molecular Biology Laboratory, Faculty Medicine and Health Sciences, Warmadewa University | Eijkman Institute for Molecular Biology, National Agency for Research and Innovation; Molecular Biology Laboratory, Faculty Medicine and Health Sciences, Warmadewa University | Hidayat Trimarsanto, Lydia V. Panggalo, Iskandar Adnan, Sukma Oktavianthi, Lidwina Priliani, Edison Johar, Frilasita A Yudhaputri, Muhammad Rezki Rasyak, Willy Agustine, Sri Masyeni, Erly Sintya, Safarina G Malik, Khin Saw |
| EPI_ISL_2101108 | Molecular Biology Laboratory, Faculty Medicine and Health Sciences, Warmadewa University | Eijkman Institute for Molecular Biology, National Agency for Research and Innovation; Molecular Biology Laboratory, Faculty Medicine and Health Sciences, Warmadewa University | Lydia V. Panggalo, Iskandar Adnan, Sukma Oktavianthi, Lidwina Priliani, Edison Johar, Frilasita A Yudhaputri, Muhammad Rezki Rasyak, Willy Agustine, Hidayat Trimarsanto, Sri Masyeni, Erly Sintya, Safarina G Malik, Khin Saw |
| EPI_ISL_2101109 | Molecular Biology Laboratory, Faculty Medicine and Health Sciences, Warmadewa University | Eijkman Institute for Molecular Biology, National Agency for Research and Innovation; Molecular Biology Laboratory, Faculty Medicine and Health Sciences, Warmadewa University | Iskandar Adnan, Sukma Oktavianthi, Lidwina Priliani, Edison Johar, Frilasita A Yudhaputri, Muhammad Rezki Rasyak, Willy Agustine, Hidayat Trimarsanto, Lydia V. Panggalo, Sri Masyeni, Erly Sintya, Safarina G Malik, Khin Saw |
| EPI_ISL_2101110 | Molecular Biology Laboratory, Faculty Medicine and Health Sciences, Warmadewa University | Eijkman Institute for Molecular Biology, National Agency for Research and Innovation; Molecular Biology Laboratory, Faculty Medicine and Health Sciences, Warmadewa University | Lidwina Priliani, Edison Johar, Frilasita A Yudhaputri, Muhammad Rezki Rasyak, Willy Agustine, Hidayat Trimarsanto, Lydia V. Panggalo, Iskandar Adnan, Sukma Oktavianthi, Sri Masyeni, Erly Sintya, Safarina G Malik, Khin Saw |
| EPI_ISL_2101111 | Molecular Biology Laboratory, Faculty Medicine and Health Sciences, Warmadewa University | Eijkman Institute for Molecular Biology, National Agency for Research and Innovation; Molecular Biology Laboratory, Faculty Medicine and Health Sciences, Warmadewa University | Edison Johar, Frilasita A Yudhaputri, Muhammad Rezki Rasyak, Willy Agustine, Hidayat Trimarsanto, Lydia V. Panggalo, Iskandar Adnan, Sukma Oktavianthi, Lidwina Priliani, Sri Masyeni, Erly Sintya, Safarina G Malik, Khin Saw |
| EPI_ISL_2101113 | Molecular Biology Laboratory, Faculty Medicine and Health Sciences, Warmadewa University | Eijkman Institute for Molecular Biology, National Agency for Research and Innovation; Molecular Biology Laboratory, Faculty Medicine and Health Sciences, Warmadewa University | Frilasita A Yudhaputri, Muhammad Rezki Rasyak, Willy Agustine, Hidayat Trimarsanto, Lydia V. Panggalo, Iskandar Adnan, Sukma Oktavianthi, Lidwina Priliani, Edison Johar, Sri Masyeni, Erly Sintya, Safarina G Malik, Khin Saw |
| EPI_ISL_2101114 | Molecular Biology Laboratory, Faculty Medicine and Health Sciences, Warmadewa University | Eijkman Institute for Molecular Biology, National Agency for Research and Innovation; Molecular Biology Laboratory, Faculty Medicine and Health Sciences, Warmadewa University | Willy Agustine, Hidayat Trimarsanto, Lydia V. Panggalo, Iskandar Adnan, Sukma Oktavianthi, Lidwina Priliani, Edison Johar, Frilasita A Yudhaputri, Muhammad Rezki Rasyak, Sri Masyeni, Erly Sintya, Safarina G Malik, Khin Saw |
| EPI_ISL_2101115 | Molecular Biology Laboratory, Faculty Medicine and Health Sciences, Warmadewa University | Eijkman Institute for Molecular Biology, National Agency for Research and Innovation; Molecular Biology Laboratory, Faculty | Hidayat Trimarsanto, Lydia V. Panggalo, Iskandar Adnan, Sukma Oktavianthi, Lidwina Priliani, Edison Johar, Frilasita A Yudhaputri, Muhammad Rezki Rasyak, Willy Agustine, Sri Masyeni, Erly Sintya, Safarina G Malik, Khin Saw |

| | | | |
|---|---|---|---|
| | | Medicine and Health Sciences, Warmadewa University | |
| EPI_ISL_2101116 | Molecular Biology Laboratory, Faculty Medicine and Health Sciences, Warmadewa University | Eijkman Institute for Molecular Biology, National Agency for Research and Innovation; Molecular Biology Laboratory, Faculty Medicine and Health Sciences, Warmadewa University | Lydia V. Panggalo, Iskandar Adnan, Sukma Oktavianthi, Lidwina Priliani, Edison Johar, Frilasita A Yudhaputri, Muhammad Rezki Rasyak, Willy Agustine, Hidayat Trimarsanto, Sri Masyeni, Erly Sintya, Safarina G Malik, Khin Saw |
| EPI_ISL_2101117 | Molecular Biology Laboratory, Faculty Medicine and Health Sciences, Warmadewa University | Eijkman Institute for Molecular Biology, National Agency for Research and Innovation; Molecular Biology Laboratory, Faculty Medicine and Health Sciences, Warmadewa University | Iskandar Adnan, Sukma Oktavianthi, Lidwina Priliani, Edison Johar, Frilasita A Yudhaputri, Muhammad Rezki Rasyak, Willy Agustine, Hidayat Trimarsanto, Lydia V. Panggalo, Sri Masyeni, Erly Sintya, Safarina G Malik, Khin Saw |
| EPI_ISL_2101118 | Molecular Biology Laboratory, Faculty Medicine and Health Sciences, Warmadewa University | Eijkman Institute for Molecular Biology, National Agency for Research and Innovation; Molecular Biology Laboratory, Faculty Medicine and Health Sciences, Warmadewa University | Sukma Oktavianthi, Lidwina Priliani, Edison Johar, Frilasita A Yudhaputri, Muhammad Rezki Rasyak, Willy Agustine, Hidayat Trimarsanto, Lydia V. Panggalo, Iskandar Adnan, Sri Masyeni, Erly Sintya, Safarina G Malik, Khin Saw |
| EPI_ISL_2101119 | Molecular Biology Laboratory, Faculty Medicine and Health Sciences, Warmadewa University | Eijkman Institute for Molecular Biology, National Agency for Research and Innovation; Molecular Biology Laboratory, Faculty Medicine and Health Sciences, Warmadewa University | Lidwina Priliani, Edison Johar, Frilasita A Yudhaputri, Muhammad Rezki Rasyak, Willy Agustine, Hidayat Trimarsanto, Lydia V. Panggalo, Iskandar Adnan, Sukma Oktavianthi, Sri Masyeni, Erly Sintya, Safarina G Malik, Khin Saw |
| EPI_ISL_2101120 | Molecular Biology Laboratory, Faculty Medicine and Health Sciences, Warmadewa University | Eijkman Institute for Molecular Biology, National Agency for Research and Innovation; Molecular Biology Laboratory, Faculty Medicine and Health Sciences, Warmadewa University | Edison Johar, Frilasita A Yudhaputri, Muhammad Rezki Rasyak, Willy Agustine, Hidayat Trimarsanto, Lydia V. Panggalo, Iskandar Adnan, Sukma Oktavianthi, Lidwina Priliani, Sri Masyeni, Erly Sintya, Safarina G Malik, Khin Saw |
| EPI_ISL_2101121 | Molecular Biology Laboratory, Faculty Medicine and Health Sciences, Warmadewa University | Eijkman Institute for Molecular Biology, National Agency for Research and Innovation; Molecular Biology Laboratory, Faculty Medicine and Health Sciences, Warmadewa University | Willy Agustine, Hidayat Trimarsanto, Lydia V. Panggalo, Iskandar Adnan, Sukma Oktavianthi, Lidwina Priliani, Edison Johar, Frilasita A Yudhaputri, Muhammad Rezki Rasyak, Sri Masyeni, Erly Sintya, Safarina G Malik, Khin Saw |
| EPI_ISL_2101122 | Molecular Biology Laboratory, Faculty Medicine and Health Sciences, Warmadewa University | Eijkman Institute for Molecular Biology, National Agency for Research and Innovation; Molecular Biology Laboratory, Faculty Medicine and Health Sciences, Warmadewa University | Hidayat Trimarsanto, Lydia V. Panggalo, Iskandar Adnan, Sukma Oktavianthi, Lidwina Priliani, Edison Johar, Frilasita A Yudhaputri, Muhammad Rezki Rasyak, Willy Agustine, Sri Masyeni, Erly Sintya, Safarina G Malik, Khin Saw |
| EPI_ISL_2101123 | Molecular Biology Laboratory, Faculty Medicine and Health Sciences, Warmadewa University | Eijkman Institute for Molecular Biology, National Agency for Research and Innovation; Molecular Biology Laboratory, Faculty Medicine and Health Sciences, Warmadewa University | Lydia V. Panggalo, Iskandar Adnan, Sukma Oktavianthi, Lidwina Priliani, Edison Johar, Frilasita A Yudhaputri, Muhammad Rezki Rasyak, Willy Agustine, Hidayat Trimarsanto, Sri Masyeni, Erly Sintya, Safarina G Malik, Khin Saw |
| EPI_ISL_2101125 | Molecular Biology Laboratory, Faculty Medicine and Health Sciences, Warmadewa University | Eijkman Institute for Molecular Biology, National Agency for Research and Innovation; Molecular Biology Laboratory, Faculty Medicine and Health Sciences, Warmadewa University | Willy Agustine, Hidayat Trimarsanto, Lydia V. Panggalo, Iskandar Adnan, Sukma Oktavianthi, Lidwina Priliani, Edison Johar, Frilasita A Yudhaputri, Muhammad Rezki Rasyak, Sri Masyeni, Erly Sintya, Safarina G Malik, Khin Saw |
| EPI_ISL_2101126 | Molecular Biology Laboratory, Faculty Medicine and Health Sciences, Warmadewa University | Eijkman Institute for Molecular Biology, National Agency for Research and Innovation; Molecular Biology Laboratory, Faculty | Hidayat Trimarsanto, Lydia V. Panggalo, Iskandar Adnan, Sukma Oktavianthi, Lidwina Priliani, Edison Johar, Frilasita A Yudhaputri, Muhammad Rezki Rasyak, Willy Agustine, Sri Masyeni, Erly Sintya, Safarina G Malik, Khin Saw |

| | | | |
|---|---|---|---|
| | | Medicine and Health Sciences, Warmadewa University | |
| EPI_ISL_2101127 | Molecular Biology Laboratory, Faculty Medicine and Health Sciences, Warmadewa University | Eijkman Institute for Molecular Biology, National Agency for Research and Innovation; Molecular Biology Laboratory, Faculty Medicine and Health Sciences, Warmadewa University | Iskandar Adnan, Sukma Oktavianthi, Lidwina Priliani, Edison Johar, Frilasita A Yudhaputri, Muhammad Rezki Rasyak, Willy Agustine, Hidayat Trimarsanto, Lydia V. Panggalo, Sri Masyeni, Erly Sintya, Safarina G Malik, Khin Saw |
| EPI_ISL_2101128 | Molecular Biology Laboratory, Faculty Medicine and Health Sciences, Warmadewa University | Eijkman Institute for Molecular Biology, National Agency for Research and Innovation; Molecular Biology Laboratory, Faculty Medicine and Health Sciences, Warmadewa University | Sukma Oktavianthi, Lidwina Priliani, Edison Johar, Frilasita A Yudhaputri, Muhammad Rezki Rasyak, Willy Agustine, Hidayat Trimarsanto, Lydia V. Panggalo, Iskandar Adnan, Sri Masyeni, Erly Sintya, Safarina G Malik, Khin Saw |
| EPI_ISL_2101129 | Molecular Biology Laboratory, Faculty Medicine and Health Sciences, Warmadewa University | Eijkman Institute for Molecular Biology, National Agency for Research and Innovation; Molecular Biology Laboratory, Faculty Medicine and Health Sciences, Warmadewa University | Edison Johar, Frilasita A Yudhaputri, Muhammad Rezki Rasyak, Willy Agustine, Hidayat Trimarsanto, Lydia V. Panggalo, Iskandar Adnan, Sukma Oktavianthi, Lidwina Priliani, Sri Masyeni, Erly Sintya, Safarina G Malik, Khin Saw |
| EPI_ISL_2101130 | Molecular Biology Laboratory, Faculty Medicine and Health Sciences, Warmadewa University | Eijkman Institute for Molecular Biology, National Agency for Research and Innovation; Molecular Biology Laboratory, Faculty Medicine and Health Sciences, Warmadewa University | Frilasita A Yudhaputri, Muhammad Rezki Rasyak, Willy Agustine, Hidayat Trimarsanto, Lydia V. Panggalo, Iskandar Adnan, Sukma Oktavianthi, Lidwina Priliani, Edison Johar, Sri Masyeni, Erly Sintya, Safarina G Malik, Khin Saw |
| EPI_ISL_2101131 | Molecular Biology Laboratory, Faculty Medicine and Health Sciences, Warmadewa University | Eijkman Institute for Molecular Biology, National Agency for Research and Innovation; Molecular Biology Laboratory, Faculty Medicine and Health Sciences, Warmadewa University | Willy Agustine, Hidayat Trimarsanto, Lydia V. Panggalo, Iskandar Adnan, Sukma Oktavianthi, Lidwina Priliani, Edison Johar, Frilasita A Yudhaputri, Muhammad Rezki Rasyak, Sri Masyeni, Erly Sintya, Safarina G Malik, Khin Saw |
| EPI_ISL_2101132 | Molecular Biology Laboratory, Faculty Medicine and Health Sciences, Warmadewa University | Eijkman Institute for Molecular Biology, National Agency for Research and Innovation; Molecular Biology Laboratory, Faculty Medicine and Health Sciences, Warmadewa University | Hidayat Trimarsanto, Lydia V. Panggalo, Iskandar Adnan, Sukma Oktavianthi, Lidwina Priliani, Edison Johar, Frilasita A Yudhaputri, Muhammad Rezki Rasyak, Willy Agustine, Sri Masyeni, Erly Sintya, Safarina G Malik, Khin Saw |
| EPI_ISL_2101133 | Molecular Biology Laboratory, Faculty Medicine and Health Sciences, Warmadewa University | Eijkman Institute for Molecular Biology, National Agency for Research and Innovation; Molecular Biology Laboratory, Faculty Medicine and Health Sciences, Warmadewa University | Lydia V. Panggalo, Iskandar Adnan, Sukma Oktavianthi, Lidwina Priliani, Edison Johar, Frilasita A Yudhaputri, Muhammad Rezki Rasyak, Willy Agustine, Hidayat Trimarsanto, Sri Masyeni, Erly Sintya, Safarina G Malik, Khin Saw |
| EPI_ISL_2101134 | Molecular Biology Laboratory, Faculty Medicine and Health Sciences, Warmadewa University | Eijkman Institute for Molecular Biology, National Agency for Research and Innovation; Molecular Biology Laboratory, Faculty Medicine and Health Sciences, Warmadewa University | Iskandar Adnan, Sukma Oktavianthi, Lidwina Priliani, Edison Johar, Frilasita A Yudhaputri, Muhammad Rezki Rasyak, Willy Agustine, Hidayat Trimarsanto, Lydia V. Panggalo, Sri Masyeni, Erly Sintya, Safarina G Malik, Khin Saw |
| EPI_ISL_2101135 | Molecular Biology Laboratory, Faculty Medicine and Health Sciences, Warmadewa University | Eijkman Institute for Molecular Biology, National Agency for Research and Innovation; Molecular Biology Laboratory, Faculty Medicine and Health Sciences, Warmadewa University | Sukma Oktavianthi, Lidwina Priliani, Edison Johar, Frilasita A Yudhaputri, Muhammad Rezki Rasyak, Willy Agustine, Hidayat Trimarsanto, Lydia V. Panggalo, Iskandar Adnan, Sri Masyeni, Erly Sintya, Safarina G Malik, Khin Saw |
| EPI_ISL_2101136 | Molecular Biology Laboratory, Faculty Medicine and Health Sciences, Warmadewa University | Eijkman Institute for Molecular Biology, National Agency for Research and Innovation; Molecular Biology Laboratory, Faculty | Edison Johar, Frilasita A Yudhaputri, Muhammad Rezki Rasyak, Willy Agustine, Hidayat Trimarsanto, Lydia V. Panggalo, Iskandar Adnan, Sukma Oktavianthi, Lidwina Priliani, Sri Masyeni, Erly Sintya, Safarina G Malik, Khin Saw |

| | | | |
|---|---|---|---|
| | | Medicine and Health Sciences, Warmadewa University | |
| EPI_ISL_2101137 | Molecular Biology Laboratory, Faculty Medicine and Health Sciences, Warmadewa University | Eijkman Institute for Molecular Biology, National Agency for Research and Innovation; Molecular Biology Laboratory, Faculty Medicine and Health Sciences, Warmadewa University | Frilasita A Yudhaputri, Muhammad Rezki Rasyak, Willy Agustine, Hidayat Trimarsanto, Lydia V. Panggalo, Iskandar Adnan, Sukma Oktavianthi, Lidwina Priliani, Edison Johar, Sri Masyeni, Erly Sintya, Safarina G Malik, Khin Saw |
| EPI_ISL_2101138 | Molecular Biology Laboratory, Faculty Medicine and Health Sciences, Warmadewa University | Eijkman Institute for Molecular Biology, National Agency for Research and Innovation; Molecular Biology Laboratory, Faculty Medicine and Health Sciences, Warmadewa University | Willy Agustine, Hidayat Trimarsanto, Lydia V. Panggalo, Iskandar Adnan, Sukma Oktavianthi, Lidwina Priliani, Edison Johar, Frilasita A Yudhaputri, Muhammad Rezki Rasyak, Sri Masyeni, Erly Sintya, Safarina G Malik, Khin Saw |
| EPI_ISL_2101139 | Molecular Biology Laboratory, Faculty Medicine and Health Sciences, Warmadewa University | Eijkman Institute for Molecular Biology, National Agency for Research and Innovation; Molecular Biology Laboratory, Faculty Medicine and Health Sciences, Warmadewa University | Hidayat Trimarsanto, Lydia V. Panggalo, Iskandar Adnan, Sukma Oktavianthi, Lidwina Priliani, Edison Johar, Frilasita A Yudhaputri, Muhammad Rezki Rasyak, Willy Agustine, Sri Masyeni, Erly Sintya, Safarina G Malik, Khin Saw |
| EPI_ISL_2101140 | Molecular Biology Laboratory, Faculty Medicine and Health Sciences, Warmadewa University | Eijkman Institute for Molecular Biology, National Agency for Research and Innovation; Molecular Biology Laboratory, Faculty Medicine and Health Sciences, Warmadewa University | Lydia V. Panggalo, Iskandar Adnan, Sukma Oktavianthi, Lidwina Priliani, Edison Johar, Frilasita A Yudhaputri, Muhammad Rezki Rasyak, Willy Agustine, Hidayat Trimarsanto, Sri Masyeni, Erly Sintya, Safarina G Malik, Khin Saw |
| EPI_ISL_2101141 | Molecular Biology Laboratory, Faculty Medicine and Health Sciences, Warmadewa University | Eijkman Institute for Molecular Biology, National Agency for Research and Innovation; Molecular Biology Laboratory, Faculty Medicine and Health Sciences, Warmadewa University | Iskandar Adnan, Sukma Oktavianthi, Lidwina Priliani, Edison Johar, Frilasita A Yudhaputri, Muhammad Rezki Rasyak, Willy Agustine, Hidayat Trimarsanto, Lydia V. Panggalo, Sri Masyeni, Erly Sintya, Safarina G Malik, Khin Saw |
| EPI_ISL_2101142 | Molecular Biology Laboratory, Faculty Medicine and Health Sciences, Warmadewa University | Eijkman Institute for Molecular Biology, National Agency for Research and Innovation; Molecular Biology Laboratory, Faculty Medicine and Health Sciences, Warmadewa University | Lidwina Priliani, Edison Johar, Frilasita A Yudhaputri, Muhammad Rezki Rasyak, Willy Agustine, Hidayat Trimarsanto, Lydia V. Panggalo, Iskandar Adnan, Sukma Oktavianthi, Sri Masyeni, Erly Sintya, Safarina G Malik, Khin Saw |
| EPI_ISL_2101143 | Molecular Biology Laboratory, Faculty Medicine and Health Sciences, Warmadewa University | Eijkman Institute for Molecular Biology, National Agency for Research and Innovation; Molecular Biology Laboratory, Faculty Medicine and Health Sciences, Warmadewa University | Frilasita A Yudhaputri, Muhammad Rezki Rasyak, Willy Agustine, Hidayat Trimarsanto, Lydia V. Panggalo, Iskandar Adnan, Sukma Oktavianthi, Lidwina Priliani, Edison Johar, Sri Masyeni, Erly Sintya, Safarina G Malik, Khin Saw |
| EPI_ISL_2101144 | Molecular Biology Laboratory, Faculty Medicine and Health Sciences, Warmadewa University | Eijkman Institute for Molecular Biology, National Agency for Research and Innovation; Molecular Biology Laboratory, Faculty Medicine and Health Sciences, Warmadewa University | Hidayat Trimarsanto, Lydia V. Panggalo, Iskandar Adnan, Sukma Oktavianthi, Lidwina Priliani, Edison Johar, Frilasita A Yudhaputri, Muhammad Rezki Rasyak, Willy Agustine, Sri Masyeni, Erly Sintya, Safarina G Malik, Khin Saw |
| EPI_ISL_2101145 | Molecular Biology Laboratory, Faculty Medicine and Health Sciences, Warmadewa University | Eijkman Institute for Molecular Biology, National Agency for Research and Innovation; Molecular Biology Laboratory, Faculty Medicine and Health Sciences, Warmadewa University | Lydia V. Panggalo, Iskandar Adnan, Sukma Oktavianthi, Lidwina Priliani, Edison Johar, Frilasita A Yudhaputri, Muhammad Rezki Rasyak, Willy Agustine, Hidayat Trimarsanto, Sri Masyeni, Erly Sintya, Safarina G Malik, Khin Saw |
| EPI_ISL_2101146 | Molecular Biology Laboratory, Faculty Medicine and Health Sciences, Warmadewa University | Eijkman Institute for Molecular Biology, National Agency for Research and Innovation; Molecular Biology Laboratory, Faculty | Iskandar Adnan, Sukma Oktavianthi, Lidwina Priliani, Edison Johar, Frilasita A Yudhaputri, Muhammad Rezki Rasyak, Willy Agustine, Hidayat Trimarsanto, Lydia V. Panggalo, Sri Masyeni, Erly Sintya, Safarina G Malik, Khin Saw |

| | | | |
|---|---|---|---|
| | | Medicine and Health Sciences, Warmadewa University | |
| EPI_ISL_2101147 | Molecular Biology Laboratory, Faculty Medicine and Health Sciences, Warmadewa University | Eijkman Institute for Molecular Biology, National Agency for Research and Innovation; Molecular Biology Laboratory, Faculty Medicine and Health Sciences, Warmadewa University | Sukma Oktavianthi, Lidwina Priliani, Edison Johar, Frilasita A Yudhaputri, Muhammad Rezki Rasyak, Willy Agustine, Hidayat Trimarsanto, Lydia V. Panggalo, Iskandar Adnan, Sri Masyeni, Erly Sintya, Safarina G Malik, Khin Saw |
| EPI_ISL_2101148 | Molecular Biology Laboratory, Faculty Medicine and Health Sciences, Warmadewa University | Eijkman Institute for Molecular Biology, National Agency for Research and Innovation; Molecular Biology Laboratory, Faculty Medicine and Health Sciences, Warmadewa University | Lidwina Priliani, Edison Johar, Frilasita A Yudhaputri, Muhammad Rezki Rasyak, Willy Agustine, Hidayat Trimarsanto, Lydia V. Panggalo, Iskandar Adnan, Sukma Oktavianthi, Sri Masyeni, Erly Sintya, Safarina G Malik, Khin Saw |
| EPI_ISL_2101149 | Molecular Biology Laboratory, Faculty Medicine and Health Sciences, Warmadewa University | Eijkman Institute for Molecular Biology, National Agency for Research and Innovation; Molecular Biology Laboratory, Faculty Medicine and Health Sciences, Warmadewa University | Edison Johar, Frilasita A Yudhaputri, Muhammad Rezki Rasyak, Willy Agustine, Hidayat Trimarsanto, Lydia V. Panggalo, Iskandar Adnan, Sukma Oktavianthi, Lidwina Priliani, Sri Masyeni, Erly Sintya, Safarina G Malik, Khin Saw |
| EPI_ISL_2101150 | Molecular Biology Laboratory, Faculty Medicine and Health Sciences, Warmadewa University | Eijkman Institute for Molecular Biology, National Agency for Research and Innovation; Molecular Biology Laboratory, Faculty Medicine and Health Sciences, Warmadewa University | Frilasita A Yudhaputri, Muhammad Rezki Rasyak, Willy Agustine, Hidayat Trimarsanto, Lydia V. Panggalo, Iskandar Adnan, Sukma Oktavianthi, Lidwina Priliani, Edison Johar, Sri Masyeni, Erly Sintya, Safarina G Malik, Khin Saw |
| EPI_ISL_2101151 | Molecular Biology Laboratory, Faculty Medicine and Health Sciences, Warmadewa University | Eijkman Institute for Molecular Biology, National Agency for Research and Innovation; Molecular Biology Laboratory, Faculty Medicine and Health Sciences, Warmadewa University | Willy Agustine, Hidayat Trimarsanto, Lydia V. Panggalo, Iskandar Adnan, Sukma Oktavianthi, Lidwina Priliani, Edison Johar, Frilasita A Yudhaputri, Muhammad Rezki Rasyak, Sri Masyeni, Erly Sintya, Safarina G Malik, Khin Saw |
| EPI_ISL_2101152 | Molecular Biology Laboratory, Faculty Medicine and Health Sciences, Warmadewa University | Eijkman Institute for Molecular Biology, National Agency for Research and Innovation; Molecular Biology Laboratory, Faculty Medicine and Health Sciences, Warmadewa University | Hidayat Trimarsanto, Lydia V. Panggalo, Iskandar Adnan, Sukma Oktavianthi, Lidwina Priliani, Edison Johar, Frilasita A Yudhaputri, Muhammad Rezki Rasyak, Willy Agustine, Sri Masyeni, Erly Sintya, Safarina G Malik, Khin Saw |
| EPI_ISL_2101153 | Molecular Biology Laboratory, Faculty Medicine and Health Sciences, Warmadewa University | Eijkman Institute for Molecular Biology, National Agency for Research and Innovation; Molecular Biology Laboratory, Faculty Medicine and Health Sciences, Warmadewa University | Frilasita A Yudhaputri, Muhammad Rezki Rasyak, Willy Agustine, Hidayat Trimarsanto, Lydia V. Panggalo, Iskandar Adnan, Sukma Oktavianthi, Lidwina Priliani, Edison Johar, Sri Masyeni, Erly Sintya, Safarina G Malik, Khin Saw |
| EPI_ISL_2101154 | Molecular Biology Laboratory, Faculty Medicine and Health Sciences, Warmadewa University | Eijkman Institute for Molecular Biology, National Agency for Research and Innovation; Molecular Biology Laboratory, Faculty Medicine and Health Sciences, Warmadewa University | Sukma Oktavianthi, Lidwina Priliani, Edison Johar, Frilasita A Yudhaputri, Muhammad Rezki Rasyak, Willy Agustine, Hidayat Trimarsanto, Lydia V. Panggalo, Iskandar Adnan, Sri Masyeni, Erly Sintya, Safarina G Malik, Khin Saw |
| EPI_ISL_2101155 | Molecular Biology Laboratory, Faculty Medicine and Health Sciences, Warmadewa University | Eijkman Institute for Molecular Biology, National Agency for Research and Innovation; Molecular Biology Laboratory, Faculty Medicine and Health Sciences, Warmadewa University | Lidwina Priliani, Edison Johar, Frilasita A Yudhaputri, Muhammad Rezki Rasyak, Willy Agustine, Hidayat Trimarsanto, Lydia V. Panggalo, Iskandar Adnan, Sukma Oktavianthi, Sri Masyeni, Erly Sintya, Safarina G Malik, Khin Saw |
| EPI_ISL_2101156 | Molecular Biology Laboratory, Faculty Medicine and Health Sciences, Warmadewa University | Eijkman Institute for Molecular Biology, National Agency for Research and Innovation; Molecular Biology Laboratory, Faculty | Edison Johar, Frilasita A Yudhaputri, Muhammad Rezki Rasyak, Willy Agustine, Hidayat Trimarsanto, Lydia V. Panggalo, Iskandar Adnan, Sukma Oktavianthi, Lidwina Priliani, Sri Masyeni, Erly Sintya, Safarina G Malik, Khin Saw |

| | | | |
|---|---|---|---|
| | | Medicine and Health Sciences, Warmadewa University | |
| EPI_ISL_2101157 | Molecular Biology Laboratory, Faculty Medicine and Health Sciences, Warmadewa University | Eijkman Institute for Molecular Biology, National Agency for Research and Innovation; Molecular Biology Laboratory, Faculty Medicine and Health Sciences, Warmadewa University | Frilasita A Yudhaputri, Muhammad Rezki Rasyak, Willy Agustine, Hidayat Trimarsanto, Lydia V. Panggalo, Iskandar Adnan, Sukma Oktavianthi, Lidwina Priliani, Edison Johar, Sri Masyeni, Erly Sintya, Safarina G Malik, Khin Saw |
| EPI_ISL_2101158 | Molecular Biology Laboratory, Faculty Medicine and Health Sciences, Warmadewa University | Eijkman Institute for Molecular Biology, National Agency for Research and Innovation; Molecular Biology Laboratory, Faculty Medicine and Health Sciences, Warmadewa University | Willy Agustine, Hidayat Trimarsanto, Lydia V. Panggalo, Iskandar Adnan, Sukma Oktavianthi, Lidwina Priliani, Edison Johar, Frilasita A Yudhaputri, Muhammad Rezki Rasyak, Sri Masyeni, Erly Sintya, Safarina G Malik, Khin Saw |
| EPI_ISL_2101159 | Molecular Biology Laboratory, Faculty Medicine and Health Sciences, Warmadewa University | Eijkman Institute for Molecular Biology, National Agency for Research and Innovation; Molecular Biology Laboratory, Faculty Medicine and Health Sciences, Warmadewa University | Hidayat Trimarsanto, Lydia V. Panggalo, Iskandar Adnan, Sukma Oktavianthi, Lidwina Priliani, Edison Johar, Frilasita A Yudhaputri, Muhammad Rezki Rasyak, Willy Agustine, Sri Masyeni, Erly Sintya, Safarina G Malik, Khin Saw |
| EPI_ISL_2101160 | Molecular Biology Laboratory, Faculty Medicine and Health Sciences, Warmadewa University | Eijkman Institute for Molecular Biology, National Agency for Research and Innovation; Molecular Biology Laboratory, Faculty Medicine and Health Sciences, Warmadewa University | Lydia V. Panggalo, Iskandar Adnan, Sukma Oktavianthi, Lidwina Priliani, Edison Johar, Frilasita A Yudhaputri, Muhammad Rezki Rasyak, Willy Agustine, Hidayat Trimarsanto, Sri Masyeni, Erly Sintya, Safarina G Malik, Khin Saw |
| EPI_ISL_2101161 | Molecular Biology Laboratory, Faculty Medicine and Health Sciences, Warmadewa University | Eijkman Institute for Molecular Biology, National Agency for Research and Innovation; Molecular Biology Laboratory, Faculty Medicine and Health Sciences, Warmadewa University | Iskandar Adnan, Sukma Oktavianthi, Lidwina Priliani, Edison Johar, Frilasita A Yudhaputri, Muhammad Rezki Rasyak, Willy Agustine, Hidayat Trimarsanto, Lydia V. Panggalo, Sri Masyeni, Erly Sintya, Safarina G Malik, Khin Saw |
| EPI_ISL_2101162 | Molecular Biology Laboratory, Faculty Medicine and Health Sciences, Warmadewa University | Eijkman Institute for Molecular Biology, National Agency for Research and Innovation; Molecular Biology Laboratory, Faculty Medicine and Health Sciences, Warmadewa University | Sukma Oktavianthi, Lidwina Priliani, Edison Johar, Frilasita A Yudhaputri, Muhammad Rezki Rasyak, Willy Agustine, Hidayat Trimarsanto, Lydia V. Panggalo, Iskandar Adnan, Sri Masyeni, Erly Sintya, Safarina G Malik, Khin Saw |
| EPI_ISL_2101163 | Molecular Biology Laboratory, Faculty Medicine and Health Sciences, Warmadewa University | Eijkman Institute for Molecular Biology, National Agency for Research and Innovation; Molecular Biology Laboratory, Faculty Medicine and Health Sciences, Warmadewa University | Iskandar Adnan, Sukma Oktavianthi, Lidwina Priliani, Edison Johar, Frilasita A Yudhaputri, Muhammad Rezki Rasyak, Willy Agustine, Hidayat Trimarsanto, Lydia V. Panggalo, Sri Masyeni, Erly Sintya, Safarina G Malik, Khin Saw |
| EPI_ISL_2101164 | Molecular Biology Laboratory, Faculty Medicine and Health Sciences, Warmadewa University | Eijkman Institute for Molecular Biology, National Agency for Research and Innovation; Molecular Biology Laboratory, Faculty Medicine and Health Sciences, Warmadewa University | Sukma Oktavianthi, Lidwina Priliani, Edison Johar, Frilasita A Yudhaputri, Muhammad Rezki Rasyak, Willy Agustine, Hidayat Trimarsanto, Lydia V. Panggalo, Iskandar Adnan, Sri Masyeni, Erly Sintya, Safarina G Malik, Khin Saw |
| EPI_ISL_2101165 | Molecular Biology Laboratory, Faculty Medicine and Health Sciences, Warmadewa University | Eijkman Institute for Molecular Biology, National Agency for Research and Innovation; Molecular Biology Laboratory, Faculty Medicine and Health Sciences, Warmadewa University | Lidwina Priliani, Edison Johar, Frilasita A Yudhaputri, Muhammad Rezki Rasyak, Willy Agustine, Hidayat Trimarsanto, Lydia V. Panggalo, Iskandar Adnan, Sukma Oktavianthi, Sri Masyeni, Erly Sintya, Safarina G Malik, Khin Saw |
| EPI_ISL_2101166 | Molecular Biology Laboratory, Faculty Medicine and Health Sciences, Warmadewa University | Eijkman Institute for Molecular Biology, National Agency for Research and Innovation; Molecular Biology Laboratory, Faculty | Edison Johar, Frilasita A Yudhaputri, Muhammad Rezki Rasyak, Willy Agustine, Hidayat Trimarsanto, Lydia V. Panggalo, Iskandar Adnan, Sukma Oktavianthi, Lidwina Priliani, Sri Masyeni, Erly Sintya, Safarina G Malik, Khin Saw |

| | | | |
|---|---|---|---|
| | | Medicine and Health Sciences, Warmadewa University | |
| EPI_ISL_2101167 | Molecular Biology Laboratory, Faculty Medicine and Health Sciences, Warmadewa University | Eijkman Institute for Molecular Biology, National Agency for Research and Innovation; Molecular Biology Laboratory, Faculty Medicine and Health Sciences, Warmadewa University | Frilasita A Yudhaputri, Muhammad Rezki Rasyak, Willy Agustine, Hidayat Trimarsanto, Lydia V. Panggalo, Iskandar Adnan, Sukma Oktavianthi, Lidwina Priliani, Edison Johar, Sri Masyeni, Erly Sintya, Safarina G Malik, Khin Saw |
| EPI_ISL_2101168 | Molecular Biology Laboratory, Faculty Medicine and Health Sciences, Warmadewa University | Eijkman Institute for Molecular Biology, National Agency for Research and Innovation; Molecular Biology Laboratory, Faculty Medicine and Health Sciences, Warmadewa University | Willy Agustine, Hidayat Trimarsanto, Lydia V. Panggalo, Iskandar Adnan, Sukma Oktavianthi, Lidwina Priliani, Edison Johar, Frilasita A Yudhaputri, Muhammad Rezki Rasyak, Sri Masyeni, Erly Sintya, Safarina G Malik, Khin Saw |
| EPI_ISL_2101169 | Molecular Biology Laboratory, Faculty Medicine and Health Sciences, Warmadewa University | Eijkman Institute for Molecular Biology, National Agency for Research and Innovation; Molecular Biology Laboratory, Faculty Medicine and Health Sciences, Warmadewa University | Hidayat Trimarsanto, Lydia V. Panggalo, Iskandar Adnan, Sukma Oktavianthi, Lidwina Priliani, Edison Johar, Frilasita A Yudhaputri, Muhammad Rezki Rasyak, Willy Agustine, Sri Masyeni, Erly Sintya, Safarina G Malik, Khin Saw |
| EPI_ISL_2101170 | Molecular Biology Laboratory, Faculty Medicine and Health Sciences, Warmadewa University | Eijkman Institute for Molecular Biology, National Agency for Research and Innovation; Molecular Biology Laboratory, Faculty Medicine and Health Sciences, Warmadewa University | Lidwina Priliani, Edison Johar, Frilasita A Yudhaputri, Muhammad Rezki Rasyak, Willy Agustine, Hidayat Trimarsanto, Lydia V. Panggalo, Iskandar Adnan, Sukma Oktavianthi, Sri Masyeni, Erly Sintya, Safarina G Malik, Khin Saw |
| EPI_ISL_2101171 | Molecular Biology Laboratory, Faculty Medicine and Health Sciences, Warmadewa University | Eijkman Institute for Molecular Biology, National Agency for Research and Innovation; Molecular Biology Laboratory, Faculty Medicine and Health Sciences, Warmadewa University | Edison Johar, Frilasita A Yudhaputri, Muhammad Rezki Rasyak, Willy Agustine, Hidayat Trimarsanto, Lydia V. Panggalo, Iskandar Adnan, Sukma Oktavianthi, Lidwina Priliani, Sri Masyeni, Erly Sintya, Safarina G Malik, Khin Saw |
| EPI_ISL_2101172 | Molecular Biology Laboratory, Faculty Medicine and Health Sciences, Warmadewa University | Eijkman Institute for Molecular Biology, National Agency for Research and Innovation; Molecular Biology Laboratory, Faculty Medicine and Health Sciences, Warmadewa University | Iskandar Adnan, Sukma Oktavianthi, Lidwina Priliani, Edison Johar, Frilasita A Yudhaputri, Muhammad Rezki Rasyak, Willy Agustine, Hidayat Trimarsanto, Lydia V. Panggalo, Sri Masyeni, Erly Sintya, Safarina G Malik, Khin Saw |
| EPI_ISL_2101173 | Molecular Biology Laboratory, Faculty Medicine and Health Sciences, Warmadewa University | Eijkman Institute for Molecular Biology, National Agency for Research and Innovation; Molecular Biology Laboratory, Faculty Medicine and Health Sciences, Warmadewa University | Sukma Oktavianthi, Lidwina Priliani, Edison Johar, Frilasita A Yudhaputri, Muhammad Rezki Rasyak, Willy Agustine, Hidayat Trimarsanto, Lydia V. Panggalo, Iskandar Adnan, Sri Masyeni, Erly Sintya, Safarina G Malik, Khin Saw |
| EPI_ISL_2101174 | Molecular Biology Laboratory, Faculty Medicine and Health Sciences, Warmadewa University | Eijkman Institute for Molecular Biology, National Agency for Research and Innovation; Molecular Biology Laboratory, Faculty Medicine and Health Sciences, Warmadewa University | Lidwina Priliani, Edison Johar, Frilasita A Yudhaputri, Muhammad Rezki Rasyak, Willy Agustine, Hidayat Trimarsanto, Lydia V. Panggalo, Iskandar Adnan, Sukma Oktavianthi, Sri Masyeni, Erly Sintya, Safarina G Malik, Khin Saw |
| EPI_ISL_2101175 | Molecular Biology Laboratory, Faculty Medicine and Health Sciences, Warmadewa University | Eijkman Institute for Molecular Biology, National Agency for Research and Innovation; Molecular Biology Laboratory, Faculty Medicine and Health Sciences, Warmadewa University | Edison Johar, Frilasita A Yudhaputri, Muhammad Rezki Rasyak, Willy Agustine, Hidayat Trimarsanto, Lydia V. Panggalo, Iskandar Adnan, Sukma Oktavianthi, Lidwina Priliani, Sri Masyeni, Erly Sintya, Safarina G Malik, Khin Saw |
| EPI_ISL_2101176 | Molecular Biology Laboratory, Faculty Medicine and Health Sciences, Warmadewa University | Eijkman Institute for Molecular Biology, National Agency for Research and Innovation; Molecular Biology Laboratory, Faculty | Frilasita A Yudhaputri, Muhammad Rezki Rasyak, Willy Agustine, Hidayat Trimarsanto, Lydia V. Panggalo, Iskandar Adnan, Sukma Oktavianthi, Lidwina Priliani, Edison Johar, Sri Masyeni, Erly Sintya, Safarina G Malik, Khin Saw |

| | | | |
|---|---|---|---|
| | | Medicine and Health Sciences, Warmadewa University | |
| EPI_ISL_2101177 | Molecular Biology Laboratory, Faculty Medicine and Health Sciences, Warmadewa University | Eijkman Institute for Molecular Biology, National Agency for Research and Innovation; Molecular Biology Laboratory, Faculty Medicine and Health Sciences, Warmadewa University | Willy Agustine, Hidayat Trimarsanto, Lydia V. Panggalo, Iskandar Adnan, Sukma Oktavianthi, Lidwina Priliani, Edison Johar, Frilasita A Yudhaputri, Muhammad Rezki Rasyak, Sri Masyeni, Erly Sintya, Safarina G Malik, Khin Saw |
| EPI_ISL_2101178 | Molecular Biology Laboratory, Faculty Medicine and Health Sciences, Warmadewa University | Eijkman Institute for Molecular Biology, National Agency for Research and Innovation; Molecular Biology Laboratory, Faculty Medicine and Health Sciences, Warmadewa University | Hidayat Trimarsanto, Lydia V. Panggalo, Iskandar Adnan, Sukma Oktavianthi, Lidwina Priliani, Edison Johar, Frilasita A Yudhaputri, Muhammad Rezki Rasyak, Willy Agustine, Sri Masyeni, Erly Sintya, Safarina G Malik, Khin Saw |
| EPI_ISL_2101179 | Molecular Biology Laboratory, Faculty Medicine and Health Sciences, Warmadewa University | Eijkman Institute for Molecular Biology, National Agency for Research and Innovation; Molecular Biology Laboratory, Faculty Medicine and Health Sciences, Warmadewa University | Lydia V. Panggalo, Iskandar Adnan, Sukma Oktavianthi, Lidwina Priliani, Edison Johar, Frilasita A Yudhaputri, Muhammad Rezki Rasyak, Willy Agustine, Hidayat Trimarsanto, Sri Masyeni, Erly Sintya, Safarina G Malik, Khin Saw |
| EPI_ISL_2101180 | Molecular Biology Laboratory, Faculty Medicine and Health Sciences, Warmadewa University | Eijkman Institute for Molecular Biology, National Agency for Research and Innovation; Molecular Biology Laboratory, Faculty Medicine and Health Sciences, Warmadewa University | Sukma Oktavianthi, Lidwina Priliani, Edison Johar, Frilasita A Yudhaputri, Muhammad Rezki Rasyak, Willy Agustine, Hidayat Trimarsanto, Lydia V. Panggalo, Iskandar Adnan, Sri Masyeni, Erly Sintya, Safarina G Malik, Khin Saw |
| EPI_ISL_2101181 | Molecular Biology Laboratory, Faculty Medicine and Health Sciences, Warmadewa University | Eijkman Institute for Molecular Biology, National Agency for Research and Innovation; Molecular Biology Laboratory, Faculty Medicine and Health Sciences, Warmadewa University | Lidwina Priliani, Edison Johar, Frilasita A Yudhaputri, Muhammad Rezki Rasyak, Willy Agustine, Hidayat Trimarsanto, Lydia V. Panggalo, Iskandar Adnan, Sukma Oktavianthi, Sri Masyeni, Erly Sintya, Safarina G Malik, Khin Saw |
| EPI_ISL_2101182 | Molecular Biology Laboratory, Faculty Medicine and Health Sciences, Warmadewa University | Eijkman Institute for Molecular Biology, National Agency for Research and Innovation; Molecular Biology Laboratory, Faculty Medicine and Health Sciences, Warmadewa University | Edison Johar, Frilasita A Yudhaputri, Muhammad Rezki Rasyak, Willy Agustine, Hidayat Trimarsanto, Lydia V. Panggalo, Iskandar Adnan, Sukma Oktavianthi, Lidwina Priliani, Sri Masyeni, Erly Sintya, Safarina G Malik, Khin Saw |
| EPI_ISL_2101183 | Molecular Biology Laboratory, Faculty Medicine and Health Sciences, Warmadewa University | Eijkman Institute for Molecular Biology, National Agency for Research and Innovation; Molecular Biology Laboratory, Faculty Medicine and Health Sciences, Warmadewa University | Frilasita A Yudhaputri, Muhammad Rezki Rasyak, Willy Agustine, Hidayat Trimarsanto, Lydia V. Panggalo, Iskandar Adnan, Sukma Oktavianthi, Lidwina Priliani, Edison Johar, Sri Masyeni, Erly Sintya, Safarina G Malik, Khin Saw |
| EPI_ISL_2101184 | Molecular Biology Laboratory, Faculty Medicine and Health Sciences, Warmadewa University | Eijkman Institute for Molecular Biology, National Agency for Research and Innovation; Molecular Biology Laboratory, Faculty Medicine and Health Sciences, Warmadewa University | Willy Agustine, Hidayat Trimarsanto, Lydia V. Panggalo, Iskandar Adnan, Sukma Oktavianthi, Lidwina Priliani, Edison Johar, Frilasita A Yudhaputri, Muhammad Rezki Rasyak, Sri Masyeni, Erly Sintya, Safarina G Malik, Khin Saw |
| EPI_ISL_2101185 | Molecular Biology Laboratory, Faculty Medicine and Health Sciences, Warmadewa University | Eijkman Institute for Molecular Biology, National Agency for Research and Innovation; Molecular Biology Laboratory, Faculty Medicine and Health Sciences, Warmadewa University | Hidayat Trimarsanto, Lydia V. Panggalo, Iskandar Adnan, Sukma Oktavianthi, Lidwina Priliani, Edison Johar, Frilasita A Yudhaputri, Muhammad Rezki Rasyak, Willy Agustine, Sri Masyeni, Erly Sintya, Safarina G Malik, Khin Saw |
| EPI_ISL_2101186 | Molecular Biology Laboratory, Faculty Medicine and Health Sciences, Warmadewa University | Eijkman Institute for Molecular Biology, National Agency for Research and Innovation; Molecular Biology Laboratory, Faculty | Lydia V. Panggalo, Iskandar Adnan, Sukma Oktavianthi, Lidwina Priliani, Edison Johar, Frilasita A Yudhaputri, Muhammad Rezki Rasyak, Willy Agustine, Hidayat Trimarsanto, Sri Masyeni, Erly Sintya, Safarina G Malik, Khin Saw |

| | | | |
|---|---|---|---|
| | | Medicine and Health Sciences, Warmadewa University | |
| EPI_ISL_2101187 | Molecular Biology Laboratory, Faculty Medicine and Health Sciences, Warmadewa University | Eijkman Institute for Molecular Biology, National Agency for Research and Innovation; Molecular Biology Laboratory, Faculty Medicine and Health Sciences, Warmadewa University | Iskandar Adnan, Sukma Oktavianthi, Lidwina Priliani, Edison Johar, Frilasita A Yudhaputri, Muhammad Rezki Rasyak, Willy Agustine, Hidayat Trimarsanto, Lydia V. Panggalo, Sri Masyeni, Erly Sintya, Safarina G Malik, Khin Saw |
| EPI_ISL_2101188 | Molecular Biology Laboratory, Faculty Medicine and Health Sciences, Warmadewa University | Eijkman Institute for Molecular Biology, National Agency for Research and Innovation; Molecular Biology Laboratory, Faculty Medicine and Health Sciences, Warmadewa University | Sukma Oktavianthi, Lidwina Priliani, Edison Johar, Frilasita A Yudhaputri, Muhammad Rezki Rasyak, Willy Agustine, Hidayat Trimarsanto, Lydia V. Panggalo, Iskandar Adnan, Sri Masyeni, Erly Sintya, Safarina G Malik, Khin Saw |
| EPI_ISL_2101189 | Molecular Biology Laboratory, Faculty Medicine and Health Sciences, Warmadewa University | Eijkman Institute for Molecular Biology, National Agency for Research and Innovation; Molecular Biology Laboratory, Faculty Medicine and Health Sciences, Warmadewa University | Lidwina Priliani, Edison Johar, Frilasita A Yudhaputri, Muhammad Rezki Rasyak, Willy Agustine, Hidayat Trimarsanto, Lydia V. Panggalo, Iskandar Adnan, Sukma Oktavianthi, Sri Masyeni, Erly Sintya, Safarina G Malik, Khin Saw |
| EPI_ISL_2101190 | Molecular Biology Laboratory, Faculty Medicine and Health Sciences, Warmadewa University | Eijkman Institute for Molecular Biology, National Agency for Research and Innovation; Molecular Biology Laboratory, Faculty Medicine and Health Sciences, Warmadewa University | Frilasita A Yudhaputri, Muhammad Rezki Rasyak, Willy Agustine, Hidayat Trimarsanto, Lydia V. Panggalo, Iskandar Adnan, Sukma Oktavianthi, Lidwina Priliani, Edison Johar, Sri Masyeni, Erly Sintya, Safarina G Malik, Khin Saw |
| EPI_ISL_2101191 | Molecular Biology Laboratory, Faculty Medicine and Health Sciences, Warmadewa University | Eijkman Institute for Molecular Biology, National Agency for Research and Innovation; Molecular Biology Laboratory, Faculty Medicine and Health Sciences, Warmadewa University | Willy Agustine, Hidayat Trimarsanto, Lydia V. Panggalo, Iskandar Adnan, Sukma Oktavianthi, Lidwina Priliani, Edison Johar, Frilasita A Yudhaputri, Muhammad Rezki Rasyak, Sri Masyeni, Erly Sintya, Safarina G Malik, Khin Saw |
| EPI_ISL_2101192 | Molecular Biology Laboratory, Faculty Medicine and Health Sciences, Warmadewa University | Eijkman Institute for Molecular Biology, National Agency for Research and Innovation; Molecular Biology Laboratory, Faculty Medicine and Health Sciences, Warmadewa University | Hidayat Trimarsanto, Lydia V. Panggalo, Iskandar Adnan, Sukma Oktavianthi, Lidwina Priliani, Edison Johar, Frilasita A Yudhaputri, Muhammad Rezki Rasyak, Willy Agustine, Sri Masyeni, Erly Sintya, Safarina G Malik, Khin Saw |
| EPI_ISL_2101193 | Molecular Biology Laboratory, Faculty Medicine and Health Sciences, Warmadewa University | Eijkman Institute for Molecular Biology, National Agency for Research and Innovation; Molecular Biology Laboratory, Faculty Medicine and Health Sciences, Warmadewa University | Lydia V. Panggalo, Iskandar Adnan, Sukma Oktavianthi, Lidwina Priliani, Edison Johar, Frilasita A Yudhaputri, Muhammad Rezki Rasyak, Willy Agustine, Hidayat Trimarsanto, Sri Masyeni, Erly Sintya, Safarina G Malik, Khin Saw |
| EPI_ISL_2101194 | Molecular Biology Laboratory, Faculty Medicine and Health Sciences, Warmadewa University | Eijkman Institute for Molecular Biology, National Agency for Research and Innovation; Molecular Biology Laboratory, Faculty Medicine and Health Sciences, Warmadewa University | Iskandar Adnan, Sukma Oktavianthi, Lidwina Priliani, Edison Johar, Frilasita A Yudhaputri, Muhammad Rezki Rasyak, Willy Agustine, Hidayat Trimarsanto, Lydia V. Panggalo, Sri Masyeni, Erly Sintya, Safarina G Malik, Khin Saw |
| EPI_ISL_2101195 | Molecular Biology Laboratory, Faculty Medicine and Health Sciences, Warmadewa University | Eijkman Institute for Molecular Biology, National Agency for Research and Innovation; Molecular Biology Laboratory, Faculty Medicine and Health Sciences, Warmadewa University | Sukma Oktavianthi, Lidwina Priliani, Edison Johar, Frilasita A Yudhaputri, Muhammad Rezki Rasyak, Willy Agustine, Hidayat Trimarsanto, Lydia V. Panggalo, Iskandar Adnan, Sri Masyeni, Erly Sintya, Safarina G Malik, Khin Saw |
| EPI_ISL_2101196 | Molecular Biology Laboratory, Faculty Medicine and Health Sciences, Warmadewa University | Eijkman Institute for Molecular Biology, National Agency for Research and Innovation; Molecular Biology Laboratory, Faculty | Lidwina Priliani, Edison Johar, Frilasita A Yudhaputri, Muhammad Rezki Rasyak, Willy Agustine, Hidayat Trimarsanto, Lydia V. Panggalo, Iskandar Adnan, Sukma Oktavianthi, Sri Masyeni, Erly Sintya, Safarina G Malik, Khin Saw |

| | | | |
|---|---|---|---|
| | | Medicine and Health Sciences, Warmadewa University | |
| EPI_ISL_2101197 | Molecular Biology Laboratory, Faculty Medicine and Health Sciences, Warmadewa University | Eijkman Institute for Molecular Biology, National Agency for Research and Innovation; Molecular Biology Laboratory, Faculty Medicine and Health Sciences, Warmadewa University | Edison Johar, Frilasita A Yudhaputri, Muhammad Rezki Rasyak, Willy Agustine, Hidayat Trimarsanto, Lydia V. Panggalo, Iskandar Adnan, Sukma Oktavianthi, Lidwina Priliani, Sri Masyeni, Erly Sintya, Safarina G Malik, Khin Saw |
| EPI_ISL_2101198 | Molecular Biology Laboratory, Faculty Medicine and Health Sciences, Warmadewa University | Eijkman Institute for Molecular Biology, National Agency for Research and Innovation; Molecular Biology Laboratory, Faculty Medicine and Health Sciences, Warmadewa University | Frilasita A Yudhaputri, Muhammad Rezki Rasyak, Willy Agustine, Hidayat Trimarsanto, Lydia V. Panggalo, Iskandar Adnan, Sukma Oktavianthi, Lidwina Priliani, Edison Johar, Sri Masyeni, Erly Sintya, Safarina G Malik, Khin Saw |
| EPI_ISL_2101199 | Molecular Biology Laboratory, Faculty Medicine and Health Sciences, Warmadewa University | Eijkman Institute for Molecular Biology, National Agency for Research and Innovation; Molecular Biology Laboratory, Faculty Medicine and Health Sciences, Warmadewa University | Willy Agustine, Hidayat Trimarsanto, Lydia V. Panggalo, Iskandar Adnan, Sukma Oktavianthi, Lidwina Priliani, Edison Johar, Frilasita A Yudhaputri, Muhammad Rezki Rasyak, Sri Masyeni, Erly Sintya, Safarina G Malik, Khin Saw |
| EPI_ISL_2101200 | Molecular Biology Laboratory, Faculty Medicine and Health Sciences, Warmadewa University | Eijkman Institute for Molecular Biology, National Agency for Research and Innovation; Molecular Biology Laboratory, Faculty Medicine and Health Sciences, Warmadewa University | Lydia V. Panggalo, Iskandar Adnan, Sukma Oktavianthi, Lidwina Priliani, Edison Johar, Frilasita A Yudhaputri, Muhammad Rezki Rasyak, Willy Agustine, Hidayat Trimarsanto, Sri Masyeni, Erly Sintya, Safarina G Malik, Khin Saw |
| EPI_ISL_2101202 | Molecular Biology Laboratory, Faculty Medicine and Health Sciences, Warmadewa University | Eijkman Institute for Molecular Biology, National Agency for Research and Innovation; Molecular Biology Laboratory, Faculty Medicine and Health Sciences, Warmadewa University | Iskandar Adnan, Sukma Oktavianthi, Lidwina Priliani, Edison Johar, Frilasita A Yudhaputri, Muhammad Rezki Rasyak, Willy Agustine, Hidayat Trimarsanto, Lydia V. Panggalo, Sri Masyeni, Erly Sintya, Safarina G Malik, Khin Saw |
| EPI_ISL_2101203 | Molecular Biology Laboratory, Faculty Medicine and Health Sciences, Warmadewa University | Eijkman Institute for Molecular Biology, National Agency for Research and Innovation; Molecular Biology Laboratory, Faculty Medicine and Health Sciences, Warmadewa University | Sukma Oktavianthi, Lidwina Priliani, Edison Johar, Frilasita A Yudhaputri, Muhammad Rezki Rasyak, Willy Agustine, Hidayat Trimarsanto, Lydia V. Panggalo, Iskandar Adnan, Sri Masyeni, Erly Sintya, Safarina G Malik, Khin Saw |
| EPI_ISL_2101204 | Molecular Biology Laboratory, Faculty Medicine and Health Sciences, Warmadewa University | Eijkman Institute for Molecular Biology, National Agency for Research and Innovation; Molecular Biology Laboratory, Faculty Medicine and Health Sciences, Warmadewa University | Lidwina Priliani, Edison Johar, Frilasita A Yudhaputri, Muhammad Rezki Rasyak, Willy Agustine, Hidayat Trimarsanto, Lydia V. Panggalo, Iskandar Adnan, Sukma Oktavianthi, Sri Masyeni, Erly Sintya, Safarina G Malik, Khin Saw |
| EPI_ISL_2101205 | Molecular Biology Laboratory, Faculty Medicine and Health Sciences, Warmadewa University | Eijkman Institute for Molecular Biology, National Agency for Research and Innovation; Molecular Biology Laboratory, Faculty Medicine and Health Sciences, Warmadewa University | Edison Johar, Frilasita A Yudhaputri, Muhammad Rezki Rasyak, Willy Agustine, Hidayat Trimarsanto, Lydia V. Panggalo, Iskandar Adnan, Sukma Oktavianthi, Lidwina Priliani, Sri Masyeni, Erly Sintya, Safarina G Malik, Khin Saw |
| EPI_ISL_2101206 | Molecular Biology Laboratory, Faculty Medicine and Health Sciences, Warmadewa University | Eijkman Institute for Molecular Biology, National Agency for Research and Innovation; Molecular Biology Laboratory, Faculty Medicine and Health Sciences, Warmadewa University | Frilasita A Yudhaputri, Muhammad Rezki Rasyak, Willy Agustine, Hidayat Trimarsanto, Lydia V. Panggalo, Iskandar Adnan, Sukma Oktavianthi, Lidwina Priliani, Edison Johar, Sri Masyeni, Erly Sintya, Safarina G Malik, Khin Saw |
| EPI_ISL_2101207 | Molecular Biology Laboratory, Faculty Medicine and Health Sciences, Warmadewa University | Eijkman Institute for Molecular Biology, National Agency for Research and Innovation; Molecular Biology Laboratory, Faculty | Willy Agustine, Hidayat Trimarsanto, Lydia V. Panggalo, Iskandar Adnan, Sukma Oktavianthi, Lidwina Priliani, Edison Johar, Frilasita A Yudhaputri, Muhammad Rezki Rasyak, Sri Masyeni, Erly Sintya, Safarina G Malik, Khin Saw |

| | | | |
|---|---|---|---|
| | | Medicine and Health Sciences, Warmadewa University | |
| EPI_ISL_2101208 | Molecular Biology Laboratory, Faculty Medicine and Health Sciences, Warmadewa University | Eijkman Institute for Molecular Biology, National Agency for Research and Innovation; Molecular Biology Laboratory, Faculty Medicine and Health Sciences, Warmadewa University | Hidayat Trimarsanto, Lydia V. Panggalo, Iskandar Adnan, Sukma Oktavianthi, Lidwina Priliani, Edison Johar, Frilasita A Yudhaputri, Muhammad Rezki Rasyak, Willy Agustine, Sri Masyeni, Erly Sintya, Safarina G Malik, Khin Saw |
| EPI_ISL_2101209 | Molecular Biology Laboratory, Faculty Medicine and Health Sciences, Warmadewa University | Eijkman Institute for Molecular Biology, National Agency for Research and Innovation; Molecular Biology Laboratory, Faculty Medicine and Health Sciences, Warmadewa University | Lydia V. Panggalo, Iskandar Adnan, Sukma Oktavianthi, Lidwina Priliani, Edison Johar, Frilasita A Yudhaputri, Muhammad Rezki Rasyak, Willy Agustine, Hidayat Trimarsanto, Sri Masyeni, Erly Sintya, Safarina G Malik, Khin Saw |
| EPI_ISL_2101210 | Molecular Biology Laboratory, Faculty Medicine and Health Sciences, Warmadewa University | Eijkman Institute for Molecular Biology, National Agency for Research and Innovation; Molecular Biology Laboratory, Faculty Medicine and Health Sciences, Warmadewa University | Iskandar Adnan, Sukma Oktavianthi, Lidwina Priliani, Edison Johar, Frilasita A Yudhaputri, Muhammad Rezki Rasyak, Willy Agustine, Hidayat Trimarsanto, Lydia V. Panggalo, Sri Masyeni, Erly Sintya, Safarina G Malik, Khin Saw |
| EPI_ISL_2101211 | Molecular Biology Laboratory, Faculty Medicine and Health Sciences, Warmadewa University | Eijkman Institute for Molecular Biology, National Agency for Research and Innovation; Molecular Biology Laboratory, Faculty Medicine and Health Sciences, Warmadewa University | Lidwina Priliani, Edison Johar, Frilasita A Yudhaputri, Muhammad Rezki Rasyak, Willy Agustine, Hidayat Trimarsanto, Lydia V. Panggalo, Iskandar Adnan, Sukma Oktavianthi, Sri Masyeni, Erly Sintya, Safarina G Malik, Khin Saw |
| EPI_ISL_2101212 | Molecular Biology Laboratory, Faculty Medicine and Health Sciences, Warmadewa University | Eijkman Institute for Molecular Biology, National Agency for Research and Innovation; Molecular Biology Laboratory, Faculty Medicine and Health Sciences, Warmadewa University | Edison Johar, Frilasita A Yudhaputri, Muhammad Rezki Rasyak, Willy Agustine, Hidayat Trimarsanto, Lydia V. Panggalo, Iskandar Adnan, Sukma Oktavianthi, Lidwina Priliani, Sri Masyeni, Erly Sintya, Safarina G Malik, Khin Saw |
| EPI_ISL_2101213 | Molecular Biology Laboratory, Faculty Medicine and Health Sciences, Warmadewa University | Eijkman Institute for Molecular Biology, National Agency for Research and Innovation; Molecular Biology Laboratory, Faculty Medicine and Health Sciences, Warmadewa University | Frilasita A Yudhaputri, Muhammad Rezki Rasyak, Willy Agustine, Hidayat Trimarsanto, Lydia V. Panggalo, Iskandar Adnan, Sukma Oktavianthi, Lidwina Priliani, Edison Johar, Sri Masyeni, Erly Sintya, Safarina G Malik, Khin Saw |
| EPI_ISL_2101214 | Molecular Biology Laboratory, Faculty Medicine and Health Sciences, Warmadewa University | Eijkman Institute for Molecular Biology, National Agency for Research and Innovation; Molecular Biology Laboratory, Faculty Medicine and Health Sciences, Warmadewa University | Willy Agustine, Hidayat Trimarsanto, Lydia V. Panggalo, Iskandar Adnan, Sukma Oktavianthi, Lidwina Priliani, Edison Johar, Frilasita A Yudhaputri, Muhammad Rezki Rasyak, Sri Masyeni, Erly Sintya, Safarina G Malik, Khin Saw |
| EPI_ISL_2101215 | Molecular Biology Laboratory, Faculty Medicine and Health Sciences, Warmadewa University | Eijkman Institute for Molecular Biology, National Agency for Research and Innovation; Molecular Biology Laboratory, Faculty Medicine and Health Sciences, Warmadewa University | Hidayat Trimarsanto, Lydia V. Panggalo, Iskandar Adnan, Sukma Oktavianthi, Lidwina Priliani, Edison Johar, Frilasita A Yudhaputri, Muhammad Rezki Rasyak, Willy Agustine, Sri Masyeni, Erly Sintya, Safarina G Malik, Khin Saw |
| EPI_ISL_2101216 | BRSU Tabanan | Eijkman Institute for Molecular Biology, National Agency for Research and Innovation; Molecular Biology Laboratory, Faculty Medicine and Health Sciences, Warmadewa University | Frilasita A Yudhaputri, Muhammad Rezki Rasyak, Willy Agustine, Hidayat Trimarsanto, Lydia V. Panggalo, Iskandar Adnan, Sukma Oktavianthi, Lidwina Priliani, Edison Johar, Sri Masyeni, Erly Sintya, I Wayan Duta Krisna, Safarina G Malik, |
| EPI_ISL_2101217 | BRSU Tabanan | Eijkman Institute for Molecular Biology, National Agency for Research and Innovation; Molecular Biology Laboratory, Faculty | Muhammad Rezki Rasyak, Willy Agustine, Hidayat Trimarsanto, Lydia V. Panggalo, Iskandar Adnan, Sukma Oktavianthi, Lidwina Priliani, Edison Johar, Frilasita A Yudhaputri, Sri Masyeni, Erly Sintya, I Wayan Duta Krisna, Safarina G Malik, |

| | | | |
|---|---|---|---|
| | | Medicine and Health Sciences, Warmadewa University | |
| EPI_ISL_2101218 | BRSU Tabanan | Eijkman Institute for Molecular Biology, National Agency for Research and Innovation; Molecular Biology Laboratory, Faculty Medicine and Health Sciences, Warmadewa University | Willy Agustine, Hidayat Trimarsanto, Lydia V. Panggalo, Iskandar Adnan, Sukma Oktavianthi, Lidwina Priliani, Edison Johar, Frilasita A Yudhaputri, Muhammad Rezki Rasyak, Sri Masyeni, Erly Sintya, I Wayan Duta Krisna, Safarina G Malik, |
| EPI_ISL_2101219 | BRSU Tabanan | Eijkman Institute for Molecular Biology, National Agency for Research and Innovation; Molecular Biology Laboratory, Faculty Medicine and Health Sciences, Warmadewa University | Hidayat Trimarsanto, Lydia V. Panggalo, Iskandar Adnan, Sukma Oktavianthi, Lidwina Priliani, Edison Johar, Frilasita A Yudhaputri, Muhammad Rezki Rasyak, Willy Agustine, Sri Masyeni, Erly Sintya, I Wayan Duta Krisna, Safarina G Malik, |
| EPI_ISL_2101220 | BRSU Tabanan | Eijkman Institute for Molecular Biology, National Agency for Research and Innovation; Molecular Biology Laboratory, Faculty Medicine and Health Sciences, Warmadewa University | Lydia V. Panggalo, Iskandar Adnan, Sukma Oktavianthi, Lidwina Priliani, Edison Johar, Frilasita A Yudhaputri, Muhammad Rezki Rasyak, Willy Agustine, Hidayat Trimarsanto, Sri Masyeni, Erly Sintya, I Wayan Duta Krisna, Safarina G Malik, |
| EPI_ISL_2101221 | BRSU Tabanan | Eijkman Institute for Molecular Biology, National Agency for Research and Innovation; Molecular Biology Laboratory, Faculty Medicine and Health Sciences, Warmadewa University | Iskandar Adnan, Sukma Oktavianthi, Lidwina Priliani, Edison Johar, Frilasita A Yudhaputri, Muhammad Rezki Rasyak, Willy Agustine, Hidayat Trimarsanto, Lydia V. Panggalo, Sri Masyeni, Erly Sintya, I Wayan Duta Krisna, Safarina G Malik, |
| EPI_ISL_2101222 | BRSU Tabanan | Eijkman Institute for Molecular Biology, National Agency for Research and Innovation; Molecular Biology Laboratory, Faculty Medicine and Health Sciences, Warmadewa University | Sukma Oktavianthi, Lidwina Priliani, Edison Johar, Frilasita A Yudhaputri, Muhammad Rezki Rasyak, Willy Agustine, Hidayat Trimarsanto, Lydia V. Panggalo, Iskandar Adnan, Sri Masyeni, Erly Sintya, I Wayan Duta Krisna, Safarina G Malik, |
| EPI_ISL_2101223 | BRSU Tabanan | Eijkman Institute for Molecular Biology, National Agency for Research and Innovation; Molecular Biology Laboratory, Faculty Medicine and Health Sciences, Warmadewa University | Lidwina Priliani, Edison Johar, Frilasita A Yudhaputri, Muhammad Rezki Rasyak, Willy Agustine, Hidayat Trimarsanto, Lydia V. Panggalo, Iskandar Adnan, Sukma Oktavianthi, Sri Masyeni, Erly Sintya, I Wayan Duta Krisna, Safarina G Malik, |
| EPI_ISL_2101224 | BRSU Tabanan | Eijkman Institute for Molecular Biology, National Agency for Research and Innovation; Molecular Biology Laboratory, Faculty Medicine and Health Sciences, Warmadewa University | Frilasita A Yudhaputri, Muhammad Rezki Rasyak, Willy Agustine, Hidayat Trimarsanto, Lydia V. Panggalo, Iskandar Adnan, Sukma Oktavianthi, Lidwina Priliani, Edison Johar, Sri Masyeni, Erly Sintya, I Wayan Duta Krisna, Safarina G Malik, |
| EPI_ISL_2101225 | BRSU Tabanan | Eijkman Institute for Molecular Biology, National Agency for Research and Innovation; Molecular Biology Laboratory, Faculty Medicine and Health Sciences, Warmadewa University | Muhammad Rezki Rasyak, Willy Agustine, Hidayat Trimarsanto, Lydia V. Panggalo, Iskandar Adnan, Sukma Oktavianthi, Lidwina Priliani, Edison Johar, Frilasita A Yudhaputri, Sri Masyeni, Erly Sintya, I Wayan Duta Krisna, Safarina G Malik, |
| EPI_ISL_2101226 | BRSU Tabanan | Eijkman Institute for Molecular Biology, National Agency for Research and Innovation; Molecular Biology Laboratory, Faculty Medicine and Health Sciences, Warmadewa University | Willy Agustine, Hidayat Trimarsanto, Lydia V. Panggalo, Iskandar Adnan, Sukma Oktavianthi, Lidwina Priliani, Edison Johar, Frilasita A Yudhaputri, Muhammad Rezki Rasyak, Sri Masyeni, Erly Sintya, I Wayan Duta Krisna, Safarina G Malik, |
| EPI_ISL_2101227 | BRSU Tabanan | Eijkman Institute for Molecular Biology, National Agency for Research and Innovation; Molecular Biology Laboratory, Faculty | Hidayat Trimarsanto, Lydia V. Panggalo, Iskandar Adnan, Sukma Oktavianthi, Lidwina Priliani, Edison Johar, Frilasita A Yudhaputri, Muhammad Rezki Rasyak, Willy Agustine, Sri Masyeni, Erly Sintya, I Wayan Duta Krisna, Safarina G Malik, |

| | | | |
|---|---|---|---|
| | | Medicine and Health Sciences, Warmadewa University | |
| EPI_ISL_2101228 | BRSU Tabanan | Eijkman Institute for Molecular Biology, National Agency for Research and Innovation; Molecular Biology Laboratory, Faculty Medicine and Health Sciences, Warmadewa University | Lydia V. Panggalo, Iskandar Adnan, Sukma Oktavianthi, Lidwina Priliani, Edison Johar, Frilasita A Yudhaputri, Muhammad Rezki Rasyak, Willy Agustine, Hidayat Trimarsanto, Sri Masyeni, Erly Sintya, I Wayan Duta Krisna, Safarina G Malik, |
| EPI_ISL_2101229 | RSUD Klungkung | Eijkman Institute for Molecular Biology, National Agency for Research and Innovation; Molecular Biology Laboratory, Faculty Medicine and Health Sciences, Warmadewa University | Iskandar Adnan, Sukma Oktavianthi, Lidwina Priliani, Edison Johar, Frilasita A Yudhaputri, Muhammad Rezki Rasyak, Willy Agustine, Hidayat Trimarsanto, Lydia V. Panggalo, Sri Masyeni, Erly Sintya, I Komang Parwata, Safarina G Malik, |
| EPI_ISL_2101230 | RSUD Klungkung | Eijkman Institute for Molecular Biology, National Agency for Research and Innovation; Molecular Biology Laboratory, Faculty Medicine and Health Sciences, Warmadewa University | Lidwina Priliani, Edison Johar, Frilasita A Yudhaputri, Muhammad Rezki Rasyak, Willy Agustine, Hidayat Trimarsanto, Lydia V. Panggalo, Iskandar Adnan, Sukma Oktavianthi, Sri Masyeni, Erly Sintya, I Komang Parwata, Safarina G Malik, |
| EPI_ISL_2101231 | RSUD Klungkung | Eijkman Institute for Molecular Biology, National Agency for Research and Innovation; Molecular Biology Laboratory, Faculty Medicine and Health Sciences, Warmadewa University | Willy Agustine, Hidayat Trimarsanto, Lydia V. Panggalo, Iskandar Adnan, Sukma Oktavianthi, Lidwina Priliani, Edison Johar, Frilasita A Yudhaputri, Muhammad Rezki Rasyak, Sri Masyeni, Erly Sintya, I Komang Parwata, Safarina G Malik, |
| EPI_ISL_2101232 | RSUD Klungkung | Eijkman Institute for Molecular Biology, National Agency for Research and Innovation; Molecular Biology Laboratory, Faculty Medicine and Health Sciences, Warmadewa University | Lidwina Priliani, Edison Johar, Frilasita A Yudhaputri, Muhammad Rezki Rasyak, Willy Agustine, Hidayat Trimarsanto, Lydia V. Panggalo, Iskandar Adnan, Sukma Oktavianthi, Sri Masyeni, Erly Sintya, I Komang Parwata, Safarina G Malik, |
| EPI_ISL_2101233 | RSUD Klungkung | Eijkman Institute for Molecular Biology, National Agency for Research and Innovation; Molecular Biology Laboratory, Faculty Medicine and Health Sciences, Warmadewa University | Frilasita A Yudhaputri, Muhammad Rezki Rasyak, Willy Agustine, Hidayat Trimarsanto, Lydia V. Panggalo, Iskandar Adnan, Sukma Oktavianthi, Lidwina Priliani, Edison Johar, Sri Masyeni, Erly Sintya, I Komang Parwata, Safarina G Malik, |
| EPI_ISL_2101234 | RSUD Klungkung | Eijkman Institute for Molecular Biology, National Agency for Research and Innovation; Molecular Biology Laboratory, Faculty Medicine and Health Sciences, Warmadewa University | Muhammad Rezki Rasyak, Willy Agustine, Hidayat Trimarsanto, Lydia V. Panggalo, Iskandar Adnan, Sukma Oktavianthi, Lidwina Priliani, Edison Johar, Frilasita A Yudhaputri, Sri Masyeni, Erly Sintya, I Komang Parwata, Safarina G Malik, |
| EPI_ISL_2101235 | RSUD Klungkung | Eijkman Institute for Molecular Biology, National Agency for Research and Innovation; Molecular Biology Laboratory, Faculty Medicine and Health Sciences, Warmadewa University | Hidayat Trimarsanto, Lydia V. Panggalo, Iskandar Adnan, Sukma Oktavianthi, Lidwina Priliani, Edison Johar, Frilasita A Yudhaputri, Muhammad Rezki Rasyak, Willy Agustine, Sri Masyeni, Erly Sintya, I Komang Parwata, Safarina G Malik, |
| EPI_ISL_2101236 | RSUD Klungkung | Eijkman Institute for Molecular Biology, National Agency for Research and Innovation; Molecular Biology Laboratory, Faculty Medicine and Health Sciences, Warmadewa University | Frilasita A Yudhaputri, Muhammad Rezki Rasyak, Willy Agustine, Hidayat Trimarsanto, Lydia V. Panggalo, Iskandar Adnan, Sukma Oktavianthi, Lidwina Priliani, Edison Johar, Sri Masyeni, Erly Sintya, I Komang Parwata, Safarina G Malik, |
| EPI_ISL_2101237 | RSUD Klungkung | Eijkman Institute for Molecular Biology, National Agency for Research and Innovation; Molecular Biology Laboratory, Faculty | Willy Agustine, Hidayat Trimarsanto, Lydia V. Panggalo, Iskandar Adnan, Sukma Oktavianthi, Lidwina Priliani, Edison Johar, Frilasita A Yudhaputri, Muhammad Rezki Rasyak, Sri Masyeni, Erly Sintya, I Komang Parwata, Safarina G Malik, |

| | | | |
|---|---|---|---|
| | | Medicine and Health Sciences, Warmadewa University | |
| EPI_ISL_2101238 | RSUD Klungkung | Eijkman Institute for Molecular Biology, National Agency for Research and Innovation; Molecular Biology Laboratory, Faculty Medicine and Health Sciences, Warmadewa University | Iskandar Adnan, Sukma Oktavianthi, Lidwina Priliani, Edison Johar, Frilasita A Yudhaputri, Muhammad Rezki Rasyak, Willy Agustine, Hidayat Trimarsanto, Lydia V. Panggalo, Sri Masyeni, Erly Sintya, I Komang Parwata, Safarina G Malik, |
| EPI_ISL_2101239 | RSUD Klungkung | Eijkman Institute for Molecular Biology, National Agency for Research and Innovation; Molecular Biology Laboratory, Faculty Medicine and Health Sciences, Warmadewa University | Lidwina Priliani, Edison Johar, Frilasita A Yudhaputri, Muhammad Rezki Rasyak, Willy Agustine, Hidayat Trimarsanto, Lydia V. Panggalo, Iskandar Adnan, Sukma Oktavianthi, Sri Masyeni, Erly Sintya, I Komang Parwata, Safarina G Malik, |
| EPI_ISL_2101240 | RSUD Klungkung | Eijkman Institute for Molecular Biology, National Agency for Research and Innovation; Molecular Biology Laboratory, Faculty Medicine and Health Sciences, Warmadewa University | Edison Johar, Frilasita A Yudhaputri, Muhammad Rezki Rasyak, Willy Agustine, Hidayat Trimarsanto, Lydia V. Panggalo, Iskandar Adnan, Sukma Oktavianthi, Lidwina Priliani, Sri Masyeni, Erly Sintya, I Komang Parwata, Safarina G Malik, |
| EPI_ISL_2101241 | RSUD Klungkung | Eijkman Institute for Molecular Biology, National Agency for Research and Innovation; Molecular Biology Laboratory, Faculty Medicine and Health Sciences, Warmadewa University | Frilasita A Yudhaputri, Muhammad Rezki Rasyak, Willy Agustine, Hidayat Trimarsanto, Lydia V. Panggalo, Iskandar Adnan, Sukma Oktavianthi, Lidwina Priliani, Edison Johar, Sri Masyeni, Erly Sintya, I Komang Parwata, Safarina G Malik, |
| EPI_ISL_2101242 | RSUD Klungkung | Eijkman Institute for Molecular Biology, National Agency for Research and Innovation; Molecular Biology Laboratory, Faculty Medicine and Health Sciences, Warmadewa University | Muhammad Rezki Rasyak, Willy Agustine, Hidayat Trimarsanto, Lydia V. Panggalo, Iskandar Adnan, Sukma Oktavianthi, Lidwina Priliani, Edison Johar, Frilasita A Yudhaputri, Sri Masyeni, Erly Sintya, I Komang Parwata, Safarina G Malik, |
| EPI_ISL_2101243 | RSUD Klungkung | Eijkman Institute for Molecular Biology, National Agency for Research and Innovation; Molecular Biology Laboratory, Faculty Medicine and Health Sciences, Warmadewa University | Willy Agustine, Hidayat Trimarsanto, Lydia V. Panggalo, Iskandar Adnan, Sukma Oktavianthi, Lidwina Priliani, Edison Johar, Frilasita A Yudhaputri, Muhammad Rezki Rasyak, Sri Masyeni, Erly Sintya, I Komang Parwata, Safarina G Malik, |
| EPI_ISL_2101244 | RSUD Klungkung | Eijkman Institute for Molecular Biology, National Agency for Research and Innovation; Molecular Biology Laboratory, Faculty Medicine and Health Sciences, Warmadewa University | Lydia V. Panggalo, Iskandar Adnan, Sukma Oktavianthi, Lidwina Priliani, Edison Johar, Frilasita A Yudhaputri, Muhammad Rezki Rasyak, Willy Agustine, Hidayat Trimarsanto, Sri Masyeni, Erly Sintya, I Komang Parwata, Safarina G Malik, |
| EPI_ISL_2101245 | RSUD Klungkung | Eijkman Institute for Molecular Biology, National Agency for Research and Innovation; Molecular Biology Laboratory, Faculty Medicine and Health Sciences, Warmadewa University | Iskandar Adnan, Sukma Oktavianthi, Lidwina Priliani, Edison Johar, Frilasita A Yudhaputri, Muhammad Rezki Rasyak, Willy Agustine, Hidayat Trimarsanto, Lydia V. Panggalo, Sri Masyeni, Erly Sintya, I Komang Parwata, Safarina G Malik, |
| EPI_ISL_2101246 | RSUD Klungkung | Eijkman Institute for Molecular Biology, National Agency for Research and Innovation; Molecular Biology Laboratory, Faculty Medicine and Health Sciences, Warmadewa University | Sukma Oktavianthi, Lidwina Priliani, Edison Johar, Frilasita A Yudhaputri, Muhammad Rezki Rasyak, Willy Agustine, Hidayat Trimarsanto, Lydia V. Panggalo, Iskandar Adnan, Sri Masyeni, Erly Sintya, I Komang Parwata, Safarina G Malik, |
| EPI_ISL_2101247 | RSUD Klungkung | Eijkman Institute for Molecular Biology, National Agency for Research and Innovation; Molecular Biology Laboratory, Faculty | Lidwina Priliani, Edison Johar, Frilasita A Yudhaputri, Muhammad Rezki Rasyak, Willy Agustine, Hidayat Trimarsanto, Lydia V. Panggalo, Iskandar Adnan, Sukma Oktavianthi, Sri Masyeni, Erly Sintya, I Komang Parwata, Safarina G Malik, |

| | | | |
|---|---|---|---|
| | | Medicine and Health Sciences, Warmadewa University | |
| EPI_ISL_2101248 | RSUD Klungkung | Eijkman Institute for Molecular Biology, National Agency for Research and Innovation; Molecular Biology Laboratory, Faculty Medicine and Health Sciences, Warmadewa University | Frilasita A Yudhaputri, Muhammad Rezki Rasyak, Willy Agustine, Hidayat Trimarsanto, Lydia V. Panggalo, Iskandar Adnan, Sukma Oktavianthi, Lidwina Priliani, Edison Johar, Sri Masyeni, Erly Sintya, I Komang Parwata, Safarina G Malik, |
| EPI_ISL_2101249 | RSUD Klungkung | Eijkman Institute for Molecular Biology, National Agency for Research and Innovation; Molecular Biology Laboratory, Faculty Medicine and Health Sciences, Warmadewa University | Muhammad Rezki Rasyak, Willy Agustine, Hidayat Trimarsanto, Lydia V. Panggalo, Iskandar Adnan, Sukma Oktavianthi, Lidwina Priliani, Edison Johar, Frilasita A Yudhaputri, Sri Masyeni, Erly Sintya, I Komang Parwata, Safarina G Malik, |
| EPI_ISL_2101250 | RSUD Klungkung | Eijkman Institute for Molecular Biology, National Agency for Research and Innovation; Molecular Biology Laboratory, Faculty Medicine and Health Sciences, Warmadewa University | Willy Agustine, Hidayat Trimarsanto, Lydia V. Panggalo, Iskandar Adnan, Sukma Oktavianthi, Lidwina Priliani, Edison Johar, Frilasita A Yudhaputri, Muhammad Rezki Rasyak, Sri Masyeni, Erly Sintya, I Komang Parwata, Safarina G Malik, |
| EPI_ISL_2101251 | RSUD Klungkung | Eijkman Institute for Molecular Biology, National Agency for Research and Innovation; Molecular Biology Laboratory, Faculty Medicine and Health Sciences, Warmadewa University | Hidayat Trimarsanto, Lydia V. Panggalo, Iskandar Adnan, Sukma Oktavianthi, Lidwina Priliani, Edison Johar, Frilasita A Yudhaputri, Muhammad Rezki Rasyak, Willy Agustine, Sri Masyeni, Erly Sintya, I Komang Parwata, Safarina G Malik, |
| EPI_ISL_2101252 | RSUD Klungkung | Eijkman Institute for Molecular Biology, National Agency for Research and Innovation; Molecular Biology Laboratory, Faculty Medicine and Health Sciences, Warmadewa University | Lydia V. Panggalo, Iskandar Adnan, Sukma Oktavianthi, Lidwina Priliani, Edison Johar, Frilasita A Yudhaputri, Muhammad Rezki Rasyak, Willy Agustine, Hidayat Trimarsanto, Sri Masyeni, Erly Sintya, I Komang Parwata, Safarina G Malik, |
| EPI_ISL_2101253 | RSUD Klungkung | Eijkman Institute for Molecular Biology, National Agency for Research and Innovation; Molecular Biology Laboratory, Faculty Medicine and Health Sciences, Warmadewa University | Edison Johar, Frilasita A Yudhaputri, Muhammad Rezki Rasyak, Willy Agustine, Hidayat Trimarsanto, Lydia V. Panggalo, Iskandar Adnan, Sukma Oktavianthi, Lidwina Priliani, Sri Masyeni, Erly Sintya, I Komang Parwata, Safarina G Malik, |
| EPI_ISL_2101254 | RSUD Klungkung | Eijkman Institute for Molecular Biology, National Agency for Research and Innovation; Molecular Biology Laboratory, Faculty Medicine and Health Sciences, Warmadewa University | Hidayat Trimarsanto, Lydia V. Panggalo, Iskandar Adnan, Sukma Oktavianthi, Lidwina Priliani, Edison Johar, Frilasita A Yudhaputri, Muhammad Rezki Rasyak, Willy Agustine, Sri Masyeni, Erly Sintya, I Komang Parwata, Safarina G Malik, |
| EPI_ISL_2101255 | RSUD Klungkung | Eijkman Institute for Molecular Biology, National Agency for Research and Innovation; Molecular Biology Laboratory, Faculty Medicine and Health Sciences, Warmadewa University | Lydia V. Panggalo, Iskandar Adnan, Sukma Oktavianthi, Lidwina Priliani, Edison Johar, Frilasita A Yudhaputri, Muhammad Rezki Rasyak, Willy Agustine, Hidayat Trimarsanto, Sri Masyeni, Erly Sintya, I Komang Parwata, Safarina G Malik, |
| EPI_ISL_2101256 | RSUD Mangusada | Eijkman Institute for Molecular Biology, National Agency for Research and Innovation; Molecular Biology Laboratory, Faculty Medicine and Health Sciences, Warmadewa University | Edison Johar, Frilasita A Yudhaputri, Muhammad Rezki Rasyak, Willy Agustine, Hidayat Trimarsanto, Lydia V. Panggalo, Iskandar Adnan, Sukma Oktavianthi, Lidwina Priliani, Sri Masyeni, Erly Sintya, Ida Ayu Wayan Mahayani, Safarina G Ma |
| EPI_ISL_2101257 | RSUD Mangusada | Eijkman Institute for Molecular Biology, National Agency for Research and Innovation; Molecular Biology Laboratory, Faculty | Frilasita A Yudhaputri, Muhammad Rezki Rasyak, Willy Agustine, Hidayat Trimarsanto, Lydia V. Panggalo, Iskandar Adnan, Sukma Oktavianthi, Lidwina Priliani, Edison Johar, Sri Masyeni, Erly Sintya, Ida Ayu Wayan Mahayani, Safarina G Ma |

| | | | |
|---|---|---|---|
| | | Medicine and Health Sciences, Warmadewa University | |
| EPI_ISL_2101258 | RSUD Mangusada | Eijkman Institute for Molecular Biology, National Agency for Research and Innovation; Molecular Biology Laboratory, Faculty Medicine and Health Sciences, Warmadewa University | Muhammad Rezki Rasyak, Willy Agustine, Hidayat Trimarsanto, Lydia V. Panggalo, Iskandar Adnan, Sukma Oktavianthi, Lidwina Priliani, Edison Johar, Frilasita A Yudhaputri, Sri Masyeni, Erly Sintya, Ida Ayu Wayan Mahayani, Safarina G Ma |
| EPI_ISL_2101259 | RSUD Mangusada | Eijkman Institute for Molecular Biology, National Agency for Research and Innovation; Molecular Biology Laboratory, Faculty Medicine and Health Sciences, Warmadewa University | Willy Agustine, Hidayat Trimarsanto, Lydia V. Panggalo, Iskandar Adnan, Sukma Oktavianthi, Lidwina Priliani, Edison Johar, Frilasita A Yudhaputri, Muhammad Rezki Rasyak, Sri Masyeni, Erly Sintya, Ida Ayu Wayan Mahayani, Safarina G Ma |
| EPI_ISL_2101260 | RSUD Mangusada | Eijkman Institute for Molecular Biology, National Agency for Research and Innovation; Molecular Biology Laboratory, Faculty Medicine and Health Sciences, Warmadewa University | Hidayat Trimarsanto, Lydia V. Panggalo, Iskandar Adnan, Sukma Oktavianthi, Lidwina Priliani, Edison Johar, Frilasita A Yudhaputri, Muhammad Rezki Rasyak, Willy Agustine, Sri Masyeni, Erly Sintya, Ida Ayu Wayan Mahayani, Safarina G Ma |
| EPI_ISL_2101261 | RSUD Mangusada | Eijkman Institute for Molecular Biology, National Agency for Research and Innovation; Molecular Biology Laboratory, Faculty Medicine and Health Sciences, Warmadewa University | Lydia V. Panggalo, Iskandar Adnan, Sukma Oktavianthi, Lidwina Priliani, Edison Johar, Frilasita A Yudhaputri, Muhammad Rezki Rasyak, Willy Agustine, Hidayat Trimarsanto, Sri Masyeni, Erly Sintya, Ida Ayu Wayan Mahayani, Safarina G Ma |
| EPI_ISL_2101262 | RSUD Mangusada | Eijkman Institute for Molecular Biology, National Agency for Research and Innovation; Molecular Biology Laboratory, Faculty Medicine and Health Sciences, Warmadewa University | Iskandar Adnan, Sukma Oktavianthi, Lidwina Priliani, Edison Johar, Frilasita A Yudhaputri, Muhammad Rezki Rasyak, Willy Agustine, Hidayat Trimarsanto, Lydia V. Panggalo, Sri Masyeni, Erly Sintya, Ida Ayu Wayan Mahayani, Safarina G Ma |
| EPI_ISL_2101263 | RSUD Mangusada | Eijkman Institute for Molecular Biology, National Agency for Research and Innovation; Molecular Biology Laboratory, Faculty Medicine and Health Sciences, Warmadewa University | Sukma Oktavianthi, Lidwina Priliani, Edison Johar, Frilasita A Yudhaputri, Muhammad Rezki Rasyak, Willy Agustine, Hidayat Trimarsanto, Lydia V. Panggalo, Iskandar Adnan, Sri Masyeni, Erly Sintya, Ida Ayu Wayan Mahayani, Safarina G Ma |
| EPI_ISL_2101264 | RSUD Mangusada | Eijkman Institute for Molecular Biology, National Agency for Research and Innovation; Molecular Biology Laboratory, Faculty Medicine and Health Sciences, Warmadewa University | Lidwina Priliani, Edison Johar, Frilasita A Yudhaputri, Muhammad Rezki Rasyak, Willy Agustine, Hidayat Trimarsanto, Lydia V. Panggalo, Iskandar Adnan, Sukma Oktavianthi, Sri Masyeni, Erly Sintya, Ida Ayu Wayan Mahayani, Safarina G Ma |
| EPI_ISL_2101265 | RSUD Mangusada | Eijkman Institute for Molecular Biology, National Agency for Research and Innovation; Molecular Biology Laboratory, Faculty Medicine and Health Sciences, Warmadewa University | Edison Johar, Frilasita A Yudhaputri, Muhammad Rezki Rasyak, Willy Agustine, Hidayat Trimarsanto, Lydia V. Panggalo, Iskandar Adnan, Sukma Oktavianthi, Lidwina Priliani, Sri Masyeni, Erly Sintya, Ida Ayu Wayan Mahayani, Safarina G Ma |
| EPI_ISL_2101266 | RSUD Mangusada | Eijkman Institute for Molecular Biology, National Agency for Research and Innovation; Molecular Biology Laboratory, Faculty Medicine and Health Sciences, Warmadewa University | Frilasita A Yudhaputri, Muhammad Rezki Rasyak, Willy Agustine, Hidayat Trimarsanto, Lydia V. Panggalo, Iskandar Adnan, Sukma Oktavianthi, Lidwina Priliani, Edison Johar, Sri Masyeni, Erly Sintya, Ida Ayu Wayan Mahayani, Safarina G Ma |
| EPI_ISL_2101267 | RSUD Mangusada | Eijkman Institute for Molecular Biology, National Agency for Research and Innovation; Molecular Biology Laboratory, Faculty | Muhammad Rezki Rasyak, Willy Agustine, Hidayat Trimarsanto, Lydia V. Panggalo, Iskandar Adnan, Sukma Oktavianthi, Lidwina Priliani, Edison Johar, Frilasita A Yudhaputri, Sri Masyeni, Erly Sintya, Ida Ayu Wayan Mahayani, Safarina G Ma |

| | | | |
|---|---|---|---|
| | | Medicine and Health Sciences, Warmadewa University | |
| EPI_ISL_2101268 | RSUD Mangusada | Eijkman Institute for Molecular Biology, National Agency for Research and Innovation; Molecular Biology Laboratory, Faculty Medicine and Health Sciences, Warmadewa University | Willy Agustine, Hidayat Trimarsanto, Lydia V. Panggalo, Iskandar Adnan, Sukma Oktavianthi, Lidwina Priliani, Edison Johar, Frilasita A Yudhaputri, Muhammad Rezki Rasyak, Sri Masyeni, Erly Sintya, Ida Ayu Wayan Mahayani, Safarina G Ma |
| EPI_ISL_2101269 | RSUD Mangusada | Eijkman Institute for Molecular Biology, National Agency for Research and Innovation; Molecular Biology Laboratory, Faculty Medicine and Health Sciences, Warmadewa University | Hidayat Trimarsanto, Lydia V. Panggalo, Iskandar Adnan, Sukma Oktavianthi, Lidwina Priliani, Edison Johar, Frilasita A Yudhaputri, Muhammad Rezki Rasyak, Willy Agustine, Sri Masyeni, Erly Sintya, Ida Ayu Wayan Mahayani, Safarina G Ma |
| EPI_ISL_2101270 | RSUD Mangusada | Eijkman Institute for Molecular Biology, National Agency for Research and Innovation; Molecular Biology Laboratory, Faculty Medicine and Health Sciences, Warmadewa University | Lydia V. Panggalo, Iskandar Adnan, Sukma Oktavianthi, Lidwina Priliani, Edison Johar, Frilasita A Yudhaputri, Muhammad Rezki Rasyak, Willy Agustine, Hidayat Trimarsanto, Sri Masyeni, Erly Sintya, Ida Ayu Wayan Mahayani, Safarina G Ma |
| EPI_ISL_2101271 | RSUD Mangusada | Eijkman Institute for Molecular Biology, National Agency for Research and Innovation; Molecular Biology Laboratory, Faculty Medicine and Health Sciences, Warmadewa University | Iskandar Adnan, Sukma Oktavianthi, Lidwina Priliani, Edison Johar, Frilasita A Yudhaputri, Muhammad Rezki Rasyak, Willy Agustine, Hidayat Trimarsanto, Lydia V. Panggalo, Sri Masyeni, Erly Sintya, Ida Ayu Wayan Mahayani, Safarina G Ma |
| EPI_ISL_2101272 | RSUD Mangusada | Eijkman Institute for Molecular Biology, National Agency for Research and Innovation; Molecular Biology Laboratory, Faculty Medicine and Health Sciences, Warmadewa University | Sukma Oktavianthi, Lidwina Priliani, Edison Johar, Frilasita A Yudhaputri, Muhammad Rezki Rasyak, Willy Agustine, Hidayat Trimarsanto, Lydia V. Panggalo, Iskandar Adnan, Sri Masyeni, Erly Sintya, Ida Ayu Wayan Mahayani, Safarina G Ma |
| EPI_ISL_2101273 | RSUD Mangusada | Eijkman Institute for Molecular Biology, National Agency for Research and Innovation; Molecular Biology Laboratory, Faculty Medicine and Health Sciences, Warmadewa University | Lidwina Priliani, Edison Johar, Frilasita A Yudhaputri, Muhammad Rezki Rasyak, Willy Agustine, Hidayat Trimarsanto, Lydia V. Panggalo, Iskandar Adnan, Sukma Oktavianthi, Sri Masyeni, Erly Sintya, Ida Ayu Wayan Mahayani, Safarina G Ma |
| EPI_ISL_2101274 | RSUD Mangusada | Eijkman Institute for Molecular Biology, National Agency for Research and Innovation; Molecular Biology Laboratory, Faculty Medicine and Health Sciences, Warmadewa University | Edison Johar, Frilasita A Yudhaputri, Muhammad Rezki Rasyak, Willy Agustine, Hidayat Trimarsanto, Lydia V. Panggalo, Iskandar Adnan, Sukma Oktavianthi, Lidwina Priliani, Sri Masyeni, Erly Sintya, Ida Ayu Wayan Mahayani, Safarina G Ma |
| EPI_ISL_2101275 | RSUD Mangusada | Eijkman Institute for Molecular Biology, National Agency for Research and Innovation; Molecular Biology Laboratory, Faculty Medicine and Health Sciences, Warmadewa University | Frilasita A Yudhaputri, Muhammad Rezki Rasyak, Willy Agustine, Hidayat Trimarsanto, Lydia V. Panggalo, Iskandar Adnan, Sukma Oktavianthi, Lidwina Priliani, Edison Johar, Sri Masyeni, Erly Sintya, Ida Ayu Wayan Mahayani, Safarina G Ma |
| EPI_ISL_2101276 | RSUD Mangusada | Eijkman Institute for Molecular Biology, National Agency for Research and Innovation; Molecular Biology Laboratory, Faculty Medicine and Health Sciences, Warmadewa University | Muhammad Rezki Rasyak, Willy Agustine, Hidayat Trimarsanto, Lydia V. Panggalo, Iskandar Adnan, Sukma Oktavianthi, Lidwina Priliani, Edison Johar, Frilasita A Yudhaputri, Sri Masyeni, Erly Sintya, Ida Ayu Wayan Mahayani, Safarina G Ma |
| EPI_ISL_2101277 | RSUD Mangusada | Eijkman Institute for Molecular Biology, National Agency for Research and Innovation; Molecular Biology Laboratory, Faculty | Willy Agustine, Hidayat Trimarsanto, Lydia V. Panggalo, Iskandar Adnan, Sukma Oktavianthi, Lidwina Priliani, Edison Johar, Frilasita A Yudhaputri, Muhammad Rezki Rasyak, Sri Masyeni, Erly Sintya, Ida Ayu Wayan Mahayani, Safarina G Ma |

| | | | |
|---|---|---|---|
| | | Medicine and Health Sciences, Warmadewa University | |
| EPI_ISL_2101278 | RSUD Mangusada | Eijkman Institute for Molecular Biology, National Agency for Research and Innovation; Molecular Biology Laboratory, Faculty Medicine and Health Sciences, Warmadewa University | Hidayat Trimarsanto, Lydia V. Panggalo, Iskandar Adnan, Sukma Oktavianthi, Lidwina Priliani, Edison Johar, Frilasita A Yudhaputri, Muhammad Rezki Rasyak, Willy Agustine, Sri Masyeni, Erly Sintya, Ida Ayu Wayan Mahayani, Safarina G Ma |
| EPI_ISL_2101279 | RSUD Mangusada | Eijkman Institute for Molecular Biology, National Agency for Research and Innovation; Molecular Biology Laboratory, Faculty Medicine and Health Sciences, Warmadewa University | Lydia V. Panggalo, Iskandar Adnan, Sukma Oktavianthi, Lidwina Priliani, Edison Johar, Frilasita A Yudhaputri, Muhammad Rezki Rasyak, Willy Agustine, Hidayat Trimarsanto, Sri Masyeni, Erly Sintya, Ida Ayu Wayan Mahayani, Safarina G Ma |
| EPI_ISL_2101280 | RSUD Mangusada | Eijkman Institute for Molecular Biology, National Agency for Research and Innovation; Molecular Biology Laboratory, Faculty Medicine and Health Sciences, Warmadewa University | Iskandar Adnan, Sukma Oktavianthi, Lidwina Priliani, Edison Johar, Frilasita A Yudhaputri, Muhammad Rezki Rasyak, Willy Agustine, Hidayat Trimarsanto, Lydia V. Panggalo, Sri Masyeni, Erly Sintya, Ida Ayu Wayan Mahayani, Safarina G Ma |
| EPI_ISL_2101281 | RSUD Mangusada | Eijkman Institute for Molecular Biology, National Agency for Research and Innovation; Molecular Biology Laboratory, Faculty Medicine and Health Sciences, Warmadewa University | Sukma Oktavianthi, Lidwina Priliani, Edison Johar, Frilasita A Yudhaputri, Muhammad Rezki Rasyak, Willy Agustine, Hidayat Trimarsanto, Lydia V. Panggalo, Iskandar Adnan, Sri Masyeni, Erly Sintya, Ida Ayu Wayan Mahayani, Safarina G Ma |
| EPI_ISL_2101282 | RSUD Mangusada | Eijkman Institute for Molecular Biology, National Agency for Research and Innovation; Molecular Biology Laboratory, Faculty Medicine and Health Sciences, Warmadewa University | Lidwina Priliani, Edison Johar, Frilasita A Yudhaputri, Muhammad Rezki Rasyak, Willy Agustine, Hidayat Trimarsanto, Lydia V. Panggalo, Iskandar Adnan, Sukma Oktavianthi, Sri Masyeni, Erly Sintya, Ida Ayu Wayan Mahayani, Safarina G Ma |
| EPI_ISL_2101283 | RSUD Mangusada | Eijkman Institute for Molecular Biology, National Agency for Research and Innovation; Molecular Biology Laboratory, Faculty Medicine and Health Sciences, Warmadewa University | Edison Johar, Frilasita A Yudhaputri, Muhammad Rezki Rasyak, Willy Agustine, Hidayat Trimarsanto, Lydia V. Panggalo, Iskandar Adnan, Sukma Oktavianthi, Lidwina Priliani, Sri Masyeni, Erly Sintya, Ida Ayu Wayan Mahayani, Safarina G Ma |
| EPI_ISL_2101284 | RSUD Mangusada | Eijkman Institute for Molecular Biology, National Agency for Research and Innovation; Molecular Biology Laboratory, Faculty Medicine and Health Sciences, Warmadewa University | Frilasita A Yudhaputri, Muhammad Rezki Rasyak, Willy Agustine, Hidayat Trimarsanto, Lydia V. Panggalo, Iskandar Adnan, Sukma Oktavianthi, Lidwina Priliani, Edison Johar, Sri Masyeni, Erly Sintya, Ida Ayu Wayan Mahayani, Safarina G Ma |
| EPI_ISL_2101285 | RSUD Mangusada | Eijkman Institute for Molecular Biology, National Agency for Research and Innovation; Molecular Biology Laboratory, Faculty Medicine and Health Sciences, Warmadewa University | Muhammad Rezki Rasyak, Willy Agustine, Hidayat Trimarsanto, Lydia V. Panggalo, Iskandar Adnan, Sukma Oktavianthi, Lidwina Priliani, Edison Johar, Frilasita A Yudhaputri, Sri Masyeni, Erly Sintya, Ida Ayu Wayan Mahayani, Safarina G Ma |
| EPI_ISL_2101286 | RSUD Mangusada | Eijkman Institute for Molecular Biology, National Agency for Research and Innovation; Molecular Biology Laboratory, Faculty Medicine and Health Sciences, Warmadewa University | Willy Agustine, Hidayat Trimarsanto, Lydia V. Panggalo, Iskandar Adnan, Sukma Oktavianthi, Lidwina Priliani, Edison Johar, Frilasita A Yudhaputri, Muhammad Rezki Rasyak, Sri Masyeni, Erly Sintya, Ida Ayu Wayan Mahayani, Safarina G Ma |
| EPI_ISL_2101287 | RSUD Mangusada | Eijkman Institute for Molecular Biology, National Agency for Research and Innovation; Molecular Biology Laboratory, Faculty | Hidayat Trimarsanto, Lydia V. Panggalo, Iskandar Adnan, Sukma Oktavianthi, Lidwina Priliani, Edison Johar, Frilasita A Yudhaputri, Muhammad Rezki Rasyak, Willy Agustine, Sri Masyeni, Erly Sintya, Ida Ayu Wayan Mahayani, Safarina G Ma |

| | | | |
|---|---|---|---|
| | | Medicine and Health Sciences, Warmadewa University | |
| EPI_ISL_2101288 | RSUD Mangusada | Eijkman Institute for Molecular Biology, National Agency for Research and Innovation; Molecular Biology Laboratory, Faculty Medicine and Health Sciences, Warmadewa University | Lydia V. Panggalo, Iskandar Adnan, Sukma Oktavianthi, Lidwina Priliani, Edison Johar, Frilasita A Yudhaputri, Muhammad Rezki Rasyak, Willy Agustine, Hidayat Trimarsanto, Sri Masyeni, Erly Sintya, Ida Ayu Wayan Mahayani, Safarina G Ma |
| EPI_ISL_2101290 | RSUD Mangusada | Eijkman Institute for Molecular Biology, National Agency for Research and Innovation; Molecular Biology Laboratory, Faculty Medicine and Health Sciences, Warmadewa University | Iskandar Adnan, Sukma Oktavianthi, Lidwina Priliani, Edison Johar, Frilasita A Yudhaputri, Muhammad Rezki Rasyak, Willy Agustine, Hidayat Trimarsanto, Lydia V. Panggalo, Sri Masyeni, Erly Sintya, Ida Ayu Wayan Mahayani, Safarina G Ma |
| EPI_ISL_2101291 | RSUD Mangusada | Eijkman Institute for Molecular Biology, National Agency for Research and Innovation; Molecular Biology Laboratory, Faculty Medicine and Health Sciences, Warmadewa University | Sukma Oktavianthi, Lidwina Priliani, Edison Johar, Frilasita A Yudhaputri, Muhammad Rezki Rasyak, Willy Agustine, Hidayat Trimarsanto, Lydia V. Panggalo, Iskandar Adnan, Sri Masyeni, Erly Sintya, Ida Ayu Wayan Mahayani, Safarina G Ma |
| EPI_ISL_2101292 | RSUD Mangusada | Eijkman Institute for Molecular Biology, National Agency for Research and Innovation; Molecular Biology Laboratory, Faculty Medicine and Health Sciences, Warmadewa University | Frilasita A Yudhaputri, Muhammad Rezki Rasyak, Willy Agustine, Hidayat Trimarsanto, Lydia V. Panggalo, Iskandar Adnan, Sukma Oktavianthi, Lidwina Priliani, Edison Johar, Sri Masyeni, Erly Sintya, Ida Ayu Wayan Mahayani, Safarina G Ma |
| EPI_ISL_2101293 | RSUD Mangusada | Eijkman Institute for Molecular Biology, National Agency for Research and Innovation; Molecular Biology Laboratory, Faculty Medicine and Health Sciences, Warmadewa University | Muhammad Rezki Rasyak, Willy Agustine, Hidayat Trimarsanto, Lydia V. Panggalo, Iskandar Adnan, Sukma Oktavianthi, Lidwina Priliani, Edison Johar, Frilasita A Yudhaputri, Sri Masyeni, Erly Sintya, Ida Ayu Wayan Mahayani, Safarina G Ma |
| EPI_ISL_2101294 | RSUD Mangusada | Eijkman Institute for Molecular Biology, National Agency for Research and Innovation; Molecular Biology Laboratory, Faculty Medicine and Health Sciences, Warmadewa University | Willy Agustine, Hidayat Trimarsanto, Lydia V. Panggalo, Iskandar Adnan, Sukma Oktavianthi, Lidwina Priliani, Edison Johar, Frilasita A Yudhaputri, Muhammad Rezki Rasyak, Sri Masyeni, Erly Sintya, Ida Ayu Wayan Mahayani, Safarina G Ma |
| EPI_ISL_2101295 | RSUD Mangusada | Eijkman Institute for Molecular Biology, National Agency for Research and Innovation; Molecular Biology Laboratory, Faculty Medicine and Health Sciences, Warmadewa University | Hidayat Trimarsanto, Lydia V. Panggalo, Iskandar Adnan, Sukma Oktavianthi, Lidwina Priliani, Edison Johar, Frilasita A Yudhaputri, Muhammad Rezki Rasyak, Willy Agustine, Sri Masyeni, Erly Sintya, Ida Ayu Wayan Mahayani, Safarina G Ma |
| EPI_ISL_2101296 | RSUD Mangusada | Eijkman Institute for Molecular Biology, National Agency for Research and Innovation; Molecular Biology Laboratory, Faculty Medicine and Health Sciences, Warmadewa University | Lydia V. Panggalo, Iskandar Adnan, Sukma Oktavianthi, Lidwina Priliani, Edison Johar, Frilasita A Yudhaputri, Muhammad Rezki Rasyak, Willy Agustine, Hidayat Trimarsanto, Sri Masyeni, Erly Sintya, Ida Ayu Wayan Mahayani, Safarina G Ma |
| EPI_ISL_2101297 | RSUD Mangusada | Eijkman Institute for Molecular Biology, National Agency for Research and Innovation; Molecular Biology Laboratory, Faculty Medicine and Health Sciences, Warmadewa University | Edison Johar, Frilasita A Yudhaputri, Muhammad Rezki Rasyak, Willy Agustine, Hidayat Trimarsanto, Lydia V. Panggalo, Iskandar Adnan, Sukma Oktavianthi, Lidwina Priliani, Sri Masyeni, Erly Sintya, Ida Ayu Wayan Mahayani, Safarina G Ma |
| EPI_ISL_2101298 | RSUD Mangusada | Eijkman Institute for Molecular Biology, National Agency for Research and Innovation; Molecular Biology Laboratory, Faculty | Muhammad Rezki Rasyak, Willy Agustine, Hidayat Trimarsanto, Lydia V. Panggalo, Iskandar Adnan, Sukma Oktavianthi, Lidwina Priliani, Edison Johar, Frilasita A Yudhaputri, Sri Masyeni, Erly Sintya, Ida Ayu Wayan Mahayani, Safarina G Ma |

| | | | |
|---|---|---|---|
| | | Medicine and Health Sciences, Warmadewa University | |
| EPI_ISL_2101299 | RSUD Mangusada | Eijkman Institute for Molecular Biology, National Agency for Research and Innovation; Molecular Biology Laboratory, Faculty Medicine and Health Sciences, Warmadewa University | Willy Agustine, Hidayat Trimarsanto, Lydia V. Panggalo, Iskandar Adnan, Sukma Oktavianthi, Lidwina Priliani, Edison Johar, Frilasita A Yudhaputri, Muhammad Rezki Rasyak, Sri Masyeni, Erly Sintya, Ida Ayu Wayan Mahayani, Safarina G Ma |
| EPI_ISL_2101300 | RSUD Mangusada | Eijkman Institute for Molecular Biology, National Agency for Research and Innovation; Molecular Biology Laboratory, Faculty Medicine and Health Sciences, Warmadewa University | Hidayat Trimarsanto, Lydia V. Panggalo, Iskandar Adnan, Sukma Oktavianthi, Lidwina Priliani, Edison Johar, Frilasita A Yudhaputri, Muhammad Rezki Rasyak, Willy Agustine, Sri Masyeni, Erly Sintya, Ida Ayu Wayan Mahayani, Safarina G Ma |
| EPI_ISL_2101301 | RSUD Mangusada | Eijkman Institute for Molecular Biology, National Agency for Research and Innovation; Molecular Biology Laboratory, Faculty Medicine and Health Sciences, Warmadewa University | Lydia V. Panggalo, Iskandar Adnan, Sukma Oktavianthi, Lidwina Priliani, Edison Johar, Frilasita A Yudhaputri, Muhammad Rezki Rasyak, Willy Agustine, Hidayat Trimarsanto, Sri Masyeni, Erly Sintya, Ida Ayu Wayan Mahayani, Safarina G Ma |
| EPI_ISL_2101302 | RSUD Mangusada | Eijkman Institute for Molecular Biology, National Agency for Research and Innovation; Molecular Biology Laboratory, Faculty Medicine and Health Sciences, Warmadewa University | Iskandar Adnan, Sukma Oktavianthi, Lidwina Priliani, Edison Johar, Frilasita A Yudhaputri, Muhammad Rezki Rasyak, Willy Agustine, Hidayat Trimarsanto, Lydia V. Panggalo, Sri Masyeni, Erly Sintya, Ida Ayu Wayan Mahayani, Safarina G Ma |
| EPI_ISL_2101303 | RSUD Mangusada | Eijkman Institute for Molecular Biology, National Agency for Research and Innovation; Molecular Biology Laboratory, Faculty Medicine and Health Sciences, Warmadewa University | Sukma Oktavianthi, Lidwina Priliani, Edison Johar, Frilasita A Yudhaputri, Muhammad Rezki Rasyak, Willy Agustine, Hidayat Trimarsanto, Lydia V. Panggalo, Iskandar Adnan, Sri Masyeni, Erly Sintya, Ida Ayu Wayan Mahayani, Safarina G Ma |
| EPI_ISL_2101304 | RSUD Mangusada | Eijkman Institute for Molecular Biology, National Agency for Research and Innovation; Molecular Biology Laboratory, Faculty Medicine and Health Sciences, Warmadewa University | Lidwina Priliani, Edison Johar, Frilasita A Yudhaputri, Muhammad Rezki Rasyak, Willy Agustine, Hidayat Trimarsanto, Lydia V. Panggalo, Iskandar Adnan, Sukma Oktavianthi, Sri Masyeni, Erly Sintya, Ida Ayu Wayan Mahayani, Safarina G Ma |
| EPI_ISL_2101305 | RSUD Mangusada | Eijkman Institute for Molecular Biology, National Agency for Research and Innovation; Molecular Biology Laboratory, Faculty Medicine and Health Sciences, Warmadewa University | Edison Johar, Frilasita A Yudhaputri, Muhammad Rezki Rasyak, Willy Agustine, Hidayat Trimarsanto, Lydia V. Panggalo, Iskandar Adnan, Sukma Oktavianthi, Lidwina Priliani, Sri Masyeni, Erly Sintya, Ida Ayu Wayan Mahayani, Safarina G Ma |
| EPI_ISL_2101306 | RSUD Mangusada | Eijkman Institute for Molecular Biology, National Agency for Research and Innovation; Molecular Biology Laboratory, Faculty Medicine and Health Sciences, Warmadewa University | Frilasita A Yudhaputri, Muhammad Rezki Rasyak, Willy Agustine, Hidayat Trimarsanto, Lydia V. Panggalo, Iskandar Adnan, Sukma Oktavianthi, Lidwina Priliani, Edison Johar, Sri Masyeni, Erly Sintya, Ida Ayu Wayan Mahayani, Safarina G Ma |
| EPI_ISL_2101307 | RSUD Mangusada | Eijkman Institute for Molecular Biology, National Agency for Research and Innovation; Molecular Biology Laboratory, Faculty Medicine and Health Sciences, Warmadewa University | Muhammad Rezki Rasyak, Willy Agustine, Hidayat Trimarsanto, Lydia V. Panggalo, Iskandar Adnan, Sukma Oktavianthi, Lidwina Priliani, Edison Johar, Frilasita A Yudhaputri, Sri Masyeni, Erly Sintya, Ida Ayu Wayan Mahayani, Safarina G Ma |
| EPI_ISL_2101308 | RSUD Mangusada | Eijkman Institute for Molecular Biology, National Agency for Research and Innovation; Molecular Biology Laboratory, Faculty | Willy Agustine, Hidayat Trimarsanto, Lydia V. Panggalo, Iskandar Adnan, Sukma Oktavianthi, Lidwina Priliani, Edison Johar, Frilasita A Yudhaputri, Muhammad Rezki Rasyak, Sri Masyeni, Erly Sintya, Ida Ayu Wayan Mahayani, Safarina G Ma |

| | | | |
|---|---|---|---|
| | | Medicine and Health Sciences, Warmadewa University | |
| EPI_ISL_2101309 | RSUD Mangusada | Eijkman Institute for Molecular Biology, National Agency for Research and Innovation; Molecular Biology Laboratory, Faculty Medicine and Health Sciences, Warmadewa University | Hidayat Trimarsanto, Lydia V. Panggalo, Iskandar Adnan, Sukma Oktavianthi, Lidwina Priliani, Edison Johar, Frilasita A Yudhaputri, Muhammad Rezki Rasyak, Willy Agustine, Sri Masyeni, Erly Sintya, Ida Ayu Wayan Mahayani, Safarina G Ma |
| EPI_ISL_2101310 | RSUD Mangusada | Eijkman Institute for Molecular Biology, National Agency for Research and Innovation; Molecular Biology Laboratory, Faculty Medicine and Health Sciences, Warmadewa University | Lydia V. Panggalo, Iskandar Adnan, Sukma Oktavianthi, Lidwina Priliani, Edison Johar, Frilasita A Yudhaputri, Muhammad Rezki Rasyak, Willy Agustine, Hidayat Trimarsanto, Sri Masyeni, Erly Sintya, Ida Ayu Wayan Mahayani, Safarina G Ma |
| EPI_ISL_2101311 | RSUD Mangusada | Eijkman Institute for Molecular Biology, National Agency for Research and Innovation; Molecular Biology Laboratory, Faculty Medicine and Health Sciences, Warmadewa University | Iskandar Adnan, Sukma Oktavianthi, Lidwina Priliani, Edison Johar, Frilasita A Yudhaputri, Muhammad Rezki Rasyak, Willy Agustine, Hidayat Trimarsanto, Lydia V. Panggalo, Sri Masyeni, Erly Sintya, Ida Ayu Wayan Mahayani, Safarina G Ma |
| EPI_ISL_2101312 | RSUD Mangusada | Eijkman Institute for Molecular Biology, National Agency for Research and Innovation; Molecular Biology Laboratory, Faculty Medicine and Health Sciences, Warmadewa University | Sukma Oktavianthi, Lidwina Priliani, Edison Johar, Frilasita A Yudhaputri, Muhammad Rezki Rasyak, Willy Agustine, Hidayat Trimarsanto, Lydia V. Panggalo, Iskandar Adnan, Sri Masyeni, Erly Sintya, Ida Ayu Wayan Mahayani, Safarina G Ma |
| EPI_ISL_2101313 | RSUD Mangusada | Eijkman Institute for Molecular Biology, National Agency for Research and Innovation; Molecular Biology Laboratory, Faculty Medicine and Health Sciences, Warmadewa University | Edison Johar, Frilasita A Yudhaputri, Muhammad Rezki Rasyak, Willy Agustine, Hidayat Trimarsanto, Lydia V. Panggalo, Iskandar Adnan, Sukma Oktavianthi, Lidwina Priliani, Sri Masyeni, Erly Sintya, Ida Ayu Wayan Mahayani, Safarina G Ma |
| EPI_ISL_2101314 | RSUD Mangusada | Eijkman Institute for Molecular Biology, National Agency for Research and Innovation; Molecular Biology Laboratory, Faculty Medicine and Health Sciences, Warmadewa University | Frilasita A Yudhaputri, Muhammad Rezki Rasyak, Willy Agustine, Hidayat Trimarsanto, Lydia V. Panggalo, Iskandar Adnan, Sukma Oktavianthi, Lidwina Priliani, Edison Johar, Sri Masyeni, Erly Sintya, Ida Ayu Wayan Mahayani, Safarina G Ma |
| EPI_ISL_2101315 | RSUD Mangusada | Eijkman Institute for Molecular Biology, National Agency for Research and Innovation; Molecular Biology Laboratory, Faculty Medicine and Health Sciences, Warmadewa University | Muhammad Rezki Rasyak, Willy Agustine, Hidayat Trimarsanto, Lydia V. Panggalo, Iskandar Adnan, Sukma Oktavianthi, Lidwina Priliani, Edison Johar, Frilasita A Yudhaputri, Sri Masyeni, Erly Sintya, Ida Ayu Wayan Mahayani, Safarina G Ma |
| EPI_ISL_2101316 | RSUD Mangusada | Eijkman Institute for Molecular Biology, National Agency for Research and Innovation; Molecular Biology Laboratory, Faculty Medicine and Health Sciences, Warmadewa University | Willy Agustine, Hidayat Trimarsanto, Lydia V. Panggalo, Iskandar Adnan, Sukma Oktavianthi, Lidwina Priliani, Edison Johar, Frilasita A Yudhaputri, Muhammad Rezki Rasyak, Sri Masyeni, Erly Sintya, Ida Ayu Wayan Mahayani, Safarina G Ma |
| EPI_ISL_2101317 | RSUD Mangusada | Eijkman Institute for Molecular Biology, National Agency for Research and Innovation; Molecular Biology Laboratory, Faculty Medicine and Health Sciences, Warmadewa University | Hidayat Trimarsanto, Lydia V. Panggalo, Iskandar Adnan, Sukma Oktavianthi, Lidwina Priliani, Edison Johar, Frilasita A Yudhaputri, Muhammad Rezki Rasyak, Willy Agustine, Sri Masyeni, Erly Sintya, Ida Ayu Wayan Mahayani, Safarina G Ma |
| EPI_ISL_2101318 | RSUD Mangusada | Eijkman Institute for Molecular Biology, National Agency for Research and Innovation; Molecular Biology Laboratory, Faculty | Iskandar Adnan, Sukma Oktavianthi, Lidwina Priliani, Edison Johar, Frilasita A Yudhaputri, Muhammad Rezki Rasyak, Willy Agustine, Hidayat Trimarsanto, Lydia V. Panggalo, Sri Masyeni, Erly Sintya, Ida Ayu Wayan Mahayani, Safarina G Ma |

| | | | |
|---|---|---|---|
| | | Medicine and Health Sciences, Warmadewa University | |
| EPI_ISL_2101319 | RSUD Mangusada | Eijkman Institute for Molecular Biology, National Agency for Research and Innovation; Molecular Biology Laboratory, Faculty Medicine and Health Sciences, Warmadewa University | Lidwina Priliani, Edison Johar, Frilasita A Yudhaputri, Muhammad Rezki Rasyak, Willy Agustine, Hidayat Trimarsanto, Lydia V. Panggalo, Iskandar Adnan, Sukma Oktavianthi, Sri Masyeni, Erly Sintya, Ida Ayu Wayan Mahayani, Safarina G Ma |
| EPI_ISL_2101320 | RSUD Mangusada | Eijkman Institute for Molecular Biology, National Agency for Research and Innovation; Molecular Biology Laboratory, Faculty Medicine and Health Sciences, Warmadewa University | Edison Johar, Frilasita A Yudhaputri, Muhammad Rezki Rasyak, Willy Agustine, Hidayat Trimarsanto, Lydia V. Panggalo, Iskandar Adnan, Sukma Oktavianthi, Lidwina Priliani, Sri Masyeni, Erly Sintya, Ida Ayu Wayan Mahayani, Safarina G Ma |
| EPI_ISL_2101321 | RSUD Mangusada | Eijkman Institute for Molecular Biology, National Agency for Research and Innovation; Molecular Biology Laboratory, Faculty Medicine and Health Sciences, Warmadewa University | Frilasita A Yudhaputri, Muhammad Rezki Rasyak, Willy Agustine, Hidayat Trimarsanto, Lydia V. Panggalo, Iskandar Adnan, Sukma Oktavianthi, Lidwina Priliani, Edison Johar, Sri Masyeni, Erly Sintya, Ida Ayu Wayan Mahayani, Safarina G Ma |
| EPI_ISL_2101322 | RSUD Mangusada | Eijkman Institute for Molecular Biology, National Agency for Research and Innovation; Molecular Biology Laboratory, Faculty Medicine and Health Sciences, Warmadewa University | Muhammad Rezki Rasyak, Willy Agustine, Hidayat Trimarsanto, Lydia V. Panggalo, Iskandar Adnan, Sukma Oktavianthi, Lidwina Priliani, Edison Johar, Frilasita A Yudhaputri, Sri Masyeni, Erly Sintya, Ida Ayu Wayan Mahayani, Safarina G Ma |
| EPI_ISL_2101323 | RSUD Mangusada | Eijkman Institute for Molecular Biology, National Agency for Research and Innovation; Molecular Biology Laboratory, Faculty Medicine and Health Sciences, Warmadewa University | Willy Agustine, Hidayat Trimarsanto, Lydia V. Panggalo, Iskandar Adnan, Sukma Oktavianthi, Lidwina Priliani, Edison Johar, Frilasita A Yudhaputri, Muhammad Rezki Rasyak, Sri Masyeni, Erly Sintya, Ida Ayu Wayan Mahayani, Safarina G Ma |
| EPI_ISL_2101324 | RSUD Mangusada | Eijkman Institute for Molecular Biology, National Agency for Research and Innovation; Molecular Biology Laboratory, Faculty Medicine and Health Sciences, Warmadewa University | Hidayat Trimarsanto, Lydia V. Panggalo, Iskandar Adnan, Sukma Oktavianthi, Lidwina Priliani, Edison Johar, Frilasita A Yudhaputri, Muhammad Rezki Rasyak, Willy Agustine, Sri Masyeni, Erly Sintya, Ida Ayu Wayan Mahayani, Safarina G Ma |
| EPI_ISL_2101325 | RSUD Mangusada | Eijkman Institute for Molecular Biology, National Agency for Research and Innovation; Molecular Biology Laboratory, Faculty Medicine and Health Sciences, Warmadewa University | Lydia V. Panggalo, Iskandar Adnan, Sukma Oktavianthi, Lidwina Priliani, Edison Johar, Frilasita A Yudhaputri, Muhammad Rezki Rasyak, Willy Agustine, Hidayat Trimarsanto, Sri Masyeni, Erly Sintya, Ida Ayu Wayan Mahayani, Safarina G Ma |
| EPI_ISL_2101326 | RSUD Mangusada | Eijkman Institute for Molecular Biology, National Agency for Research and Innovation; Molecular Biology Laboratory, Faculty Medicine and Health Sciences, Warmadewa University | Iskandar Adnan, Sukma Oktavianthi, Lidwina Priliani, Edison Johar, Frilasita A Yudhaputri, Muhammad Rezki Rasyak, Willy Agustine, Hidayat Trimarsanto, Lydia V. Panggalo, Sri Masyeni, Erly Sintya, Ida Ayu Wayan Mahayani, Safarina G Ma |
| EPI_ISL_2101327 | RSUD Mangusada | Eijkman Institute for Molecular Biology, National Agency for Research and Innovation; Molecular Biology Laboratory, Faculty Medicine and Health Sciences, Warmadewa University | Sukma Oktavianthi, Lidwina Priliani, Edison Johar, Frilasita A Yudhaputri, Muhammad Rezki Rasyak, Willy Agustine, Hidayat Trimarsanto, Lydia V. Panggalo, Iskandar Adnan, Sri Masyeni, Erly Sintya, Ida Ayu Wayan Mahayani, Safarina G Ma |
| EPI_ISL_2101328 | RSUD Mangusada | Eijkman Institute for Molecular Biology, National Agency for Research and Innovation; Molecular Biology Laboratory, Faculty | Lidwina Priliani, Edison Johar, Frilasita A Yudhaputri, Muhammad Rezki Rasyak, Willy Agustine, Hidayat Trimarsanto, Lydia V. Panggalo, Iskandar Adnan, Sukma Oktavianthi, Sri Masyeni, Erly Sintya, Ida Ayu Wayan Mahayani, Safarina G Ma |

| | | | |
|---|---|---|---|
| | | Medicine and Health Sciences, Warmadewa University | |
| EPI_ISL_2101329 | RSUD Mangusada | Eijkman Institute for Molecular Biology, National Agency for Research and Innovation; Molecular Biology Laboratory, Faculty Medicine and Health Sciences, Warmadewa University | Edison Johar, Frilasita A Yudhaputri, Muhammad Rezki Rasyak, Willy Agustine, Hidayat Trimarsanto, Lydia V. Panggalo, Iskandar Adnan, Sukma Oktavianthi, Lidwina Priliani, Sri Masyeni, Erly Sintya, Ida Ayu Wayan Mahayani, Safarina G Ma |
| EPI_ISL_2101330 | RSUD Mangusada | Eijkman Institute for Molecular Biology, National Agency for Research and Innovation; Molecular Biology Laboratory, Faculty Medicine and Health Sciences, Warmadewa University | Frilasita A Yudhaputri, Muhammad Rezki Rasyak, Willy Agustine, Hidayat Trimarsanto, Lydia V. Panggalo, Iskandar Adnan, Sukma Oktavianthi, Lidwina Priliani, Edison Johar, Sri Masyeni, Erly Sintya, Ida Ayu Wayan Mahayani, Safarina G Ma |
| EPI_ISL_2101331 | RSUD Mangusada | Eijkman Institute for Molecular Biology, National Agency for Research and Innovation; Molecular Biology Laboratory, Faculty Medicine and Health Sciences, Warmadewa University | Muhammad Rezki Rasyak, Willy Agustine, Hidayat Trimarsanto, Lydia V. Panggalo, Iskandar Adnan, Sukma Oktavianthi, Lidwina Priliani, Edison Johar, Frilasita A Yudhaputri, Sri Masyeni, Erly Sintya, Ida Ayu Wayan Mahayani, Safarina G Ma |
| EPI_ISL_2101332 | RSUD Mangusada | Eijkman Institute for Molecular Biology, National Agency for Research and Innovation; Molecular Biology Laboratory, Faculty Medicine and Health Sciences, Warmadewa University | Hidayat Trimarsanto, Lydia V. Panggalo, Iskandar Adnan, Sukma Oktavianthi, Lidwina Priliani, Edison Johar, Frilasita A Yudhaputri, Muhammad Rezki Rasyak, Willy Agustine, Sri Masyeni, Erly Sintya, Ida Ayu Wayan Mahayani, Safarina G Ma |
| EPI_ISL_2101333 | RSUD Mangusada | Eijkman Institute for Molecular Biology, National Agency for Research and Innovation; Molecular Biology Laboratory, Faculty Medicine and Health Sciences, Warmadewa University | Lydia V. Panggalo, Iskandar Adnan, Sukma Oktavianthi, Lidwina Priliani, Edison Johar, Frilasita A Yudhaputri, Muhammad Rezki Rasyak, Willy Agustine, Hidayat Trimarsanto, Sri Masyeni, Erly Sintya, Ida Ayu Wayan Mahayani, Safarina G Ma |
| EPI_ISL_2101334 | RSUD Mangusada | Eijkman Institute for Molecular Biology, National Agency for Research and Innovation; Molecular Biology Laboratory, Faculty Medicine and Health Sciences, Warmadewa University | Iskandar Adnan, Sukma Oktavianthi, Lidwina Priliani, Edison Johar, Frilasita A Yudhaputri, Muhammad Rezki Rasyak, Willy Agustine, Hidayat Trimarsanto, Lydia V. Panggalo, Sri Masyeni, Erly Sintya, Ida Ayu Wayan Mahayani, Safarina G Ma |
| EPI_ISL_2101335 | RSUD Mangusada | Eijkman Institute for Molecular Biology, National Agency for Research and Innovation; Molecular Biology Laboratory, Faculty Medicine and Health Sciences, Warmadewa University | Sukma Oktavianthi, Lidwina Priliani, Edison Johar, Frilasita A Yudhaputri, Muhammad Rezki Rasyak, Willy Agustine, Hidayat Trimarsanto, Lydia V. Panggalo, Iskandar Adnan, Sri Masyeni, Erly Sintya, Ida Ayu Wayan Mahayani, Safarina G Ma |
| EPI_ISL_2101336 | RSUD Mangusada | Eijkman Institute for Molecular Biology, National Agency for Research and Innovation; Molecular Biology Laboratory, Faculty Medicine and Health Sciences, Warmadewa University | Edison Johar, Frilasita A Yudhaputri, Muhammad Rezki Rasyak, Willy Agustine, Hidayat Trimarsanto, Lydia V. Panggalo, Iskandar Adnan, Sukma Oktavianthi, Lidwina Priliani, Sri Masyeni, Erly Sintya, Ida Ayu Wayan Mahayani, Safarina G Ma |
| EPI_ISL_2101337 | RSUD Mangusada | Eijkman Institute for Molecular Biology, National Agency for Research and Innovation; Molecular Biology Laboratory, Faculty Medicine and Health Sciences, Warmadewa University | Frilasita A Yudhaputri, Muhammad Rezki Rasyak, Willy Agustine, Hidayat Trimarsanto, Lydia V. Panggalo, Iskandar Adnan, Sukma Oktavianthi, Lidwina Priliani, Edison Johar, Sri Masyeni, Erly Sintya, Ida Ayu Wayan Mahayani, Safarina G Ma |
| EPI_ISL_2101338 | RSUD Mangusada | Eijkman Institute for Molecular Biology, National Agency for Research and Innovation; Molecular Biology Laboratory, Faculty | Muhammad Rezki Rasyak, Willy Agustine, Hidayat Trimarsanto, Lydia V. Panggalo, Iskandar Adnan, Sukma Oktavianthi, Lidwina Priliani, Edison Johar, Frilasita A Yudhaputri, Sri Masyeni, Erly Sintya, Ida Ayu Wayan Mahayani, Safarina G Ma |

| | | | |
|---|---|---|---|
| | | Medicine and Health Sciences, Warmadewa University | |
| EPI_ISL_2101339 | RSUD Mangusada | Eijkman Institute for Molecular Biology, National Agency for Research and Innovation; Molecular Biology Laboratory, Faculty Medicine and Health Sciences, Warmadewa University | Willy Agustine, Hidayat Trimarsanto, Lydia V. Panggalo, Iskandar Adnan, Sukma Oktavianthi, Lidwina Priliani, Edison Johar, Frilasita A Yudhaputri, Muhammad Rezki Rasyak, Sri Masyeni, Erly Sintya, Ida Ayu Wayan Mahayani, Safarina G Ma |
| EPI_ISL_2101340 | RSUD Mangusada | Eijkman Institute for Molecular Biology, National Agency for Research and Innovation; Molecular Biology Laboratory, Faculty Medicine and Health Sciences, Warmadewa University | Hidayat Trimarsanto, Lydia V. Panggalo, Iskandar Adnan, Sukma Oktavianthi, Lidwina Priliani, Edison Johar, Frilasita A Yudhaputri, Muhammad Rezki Rasyak, Willy Agustine, Sri Masyeni, Erly Sintya, Ida Ayu Wayan Mahayani, Safarina G Ma |
| EPI_ISL_2101341 | RSUD Mangusada | Eijkman Institute for Molecular Biology, National Agency for Research and Innovation; Molecular Biology Laboratory, Faculty Medicine and Health Sciences, Warmadewa University | Lydia V. Panggalo, Iskandar Adnan, Sukma Oktavianthi, Lidwina Priliani, Edison Johar, Frilasita A Yudhaputri, Muhammad Rezki Rasyak, Willy Agustine, Hidayat Trimarsanto, Sri Masyeni, Erly Sintya, Ida Ayu Wayan Mahayani, Safarina G Ma |
| EPI_ISL_2101342 | RSUD Mangusada | Eijkman Institute for Molecular Biology, National Agency for Research and Innovation; Molecular Biology Laboratory, Faculty Medicine and Health Sciences, Warmadewa University | Iskandar Adnan, Sukma Oktavianthi, Lidwina Priliani, Edison Johar, Frilasita A Yudhaputri, Muhammad Rezki Rasyak, Willy Agustine, Hidayat Trimarsanto, Lydia V. Panggalo, Sri Masyeni, Erly Sintya, Ida Ayu Wayan Mahayani, Safarina G Ma |
| EPI_ISL_2106975 | BPOM Gorontalo | National Institute of Health Research and Development | Hartanti Dian Ikawati, Hana Apsari Pawestri, Subangkit, Kartika Dewi Puspa, Arie Ardiansyah Nugraha, Triyani Soekarso, Krisna Pangesti, Nelly Puspandari, Vivi Setiawaty |
| EPI_ISL_2106976, EPI_ISL_2106977, EPI_ISL_2106978 | RSUP Sanglah | National Institute of Health Research and Development | Vivi Setiawaty, Hana Apsari Pawestri, Subangkit, Kartika Dewi Puspa, Arie Ardiansyah Nugraha, Hartanti Dian Ikawati, Krisna Pangesti, Triyani Soekarso, Nelly Puspandari, I Nengah Tony Rustawan, I Gede Ketut Sa |
| EPI_ISL_2106979 | RSUD Bali Mandara | National Institute of Health Research and Development | Hana Apsari Pawestri, Kartika Dewi Puspa, Arie Ardiansyah Nugraha, Hartanti Dian Ikawati, Krisna Pangesti, Triyani Soekarso, Nelly Puspandari, Subangkit, I Wayan Agus Gede Manik Saputra, Vivi Setiawa |
| EPI_ISL_2106980 | RS Mandaya Karawang | National Institute of Health Research and Development | Vivi Setiawaty, Hana Apsari Pawestri, Subangkit, Kartika Dewi Puspa, Arie Ardiansyah Nugraha, Hartanti Dian Ikawati, Nelly Puspandari, Krisna Pangesti |
| EPI_ISL_2106981 | Labkesda Karawang | National Institute of Health Research and Development | Hana Apsari Pawestri, Subangkit, Kartika Dewi Puspa, Arie Ardiansyah Nugraha, Hartanti Dian Ikawati, Nelly Puspandari, Krisna Pangesti, Vivi Setiawaty |
| EPI_ISL_2106982 | Ciputra Hospital Citra Garden City | National Institute of Health Research and Development | Vivi Setiawaty, Hana Apsari Pawestri, Subangkit, Kartika Dewi Puspa, Arie Ardiansyah Nugraha, Hartanti Dian Ikawati, Nelly Puspandari, Krisna Pangesti |
| EPI_ISL_2106983 | PKC Kramat Jati | National Institute of Health Research and Development | Hana Apsari Pawestri, Kartika Dewi Puspa, Arie Ardiansyah Nugraha, Hartanti Dian Ikawati, Krisna Pangesti, Triyani Soekarso, Nelly Puspandari, Subangkit, Vivi Setiawaty |
| EPI_ISL_2106984, EPI_ISL_2106985, EPI_ISL_2106986 | RSPI Prof. Dr. Sulianti Saroso | National Institute of Health Research and Development | Vivi Setiawaty, Hana Apsari Pawestri, Subangkit, Kartika Dewi Puspa, Arie Ardiansyah Nugraha, Hartanti Dian Ikawati, Krisna Pangesti, Triyani Soekarso, Nelly Puspandari |
| EPI_ISL_2106987 | RSUP H Adam Malik Medan | National Institute of Health Research and Development | Kartika Dewi Puspa, Hana Apsari Pawestri, Arie Ardiansyah Nugraha, Hartanti Dian Ikawati, Krisna Pangesti, Triyani Soekarso, Nelly Puspandari, Subangkit, Maliana, Ade Rahmaini, Nelly Elfrida Samosir, Rina Yunita, V |
| EPI_ISL_2106988 | National Institute of Health Research and Development | National Institute of Health Research and Development | Subangkit, Hana Apsari Pawestri, Kartika Dewi Puspa, Arie Ardiansyah Nugraha, Hartanti Dian Ikawati, Krisna Nur Andriana Pangesti, Yuni Rukminiati, Ririn Ramadhany, Agustiningsih, Kindi Adam, Holy Arif Wibowo, Triyani Soekarso, Ni Ketut Susilarini, Nurika Har Tati Febriyanti, Natalie Laurencia Kipuw, Fauzul Muna, Irene Lorinda Indalao, Nelly Puspandari, Vivi Setiawaty. |
| EPI_ISL_2226647 | RSUD Dr. Soetomo | Institute of Tropical Disease, Universitas Airlangga | Krisnoadi Rahardjo, Aldise M Nastri, Jezzy R Dewantari, Rima R Prasetya, Acub Zaenal, Joni Wahyuhadi, Neneng D Kurniati, Gatot Soegiarto, Laksmi Wulandari, Resti Yudhawati, Yasuko Mori, Soetjipto, Kazufumi Shimiz |
| EPI_ISL_2226648 | BBTKLPP Surabaya | Institute of Tropical Disease, Universitas Airlangga | Aldise M Nastri, Krisnoadi Rahardjo, Jezzy R Dewantari, Rima R Prasetya, Acub Zaenal, Rosidi Roslan, Zahrotunnisa, Gatot Soegiarto, Laksmi Wulandari, Resti Yudhawati, Yasuko Mori, Soetjipto, Kazufumi Shimizu, M |
| EPI_ISL_2233087 | Hotel Horison | National Institute of Health Research and Development | Hartanti Dian Ikawati, Hana Apsari Pawestri, Subangkit, Kartika Dewi Puspa, Arie Ardiansyah Nugraha, Triyani Soekarso, Krisna Pangesti, Nelly Puspandari, Vivi Setiawaty |
| EPI_ISL_2233088 | BBTKLPP Kelas I Batam | National Institute of Health Research and Development | Vivi Setiawaty, Hartanti Dian Ikawati, Hana Apsari Pawestri, Subangkit, Kartika Dewi Puspa, Arie Ardiansyah Nugraha, Triyani Soekarso, Krisna Pangesti, Budi Santosa, Ismail, Ida Neni Haryanti, Nelly Puspan |
| EPI_ISL_2233089 | RSUD Abdul Moeis | National Institute of Health Research and Development | Vivi Setiawaty, Hartanti Dian Ikawati, Hana Apsari Pawestri, Subangkit, Kartika Dewi Puspa, Arie Ardiansyah Nugraha, Triyani Soekarso, Krisna Pangesti, Nelly Puspandari |
| EPI_ISL_2233090 | Dinkes Samarinda | National Institute of Health Research and Development | Vivi Setiawaty, Hartanti Dian Ikawati, Hana Apsari Pawestri, Subangkit, Kartika Dewi Puspa, Arie Ardiansyah Nugraha, Triyani Soekarso, Krisna Pangesti, Nelly Puspandari |
| EPI_ISL_2233091, EPI_ISL_2233092, EPI_ISL_2233093 | Kantor Kesehatan Pelabuhan Kelas II Cilacap | National Institute of Health Research and Development | Kartika Dewi Puspa, Subangkit, Arie Ardiansyah Nugraha, Hartanti Dian Ikawati, Hana Apsari Pawestri, Triyani Soekarso, Krisna Pangesti, Nelly Puspandari, Vivi Setiawaty |
| EPI_ISL_2233094 | Labkesda Prov Kal | National Institute of Health | Kartika Dewi Puspa, Subangkit, Arie Ardiansyah Nugraha, Hartanti Dian Ikawati, Hana Apsari Pawestri, Triyani Soekarso, Krisna Pangesti, Nelly Puspandari, Vivi Setiawaty |

| | | | |
|---|---|---|---|
| | Tim | Research and Development | |
| EPI_ISL_2233095 | Hotel Horison | National Institute of Health Research and Development | Kartika Dewi Puspa, Subangkit, Arie Ardiansyah Nugraha, Hartanti Dian Ikawati, Hana Apsari Pawestri, Triyani Soekarso, Krisna Pangesti, Nelly Puspandari, Vivi Setiawaty |
| EPI_ISL_2233096, EPI_ISL_2233097, EPI_ISL_2233098, EPI_ISL_2233099, EPI_ISL_2233100, EPI_ISL_2233101, EPI_ISL_2233102, EPI_ISL_2233103, EPI_ISL_2233104, EPI_ISL_2233105 | Kantor Kesehatan Pelabuhan Kelas II Cilacap | National Institute of Health Research and Development | Subangkit, Hana Apsari Pawestri, Hartanti Dian Ikawati, Kartika Dewi Puspa, Arie Ardiansyah Nugraha, Triyani Soekarso, Krisna Pangesti, Nelly Puspandari, Vivi Setiawaty |
| EPI_ISL_2233106, EPI_ISL_2233107, EPI_ISL_2233108, EPI_ISL_2233109 | RSUD Arifin Ahmad Riau | National Institute of Health Research and Development | Hana Apsari Pawestri, Hartanti Dian Ikawati, Subangkit, Kartika Dewi Puspa, Arie Ardiansyah Nugraha, Triyani Soekarso, Krisna Pangesti, Nelly Puspandari, Fajri Marinda, Vivi Setiawaty |
| EPI_ISL_2258207 | RSUD dr Soetomo Surabaya | National Institute of Health Research and Development | Hana Apsari Pawestri, Kartika Dewi Puspa, Arie Ardiansyah Nugraha, Subangkit, Hartanti Dian Ikawati, Triyani Soekarso, Krisna Pangesti, Nelly Puspandari, Vivi Setiawaty |
| EPI_ISL_2258213, EPI_ISL_2258218 | RSUD Arifin Ahmad Riau | National Institute of Health Research and Development | Hana Apsari Pawestri, Hartanti Dian Ikawati, Subangkit, Kartika Dewi Puspa, Arie Ardiansyah Nugraha, Triyani Soekarso, Krisna Pangesti, Nelly Puspandari, Fajri Marinda, Vivi Setiawaty |
| EPI_ISL_2262252 | RSUP Fatmawati | National Institute of Health Research and Development | Vivi Setiawaty, Hana Apsari Pawestri, Hartanti Dian Ikawati, Subangkit, Kartika Dewi Puspa, Arie Ardiansyah Nugraha, Triyani Soekarso, Krisna Pangesti, Nelly Puspandari |
| EPI_ISL_2262254, EPI_ISL_2262255, EPI_ISL_2262257, EPI_ISL_2262260, EPI_ISL_2262261, EPI_ISL_2262262, EPI_ISL_2262263, EPI_ISL_2262265 | RS Umum Kartika Pulomas | National Institute of Health Research and Development | Vivi Setiawaty, Hana Apsari Pawestri, Hartanti Dian Ikawati, Subangkit, Kartika Dewi Puspa, Arie Ardiansyah Nugraha, Triyani Soekarso, Krisna Pangesti, Nelly Puspandari |
| EPI_ISL_2262267 | Dinkes Kepulauan Seribu | National Institute of Health Research and Development | Vivi Setiawaty, Hana Apsari Pawestri, Hartanti Dian Ikawati, Subangkit, Kartika Dewi Puspa, Arie Ardiansyah Nugraha, Triyani Soekarso, Krisna Pangesti, Nelly Puspandari |
| EPI_ISL_2262269, EPI_ISL_2262271, EPI_ISL_2262273 | RSPI Prof. Dr. Sulianti Saroso | National Institute of Health Research and Development | Subangkit, Hana Apsari Pawestri, Hartanti Dian Ikawati, Kartika Dewi Puspa, Arie Ardiansyah Nugraha, Triyani Soekarso, Krisna Pangesti, Nelly Puspandari, Vivi Setiawaty |
| EPI_ISL_2262276, EPI_ISL_2262277 | RSUD dr Soetomo Surabaya | National Institute of Health Research and Development | Subangkit, Hana Apsari Pawestri, Hartanti Dian Ikawati, Kartika Dewi Puspa, Arie Ardiansyah Nugraha, Triyani Soekarso, Krisna Pangesti, Nelly Puspandari, Vivi Setiawaty |
| EPI_ISL_2262280 | Dinkes Kab Kota Waringin Barat | National Institute of Health Research and Development | Subangkit, Hana Apsari Pawestri, Hartanti Dian Ikawati, Kartika Dewi Puspa, Arie Ardiansyah Nugraha, Triyani Soekarso, Krisna Pangesti, Nelly Puspandari, Vivi Setiawaty |
| EPI_ISL_2262281 | Dinkes Kab Kota Waringin Timur | National Institute of Health Research and Development | Subangkit, Hana Apsari Pawestri, Hartanti Dian Ikawati, Kartika Dewi Puspa, Arie Ardiansyah Nugraha, Triyani Soekarso, Krisna Pangesti, Nelly Puspandari, Vivi Setiawaty |
| EPI_ISL_2262283, EPI_ISL_2262285, EPI_ISL_2262288, EPI_ISL_2262289, EPI_ISL_2262291 | RSPI Prof. Dr. Sulianti Saroso | National Institute of Health Research and Development | Hana Apsari Pawestri, Kartika Dewi Puspa, Arie Ardiansyah Nugraha, Subangkit, Hartanti Dian Ikawati, Triyani Soekarso, Krisna Pangesti, Nelly Puspandari, Vivi Setiawaty |
| EPI_ISL_2262294 | RSUD Brebes | National Institute of Health Research and Development | Hana Apsari Pawestri, Kartika Dewi Puspa, Arie Ardiansyah Nugraha, Subangkit, Hartanti Dian Ikawati, Triyani Soekarso, Krisna Pangesti, Nelly Puspandari, Vivi Setiawaty |
| EPI_ISL_2262295 | RSPI Prof. Dr. Sulianti Saroso | National Institute of Health Research and Development | Hana Apsari Pawestri, Kartika Dewi Puspa, Arie Ardiansyah Nugraha, Subangkit, Hartanti Dian Ikawati, Triyani Soekarso, Krisna Pangesti, Nelly Puspandari, Vivi Setiawaty |
| EPI_ISL_2274436 | Anwar Medika General Hospital | Institute of Tropical Disease, Universitas Airlangga | Rima R Prasetya, Krisnoadi Rahardjo, Aldise M Nastri, Jezzy R Dewantari, Nungky Taniasari, Gatot Soegiarto, Laksmi Wulandari, Resti Yudhawati, Yasuko Mori, Soetjipto, Kazufumi Shimizu, Maria I Lusida |
| EPI_ISL_2274437 | RSUD Dr. Soetomo | Institute of Tropical Disease, Universitas Airlangga | Kazufumi Shimizu, Krisnoadi Rahardjo, Aldise M Nastri, Jezzy R Dewantari, Rima R Prasetya, Joni Wahyuhadi, Neneng D Kurniati, Gatot Soegiarto, Laksmi Wulandari, Resti Yudhawati, Yasuko Mori, Soetjipto, Mari |
| EPI_ISL_2274438 | Adi Husada Kapasari Hospital | Institute of Tropical Disease, Universitas Airlangga | Aldise M Nastri, Krisnoadi Rahardjo, Jezzy R Dewantari, Rima R Prasetya, Hermanto Wijaya, Gatot Soegiarto, Laksmi Wulandari, Resti Yudhawati, Yasuko Mori, Soetjipto, Kazufumi Shimizu, Maria I Lusida |
| EPI_ISL_2284868 | Primasatya Husada Citra Hospital | Institute of Tropical Disease, Universitas Airlangga | Jezzy R Dewantari, Rima R Prasetya, Krisnoadi Rahardjo, Aldise M Nastri, Pudji Djanuartono, Gatot Soegiarto, Laksmi Wulandari, Resti Yudhawati, Yasuko Mori, Soetjipto, Kazufumi Shimizu, Maria I Lusida |
| EPI_ISL_2284869 | RSUD Dr. Soetomo | Institute of Tropical Disease, Universitas Airlangga | Rima R Prasetya, Krisnoadi Rahardjo, Aldise M Nastri, Jezzy R Dewantari, Joni Wahyuhadi, Neneng D Kurniati, Gatot Soegiarto, Laksmi Wulandari, Resti Yudhawati, Yasuko Mori, Soetjipto, Kazufumi Shimizu, Mari |
| EPI_ISL_2284870 | RSUD Dr. Soetomo | Institute of Tropical Disease, Universitas Airlangga | Kazufumi Shimizu, Krisnoadi Rahardjo, Aldise M Nastri, Jezzy R Dewantari, Rima R Prasetya, Joni Wahyuhadi, Neneng D Kurniati, Gatot Soegiarto, Laksmi Wulandari, Resti Yudhawati, Yasuko Mori, Soetjipto, Mari |
| EPI_ISL_2284871 | RSUD Dr. Soetomo | Institute of Tropical Disease, Universitas Airlangga | Krisnoadi Rahardjo, Aldise M Nastri, Jezzy R Dewantari, Rima R Prasetya, Joni Wahyuhadi, Neneng D Kurniati, Gatot Soegiarto, Laksmi Wulandari, Resti Yudhawati, Yasuko Mori, Soetjipto, Kazufumi Shimizu, Mari |
| EPI_ISL_2284872 | RSUD Dr. Soetomo | Institute of Tropical Disease, Universitas Airlangga | Aldise M Nastri, Krisnoadi Rahardjo, Jezzy R Dewantari, Rima R Prasetya, Joni Wahyuhadi, Neneng D Kurniati, Gatot Soegiarto, Laksmi Wulandari, Resti Yudhawati, Yasuko Mori, Soetjipto, Kazufumi Shimizu, Mari |

| | | | |
|---|---|---|---|
| EPI_ISL_2284873 | National Hospital | Institute of Tropical Disease, Universitas Airlangga | Jezzy R Dewantari, Rima R Prasetya, Krisnoadi Rahardjo, Aldise M Nastri, Azwan H Lubis, Gatot Soegiarto, Laksmi Wulandari, Resti Yudhawati, Yasuko Mori, Soetjipto, Kazufumi Shimizu, Maria I Lusida |
| EPI_ISL_2284874 | Siti Khodijah Hospital | Institute of Tropical Disease, Universitas Airlangga | Kazufumi Shimizu, Krisnoadi Rahardjo, Aldise M Nastri, Jezzy R Dewantari, Rima R Prasetya, Muhammad Hamdan, Gatot Soegiarto, Laksmi Wulandari, Resti Yudhawati, Yasuko Mori, Soetjipto, Maria I Lusi |
| EPI_ISL_2284875 | Darus Syifa' Islamic Hospital | Institute of Tropical Disease, Universitas Airlangga | Krisnoadi Rahardjo, Aldise M Nastri, Jezzy R Dewantari, Rima R Prasetya, HM Faiz, Gatot Soegiarto, Laksmi Wulandari, Resti Yudhawati, Yasuko Mori, Soetjipto, Kazufumi Shimizu, Maria I Lusida |
| EPI_ISL_2284876 | Institute of Tropical Disease, Universitas Airlangga | Institute of Tropical Disease, Universitas Airlangga | Aldise M Nastri, Krisnoadi Rahardjo, Jezzy R Dewantari, Rima R Prasetya, Gatot Soegiarto, Laksmi Wulandari, Resti Yudhawati, Yasuko Mori, Soetjipto, Kazufumi Shimizu, Maria I Lusida |
| EPI_ISL_2284877 | Graha Medika Maternity Hospital | Institute of Tropical Disease, Universitas Airlangga | Jezzy R Dewantari, Rima R Prasetya, Krisnoadi Rahardjo, Aldise M Nastri, Diah R Kusumawati, Gatot Soegiarto, Laksmi Wulandari, Resti Yudhawati, Yasuko Mori, Soetjipto, Kazufumi Shimizu, Maria I Lusic |
| EPI_ISL_2284878 | Institute of Tropical Disease, Universitas Airlangga | Institute of Tropical Disease, Universitas Airlangga | Rima R Prasetya, Krisnoadi Rahardjo, Aldise M Nastri, Jezzy R Dewantari, Gatot Soegiarto, Laksmi Wulandari, Resti Yudhawati, Yasuko Mori, Soetjipto, Kazufumi Shimizu, Maria I Lusida |
| EPI_ISL_2284879 | Siti Khodijah Hospital | Institute of Tropical Disease, Universitas Airlangga | Kazufumi Shimizu, Krisnoadi Rahardjo, Aldise M Nastri, Jezzy R Dewantari, Rima R Prasetya, Muhammad Hamdan, Gatot Soegiarto, Laksmi Wulandari, Resti Yudhawati, Yasuko Mori, Soetjipto, Maria I Lusi |
| EPI_ISL_2284880 | Institute of Tropical Disease, Universitas Airlangga | Institute of Tropical Disease, Universitas Airlangga | Krisnoadi Rahardjo, Aldise M Nastri, Jezzy R Dewantari, Rima R Prasetya, Gatot Soegiarto, Laksmi Wulandari, Resti Yudhawati, Yasuko Mori, Soetjipto, Kazufumi Shimizu, Maria I Lusida |
| EPI_ISL_2284881 | Graha Medika Maternity Hospital | Institute of Tropical Disease, Universitas Airlangga | Aldise M Nastri, Krisnoadi Rahardjo, Jezzy R Dewantari, Rima R Prasetya, Diah R Kusumawati, Gatot Soegiarto, Laksmi Wulandari, Resti Yudhawati, Yasuko Mori, Soetjipto, Kazufumi Shimizu, Maria I Lusic |
| EPI_ISL_2284882 | Primasatya Husada Citra Hospital | Institute of Tropical Disease, Universitas Airlangga | Jezzy R Dewantari, Rima R Prasetya, Krisnoadi Rahardjo, Aldise M Nastri, Pudji Djanuartono, Gatot Soegiarto, Laksmi Wulandari, Resti Yudhawati, Yasuko Mori, Soetjipto, Kazufumi Shimizu, Maria I Lusida |
| EPI_ISL_2284883 | Siti Khodijah Hospital | Institute of Tropical Disease, Universitas Airlangga | Rima R Prasetya, Krisnoadi Rahardjo, Aldise M Nastri, Jezzy R Dewantari, Muhammad Hamdan, Gatot Soegiarto, Laksmi Wulandari, Resti Yudhawati, Yasuko Mori, Soetjipto, Kazufumi Shimizu, Maria I Lusi |
| EPI_ISL_2284884 | RSUD Dr. Soetomo | Institute of Tropical Disease, Universitas Airlangga | Krisnoadi Rahardjo, Aldise M Nastri, Jezzy R Dewantari, Rima R Prasetya, Joni Wahyuhadi, Neneng D Kurniati, Gatot Soegiarto, Laksmi Wulandari, Resti Yudhawati, Yasuko Mori, Soetjipto, Kazufumi Shimizu, Mari |
| EPI_ISL_2296526 | Genomik Solidaritas Indonesia Laboratorium | Genomik Solidaritas Indonesia Laboratorium | Carissa Sintca Wijaya, Vania Gavrila Wikasa, Normastuti Adhini Tantyo, Gracia Felias Enos Korompis, Alfin Mohammad Abdillah, Dhahlia Agustina Cahyono, Annisa Muthiah Sukirman, Meutia Ayuputeri Kumaheri, Anindya Pradipt |
| EPI_ISL_2333654 | RSUD Cilacap | Genetics Working Group (Pokja Genetik) Faculty of Medicine, Public Health and Nursing Universitas Gadjah Mada (FK-KMK UGM); Disease Investigation Center Wates Ministry of Agriculture Indonesia; Department of Microbiology FK-KMK UGM; Laboratorium Diagnostik Yayasan Tahija World Mosquito Program (WMP) Yogyakarta Center for Tropical Medicine FK-KMK UGM; Integrated Research Center FK-KMK UGM; Department of Computer Science and Electronics FMIPA UGM; RSUP Dr. Sardjito | Gunadi, Hendra Wibawa, Marcellus, Mohamad S. Hakim, Edwin W. Daniwijaya, Ludhang P. Rizki, Endah Supriyati, Eggi Arguni, Titik Nuryastuti, Tri Wibawa, Dwi AA Nugrahaningsih, Afiahayati, Siswanto,Kristy Iskandar,Nungki Anggorowati, Pramesti G Dewi, Dw Puspitarani |
| EPI_ISL_2333810 | RSUD Cilacap | Genetics Working Group (Pokja Genetik) Faculty of Medicine, Public Health and Nursing Universitas Gadjah Mada (FK-KMK UGM); Disease Investigation Center Wates Ministry of Agriculture Indonesia; Department of Microbiology FK-KMK UGM; Laboratorium Diagnostik Yayasan Tahija World Mosquito Program (WMP) Yogyakarta Center for Tropical Medicine FK-KMK UGM; Integrated Research Center FK-KMK UGM; Department of Computer | Gunadi, Hendra Wibawa, Marcellus, Mohamad S. Hakim, Edwin W. Daniwijaya, Ludhang P. Rizki, Endah Supriyati, Eggi Arguni, Titik Nuryastuti, Tri Wibawa, Dwi AA Nugrahaningsih, Afiahayati, Siswanto,Kristy Iskandar,Nungki Anggorowati, Pramesti G Dewi, D Inggriani |

| | | | |
|---|---|---|---|
| | | Science and Electronics FMIPA UGM; RSUP Dr. Sardjito | |
| EPI_ISL_2334058 | RSUD Cilacap | Genetics Working Group (Pokja Genetik) Faculty of Medicine, Public Health and Nursing Universitas Gadjah Mada (FK-KMK UGM); Disease Investigation Center Wates Ministry of Agriculture Indonesia; Department of Microbiology FK-KMK UGM; Laboratorium Diagnostik Yayasan Tahija World Mosquito Program (WMP) Yogyakarta Center for Tropical Medicine FK-KMK UGM; Integrated Research Center FK-KMK UGM; Department of Computer Science and Electronics FMIPA UGM; RSUP Dr. Sardjito | Gunadi, Hendra Wibawa, Marcellus, Mohamad S. Hakim, Edwin W. Daniwijaya, Ludhang P. Rizki, Endah Supriyati, Eggi Arguni, Titik Nuryastuti, Tri Wibawa, Dwi AA Nugrahaningsih, Afiahayati, Siswanto,Kristy Iskandar,Nungki Anggorowati, Pramesti G Dewi, Dw |
| EPI_ISL_2334094 | RSUD Cilacap | Genetics Working Group (Pokja Genetik) Faculty of Medicine, Public Health and Nursing Universitas Gadjah Mada (FK-KMK UGM); Disease Investigation Center Wates Ministry of Agriculture Indonesia; Department of Microbiology FK-KMK UGM; Laboratorium Diagnostik Yayasan Tahija World Mosquito Program (WMP) Yogyakarta Center for Tropical Medicine FK-KMK UGM; Integrated Research Center FK-KMK UGM; Department of Computer Science and Electronics FMIPA UGM; RSUP Dr. Sardjito | Gunadi, Hendra Wibawa, Marcellus, Mohamad S. Hakim, Edwin W. Daniwijaya, Ludhang P. Rizki, Endah Supriyati, Eggi Arguni, Titik Nuryastuti, Tri Wibawa, Dwi AA Nugrahaningsih, Afiahayati, Siswanto,Kristy Iskandar,Nungki Anggorowati, Pramesti G Dewi, D Setiawan |
| EPI_ISL_2334474 | RSUD Cilacap | Genetics Working Group (Pokja Genetik) Faculty of Medicine, Public Health and Nursing Universitas Gadjah Mada (FK-KMK UGM); Disease Investigation Center Wates Ministry of Agriculture Indonesia; Department of Microbiology FK-KMK UGM; Laboratorium Diagnostik Yayasan Tahija World Mosquito Program (WMP) Yogyakarta Center for Tropical Medicine FK-KMK UGM; Integrated Research Center FK-KMK UGM; Department of Computer Science and Electronics FMIPA UGM; RSUP Dr. Sardjito | Gunadi, Hendra Wibawa, Marcellus, Mohamad S. Hakim, Edwin W. Daniwijaya, Ludhang P. Rizki, Endah Supriyati, Eggi Arguni, Titik Nuryastuti, Tri Wibawa, Dwi AA Nugrahaningsih, Afiahayati, Siswanto,Kristy Iskandar,Nungki Anggorowati, Pramesti G Dewi, Dwi I A |
| EPI_ISL_2334587 | RSUD Cilacap | Genetics Working Group (Pokja Genetik) Faculty of Medicine, Public Health and Nursing Universitas Gadjah Mada (FK-KMK UGM); Disease Investigation Center Wates Ministry of Agriculture Indonesia; Department of Microbiology FK-KMK UGM; Laboratorium Diagnostik Yayasan Tahija World Mosquito Program (WMP) Yogyakarta Center for Tropical Medicine FK-KMK UGM; Integrated Research Center FK-KMK UGM; Department of Computer Science and Electronics | Gunadi, Hendra Wibawa, Marcellus, Mohamad S. Hakim, Edwin W. Daniwijaya, Ludhang P. Rizki, Endah Supriyati, Eggi Arguni, Titik Nuryastuti, Tri Wibawa, Dwi AA Nugrahaningsih, Afiahayati, Siswanto,Kristy Iskandar,Nungki Anggorowati, Pramesti G Dewi, Dw Puspitarani |

| | | | |
|---|---|---|---|
| | | FMIPA UGM; RSUP Dr. Sardjito | |
| EPI_ISL_2335021 | RSUD Cilacap | Genetics Working Group (Pokja Genetik) Faculty of Medicine, Public Health and Nursing Universitas Gadjah Mada (FK-KMK UGM); Disease Investigation Center Wates Ministry of Agriculture Indonesia; Department of Microbiology FK-KMK UGM; Laboratorium Diagnostik Yayasan Tahija World Mosquito Program (WMP) Yogyakarta Center for Tropical Medicine FK-KMK UGM; Integrated Research Center FK-KMK UGM; Department of Computer Science and Electronics FMIPA UGM; RSUP Dr. Sardjito | Gunadi, Hendra Wibawa, Marcellus, Mohamad S. Hakim, Edwin W. Daniwijaya, Ludhang P. Rizki, Endah Supriyati, Eggi Arguni, Titik Nuryastuti, Tri Wibawa, Dwi AA Nugrahaningsih, Afiahayati, Siswanto,Kristy Iskandar,Nungki Anggorowati, Pramesti G Dewi, D Inggriani |
| EPI_ISL_2335090 | RSUD CILACAP | Genetics Working Group (Pokja Genetik) Faculty of Medicine, Public Health and Nursing Universitas Gadjah Mada (FK-KMK UGM); Disease Investigation Center Wates Ministry of Agriculture Indonesia; Department of Microbiology FK-KMK UGM; Laboratorium Diagnostik Yayasan Tahija World Mosquito Program (WMP) Yogyakarta Center for Tropical Medicine FK-KMK UGM; Integrated Research Center FK-KMK UGM; Department of Computer Science and Electronics FMIPA UGM; RSUP Dr. Sardjito | Gunadi, Hendra Wibawa, Marcellus, Mohamad S. Hakim, Edwin W. Daniwijaya, Ludhang P. Rizki, Endah Supriyati, Eggi Arguni, Titik Nuryastuti, Tri Wibawa, Dwi AA Nugrahaningsih, Afiahayati, Siswanto,Kristy Iskandar,Nungki Anggorowati, Pramesti G Dewi, Dw |
| EPI_ISL_2335146 | RSUD CILACAP | Genetics Working Group (Pokja Genetik) Faculty of Medicine, Public Health and Nursing Universitas Gadjah Mada (FK-KMK UGM); Disease Investigation Center Wates Ministry of Agriculture Indonesia; Department of Microbiology FK-KMK UGM; Laboratorium Diagnostik Yayasan Tahija World Mosquito Program (WMP) Yogyakarta Center for Tropical Medicine FK-KMK UGM; Integrated Research Center FK-KMK UGM; Department of Computer Science and Electronics FMIPA UGM; RSUP Dr. Sardjito | Gunadi, Hendra Wibawa, Marcellus, Mohamad S. Hakim, Edwin W. Daniwijaya, Ludhang P. Rizki, Endah Supriyati, Eggi Arguni, Titik Nuryastuti, Tri Wibawa, Dwi AA Nugrahaningsih, Afiahayati, Siswanto,Kristy Iskandar,Nungki Anggorowati, Pramesti G Dewi, D Setiawan |
| EPI_ISL_2335456 | RSUD CILACAP | Genetics Working Group (Pokja Genetik) Faculty of Medicine, Public Health and Nursing Universitas Gadjah Mada (FK-KMK UGM); Disease Investigation Center Wates Ministry of Agriculture Indonesia; Department of Microbiology FK-KMK UGM; Laboratorium Diagnostik Yayasan Tahija World Mosquito Program (WMP) Yogyakarta Center for Tropical Medicine FK-KMK UGM; Integrated Research Center FK-KMK UGM; Department of Computer | Gunadi, Hendra Wibawa, Marcellus, Mohamad S. Hakim, Edwin W. Daniwijaya, Ludhang P. Rizki, Endah Supriyati, Eggi Arguni, Titik Nuryastuti, Tri Wibawa, Dwi AA Nugrahaningsih, Afiahayati, Siswanto,Kristy Iskandar,Nungki Anggorowati, Pramesti G Dewi, Dwi I A |

| | | | |
|---|---|---|---|
| | | Science and Electronics FMIPA UGM; RSUP Dr. Sardjito | |
| EPI_ISL_2335872 | RSUD CILACAP | Genetics Working Group (Pokja Genetik) Faculty of Medicine, Public Health and Nursing Universitas Gadjah Mada (FK-KMK UGM); Disease Investigation Center Wates Ministry of Agriculture Indonesia; Department of Microbiology FK-KMK UGM; Laboratorium Diagnostik Yayasan Tahija World Mosquito Program (WMP) Yogyakarta Center for Tropical Medicine FK-KMK UGM; Integrated Research Center FK-KMK UGM; Department of Computer Science and Electronics FMIPA UGM; RSUP Dr. Sardjito | Gunadi, Hendra Wibawa, Marcellus, Mohamad S. Hakim, Edwin W. Daniwijaya, Ludhang P. Rizki, Endah Supriyati, Eggi Arguni, Titik Nuryastuti, Tri Wibawa, Dwi AA Nugrahaningsih, Afiahayati, Siswanto,Kristy Iskandar,Nungki Anggorowati, Pramesti G Dewi, Dw Puspitarani |
| EPI_ISL_2336149 | RSUD CILACAP | Genetics Working Group (Pokja Genetik) Faculty of Medicine, Public Health and Nursing Universitas Gadjah Mada (FK-KMK UGM); Disease Investigation Center Wates Ministry of Agriculture Indonesia; Department of Microbiology FK-KMK UGM; Laboratorium Diagnostik Yayasan Tahija World Mosquito Program (WMP) Yogyakarta Center for Tropical Medicine FK-KMK UGM; Integrated Research Center FK-KMK UGM; Department of Computer Science and Electronics FMIPA UGM; RSUP Dr. Sardjito | Gunadi, Hendra Wibawa, Marcellus, Mohamad S. Hakim, Edwin W. Daniwijaya, Ludhang P. Rizki, Endah Supriyati, Eggi Arguni, Titik Nuryastuti, Tri Wibawa, Dwi AA Nugrahaningsih, Afiahayati, Siswanto,Kristy Iskandar,Nungki Anggorowati, Pramesti G Dewi, D Inggriani |
| EPI_ISL_2337254 | RS Dr Oen Solo Baru | Genetics Working Group (Pokja Genetik) Faculty of Medicine, Public Health and Nursing Universitas Gadjah Mada (FK-KMK UGM); Disease Investigation Center Wates Ministry of Agriculture Indonesia; Department of Microbiology FK-KMK UGM; Laboratorium Diagnostik Yayasan Tahija World Mosquito Program (WMP) Yogyakarta Center for Tropical Medicine FK-KMK UGM; Integrated Research Center FK-KMK UGM; Department of Computer Science and Electronics FMIPA UGM; RSUP Dr. Sardjito | Gunadi, Hendra Wibawa, Marcellus, Mohamad S. Hakim, Edwin W. Daniwijaya, Ludhang P. Rizki, Endah Supriyati, Eggi Arguni, Titik Nuryastuti, Tri Wibawa, Dwi AA Nugrahaningsih, Afiahayati, Siswanto,Kristy Iskandar,Nungki Anggorov |
| EPI_ISL_2337255 | PUSKESMAS JEPARA | Genetics Working Group (Pokja Genetik) Faculty of Medicine, Public Health and Nursing Universitas Gadjah Mada (FK-KMK UGM); Disease Investigation Center Wates Ministry of Agriculture Indonesia; Department of Microbiology FK-KMK UGM; Laboratorium Diagnostik Yayasan Tahija World Mosquito Program (WMP) Yogyakarta Center for Tropical Medicine FK-KMK UGM; Integrated Research Center FK-KMK UGM; Department of Computer Science and Electronics | Gunadi, Hendra Wibawa, Marcellus, Mohamad S. Hakim, Edwin W. Daniwijaya, Ludhang P. Rizki, Endah Supriyati, Eggi Arguni, Titik Nuryastuti, Tri Wibawa, Dwi AA Nugrahaningsih, Afiahayati, Siswanto,Kristy Iskandar,Nungki Anggorowat |

| | | | |
|---|---|---|---|
| | | FMIPA UGM; RSUP Dr. Sardjito | |
| EPI_ISL_2338534 | RSUP DR SARDJITO | Genetics Working Group (Pokja Genetik) Faculty of Medicine, Public Health and Nursing Universitas Gadjah Mada (FK-KMK UGM); Disease Investigation Center Wates Ministry of Agriculture Indonesia; Department of Microbiology FK-KMK UGM; Laboratorium Diagnostik Yayasan Tahija World Mosquito Program (WMP) Yogyakarta Center for Tropical Medicine FK-KMK UGM; Integrated Research Center FK-KMK UGM; Department of Computer Science and Electronics FMIPA UGM; RSUP Dr. Sardjito | Gunadi, Hendra Wibawa, . Marcellus, Mohamad S. Hakim, Edwin W. Daniwijaya, Ludhang P. Rizki, Endah Supriyati, Eggi Arguni, Titik Nuryastuti, Tri Wibawa, Dwi AA Nugrahaningsih, Afiahayati, Siswanto, Kristy Iskandar, Nungki Anggorowati, Ika Trisnawati, Suma Munawar Gani, Satria Maulana, Nur Rahmi Ananda, Riat El Khair, Yunika Puspadewi, Osman Sianipar, Umi Solekhah Intansari, Elizabeth Henny Herningtiyas, Ira Puspitawati, Nur Imma Fatimah Harahap, Dyah A Puspitara |
| EPI_ISL_2364568 | RS Medirossa Cikarang | Eijkman Institute for Molecular Biology, National Agency for Research and Innovation | Willy Agustine, Hidayat Trimarsanto, Lydia V. Panggalo, Iskandar Adnan, Sukma Oktavianthi, Lidwina Priliani, Edison Johar, Frilasita A Yudhaputri, Muhammad Rezki Rasyak, Safarina G Malik, Khin Saw Myint, Amin |
| EPI_ISL_2364569, EPI_ISL_2364570, EPI_ISL_2364571 | RS Sentra Medika Cikarang | Eijkman Institute for Molecular Biology, National Agency for Research and Innovation | Lydia V. Panggalo, Iskandar Adnan, Sukma Oktavianthi, Lidwina Priliani, Edison Johar, Frilasita A Yudhaputri, Muhammad Rezki Rasyak, Willy Agustine, Hidayat Trimarsanto, Safarina G Malik, Khin Saw Myint, Amin |
| EPI_ISL_2364572 | RS Sentra Medika Cikarang | Eijkman Institute for Molecular Biology, National Agency for Research and Innovation | Iskandar Adnan, Sukma Oktavianthi, Lidwina Priliani, Edison Johar, Frilasita A Yudhaputri, Muhammad Rezki Rasyak, Willy Agustine, Hidayat Trimarsanto, Lydia V. Panggalo, Safarina G Malik, Khin Saw Myint, Amin |
| EPI_ISL_2364573 | RS Bhakti Husada Cikarang | Eijkman Institute for Molecular Biology, National Agency for Research and Innovation | Iskandar Adnan, Sukma Oktavianthi, Lidwina Priliani, Edison Johar, Frilasita A Yudhaputri, Muhammad Rezki Rasyak, Willy Agustine, Hidayat Trimarsanto, Lydia V. Panggalo, Safarina G Malik, Khin Saw Myint, Amin |
| EPI_ISL_2364574 | RS Bhakti Husada Cikarang | Eijkman Institute for Molecular Biology, National Agency for Research and Innovation | Sukma Oktavianthi, Lidwina Priliani, Edison Johar, Frilasita A Yudhaputri, Muhammad Rezki Rasyak, Willy Agustine, Hidayat Trimarsanto, Lydia V. Panggalo, Iskandar Adnan, Safarina G Malik, Khin Saw Myint, Amin |
| EPI_ISL_2364575 | RS Masmitra | Eijkman Institute for Molecular Biology, National Agency for Research and Innovation | Iskandar Adnan, Sukma Oktavianthi, Lidwina Priliani, Edison Johar, Frilasita A Yudhaputri, Muhammad Rezki Rasyak, Willy Agustine, Hidayat Trimarsanto, Lydia V. Panggalo, Safarina G Malik, Khin Saw Myint, Amin |
| EPI_ISL_2364576 | RS Harapan Bunda | Eijkman Institute for Molecular Biology, National Agency for Research and Innovation | Hidayat Trimarsanto, Lydia V. Panggalo, Iskandar Adnan, Sukma Oktavianthi, Lidwina Priliani, Edison Johar, Frilasita A Yudhaputri, Muhammad Rezki Rasyak, Willy Agustine, Safarina G Malik, Khin Saw Myint, Amin |
| EPI_ISL_2364577 | RS Permata Bekasi | Eijkman Institute for Molecular Biology, National Agency for Research and Innovation | Sukma Oktavianthi, Lidwina Priliani, Edison Johar, Frilasita A Yudhaputri, Muhammad Rezki Rasyak, Willy Agustine, Hidayat Trimarsanto, Lydia V. Panggalo, Iskandar Adnan, Safarina G Malik, Khin Saw Myint, Amin |
| EPI_ISL_2364578 | RS Asy-Syifaa Leuwiliang | Eijkman Institute for Molecular Biology, National Agency for Research and Innovation | Sukma Oktavianthi, Lidwina Priliani, Edison Johar, Frilasita A Yudhaputri, Muhammad Rezki Rasyak, Willy Agustine, Hidayat Trimarsanto, Lydia V. Panggalo, Iskandar Adnan, Safarina G Malik, Khin Saw Myint, Amin |
| EPI_ISL_2364579 | RS Mitra Keluarga Cibubur | Eijkman Institute for Molecular Biology, National Agency for Research and Innovation | Lidwina Priliani, Edison Johar, Frilasita A Yudhaputri, Muhammad Rezki Rasyak, Willy Agustine, Hidayat Trimarsanto, Lydia V. Panggalo, Iskandar Adnan, Sukma Oktavianthi, Safarina G Malik, Khin Saw Myint, Amin |
| EPI_ISL_2364580 | RSUD Cileungsi | Eijkman Institute for Molecular Biology, National Agency for Research and Innovation | Edison Johar, Frilasita A Yudhaputri, Muhammad Rezki Rasyak, Willy Agustine, Hidayat Trimarsanto, Lydia V. Panggalo, Iskandar Adnan, Sukma Oktavianthi, Lidwina Priliani, Safarina G Malik, Khin Saw Myint, Amin |
| EPI_ISL_2364581 | RS Sentra Medika Cikarang | Eijkman Institute for Molecular Biology, National Agency for Research and Innovation | Iskandar Adnan, Sukma Oktavianthi, Lidwina Priliani, Edison Johar, Frilasita A Yudhaputri, Muhammad Rezki Rasyak, Willy Agustine, Hidayat Trimarsanto, Lydia V. Panggalo, Safarina G Malik, Khin Saw Myint, Amin |
| EPI_ISL_2364582 | RS Sentra Medika Cikarang | Eijkman Institute for Molecular Biology, National Agency for Research and Innovation | Frilasita A Yudhaputri, Muhammad Rezki Rasyak, Willy Agustine, Hidayat Trimarsanto, Lydia V. Panggalo, Iskandar Adnan, Sukma Oktavianthi, Lidwina Priliani, Edison Johar, Safarina G Malik, Khin Saw Myint, Amin |
| EPI_ISL_2364583 | RSJ Dr. Soeharto Heerdjan | Eijkman Institute for Molecular Biology, National Agency for Research and Innovation | Muhammad Rezki Rasyak, Willy Agustine, Hidayat Trimarsanto, Lydia V. Panggalo, Iskandar Adnan, Sukma Oktavianthi, Lidwina Priliani, Edison Johar, Frilasita A Yudhaputri, Safarina G Malik, Khin Saw Myint, Amin |

| | | | |
|---|---|---|---|
| EPI_ISL_2364584 | Ciputra Hospital Citra Raya Tangerang | Eijkman Institute for Molecular Biology, National Agency for Research and Innovation | Edison Johar, Frilasita A Yudhaputri, Muhammad Rezki Rasyak, Willy Agustine, Hidayat Trimarsanto, Lydia V. Panggalo, Iskandar Adnan, Sukma Oktavianthi, Lidwina Priliani, Safarina G Malik, Khin Saw Myint, Amin |
| EPI_ISL_2364585 | RSUD Cileungsi | Eijkman Institute for Molecular Biology, National Agency for Research and Innovation | Frilasita A Yudhaputri, Muhammad Rezki Rasyak, Willy Agustine, Hidayat Trimarsanto, Lydia V. Panggalo, Iskandar Adnan, Sukma Oktavianthi, Lidwina Priliani, Edison Johar, Safarina G Malik, Khin Saw Myint, Amin |
| EPI_ISL_2364586 | RS Graha Juanda | Eijkman Institute for Molecular Biology, National Agency for Research and Innovation | Muhammad Rezki Rasyak, Willy Agustine, Hidayat Trimarsanto, Lydia V. Panggalo, Iskandar Adnan, Sukma Oktavianthi, Lidwina Priliani, Edison Johar, Frilasita A Yudhaputri, Safarina G Malik, Khin Saw Myint, Amin |
| EPI_ISL_2364587 | RS Mitra Keluarga Gading Serpong | Eijkman Institute for Molecular Biology, National Agency for Research and Innovation | Willy Agustine, Hidayat Trimarsanto, Lydia V. Panggalo, Iskandar Adnan, Sukma Oktavianthi, Lidwina Priliani, Edison Johar, Frilasita A Yudhaputri, Muhammad Rezki Rasyak, Safarina G Malik, Khin Saw Myint, Amin |
| EPI_ISL_2364588 | RS Mitra Keluarga Gading Serpong | Eijkman Institute for Molecular Biology, National Agency for Research and Innovation | Hidayat Trimarsanto, Lydia V. Panggalo, Iskandar Adnan, Sukma Oktavianthi, Lidwina Priliani, Edison Johar, Frilasita A Yudhaputri, Muhammad Rezki Rasyak, Willy Agustine, Safarina G Malik, Khin Saw Myint, Amin |
| EPI_ISL_2364589 | RS Medika Dramaga | Eijkman Institute for Molecular Biology, National Agency for Research and Innovation | Lydia V. Panggalo, Iskandar Adnan, Sukma Oktavianthi, Lidwina Priliani, Edison Johar, Frilasita A Yudhaputri, Muhammad Rezki Rasyak, Willy Agustine, Hidayat Trimarsanto, Safarina G Malik, Khin Saw Myint, Amin |
| EPI_ISL_2364590 | RS Bhakti Husada Cikarang | Eijkman Institute for Molecular Biology, National Agency for Research and Innovation | Lidwina Priliani, Edison Johar, Frilasita A Yudhaputri, Muhammad Rezki Rasyak, Willy Agustine, Hidayat Trimarsanto, Lydia V. Panggalo, Iskandar Adnan, Sukma Oktavianthi, Safarina G Malik, Khin Saw Myint, Amin |
| EPI_ISL_2364591 | RS Graha Juanda | Eijkman Institute for Molecular Biology, National Agency for Research and Innovation | Edison Johar, Frilasita A Yudhaputri, Muhammad Rezki Rasyak, Willy Agustine, Hidayat Trimarsanto, Lydia V. Panggalo, Iskandar Adnan, Sukma Oktavianthi, Lidwina Priliani, Safarina G Malik, Khin Saw Myint, Amin |
| EPI_ISL_2364592 | RS Mitra Keluarga Gading Serpong | Eijkman Institute for Molecular Biology, National Agency for Research and Innovation | Frilasita A Yudhaputri, Muhammad Rezki Rasyak, Willy Agustine, Hidayat Trimarsanto, Lydia V. Panggalo, Iskandar Adnan, Sukma Oktavianthi, Lidwina Priliani, Edison Johar, Safarina G Malik, Khin Saw Myint, Amin |
| EPI_ISL_2364593 | RS An-Nisa Tangerang | Eijkman Institute for Molecular Biology, National Agency for Research and Innovation | Frilasita A Yudhaputri, Muhammad Rezki Rasyak, Willy Agustine, Hidayat Trimarsanto, Lydia V. Panggalo, Iskandar Adnan, Sukma Oktavianthi, Lidwina Priliani, Edison Johar, Safarina G Malik, Khin Saw Myint, Amin |
| EPI_ISL_2364594 | RSUD Cileungsi | Eijkman Institute for Molecular Biology, National Agency for Research and Innovation | Muhammad Rezki Rasyak, Willy Agustine, Hidayat Trimarsanto, Lydia V. Panggalo, Iskandar Adnan, Sukma Oktavianthi, Lidwina Priliani, Edison Johar, Frilasita A Yudhaputri, Safarina G Malik, Khin Saw Myint, Amin |
| EPI_ISL_2364595 | RS Bhakti Mulia | Eijkman Institute for Molecular Biology, National Agency for Research and Innovation | Willy Agustine, Hidayat Trimarsanto, Lydia V. Panggalo, Iskandar Adnan, Sukma Oktavianthi, Lidwina Priliani, Edison Johar, Frilasita A Yudhaputri, Muhammad Rezki Rasyak, Safarina G Malik, Khin Saw Myint, Amin |
| EPI_ISL_2364596 | RS Mitra Keluarga Kelapa Gading | Eijkman Institute for Molecular Biology, National Agency for Research and Innovation | Hidayat Trimarsanto, Lydia V. Panggalo, Iskandar Adnan, Sukma Oktavianthi, Lidwina Priliani, Edison Johar, Frilasita A Yudhaputri, Muhammad Rezki Rasyak, Willy Agustine, Safarina G Malik, Khin Saw Myint, Amin |
| EPI_ISL_2364597 | RS Permata Cibubur | Eijkman Institute for Molecular Biology, National Agency for Research and Innovation | Edison Johar, Frilasita A Yudhaputri, Muhammad Rezki Rasyak, Willy Agustine, Hidayat Trimarsanto, Lydia V. Panggalo, Iskandar Adnan, Sukma Oktavianthi, Lidwina Priliani, Safarina G Malik, Khin Saw Myint, Amin |
| EPI_ISL_2364598 | RSIA Bunda Sejahtera | Eijkman Institute for Molecular Biology, National Agency for Research and Innovation | Muhammad Rezki Rasyak, Willy Agustine, Hidayat Trimarsanto, Lydia V. Panggalo, Iskandar Adnan, Sukma Oktavianthi, Lidwina Priliani, Edison Johar, Frilasita A Yudhaputri, Safarina G Malik, Khin Saw Myint, Amin |
| EPI_ISL_2364599 | RS Sari Asih Ciledug | Eijkman Institute for Molecular Biology, National Agency for Research and Innovation | Lydia V. Panggalo, Iskandar Adnan, Sukma Oktavianthi, Lidwina Priliani, Edison Johar, Frilasita A Yudhaputri, Muhammad Rezki Rasyak, Willy Agustine, Hidayat Trimarsanto, Safarina G Malik, Khin Saw Myint, Amin |
| EPI_ISL_2364600 | RS Mitra Keluarga Cibubur | Eijkman Institute for Molecular Biology, National Agency for Research and Innovation | Willy Agustine, Hidayat Trimarsanto, Lydia V. Panggalo, Iskandar Adnan, Sukma Oktavianthi, Lidwina Priliani, Edison Johar, Frilasita A Yudhaputri, Muhammad Rezki Rasyak, Safarina G Malik, Khin Saw Myint, Amin |
| EPI_ISL_2364601 | RS Mary Cileungsi Hijau | Eijkman Institute for Molecular Biology, National Agency for Research and Innovation | Hidayat Trimarsanto, Lydia V. Panggalo, Iskandar Adnan, Sukma Oktavianthi, Lidwina Priliani, Edison Johar, Frilasita A Yudhaputri, Muhammad Rezki Rasyak, Willy Agustine, Safarina G Malik, Khin Saw Myint, Amin |
| EPI_ISL_2364602 | RS Sumber Waras | Eijkman Institute for Molecular Biology, National Agency for Research and Innovation | Iskandar Adnan, Sukma Oktavianthi, Lidwina Priliani, Edison Johar, Frilasita A Yudhaputri, Muhammad Rezki Rasyak, Willy Agustine, Hidayat Trimarsanto, Lydia V. Panggalo, Safarina G Malik, Khin Saw Myint, Amin |
| EPI_ISL_2364603 | RS Mitra Keluarga Gading Serpong | Eijkman Institute for Molecular Biology, National | Sukma Oktavianthi, Lidwina Priliani, Edison Johar, Frilasita A Yudhaputri, Muhammad Rezki Rasyak, Willy Agustine, Hidayat Trimarsanto, Lydia V. Panggalo, Iskandar Adnan, Safarina G Malik, Khin Saw Myint, Amin |

| | | | |
|---|---|---|---|
| | | Agency for Research and Innovation | |
| EPI_ISL_2364604 | RS Bhakti Husada Cikarang | Eijkman Institute for Molecular Biology, National Agency for Research and Innovation | Lidwina Priliani, Edison Johar, Frilasita A Yudhaputri, Muhammad Rezki Rasyak, Willy Agustine, Hidayat Trimarsanto, Lydia V. Panggalo, Iskandar Adnan, Sukma Oktavianthi, Safarina G Malik, Khin Saw Myint, Amin |
| EPI_ISL_2364605 | RS Islam Sari Asih Ar-Rahmah | Eijkman Institute for Molecular Biology, National Agency for Research and Innovation | Willy Agustine, Hidayat Trimarsanto, Lydia V. Panggalo, Iskandar Adnan, Sukma Oktavianthi, Lidwina Priliani, Edison Johar, Frilasita A Yudhaputri, Muhammad Rezki Rasyak, Safarina G Malik, Khin Saw Myint, Amin |
| EPI_ISL_2364606 | RS Medika Dramaga | Eijkman Institute for Molecular Biology, National Agency for Research and Innovation | Edison Johar, Frilasita A Yudhaputri, Muhammad Rezki Rasyak, Willy Agustine, Hidayat Trimarsanto, Lydia V. Panggalo, Iskandar Adnan, Sukma Oktavianthi, Lidwina Priliani, Safarina G Malik, Khin Saw Myint, Amin |
| EPI_ISL_2364607 | RS Medika Dramaga | Eijkman Institute for Molecular Biology, National Agency for Research and Innovation | Lydia V. Panggalo, Iskandar Adnan, Sukma Oktavianthi, Lidwina Priliani, Edison Johar, Frilasita A Yudhaputri, Muhammad Rezki Rasyak, Willy Agustine, Hidayat Trimarsanto, Safarina G Malik, Khin Saw Myint, Amin |
| EPI_ISL_2364608 | RS Harapan Bunda | Eijkman Institute for Molecular Biology, National Agency for Research and Innovation | Iskandar Adnan, Sukma Oktavianthi, Lidwina Priliani, Edison Johar, Frilasita A Yudhaputri, Muhammad Rezki Rasyak, Willy Agustine, Hidayat Trimarsanto, Lydia V. Panggalo, Safarina G Malik, Khin Saw Myint, Amin |
| EPI_ISL_2364609 | RS Harapan Bunda | Eijkman Institute for Molecular Biology, National Agency for Research and Innovation | Frilasita A Yudhaputri, Muhammad Rezki Rasyak, Willy Agustine, Hidayat Trimarsanto, Lydia V. Panggalo, Iskandar Adnan, Sukma Oktavianthi, Lidwina Priliani, Edison Johar, Safarina G Malik, Khin Saw Myint, Amin |
| EPI_ISL_2364610 | RS Mitra Keluarga Kalideres | Eijkman Institute for Molecular Biology, National Agency for Research and Innovation | Muhammad Rezki Rasyak, Willy Agustine, Hidayat Trimarsanto, Lydia V. Panggalo, Iskandar Adnan, Sukma Oktavianthi, Lidwina Priliani, Edison Johar, Frilasita A Yudhaputri, Safarina G Malik, Khin Saw Myint, Amin |
| EPI_ISL_2364611 | RSUD Cileungsi | Eijkman Institute for Molecular Biology, National Agency for Research and Innovation | Hidayat Trimarsanto, Lydia V. Panggalo, Iskandar Adnan, Sukma Oktavianthi, Lidwina Priliani, Edison Johar, Frilasita A Yudhaputri, Muhammad Rezki Rasyak, Willy Agustine, Safarina G Malik, Khin Saw Myint, Amin |
| EPI_ISL_2364612 | RSUD Cileungsi | Eijkman Institute for Molecular Biology, National Agency for Research and Innovation | Sukma Oktavianthi, Lidwina Priliani, Edison Johar, Frilasita A Yudhaputri, Muhammad Rezki Rasyak, Willy Agustine, Hidayat Trimarsanto, Lydia V. Panggalo, Iskandar Adnan, Safarina G Malik, Khin Saw Myint, Amin |
| EPI_ISL_2364613 | RSUD Cileungsi | Eijkman Institute for Molecular Biology, National Agency for Research and Innovation | Frilasita A Yudhaputri, Muhammad Rezki Rasyak, Willy Agustine, Hidayat Trimarsanto, Lydia V. Panggalo, Iskandar Adnan, Sukma Oktavianthi, Lidwina Priliani, Edison Johar, Safarina G Malik, Khin Saw Myint, Amin |
| EPI_ISL_2364614 | RS Mary Cileungsi Hijau | Eijkman Institute for Molecular Biology, National Agency for Research and Innovation | Lydia V. Panggalo, Iskandar Adnan, Sukma Oktavianthi, Lidwina Priliani, Edison Johar, Frilasita A Yudhaputri, Muhammad Rezki Rasyak, Willy Agustine, Hidayat Trimarsanto, Safarina G Malik, Khin Saw Myint, Amin |
| EPI_ISL_2364615 | RS Sentra Medika Cibinong | Eijkman Institute for Molecular Biology, National Agency for Research and Innovation | Lidwina Priliani, Edison Johar, Frilasita A Yudhaputri, Muhammad Rezki Rasyak, Willy Agustine, Hidayat Trimarsanto, Lydia V. Panggalo, Iskandar Adnan, Sukma Oktavianthi, Safarina G Malik, Khin Saw Myint, Amin |
| EPI_ISL_2364616 | RS Mitra Keluarga Gading Serpong | Eijkman Institute for Molecular Biology, National Agency for Research and Innovation | Muhammad Rezki Rasyak, Willy Agustine, Hidayat Trimarsanto, Lydia V. Panggalo, Iskandar Adnan, Sukma Oktavianthi, Lidwina Priliani, Edison Johar, Frilasita A Yudhaputri, Safarina G Malik, Khin Saw Myint, Amin |
| EPI_ISL_2364617 | RS Mitra Keluarga Gading Serpong | Eijkman Institute for Molecular Biology, National Agency for Research and Innovation | Hidayat Trimarsanto, Lydia V. Panggalo, Iskandar Adnan, Sukma Oktavianthi, Lidwina Priliani, Edison Johar, Frilasita A Yudhaputri, Muhammad Rezki Rasyak, Willy Agustine, Safarina G Malik, Khin Saw Myint, Amin |
| EPI_ISL_2364618 | RS Mitra Keluarga Kelapa Gading | Eijkman Institute for Molecular Biology, National Agency for Research and Innovation | Lydia V. Panggalo, Iskandar Adnan, Sukma Oktavianthi, Lidwina Priliani, Edison Johar, Frilasita A Yudhaputri, Muhammad Rezki Rasyak, Willy Agustine, Hidayat Trimarsanto, Safarina G Malik, Khin Saw Myint, Amin |
| EPI_ISL_2364619 | RS Mitra Keluarga Kelapa Gading | Eijkman Institute for Molecular Biology, National Agency for Research and Innovation | Iskandar Adnan, Sukma Oktavianthi, Lidwina Priliani, Edison Johar, Frilasita A Yudhaputri, Muhammad Rezki Rasyak, Willy Agustine, Hidayat Trimarsanto, Lydia V. Panggalo, Safarina G Malik, Khin Saw Myint, Amin |
| EPI_ISL_2364621 | RS Sentra Medika Cikarang | Eijkman Institute for Molecular Biology, National Agency for Research and Innovation | Sukma Oktavianthi, Lidwina Priliani, Edison Johar, Frilasita A Yudhaputri, Muhammad Rezki Rasyak, Willy Agustine, Hidayat Trimarsanto, Lydia V. Panggalo, Iskandar Adnan, Safarina G Malik, Khin Saw Myint, Amin |
| EPI_ISL_2364624 | RS Sentra Medika Cikarang | Eijkman Institute for Molecular Biology, National Agency for Research and Innovation | Lidwina Priliani, Edison Johar, Frilasita A Yudhaputri, Muhammad Rezki Rasyak, Willy Agustine, Hidayat Trimarsanto, Lydia V. Panggalo, Iskandar Adnan, Sukma Oktavianthi, Safarina G Malik, Khin Saw Myint, Amin |
| EPI_ISL_2364626 | RS Sentra Medika Cikarang | Eijkman Institute for Molecular Biology, National Agency for Research and Innovation | Edison Johar, Frilasita A Yudhaputri, Muhammad Rezki Rasyak, Willy Agustine, Hidayat Trimarsanto, Lydia V. Panggalo, Iskandar Adnan, Sukma Oktavianthi, Lidwina Priliani, Safarina G Malik, Khin Saw Myint, Amin |

| | | | |
|---|---|---|---|
| EPI_ISL_2364629 | RS Bhakti Husada Cikarang | Eijkman Institute for Molecular Biology, National Agency for Research and Innovation | Frilasita A Yudhaputri, Muhammad Rezki Rasyak, Willy Agustine, Hidayat Trimarsanto, Lydia V. Panggalo, Iskandar Adnan, Sukma Oktavianthi, Lidwina Priliani, Edison Johar, Safarina G Malik, Khin Saw Myint, Amin |
| EPI_ISL_2364632 | RS Mitra Keluarga Bekasi Barat | Eijkman Institute for Molecular Biology, National Agency for Research and Innovation | Edison Johar, Frilasita A Yudhaputri, Muhammad Rezki Rasyak, Willy Agustine, Hidayat Trimarsanto, Lydia V. Panggalo, Iskandar Adnan, Sukma Oktavianthi, Lidwina Priliani, Safarina G Malik, Khin Saw Myint, Amin |
| EPI_ISL_2364635 | RS Harapan Bunda | Eijkman Institute for Molecular Biology, National Agency for Research and Innovation | Iskandar Adnan, Sukma Oktavianthi, Lidwina Priliani, Edison Johar, Frilasita A Yudhaputri, Muhammad Rezki Rasyak, Willy Agustine, Hidayat Trimarsanto, Lydia V. Panggalo, Safarina G Malik, Khin Saw Myint, Amin |
| EPI_ISL_2364638 | RS Mary Cileungsi Hijau | Eijkman Institute for Molecular Biology, National Agency for Research and Innovation | Willy Agustine, Hidayat Trimarsanto, Lydia V. Panggalo, Iskandar Adnan, Sukma Oktavianthi, Lidwina Priliani, Edison Johar, Frilasita A Yudhaputri, Muhammad Rezki Rasyak, Safarina G Malik, Khin Saw Myint, Amin |
| EPI_ISL_2364641 | RS Mary Cileungsi Hijau | Eijkman Institute for Molecular Biology, National Agency for Research and Innovation | Frilasita A Yudhaputri, Muhammad Rezki Rasyak, Willy Agustine, Hidayat Trimarsanto, Lydia V. Panggalo, Iskandar Adnan, Sukma Oktavianthi, Lidwina Priliani, Edison Johar, Safarina G Malik, Khin Saw Myint, Amin |
| EPI_ISL_2364644 | RSUD Cileungsi | Eijkman Institute for Molecular Biology, National Agency for Research and Innovation | Sukma Oktavianthi, Lidwina Priliani, Edison Johar, Frilasita A Yudhaputri, Muhammad Rezki Rasyak, Willy Agustine, Hidayat Trimarsanto, Lydia V. Panggalo, Iskandar Adnan, Safarina G Malik, Khin Saw Myint, Amin |
| EPI_ISL_2364646 | RS Permata Cibubur | Eijkman Institute for Molecular Biology, National Agency for Research and Innovation | Lidwina Priliani, Edison Johar, Frilasita A Yudhaputri, Muhammad Rezki Rasyak, Willy Agustine, Hidayat Trimarsanto, Lydia V. Panggalo, Iskandar Adnan, Sukma Oktavianthi, Safarina G Malik, Khin Saw Myint, Amin |
| EPI_ISL_2364650 | RS Permata Cibubur | Eijkman Institute for Molecular Biology, National Agency for Research and Innovation | Edison Johar, Frilasita A Yudhaputri, Muhammad Rezki Rasyak, Willy Agustine, Hidayat Trimarsanto, Lydia V. Panggalo, Iskandar Adnan, Sukma Oktavianthi, Lidwina Priliani, Safarina G Malik, Khin Saw Myint, Amin |
| EPI_ISL_2364653 | RSU HGA Depok | Eijkman Institute for Molecular Biology, National Agency for Research and Innovation | Willy Agustine, Hidayat Trimarsanto, Lydia V. Panggalo, Iskandar Adnan, Sukma Oktavianthi, Lidwina Priliani, Edison Johar, Frilasita A Yudhaputri, Muhammad Rezki Rasyak, Safarina G Malik, Khin Saw Myint, Amin |
| EPI_ISL_2364656 | RSIA Assyifa | Eijkman Institute for Molecular Biology, National Agency for Research and Innovation | Hidayat Trimarsanto, Lydia V. Panggalo, Iskandar Adnan, Sukma Oktavianthi, Lidwina Priliani, Edison Johar, Frilasita A Yudhaputri, Muhammad Rezki Rasyak, Willy Agustine, Safarina G Malik, Khin Saw Myint, Amin |
| EPI_ISL_2364659 | RS Medirossa Cikarang | Eijkman Institute for Molecular Biology, National Agency for Research and Innovation | Hidayat Trimarsanto, Lydia V. Panggalo, Iskandar Adnan, Sukma Oktavianthi, Lidwina Priliani, Edison Johar, Frilasita A Yudhaputri, Muhammad Rezki Rasyak, Willy Agustine, Safarina G Malik, Khin Saw Myint, Amin |
| EPI_ISL_2364663 | RS Medirossa Cikarang | Eijkman Institute for Molecular Biology, National Agency for Research and Innovation | Frilasita A Yudhaputri, Muhammad Rezki Rasyak, Willy Agustine, Hidayat Trimarsanto, Lydia V. Panggalo, Iskandar Adnan, Sukma Oktavianthi, Lidwina Priliani, Edison Johar, Safarina G Malik, Khin Saw Myint, Amin |
| EPI_ISL_2364666 | RSIA PKU Muhammadiyah | Eijkman Institute for Molecular Biology, National Agency for Research and Innovation | Willy Agustine, Hidayat Trimarsanto, Lydia V. Panggalo, Iskandar Adnan, Sukma Oktavianthi, Lidwina Priliani, Edison Johar, Frilasita A Yudhaputri, Muhammad Rezki Rasyak, Safarina G Malik, Khin Saw Myint, Amin |
| EPI_ISL_2364669 | RSIA PKU Muhammadiyah | Eijkman Institute for Molecular Biology, National Agency for Research and Innovation | Muhammad Rezki Rasyak, Willy Agustine, Hidayat Trimarsanto, Lydia V. Panggalo, Iskandar Adnan, Sukma Oktavianthi, Lidwina Priliani, Edison Johar, Frilasita A Yudhaputri, Safarina G Malik, Khin Saw Myint, Amin |
| EPI_ISL_2364672 | RSJ Dr. Soeharto Heerdjan | Eijkman Institute for Molecular Biology, National Agency for Research and Innovation | Hidayat Trimarsanto, Lydia V. Panggalo, Iskandar Adnan, Sukma Oktavianthi, Lidwina Priliani, Edison Johar, Frilasita A Yudhaputri, Muhammad Rezki Rasyak, Willy Agustine, Safarina G Malik, Khin Saw Myint, Amin |
| EPI_ISL_2364675 | RS Yarsi | Eijkman Institute for Molecular Biology, National Agency for Research and Innovation | Lidwina Priliani, Edison Johar, Frilasita A Yudhaputri, Muhammad Rezki Rasyak, Willy Agustine, Hidayat Trimarsanto, Lydia V. Panggalo, Iskandar Adnan, Sukma Oktavianthi, Safarina G Malik, Khin Saw Myint, Amin |
| EPI_ISL_2382402, EPI_ISL_2382403, EPI_ISL_2382404, EPI_ISL_2382405, EPI_ISL_2382406, EPI_ISL_2382407 | FarmaLab Indoutama Halim | National Institute of Health Research and Development | Vivi Setiawaty, Hana Apsari Pawestri, Hartanti Dian Ikawati, Subangkit, Kartika Dewi Puspa, Arie Ardiansyah Nugraha, Triyani Soekarso, Krisna Pangesti, Nelly Puspandari |
| EPI_ISL_2382408, EPI_ISL_2382409, EPI_ISL_2382410 | Wisma Atlet | National Institute of Health Research and Development | Vivi Setiawaty, Hana Apsari Pawestri, Hartanti Dian Ikawati, Subangkit, Kartika Dewi Puspa, Arie Ardiansyah Nugraha, Triyani Soekarso, Krisna Pangesti, Nelly Puspandari |
| EPI_ISL_2382411, EPI_ISL_2382412 | FarmaLab Indoutama (Wisma Atlet) | National Institute of Health Research and Development | Vivi Setiawaty, Hana Apsari Pawestri, Hartanti Dian Ikawati, Subangkit, Kartika Dewi Puspa, Arie Ardiansyah Nugraha, Triyani Soekarso, Krisna Pangesti, Nelly Puspandari |
| EPI_ISL_2382413, EPI_ISL_2382414, | Smartco Lab | National Institute of Health Research and Development | Vivi Setiawaty, Hana Apsari Pawestri, Hartanti Dian Ikawati, Subangkit, Kartika Dewi Puspa, Arie Ardiansyah Nugraha, Triyani Soekarso, Krisna Pangesti, Nelly Puspandari |

| | | | |
|---|---|---|---|
| EPI_ISL_2382415 | | | |
| EPI_ISL_2382416 | RSPI Prof. Dr. Sulianti Saroso | National Institute of Health Research and Development | Vivi Setiawaty, Hana Apsari Pawestri, Hartanti Dian Ikawati, Subangkit, Kartika Dewi Puspa, Arie Ardiansyah Nugraha, Triyani Soekarso, Krisna Pangesti, Nelly Puspandari |
| EPI_ISL_2429144 | National Institute of Health Research and Development | National Institute of Health Research and Development | Vivi Setiawaty, Hana Apsari Pawestri, Hartanti Dian Ikawati, Subangkit, Kartika Dewi Puspa, Arie Ardiansyah Nugraha, Triyani Soekarso, Krisna Pangesti, Nelly Puspandari |
| EPI_ISL_2429146 | FarmaLab Indoutama Halim | National Institute of Health Research and Development | Vivi Setiawaty, Hana Apsari Pawestri, Hartanti Dian Ikawati, Subangkit, Kartika Dewi Puspa, Arie Ardiansyah Nugraha, Triyani Soekarso, Krisna Pangesti, Nelly Puspandari |
| EPI_ISL_2434061, EPI_ISL_2434123, EPI_ISL_2434157, EPI_ISL_2434161, EPI_ISL_2434168 | Genomik Solidaritas Indonesia Laboratorium | Genomik Solidaritas Indonesia Laboratorium | Alfin Mohammad Abdillah, Dhahlia Agustina Cahyono, Normastuti Adhini Tantyo, Annisa Muthiah Sukirman, Carissa Sintca Wijaya, Vania Gavrila Wikasa, Gracia Felias Enos Korompis, Meutia Ayuputeri Kumaheri, Anindya Pradipt |
| EPI_ISL_2434173, EPI_ISL_2434175, EPI_ISL_2434973, EPI_ISL_2434978 | Genomik Solidaritas Indonesia Laboratorium | Genomik Solidaritas Indonesia Laboratorium | Dhahlia Agustina Cahyono, Alfin Mohammad Abdillah, Normastuti Adhini Tantyo, Annisa Muthiah Sukirman, Carissa Sintca Wijaya, Vania Gavrila Wikasa, Gracia Felias Enos Korompis, Meutia Ayuputeri Kumaheri, Anindya Pradipt |
| EPI_ISL_2434988 | Genomik Solidaritas Indonesia Laboratorium | Genomik Solidaritas Indonesia Laboratorium | Alfin Mohammad Abdillah, Dhahlia Agustina Cahyono, Normastuti Adhini Tantyo, Annisa Muthiah Sukirman, Carissa Sintca Wijaya, Vania Gavrila Wikasa, Gracia Felias Enos Korompis, Meutia Ayuputeri Kumaheri, Anindya Pradipt |
| EPI_ISL_413219 | National Institute of Health Research and Development | National Institute of Health Research and Development | Setiawaty,V; Subangkit; Pawestri,HA; Puspa,KD; Ikawati,HD; Nugraha,AA; Hariastuti,NI; Ramadhany,R; Susilarini,NK; Pratiwi,E; Agustiningsih; Kurniawati,J; Siswanto |
| EPI_ISL_414375, EPI_ISL_414376, EPI_ISL_414377 | National Institute of Health Research and Development | National Institute of Health Research and Development | Setiawaty,V; Subangkit; Puspa,KD; Ikawati,HD; Nugraha, AA; Susilarini, NK; Agustiningsih; Ramadhany,R; Pratiwi,E; Hariastuti,NI; Kurniawati,J; Pawestri,HA; Siswanto |
| EPI_ISL_428207, EPI_ISL_428208 | National Institute of Health Research and Development | National Institute of Health Research and Development | Setiawaty,V;Subangkit;Puspa,KD;Ikawati,HD;Nugraha,AA;Hariastuti,NI;Ramadhany,R;Susilarini,NK;Pratiwi,E;Agustiningsih;Kurniawati,J;Pawestri,HA;Siswanto |
| EPI_ISL_435281 | Medistra Hospital Jakarta | Eijkman Institute for Molecular Biology, Ministry of Research and Technology/National Agency for Research and Innovation | Edison Johar, Frilasita A Yudhaputri, Hidayat Trimarsanto, David H Muljono, Safarina G Malik, Khin Saw Myint, Amin Soebandrio |
| EPI_ISL_435282, EPI_ISL_435283 | RS Pondok Indah Hospital - Pondok Indah | Eijkman Institute for Molecular Biology, Ministry of Research and Technology/National Agency for Research and Innovation | Edison Johar, Frilasita A Yudhaputri, Hidayat Trimarsanto, David H Muljono, Safarina G Malik, Khin Saw Myint, Amin Soebandrio |
| EPI_ISL_437187 | Siloam Hospitals | Institute of Tropical Disease, Universitas Airlangga | Kazufumi Shimizu, Krisnoadi Rahardjo, Aldise M Nastri, Jezzy R Dewantari, Rima R Prasetya, Maria M Padmidewi, Gatot Soegiarto, Laksmi Wulandari, Retno A Setyoningrum, Resti Y Meliana, Yohko K Shimizu, Mitsuhiro Nishimura, Yasu |
| EPI_ISL_437188 | RSUD Dr. Soetomo | Institute of Tropical Disease, Universitas Airlangga | Krisnoadi Rahardjo, Aldise M Nastri, Jezzy R Dewantari, Rima R Prasetya, Joni Wahyuhadi, Gatot Soegiarto, Laksmi Wulandari, Retno A Setyoningrum, Resti Y Meliana, Yohko K Shimizu, Mitsuhiro Nishimura, Yasuko Mori, Soetjipto, Ka |
| EPI_ISL_437189 | Pusat Pertamina Hospital | Eijkman Institute for Molecular Biology, Ministry of Research and Technology/National Agency for Research and Innovation | Edison Johar, Frilasita A Yudhaputri, Hidayat Trimarsanto, David H Muljono, Safarina G Malik, Khin Saw Myint, Amin Soebandrio |
| EPI_ISL_437190, EPI_ISL_437191 | RS Pondok Indah Hospital - Pondok Indah | Eijkman Institute for Molecular Biology, Ministry of Research and Technology/National Agency for Research and Innovation | Edison Johar, Frilasita A Yudhaputri, Hidayat Trimarsanto, David H Muljono, Safarina G Malik, Khin Saw Myint, Amin Soebandrio |
| EPI_ISL_437192 | Mitra Keluarga Kelapa Gading Hospital | Eijkman Institute for Molecular Biology, Ministry of Research and Technology/National Agency for Research and Innovation | Edison Johar, Frilasita A Yudhaputri, Hidayat Trimarsanto, David H Muljono, Safarina G Malik, Khin Saw Myint, Amin Soebandrio |
| EPI_ISL_438546, EPI_ISL_438547, EPI_ISL_438548, EPI_ISL_438549 | Siloam Hospital Lippo Village | Mochtar Riady Institute for Nanotechnology, Universitas Pelita Harapan | Aksar C Lages, David Rustandi, Febi Andriani, Ivet M Suriapranata, Riska N Taufik, Tri Shinta Kurniasih, Irawan Yusuf |
| EPI_ISL_458079 | Mitra Keluarga Hospital Kenjeran | Institute of Tropical Disease, Universitas Airlangga | Aldise M Nastri, Jezzy R Dewantari, Rima R Prasetya, Krisnoadi Rahardjo, Anastasia W Jefuna, Gatot Soegiarto, Laksmi Wulandari, Retno A Setyoningrum, Resti Yudhawati, Yohko K Shimizu, Mitsuhiro Nishimura, Yasuko Mori, Soetjipto, |
| EPI_ISL_458081 | RSUD Bangil Pasuruan | Institute of Tropical Disease, Universitas Airlangga | Jezzy R Dewantari, Rima R Prasetya, Krisnoadi Rahardjo, Aldise M Nastri, Arma Roosalina, Gatot Soegiarto, Laksmi Wulandari, Retno A Setyoningrum, Resti Yudhawati, Yohko K Shimizu, Mitsuhiro Nishimura, Yasuko Mori, Soetjipto, Ka |
| EPI_ISL_458082 | Universitas Airlangga Hospital | Institute of Tropical Disease, Universitas | Kazufumi Shimizu, Krisnoadi Rahardjo, Aldise M Nastri, Jezzy R Dewantari, Rima R Prasetya, Nasronudin, Gatot Soegiarto, Laksmi Wulandari, Retno A Setyoningrum, Resti Yudhawati, Yohko K Shimizu, Mitsuhiro Nishimura, Yasuko I |

| | | | |
|---|---|---|---|
| | | Airlangga | |
| EPI_ISL_458083 | Adi Husada Undaan Hospital | Institute of Tropical Disease, Universitas Airlangga | Rima R Prasetya, Krisnoadi Rahardjo, Aldise M Nastri, Jezzy R Dewantari, Irawati Marga, Gatot Soegiarto, Laksmi Wulandari, Retno A Setyoningrum, Resti Yudhawati, Yohko K Shimizu, Mitsuhiro Nishimura, Yasuko Mori, Soetjipto, Ka |
| EPI_ISL_467374 | Dinkes Samarinda | Eijkman Institute for Molecular Biology, Ministry of Research and Technology/National Agency for Research and Innovation | Edison Johar, Frilasita A Yudhaputri, Hidayat Trimarsanto, David H Muljono, Safarina G Malik, Khin Saw Myint, Amin Soebandrio |
| EPI_ISL_467375 | RSUP Prof. Dr. R. Kandou Manado | Eijkman Institute for Molecular Biology, Ministry of Research and Technology/National Agency for Research and Innovation | Edison Johar, Frilasita A Yudhaputri, Hidayat Trimarsanto, David H Muljono, Safarina G Malik, Khin Saw Myint, Amin Soebandrio |
| EPI_ISL_467376 | RSUP Fatmawati | Eijkman Institute for Molecular Biology, Ministry of Research and Technology/National Agency for Research and Innovation | Edison Johar, Frilasita A Yudhaputri, Hidayat Trimarsanto, David H Muljono, Safarina G Malik, Khin Saw Myint, Amin Soebandrio |
| EPI_ISL_511878 | Santosa Hospital Bandung Kopo | Laboratorium Kesehatan Provinsi Jawa Barat; Molecular Genetics Laboratory-Faculty of Medicine-Universitas Padjadjaran; School of Life Sciences and Technology & School of Pharmacy-Institut Teknologi Bandung | Ema Rahmawati, Marselina Irasonia Tan, Yunia Sribudiani, Catur Riani, Azzania Fibriani, Husna Nugrahapraja, Tarwadi, Hesti Lina Wiraswati, Lia Faridah, Savira Ekawardhani, Ryan Bayusantika Ristandi, Rifky Waluyajati Rachman, Cut Nur Cinthia Alamanda, H |
| EPI_ISL_511879 | Laboratorium Kesehatan Provinsi Jawa Barat | Molecular Genetics Laboratory-Faculty of Medicine-Universitas Padjadjaran; School of Life Sciences and Technology & School of Pharmacy-Institut Teknologi Bandung; Laboratorium Kesehatan Provinsi Jawa Barat | Marselina Irasonia Tan, Yunia Sribudiani, Catur Riani, Azzania Fibriani, Husna Nugrahapraja, Tarwadi, Ema Rahmawati, Hesti Lina Wiraswati, Lia Faridah, Savira Ekawardhani, Ryan Bayusantika Ristandi, Rifky Waluyajati Rachman, Cut Nur Cinthia Alamanda, H |
| EPI_ISL_516800 | Rumah Sakit Akademik Universitas Gadjah Mada | Genetics Working Group (Pokja Genetik) Faculty of Medicine, Public Health and Nursing Universitas Gadjah Mada (FK-KMK UGM); Disease Investigation Center Wates Ministry of Agriculture Indonesia; Department of Microbiology FK-KMK UGM; Laboratorium Diagnostik Yayasan Tahija World Mosquito Program (WMP) Yogyakarta Center for Tropical Medicine FK-KMK UGM; Integrated Research center FK-KMK UGM; Department of Computer Science and Electronics FMIPA UGM | Gunadi, Hendra Wibawa, . Marcellus, Mohamad S. Hakim, Edwin W. Daniwijaya, Ludhang P. Rizki, Endah Supriyati, Eggi Arguni, Titik Nuryastuti, Tri Wibawa, Dwi AA Nugrahaningsih, Afiahayati, Siswanto, Alvin S. Kalim, |
| EPI_ISL_516806 | Rumah Sakit PKU Gamping | Genetics Working Group (Pokja Genetik) Faculty of Medicine, Public Health and Nursing Universitas Gadjah Mada (FK-KMK UGM); Disease Investigation Center Wates Ministry of Agriculture Indonesia; Department of Microbiology FK-KMK UGM; Laboratorium Diagnostik Yayasan Tahija World Mosquito Program (WMP) Yogyakarta Center for Tropical Medicine FK-KMK UGM; Integrated Research center FK-KMK UGM; Department of Computer Science and Electronics FMIPA UGM | Gunadi, Hendra Wibawa, . Marcellus, Mohamad S. Hakim, Edwin W. Daniwijaya, Ludhang P. Rizki, Endah Supriyati, Eggi Arguni, Titik Nuryastuti, Tri Wibawa, Dwi AA Nugrahaningsih, . Afiahayati , . Siswanto, Ardorisye Saptaty F |
| EPI_ISL_516829 | RSUD Nyi Ageng Serang | Genetics Working Group (Pokja Genetik) Faculty of Medicine, Public Health and | Gunadi, Hendra Wibawa, . Marcellus, Mohamad S. Hakim, Edwin W. Daniwijaya, Ludhang P. Rizki, Endah Supriyati, Eggi Arguni, Titik Nuryastuti, Tri Wibawa, Dwi AA Nugrahaningsih, Afiahayati , . Siswanto, Beby Dewi Sartika, |

| | | | |
|---|---|---|---|
| | | Nursing Universitas Gadjah Mada (FK-KMK UGM), Disease Investigation Center Wates Ministry of Agriculture Indonesia, Department of Microbiology FK-KMK UGM, Laboratorium Diagnostik Yayasan Tahija World Mosquito Program (WMP) Yogyakarta Center for Tropical Medicine FK-KMK UGM, Integrated Research Center FK-KMK UGM, Department of Computer Science and Electronics FMIPA UGM | |
| EPI_ISL_518819 | Qadr Hospital, Tangerang, Banten | Biosafety Level-3 Laboratory, Indonesian Institute of Sciences (LIPI) | Anik Budhi Dharmayanthi, Syam Budi Iryanto, Andri Wardiana, Anggia Prasetyoputri, Isa Nuryana, Ade Andriani, Ario Betha Juanssilfero, Asep M Ridwanuloh, Ahmad Fathoni, Rifki Sadikin, Ratih Asmana Ningrum, Wien Kushar |
| EPI_ISL_518820 | Klinik Apotek Dein, Jakarta, Indonesia | Biosafety Level-3 Laboratory, Indonesian Institute of Sciences (LIPI) | Anik Budhi Dharmayanthi, Syam Budi Iryanto, Andri Wardiana, Anggia Prasetyoputri, Isa Nuryana, Ade Andriani, Ario Betha Juanssilfero, Asep M Ridwanuloh, Ahmad Fathoni, Rifki Sadikin, Ratih Asmana Ningrum, Wien Kushar |
| EPI_ISL_525492 | RSUP dr. SOERADJI TIRTONEGORO | Genetics Working Group (Pokja Genetik) Faculty of Medicine, Public Health and Nursing Universitas Gadjah Mada (FK-KMK UGM), Disease Investigation Center Wates Ministry of Agriculture Indonesia, Department of Microbiology FK-KMK UGM, Laboratorium Diagnostik Yayasan Tahija World Mosquito Program (WMP) Yogyakarta Center for Tropical Medicine FK-KMK UGM, Integrated Research Center FK-KMK UGM, Department of Computer Science and Electronics FMIPA UGM | Gunadi, Hendra Wibawa, . Marcellus, Mohamad S. Hakim, Edwin W. Daniwijaya, Ludhang P. Rizki, Endah Supriyati, Eggi Arguni, Titik Nuryastuti, Tri Wibawa, Dwi AA Nugrahaningsih, Afiahayati , . Siswanto, Kurniyanto, Indah Juliana, |
| EPI_ISL_528745 | Laboratorium Kesehatan Provinsi Jawa Barat | School of Life Sciences and Technology & School of Pharmacy-Institut Teknologi Bandung; Molecular Genetics Laboratory-Faculty of Medicine-Universitas Padjadjaran; Laboratorium Kesehatan Provinsi Jawa Barat | Marselina Irasonia Tan, Yunia Sribudiani, Catur Riani, Azzania Fibriani, Husna Nugrahapraja, Tarwadi, Ema Rahmawati, Savira Ekawardhani, Hesti Lina Wiraswati, Ryan Bayusantika Ristandi, Rifky Waluyajati Rachman, Cut Nur Cinthia Alamanda, Lia Faridah, Tri Amalina, Hammam Riza, Sony Solistia Wirawan, Agung Eru Wibowo, Irvan Faizal |
| EPI_ISL_528746 | Immanuel Hospital | Molecular Genetics Laboratory-Faculty of Medicine-Universitas Padjadjaran; School of Life Sciences and Technology & School of Pharmacy-Institut Teknologi Bandung; Laboratorium Kesehatan Provinsi Jawa Barat | Yunia Sribudiani,Tri Hanggono Achmad, Mas Rizky A.A. Syamsunarno, Fensi Amalina, Catur Riani, Azzania Fibriani, Husna Nugrahapraja, Marselina Irasonia Tan, Tarwadi, Ema Rahmawati, Savira Ekawardhani, Hesti Lina Wiraswati, Ryan Bayusantika Ristandi, Faridah, Hammam Riza, Sony Solistia Wirawan, Agung Eru Wibowo, Irvan Faizal |
| EPI_ISL_528747 | Santo Borromeus Hospital | School of Pharmacy & School of Life Sciences and Technology - Institut Teknologi Bandung; Molecular Genetics Laboratory-Faculty of Medicine-Universitas Padjadjaran; Laboratorium Kesehatan Provinsi Jawa Barat | Catur Riani, Marselina Irasonia Tan, Yunia Sribudiani, Azzania Fibriani, Husna Nugrahapraja, Tarwadi, Ema Rahmawati, Savira Ekawardhani, Hesti Lina Wiraswati, Ryan Bayusantika Ristandi, Rifky Waluyajati Rachman, Cut Nur Cinthia Alamanda, Lia Faridah, M Solistia Wirawan, Agung Eru Wibowo, Irvan Faizal |
| EPI_ISL_528748 | Dinkes Provinsi Jawa Barat | School of Life Sciences and Technology & School of Pharmacy-Institut Teknologi Bandung; Molecular Genetics Laboratory-Faculty of Medicine-Universitas Padjadjaran; Laboratorium Kesehatan Provinsi Jawa Barat | Azzania Fibriani, Catur Riani, Marselina Irasonia Tan, Yunia Sribudiani, Husna Nugrahapraja, Tarwadi, Ema Rahmawati, Savira Ekawardhani, Hesti Lina Wiraswati, Ryan Bayusantika Ristandi, Rifky Waluyajati Rachman, Cut Nur Cinthia Alamanda, Lia Faridah, Gu Riza, Sony Solistia Wirawan, Agung Eru Wibowo, Irvan Faizal |
| EPI_ISL_528749 | Santosa Hospital | School of Life Sciences and | Husna Nugrahapraja, Azzania Fibriani, Catur Riani, Marselina Irasonia Tan, Yunia Sribudiani, Tarwadi, Ema Rahmawati, Savira Ekawardhani, Hesti Lina Wiraswati, Ryan Bayusantika Ristandi, Rifky Waluyajati Rachman, Cut Nur Cinthia Alamanda, Lia Faridah, Da |

| | | | |
|---|---|---|---|
| | Bandung Central | Technology & School of Pharmacy-Institut Teknologi Bandung; Molecular Genetics Laboratory-Faculty of Medicine-Universitas Padjadjaran; Laboratorium Kesehatan Provinsi Jawa Barat | Solistia Wirawan, Agung Eru Wibowo, Irvan Faizal |
| EPI_ISL_528750 | Santo Borromeus Hospital | School of Life Sciences and Technology & School of Pharmacy-Institut Teknologi Bandung; Molecular Genetics Laboratory-Faculty of Medicine-Universitas Padjadjaran; Laboratorium Kesehatan Provinsi Jawa Barat | Marselina Irasonia Tan, Yunia Sribudiani, Catur Riani, Azzania Fibriani, Husna Nugrahapraja, Tarwadi, Ema Rahmawati, Savira Ekawardhani, Hesti Lina Wiraswati, Ryan Bayusantika Ristandi, Rifky Waluyajati Rachman, Cut Nur Cinthia Alamanda, Lia Faridah, M Solistia Wirawan, Agung Eru Wibowo, Irvan Faizal |
| EPI_ISL_528751 | Santo Borromeus Hospital | Molecular Genetics Laboratory-Faculty of Medicine-Universitas Padjadjaran; School of Life Sciences and Technology & School of Pharmacy-Institut Teknologi Bandung; Laboratorium Kesehatan Provinsi Jawa Barat | Yunia Sribudiani, Tri Hanggono Achmad, Mas Rizky A.A. Syamsunarno, Fensi Amalina, Catur Riani, Azzania Fibriani, Husna Nugrahapraja, Marselina Irasonia Tan, Tarwadi, Ema Rahmawati, Savira Ekawardhani, Hesti Lina Wiraswati, Ryan Bayusantika Ristandi, Faridah, Hammam Riza, Sony Solistia Wirawan, Agung Eru Wibowo, Irvan Faizal |
| EPI_ISL_528752 | Dr. H. A. Rotinsulu Lung Hospital | School of Pharmacy & School of Life Sciences and Technology - Institut Teknologi Bandung; Molecular Genetics Laboratory-Faculty of Medicine-Universitas Padjadjaran; Laboratorium Kesehatan Provinsi Jawa Barat | Catur Riani, Marselina Irasonia Tan, Yunia Sribudiani, Azzania Fibriani, Husna Nugrahapraja, Tarwadi, Ema Rahmawati, Savira Ekawardhani, Hesti Lina Wiraswati, Ryan Bayusantika Ristandi, Rifky Waluyajati Rachman, Cut Nur Cinthia Alamanda, Lia Faridah, Gu Riza, Sony Solistia Wirawan, Agung Eru Wibowo, Irvan Faizal |
| EPI_ISL_528753 | Dinkes Kota Bogor | School of Life Sciences and Technology & School of Pharmacy-Institut Teknologi Bandung; Molecular Genetics Laboratory-Faculty of Medicine-Universitas Padjadjaran; Laboratorium Kesehatan Provinsi Jawa Barat | Azzania Fibriani, Catur Riani, Marselina Irasonia Tan, Yunia Sribudiani, Husna Nugrahapraja, Tarwadi, Ema Rahmawati, Savira Ekawardhani, Hesti Lina Wiraswati, Ryan Bayusantika Ristandi, Rifky Waluyajati Rachman, Cut Nur Cinthia Alamanda, Lia Faridah, Da Solistia Wirawan, Agung Eru Wibowo, Irvan Faizal |
| EPI_ISL_528759 | Santo Borromeus Hospital | School of Life Sciences and Technology & School of Pharmacy-Institut Teknologi Bandung; Molecular Genetics Laboratory-Faculty of Medicine-Universitas Padjadjaran; Laboratorium Kesehatan Provinsi Jawa Barat | Husna Nugrahapraja, Azzania Fibriani, Catur Riani, Marselina Irasonia Tan, Yunia Sribudiani, Tarwadi, Ema Rahmawati, Savira Ekawardhani, Hesti Lina Wiraswati, Ryan Bayusantika Ristandi, Rifky Waluyajati Rachman, Cut Nur Cinthia Alamanda, Lia Faridah, Tri Amalina, Hammam Riza, Sony Solistia Wirawan, Agung Eru Wibowo, Irvan Faizal |
| EPI_ISL_529138 | RSAL Dr. Ramelan Surabaya | Institute of Tropical Disease, Universitas Airlangga | Jezzy R Dewantari, Rima R Prasetya, Krisnoadi Rahardjo, Aldise M Nastri, Radito Soesanto, Gatot Soegiarto, Laksmi Wulandari, Retno A Setyoningrum, Resti Yudhawati, Yohko K Shimizu, Mitsuhiro Nishimura, Yasuko Mori, Soetjipto, K |
| EPI_ISL_529718 | Santo Borromeus Hospital | Laboratorium Kesehatan Provinsi Jawa Barat; School of Life Sciences and Technology & School of Pharmacy-Institut Teknologi Bandung; Molecular Genetics Laboratory-Faculty of Medicine-Universitas Padjadjaran | Ema Rahmawati, Marselina Irasonia Tan, Yunia Sribudiani, Catur Riani, Azzania Fibriani, Husna Nugrahapraja, Tarwadi, Savira Ekawardhani, Hesti Lina Wiraswati, Ryan Bayusantika Ristandi, Rifky Waluyajati Rachman, Cut Nur Cinthia Alamanda, Lia Faridah, Gu Riza, Sony Solistia Wirawan, Agung Eru Wibowo, Irvan Faizal |
| EPI_ISL_529719 | Sumedang Regional General Hospital | Laboratorium Kesehatan Provinsi Jawa Barat; School of Life Sciences and Technology & School of Pharmacy-Institut Teknologi Bandung; Molecular Genetics Laboratory-Faculty of Medicine-Universitas Padjadjaran | Ema Rahmawati, Marselina Irasonia Tan, Yunia Sribudiani, Catur Riani, Azzania Fibriani, Husna Nugrahapraja, Tarwadi, Savira Ekawardhani, Hesti Lina Wiraswati, Ryan Bayusantika Ristandi, Rifky Waluyajati Rachman, Cut Nur Cinthia Alamanda, Lia Faridah, Gu Riza, Sony Solistia Wirawan, Agung Eru Wibowo, Irvan Faizal |
| EPI_ISL_529961 | Husada Utama Hospital | Institute of Tropical Disease, Universitas | Krisnoadi Rahardjo, Aldise M Nastri, Jezzy R Dewantari, Rima R Prasetya, Didi Dewanto, Gatot Soegiarto, Laksmi Wulandari, Retno A Setyoningrum, Resti Yudhawati, Yohko K Shimizu, Mitsuhiro Nishimura, Yasuko Mori, Soetjipto, Ka |

| | | | |
|---|---|---|---|
| | | Airlangga | |
| EPI_ISL_529962 | Universitas Airlangga Hospital | Institute of Tropical Disease, Universitas Airlangga | Jezzy R Dewantari, Rima R Prasetya, Krisnoadi Rahardjo, Aldise M Nastri, Nasronudin, Gatot Soegiarto, Laksmi Wulandari, Retno A Setyoningrum, Resti Yudhawati, Yohko K Shimizu, Mitsuhiro Nishimura, Yasuko Mori, Soetjipto, Kaz |
| EPI_ISL_529963 | Universitas Airlangga Hospital | Institute of Tropical Disease, Universitas Airlangga | Rima R Prasetya, Krisnoadi Rahardjo, Aldise M Nastri, Jezzy R Dewantari, Nasronudin, Gatot Soegiarto, Laksmi Wulandari, Retno A Setyoningrum, Resti Yudhawati, Yohko K Shimizu, Mitsuhiro Nishimura, Yasuko Mori, Soetjipto, Kaz |
| EPI_ISL_529964 | RSUD Bangil Pasuruan | Institute of Tropical Disease, Universitas Airlangga | Aldise M Nastri, Jezzy R Dewantari, Rima R Prasetya, Krisnoadi Rahardjo, Arma Roosalina, Gatot Soegiarto, Laksmi Wulandari, Retno A Setyoningrum, Resti Yudhawati, Yohko K Shimizu, Mitsuhiro Nishimura, Yasuko Mori, Soetjipto, Ka |
| EPI_ISL_529965 | RSUD Dr. Soetomo | Institute of Tropical Disease, Universitas Airlangga | Krisnoadi Rahardjo, Aldise M Nastri, Jezzy R Dewantari, Rima R Prasetya, Joni Wahyuhadi, Gatot Soegiarto, Laksmi Wulandari, Retno A Setyoningrum, Resti Yudhawati, Yohko K Shimizu, Mitsuhiro Nishimura, Yasuko Mori, Soetjipto, Ka |
| EPI_ISL_529966 | RSUD Sidoarjo | Institute of Tropical Disease, Universitas Airlangga | Kazufumi Shimizu, Krisnoadi Rahardjo, Aldise M Nastri, Jezzy R Dewantari, Rima R Prasetya, Atok Irawan, Gatot Soegiarto, Laksmi Wulandari, Retno A Setyoningrum, Resti Yudhawati, Yohko K Shimizu, Mitsuhiro Nishimura, Yasuko |
| EPI_ISL_529967 | RSUD Sidoarjo | Institute of Tropical Disease, Universitas Airlangga | Aldise M Nastri, Jezzy R Dewantari, Rima R Prasetya, Krisnoadi Rahardjo, Atok Irawan, Gatot Soegiarto, Laksmi Wulandari, Retno A Setyoningrum, Resti Yudhawati, Yohko K Shimizu, Mitsuhiro Nishimura, Yasuko Mori, Soetjipto, Kaz |
| EPI_ISL_538498 | RSUD Soediono Madiun East Java | National Institute of Health Research and Development | Pawestri, HA; Subangkit; Puspa, KD; Nugraha, AA; Ikawati, HD; Pangesti, KNA; Soekarso, T; Paisal; Setiawaty,V. |
| EPI_ISL_538499 | RS Lavallete Malang East Java | National Institute of Health Research and Development | Pawestri, HA; Subangkit; Puspa, KD; Nugraha, AA; Ikawati, HD; Pangesti, KNA; Soekarso, T; Paisal; Setiawaty,V. |
| EPI_ISL_538500, EPI_ISL_538501 | RS Prima Husada Cipta Medan North Sumatra | National Institute of Health Research and Development | Pawestri, HA; Subangkit; Puspa, KD; Nugraha, AA; Ikawati, HD; Pangesti, KNA; Soekarso, T; Paisal; Setiawaty,V. |
| EPI_ISL_538502 | RSUD Sultan Imanudin Pangkalan Bun Center Kalimantan | National Institute of Health Research and Development | Pawestri, HA; Subangkit; Puspa, KD; Nugraha, AA; Ikawati, HD; Pangesti, KNA; Soekarso, T; Paisal; Setiawaty,V. |
| EPI_ISL_538503 | RSUD Bali Mandara Denpasar Bali | National Institute of Health Research and Development | Pawestri, HA; Subangkit; Puspa, KD; Nugraha, AA; Ikawati, HD; Pangesti, KNA; Soekarso, T; Paisal; Setiawaty,V. |
| EPI_ISL_538504, EPI_ISL_538505 | National Institute of Health Research and Development | National Institute of Health Research and Development | Pawestri, HA; Subangkit; Puspa, KD; Nugraha, AA; Ikawati, HD; Pangesti, KNA; Soekarso, T; Susilarini, NK; Hariastuti, NI; Nikmah, UA; Mursinah; Febriyani, A; Herman, R; Susanti, N; Herna; Febriyanti, T; Nurhadi, M; Paisal; Ramadhany, R; Agustiningsih; Kurniav HA; Rizki, A; Puspandary, N; Setiawaty,V. |
| EPI_ISL_538506, EPI_ISL_538507 | Balai Penelitian dan Pengembangan Biomedis Papua | National Institute of Health Research and Development | Pawestri, HA; Subangkit; Puspa, KD; Nugraha, AA; Ikawati, HD; Pangesti, KNA; Soekarso, T; Paisal; Oktavian, A; Hutapea, HML; Setiawaty,V. |
| EPI_ISL_538508, EPI_ISL_538509 | National Institute of Health Research and Development | National Institute of Health Research and Development | Pawestri, HA; Subangkit; Puspa, KD; Nugraha, AA; Ikawati, HD; Pangesti, KNA; Soekarso, T; Susilarini, NK; Hariastuti, NI; Nikmah, UA; Mursinah; Febriyani, A; Herman, R; Susanti, N; Herna; Febriyanti, T; Nurhadi, M; Paisal; Ramadhany, R; Agustiningsih; Kurniav HA; Rizki, A; Puspandary, N; Setiawaty,V. |
| EPI_ISL_538510 | RSUD Ulin Banjarmasin South Kalimantan | National Institute of Health Research and Development | Pawestri, HA; Subangkit; Puspa, KD; Nugraha, AA; Ikawati, HD; Pangesti, KNA; Soekarso, T; Paisal; Pasaribu, M; Setiawaty,V. |
| EPI_ISL_538511 | RSUD Wahidin Sudirohusodo Mojokerto East Java | National Institute of Health Research and Development | Pawestri, HA; Subangkit; Puspa, KD; Nugraha, AA; Ikawati, HD; Pangesti, KNA; Soekarso, T; Paisal; Setiawaty,V. |
| EPI_ISL_538512 | Balai Penelitian dan Pengembangan Biomedis Papua | National Institute of Health Research and Development | Pawestri, HA; Subangkit; Puspa, KD; Nugraha, AA; Ikawati, HD; Pangesti, KNA; Soekarso, T; Paisal; Pasaribu, M; Setiawaty,V. |
| EPI_ISL_538513 | Provincial Health Laboratory Bekasi West Java | National Institute of Health Research and Development | Pawestri, HA; Subangkit; Puspa, KD; Nugraha, AA; Ikawati, HD; Pangesti, KNA; Soekarso, T; Paisal; Setiawaty,V. |
| EPI_ISL_560991 | RSUD Dr. Soetomo | Institute of Tropical Disease, Universitas Airlangga | Rima R Prasetya, Krisnoadi Rahardjo, Aldise M Nastri, Jezzy R Dewantari, Joni Wahyuhadi, Gatot Soegiarto, Laksmi Wulandari, Retno A Setyoningrum, Resti Yudhawati, Yohko K Shimizu, Mitsuhiro Nishimura, Yasuko Mori, Soetjipto, Ka |
| EPI_ISL_568686 | RSUD Prof. DR. Margono Soekarjo | Eijkman Institute for Molecular Biology, Ministry of Research and Technology/National Agency for Research and Innovation | Frilasita A Yudhaputri, Edison Johar, Hidayat Trimarsanto, Iskandar A Adnan, Willy Agustine, David H Muljono, Safarina G Malik, Herawati Sudoyo, Khin Saw Myint, Amin Soebandrio |
| EPI_ISL_568687 | RSUP Fatmawati | Eijkman Institute for Molecular Biology, Ministry of Research and Technology/National Agency for Research and Innovation | Frilasita A Yudhaputri, Edison Johar, Hidayat Trimarsanto, Iskandar A Adnan, Willy Agustine, David H Muljono, Safarina G Malik, Herawati Sudoyo, Khin Saw Myint, Amin Soebandrio |
| EPI_ISL_568688 | RS Pelni | Eijkman Institute for Molecular Biology, Ministry of Research and Technology/National Agency for Research and Innovation | Frilasita A Yudhaputri, Edison Johar, Hidayat Trimarsanto, Iskandar A Adnan, Willy Agustine, David H Muljono, Safarina G Malik, Herawati Sudoyo, Khin Saw Myint, Amin Soebandrio |
| EPI_ISL_568689 | RS Pondok Indah Puri Indah | Eijkman Institute for Molecular Biology, Ministry | Frilasita A Yudhaputri, Edison Johar, Hidayat Trimarsanto, Iskandar A Adnan, Willy Agustine, David H Muljono, Safarina G Malik, Herawati Sudoyo, Khin Saw Myint, Amin Soebandrio |

| | | | |
|---|---|---|---|
| | | of Research and Technology/National Agency for Research and Innovation | |
| EPI_ISL_568690, EPI_ISL_568691 | RSUP Fatmawati | Eijkman Institute for Molecular Biology, Ministry of Research and Technology/National Agency for Research and Innovation | Frilasita A Yudhaputri, Edison Johar, Hidayat Trimarsanto, Iskandar A Adnan, Willy Agustine, David H Muljono, Safarina G Malik, Herawati Sudoyo, Khin Saw Myint, Amin Soebandrio |
| EPI_ISL_568692 | RS Kramat 128 | Eijkman Institute for Molecular Biology, Ministry of Research and Technology/National Agency for Research and Innovation | Frilasita A Yudhaputri, Edison Johar, Hidayat Trimarsanto, Iskandar A Adnan, Willy Agustine, David H Muljono, Safarina G Malik, Herawati Sudoyo, Khin Saw Myint, Amin Soebandrio |
| EPI_ISL_568693 | RS Freeport Tembagapura | Eijkman Institute for Molecular Biology, Ministry of Research and Technology/National Agency for Research and Innovation | Frilasita A Yudhaputri, Edison Johar, Hidayat Trimarsanto, Iskandar A Adnan, Willy Agustine, David H Muljono, Safarina G Malik, Herawati Sudoyo, Khin Saw Myint, Amin Soebandrio |
| EPI_ISL_568694 | RSUP Prof. Dr. R. Kandou Manado | Eijkman Institute for Molecular Biology, Ministry of Research and Technology/National Agency for Research and Innovation | Frilasita A Yudhaputri, Edison Johar, Hidayat Trimarsanto, Iskandar A Adnan, Willy Agustine, David H Muljono, Safarina G Malik, Herawati Sudoyo, Khin Saw Myint, Amin Soebandrio |
| EPI_ISL_574433 | National Institute of Health Research and Development | National Institute of Health Research and Development | Pawestri,HA;Subangkit;Puspa,KD;Nugraha,AA;Ikawati,HD;Pangesti, KNA;Soekarso,T;Susilarini,NK;Hariastuti,NI;Nikmah,UA;Mursinah;Febriyani,A;Herman,R;Susanti,N;Herna;Febriyanti,T; Nurhadi,M; Paisal;Ramadhany,R;Agustiningsih;Kurniawati,J;Kipuw,NL;Muna,F;Indalau,IL;Adam,K;Wibowo,HA;Rizki,A;Puspandari,N;Setiawaty,V |
| EPI_ISL_574603 | RS Freeport Tembagapura | Eijkman Institute for Molecular Biology, Ministry of Research and Technology/National Agency for Research and Innovation | Frilasita A Yudhaputri, Edison Johar, Hidayat Trimarsanto, Iskandar A Adnan, Willy Agustine, David H Muljono, Safarina G Malik, Herawati Sudoyo, Khin Saw Myint, Amin Soebandrio |
| EPI_ISL_574604 | RSUP Prof. Dr. R. Kandou Manado | Eijkman Institute for Molecular Biology, Ministry of Research and Technology/National Agency for Research and Innovation | Frilasita A Yudhaputri, Edison Johar, Hidayat Trimarsanto, Iskandar A Adnan, Willy Agustine, David H Muljono, Safarina G Malik, Herawati Sudoyo, Khin Saw Myint, Amin Soebandrio |
| EPI_ISL_574605 | RSUP dr. Sardjito | Eijkman Institute for Molecular Biology, Ministry of Research and Technology/National Agency for Research and Innovation | Frilasita A Yudhaputri, Edison Johar, Hidayat Trimarsanto, Iskandar A Adnan, Willy Agustine, David H Muljono, Safarina G Malik, Herawati Sudoyo, Khin Saw Myint, Amin Soebandrio |
| EPI_ISL_574606 | RS Siloam Purwakarta | Eijkman Institute for Molecular Biology, Ministry of Research and Technology/National Agency for Research and Innovation | Frilasita A Yudhaputri, Edison Johar, Hidayat Trimarsanto, Iskandar A Adnan, Willy Agustine, David H Muljono, Safarina G Malik, Herawati Sudoyo, Khin Saw Myint, Amin Soebandrio |
| EPI_ISL_574607 | RS Husada | Eijkman Institute for Molecular Biology, Ministry of Research and Technology/National Agency for Research and Innovation | Frilasita A Yudhaputri, Edison Johar, Hidayat Trimarsanto, Iskandar A Adnan, Willy Agustine, David H Muljono, Safarina G Malik, Herawati Sudoyo, Khin Saw Myint, Amin Soebandrio |
| EPI_ISL_574608 | RS Mitra Keluarga Kelapa Gading | Eijkman Institute for Molecular Biology, Ministry of Research and Technology/National Agency for Research and Innovation | Frilasita A Yudhaputri, Edison Johar, Hidayat Trimarsanto, Iskandar A Adnan, Willy Agustine, David H Muljono, Safarina G Malik, Herawati Sudoyo, Khin Saw Myint, Amin Soebandrio |
| EPI_ISL_574609 | RSJPD Harapan Kita | Eijkman Institute for Molecular Biology, Ministry of Research and Technology/National Agency for Research and Innovation | Frilasita A Yudhaputri, Edison Johar, Hidayat Trimarsanto, Iskandar A Adnan, Willy Agustine, David H Muljono, Safarina G Malik, Herawati Sudoyo, Khin Saw Myint, Amin Soebandrio |
| EPI_ISL_574610 | Puskesmas Tambora | Eijkman Institute for Molecular Biology, Ministry of Research and Technology/National Agency for Research and Innovation | Frilasita A Yudhaputri, Edison Johar, Hidayat Trimarsanto, Iskandar A Adnan, Willy Agustine, David H Muljono, Safarina G Malik, Herawati Sudoyo, Khin Saw Myint, Amin Soebandrio |
| EPI_ISL_574611 | RS Premier Bintaro | Eijkman Institute for | Frilasita A Yudhaputri, Edison Johar, Hidayat Trimarsanto, Iskandar A Adnan, Willy Agustine, David H Muljono, Safarina G Malik, Herawati Sudoyo, Khin Saw Myint, Amin Soebandrio |

| | | | |
|---|---|---|---|
| | | Molecular Biology, Ministry of Research and Technology/National Agency for Research and Innovation | |
| EPI_ISL_574612 | RS Hermina Mekarsari | Eijkman Institute for Molecular Biology, Ministry of Research and Technology/National Agency for Research and Innovation | Frilasita A Yudhaputri, Edison Johar, Hidayat Trimarsanto, Iskandar A Adnan, Willy Agustine, David H Muljono, Safarina G Malik, Herawati Sudoyo, Khin Saw Myint, Amin Soebandrio |
| EPI_ISL_574613 | RS Harapan Bunda | Eijkman Institute for Molecular Biology, Ministry of Research and Technology/National Agency for Research and Innovation | Frilasita A Yudhaputri, Edison Johar, Hidayat Trimarsanto, Iskandar A Adnan, Willy Agustine, David H Muljono, Safarina G Malik, Herawati Sudoyo, Khin Saw Myint, Amin Soebandrio |
| EPI_ISL_574614 | RS Kramat 128 | Eijkman Institute for Molecular Biology, Ministry of Research and Technology/National Agency for Research and Innovation | Frilasita A Yudhaputri, Edison Johar, Hidayat Trimarsanto, Iskandar A Adnan, Willy Agustine, David H Muljono, Safarina G Malik, Herawati Sudoyo, Khin Saw Myint, Amin Soebandrio |
| EPI_ISL_574615 | RS Harapan Bunda | Eijkman Institute for Molecular Biology, Ministry of Research and Technology/National Agency for Research and Innovation | Frilasita A Yudhaputri, Edison Johar, Hidayat Trimarsanto, Iskandar A Adnan, Willy Agustine, David H Muljono, Safarina G Malik, Herawati Sudoyo, Khin Saw Myint, Amin Soebandrio |
| EPI_ISL_574616, EPI_ISL_574617 | RS Kramat 128 | Eijkman Institute for Molecular Biology, Ministry of Research and Technology/National Agency for Research and Innovation | Frilasita A Yudhaputri, Edison Johar, Hidayat Trimarsanto, Iskandar A Adnan, Willy Agustine, David H Muljono, Safarina G Malik, Herawati Sudoyo, Khin Saw Myint, Amin Soebandrio |
| EPI_ISL_574618 | RS Kramat 128 | Eijkman Institute for Molecular Biology, Ministry of Research and Technology/National Agency for Research and Innovation | Hidayat Trimarsanto, Frilasita A Yudhaputri, Edison Johar, Iskandar A Adnan, Willy Agustine, David H Muljono, Safarina G Malik, Herawati Sudoyo, Khin Saw Myint, Amin Soebandrio |
| EPI_ISL_574619, EPI_ISL_574620, EPI_ISL_574621, EPI_ISL_574622, EPI_ISL_574623 | BTKLPP Kelas I Manado | Eijkman Institute for Molecular Biology, Ministry of Research and Technology/National Agency for Research and Innovation | Hidayat Trimarsanto, Frilasita A Yudhaputri, Edison Johar, Iskandar A Adnan, Willy Agustine, David H Muljono, Safarina G Malik, Herawati Sudoyo, Khin Saw Myint, Amin Soebandrio |
| EPI_ISL_575331 | RSUD Wates | Genetics Working Group (Pokja Genetik) Faculty of Medicine, Public Health and Nursing Universitas Gadjah Mada (FK-KMK UGM); Disease Investigation Center Wates Ministry of Agriculture Indonesia; Department of Microbiology FK-KMK UGM; Laboratorium Diagnostik Yayasan Tahija World Mosquito Program (WMP) Yogyakarta Center for Tropical Medicine FK-KMK UGM; Integrated Research Center FK-KMK UGM; Department of Computer Science and Electronics FMIPA UGM | Gunadi, Hendra Wibawa, . Marcellus, Mohamad S. Hakim, Edwin W. Daniwijaya, Ludhang P. Rizki, Endah Supriyati, Eggi Arguni, Titik Nuryastuti, Tri Wibawa, Dwi AA Nugrahaningsih, Afiahayati, Siswanto, Kristy Iskandar, Nungki Anggorowati, Bastianto Kusum |
| EPI_ISL_576113 | RSUP Dr. Sardjito | Genetics Working Group (Pokja Genetik) Faculty of Medicine, Public Health and Nursing Universitas Gadjah Mada (FK-KMK UGM); Disease Investigation Center Wates Ministry of Agriculture Indonesia; Department of Microbiology FK-KMK UGM; Laboratorium Diagnostik Yayasan Tahija World Mosquito Program (WMP) Yogyakarta Center for Tropical Medicine FK-KMK UGM; Integrated Research | Gunadi, Hendra Wibawa, . Marcellus, Mohamad S. Hakim, Edwin W. Daniwijaya, Ludhang P. Rizki, Endah Supriyati, Eggi Arguni, Titik Nuryastuti, Tri Wibawa, Dwi AA Nugrahaningsih, Afiahayati, Siswanto, Kristy Iskandar, Nungki Anggorowati, Ika Trisna |

| | | | |
|---|---|---|---|
| | | Center FK-KMK UGM; Department of Computer Science and Electronics FMIPA UGM | |
| EPI_ISL_576114 | RSUP Dr. Sardjito | Genetics Working Group (Pokja Genetik) Faculty of Medicine, Public Health and Nursing Universitas Gadjah Mada (FK-KMK UGM); Disease Investigation Center Wates Ministry of Agriculture Indonesia; Department of Microbiology FK-KMK UGM; Laboratorium Diagnostik Yayasan Tahija World Mosquito Program (WMP) Yogyakarta Center for Tropical Medicine FK-KMK UGM; Integrated Research Center FK-KMK UGM; Department of Computer Science and Electronics FMIPA UGM | Gunadi, Hendra Wibawa, . Marcellus, Mohamad S. Hakim, Edwin W. Daniwijaya, Ludhang P. Rizki, Endah Supriyati, Eggi Arguni, Titik Nuryastuti, Tri Wibawa, Dwi AA Nugrahaningsih, Afiahayati, Siswanto, Kristy Iskandar, Nungki Anggorowati, Ika Trisnawati, |
| EPI_ISL_576115 | RSUP Dr Sardjito | Genetics Working Group (Pokja Genetik) Faculty of Medicine, Public Health and Nursing Universitas Gadjah Mada (FK-KMK UGM); Disease Investigation Center Wates Ministry of Agriculture Indonesia; Department of Microbiology FK-KMK UGM; Laboratorium Diagnostik Yayasan Tahija World Mosquito Program (WMP) Yogyakarta Center for Tropical Medicine FK-KMK UGM; Integrated Research Center FK-KMK UGM; Department of Computer Science and Electronics FMIPA UGM | Gunadi, Hendra Wibawa, . Marcellus, Mohamad S. Hakim, Edwin W. Daniwijaya, Ludhang P. Rizki, Endah Supriyati, Eggi Arguni, Titik Nuryastuti, Tri Wibawa, Dwi AA Nugrahaningsih, Afiahayati, Siswanto, Kristy Iskandar, Nungki Anggorowati, Ika Trisnawati, Riat E Intansari, Elizabeth Henny Herningtiyas, Ira Puspitawati, Nur Imma Fatimah Harahap, Untung Wirawan, Maria Patricia Inggriani |
| EPI_ISL_576116 | RSUP Dr. Sardjito | Genetics Working Group (Pokja Genetik) Faculty of Medicine, Public Health and Nursing Universitas Gadjah Mada (FK-KMK UGM); Disease Investigation Center Wates Ministry of Agriculture Indonesia; Department of Microbiology FK-KMK UGM; Laboratorium Diagnostik Yayasan Tahija World Mosquito Program (WMP) Yogyakarta Center for Tropical Medicine FK-KMK UGM; Integrated Research Center FK-KMK UGM; Department of Computer Science and Electronics FMIPA UGM | Gunadi, Hendra Wibawa, . Marcellus, Mohamad S. Hakim, Edwin W. Daniwijaya, Ludhang P. Rizki, Endah Supriyati, Eggi Arguni, Titik Nuryastuti, Tri Wibawa, Dwi AA Nugrahaningsih, Afiahayati, Siswanto, Kristy Iskandar, Nungki Anggorowati, Ika Trisnawati, Suma Munawar Gani, Satria Maulana, Nur Rahmi Ananda, Riat El Khair, Yunika Puspadewi, Osman Sianipar, Umi Solekhah Intansari, Elizabeth Henny Herningtiyas, Ira Puspitawati, Nur Imma Fatimah Harahap, Ririn Enggy Yuliyanti |
| EPI_ISL_576128 | RSUP Dr. Sardjito | Genetics Working Group (Pokja Genetik) Faculty of Medicine, Public Health and Nursing Universitas Gadjah Mada (FK-KMK UGM); Disease Investigation Center Wates Ministry of Agriculture Indonesia; Department of Microbiology FK-KMK UGM; Laboratorium Diagnostik Yayasan Tahija World Mosquito Program (WMP) Yogyakarta Center for Tropical Medicine FK-KMK UGM; Integrated Research Center FK-KMK UGM; Department of Computer Science and Electronics FMIPA UGM | Gunadi, Hendra Wibawa, Marcellus, Mohamad S. Hakim, Edwin W. Daniwijaya, Ludhang P. Rizki, Endah Supriyati, Eggi Arguni, Titik Nuryastuti, Tri Wibawa, Dwi AA Nugrahaningsih, Afiahayati, Siswanto, Kristy Iskandar, Nungki Anggorowati, Ika Trisnawati, Sumar Munawar Gani, Satria Maulana, Nur Rahmi Ananda, Riat El Khair, Yunika Puspadewi, Osman Sianipar, Umi Solekhah Intansari, Elizabeth Henny Herningtiyas, Ira Puspitawati, Nur Imma Fatimah Harahap, Dyah Ayu Puspitara |
| EPI_ISL_576130 | RSUP Dr. Sardjito | Genetics Working Group | Gunadi, Hendra Wibawa, Marcellus, Mohamad S. Hakim, Edwin W. Daniwijaya, Ludhang P. Rizki, Endah Supriyati, Eggi Arguni, Titik Nuryastuti, Tri Wibawa, Dwi AA Nugrahaningsih, Afiahayati, Siswanto, Kristy Iskandar, Nungki Anggorowati, Rina Triasih, Amalia |

| | | | |
|---|---|---|---|
| | | (Pokja Genetik) Faculty of Medicine, Public Health and Nursing Universitas Gadjah Mada (FK-KMK UGM); Disease Investigation Center Wates Ministry of Agriculture Indonesia; Department of Microbiology FK-KMK UGM; Laboratorium Diagnostik Yayasan Tahija World Mosquito Program (WMP) Yogyakarta Center for Tropical Medicine FK-KMK UGM; Integrated Research Center FK-KMK UGM; Department of Computer Science and Electronics FMIPA UGM | Cahya Dewi Satria, Titis Widowati, Elisabeth Siti Herini, Riat El Khair, Yunika Puspadewi, Osman Sianipar, Umi Solekhah Intansari, Elizabeth Henny Herningtiyas, Ira Puspitawati, Nur Imma Fatimah Harahap, Alvin S. Kalim, Susan S |
| EPI_ISL_576145 | RSA Universitas Gadjah Mada | Genetics Working Group (Pokja Genetik) Faculty of Medicine, Public Health and Nursing Universitas Gadjah Mada (FK-KMK UGM); Disease Investigation Center Wates Ministry of Agriculture Indonesia; Department of Microbiology FK-KMK UGM; Laboratorium Diagnostik Yayasan Tahija World Mosquito Program (WMP) Yogyakarta Center for Tropical Medicine FK-KMK UGM; Integrated Research Center FK-KMK UGM; Department of Computer Science and Electronics FMIPA UGM | Gunadi, Hendra Wibawa, Marcellus, Mohamad S. Hakim, Edwin W. Daniwijaya, Ludhang P. Rizki, Endah Supriyati, Eggi Arguni, Titik Nuryastuti, Tri Wibawa, Dwi AA Nugrahaningsih, Afiahayati, Siswanto, Kristy Iskandar, Nungki Anggorow |
| EPI_ISL_576383 | RSUD Budi Rahayu Kota Magelang | Genetics Working Group (Pokja Genetik) Faculty of Medicine, Public Health and Nursing Universitas Gadjah Mada (FK-KMK UGM); Disease Investigation Center Wates Ministry of Agriculture Indonesia; Department of Microbiology FK-KMK UGM; Laboratorium Diagnostik Yayasan Tahija World Mosquito Program (WMP) Yogyakarta Center for Tropical Medicine FK-KMK UGM; Integrated Research Center FK-KMK UGM; Department of Computer Science and Electronics FMIPA UGM; Balai Besar Teknik Kesehatan Lingkungan dan Pengendalian Penyakit (BBTKLPP) Yogyakarta | Gunadi, Hendra Wibawa, Marcellus, Mohamad S. Hakim, Edwin W. Daniwijaya, Ludhang P. Rizki, Endah Supriyati, Eggi Arguni, Titik Nuryastuti, Tri Wibawa, Dwi AA Nugrahaningsih, Afiahayati, Siswanto, Kristy Iskandar, Nungki Anggorowati, Irene, Indaryati, Havi Kenny Tedja, Aditya Rifqi Fauzi |
| EPI_ISL_576385 | Tirta Medical Center Angsana, Banjarmasin Kalsel | National Institute of Health Research and Development | Pawestri,HA;Subangkit;Puspa,KD;Nugraha,AA;Ikawati,HD;Pangesti,KNA;Soekarso,T;Paisal;Setiawaty,V |
| EPI_ISL_576386, EPI_ISL_576387 | National Institute of Health Research and Development | National Institute of Health Research and Development | Pawestri,HA;Subangkit;Puspa,KD;Nugraha,AA;Ikawati,HD;Pangesti, KNA;Soekarso,T;Susilarini,NK;Hariastuti,NI;Nikmah,UA;Mursinah;Febriyani,A;Herman,R;Susanti,N;Herna;Febriyanti,T; Nurhadi,M; Paisal;Ramadhany,R;Agustiningsih;Kurniawati,J;Kipuw,NL;Muna,F;Indalau,IL;Adam,K;Wibowo,HA;Rizki,A;Puspandari,N;Setiawaty,V |
| EPI_ISL_610155 | Puskesmas Cangkrep | Genetics Working Group (Pokja Genetik) Faculty of Medicine, Public Health and Nursing Universitas Gadjah Mada (FK-KMK UGM); Disease Investigation Center Wates Ministry of Agriculture Indonesia; Department of Microbiology FK-KMK UGM; Laboratorium Diagnostik Yayasan Tahija World Mosquito Program (WMP) Yogyakarta Center for Tropical Medicine FK-KMK UGM; Integrated Research | Gunadi, Hendra Wibawa, Marcellus, Mohamad S. Hakim, Edwin W. Daniwijaya, Ludhang P. Rizki, Endah Supriyati, Eggi Arguni, Titik Nuryastuti, Tri Wibawa, Dwi AA Nugrahaningsih, Afiahayati, Siswanto, Kristy Iskandar, Nungki Anggorowati, Irene, Indaryati, Havid Yuliyanti |

| | | | |
|---|---|---|---|
| | | Center FK-KMK UGM; Department of Computer Science and Electronics FMIPA UGM; Balai Besar Teknik Kesehatan Lingkungan dan Pengendalian Penyakit (BBTKLPP) Yogyakarta | |
| EPI_ISL_610158 | RS Panti Rini | Genetics Working Group (Pokja Genetik) Faculty of Medicine, Public Health and Nursing Universitas Gadjah Mada (FK-KMK UGM); Disease Investigation Center Wates Ministry of Agriculture Indonesia; Department of Microbiology FK-KMK UGM; Laboratorium Diagnostik Yayasan Tahija World Mosquito Program (WMP) Yogyakarta Center for Tropical Medicine FK-KMK UGM; Integrated Research Center FK-KMK UGM; Department of Computer Science and Electronics FMIPA UGM | Gunadi, Hendra Wibawa, Marcellus, Mohamad S. Hakim, Edwin W. Daniwijaya, Ludhang P. Rizki, Endah Supriyati, Eggi Arguni, Titik Nuryastuti, Tri Wibawa, Dwi AA Nugrahaningsih, Afiahayati, Siswanto, Kristy Iskandar, Nungki Anggorowati, V. Noegroho Isti I Puspitarani |
| EPI_ISL_610161 | RSUD Dr. Tjitrowardojo | Genetics Working Group (Pokja Genetik) Faculty of Medicine, Public Health and Nursing Universitas Gadjah Mada (FK-KMK UGM); Disease Investigation Center Wates Ministry of Agriculture Indonesia; Department of Microbiology FK-KMK UGM; Laboratorium Diagnostik Yayasan Tahija World Mosquito Program (WMP) Yogyakarta Center for Tropical Medicine FK-KMK UGM; Integrated Research Center FK-KMK UGM; Department of Computer Science and Electronics FMIPA UGM; Balai Besar Teknik Kesehatan Lingkungan dan Pengendalian Penyakit (BBTKLPP) Yogyakarta | Gunadi, Hendra Wibawa, Marcellus, Mohamad S. Hakim, Edwin W. Daniwijaya, Ludhang P. Rizki, Endah Supriyati, Eggi Arguni, Titik Nuryastuti, Tri Wibawa, Dwi AA Nugrahaningsih, Afiahayati, Siswanto, Kristy Iskandar, Nungki Anggorowati, Irene, Indaryati, Havi |
| EPI_ISL_610162 | RSUD Dr. Tjitrowardojo | Genetics Working Group (Pokja Genetik) Faculty of Medicine, Public Health and Nursing Universitas Gadjah Mada (FK-KMK UGM); Disease Investigation Center Wates Ministry of Agriculture Indonesia; Department of Microbiology FK-KMK UGM; Laboratorium Diagnostik Yayasan Tahija World Mosquito Program (WMP) Yogyakarta Center for Tropical Medicine FK-KMK UGM; Integrated Research Center FK-KMK UGM; Department of Computer Science and Electronics FMIPA UGM; Balai Besar Teknik Kesehatan Lingkungan dan Pengendalian Penyakit (BBTKLPP) Yogyakarta | Gunadi, Hendra Wibawa, Marcellus, Mohamad S. Hakim, Edwin W. Daniwijaya, Ludhang P. Rizki, Endah Supriyati, Eggi Arguni, Titik Nuryastuti, Tri Wibawa, Dwi AA Nugrahaningsih, Afiahayati, Siswanto, Kristy Iskandar, Nungki Anggorowati, Irene, Indaryati, Hav Kalim |
| EPI_ISL_632936 | Puskesmas Mlati 1 Sleman | Genetics Working Group (Pokja Genetik) Faculty of Medicine, Public Health and Nursing Universitas Gadjah Mada (FK-KMK UGM); Disease Investigation Center Wates Ministry of Agriculture Indonesia; Department of Microbiology | Gunadi, Hendra Wibawa, Marcellus, Mohamad S. Hakim, Edwin W. Daniwijaya, Ludhang P. Rizki, Endah Supriyati, Eggi Arguni, Titik Nuryastuti, Tri Wibawa, Dwi AA Nugrahaningsih, Afiahayati, Siswanto, Kristy Iskandar, Nungki Anggorowati, Ika Trisnawati, Sumar Munawar Gani, Satria Maulana, Nur Rahmi Ananda, Riat El Khair, Yunika Puspadewi, Osman Sianipar, Umi Solekhah Intansari, Elizabeth Henny Herningtiyas, Ira Puspitawati, Nur Imma Fatimah Harahap, Ernawati, Untung Wirav |

| | | | |
|---|---|---|---|
| | | FK-KMK UGM; Laboratorium Diagnostik Yayasan Tahija World Mosquito Program (WMP) Yogyakarta Center for Tropical Medicine FK-KMK UGM; Integrated Research Center FK-KMK UGM; Department of Computer Science and Electronics FMIPA UGM; RSUP Dr. Sardjito | |
| EPI_ISL_632937 | RSUD Saptosari Gunung Kidul | Genetics Working Group (Pokja Genetik) Faculty of Medicine, Public Health and Nursing Universitas Gadjah Mada (FK-KMK UGM); Disease Investigation Center Wates Ministry of Agriculture Indonesia; Department of Microbiology FK-KMK UGM; Laboratorium Diagnostik Yayasan Tahija World Mosquito Program (WMP) Yogyakarta Center for Tropical Medicine FK-KMK UGM; Integrated Research Center FK-KMK UGM; Department of Computer Science and Electronics FMIPA UGM; Balai Besar Teknik Kesehatan Lingkungan dan Pengendalian Penyakit (BBTKLPP) Yogyakarta | Gunadi, Hendra Wibawa, Marcellus, Mohamad S. Hakim, Edwin W. Daniwijaya, Ludhang P. Rizki, Endah Supriyati, Eggi Arguni, Titik Nuryastuti, Tri Wibawa, Dwi AA Nugrahaningsih, Afiahayati, Siswanto, Kristy Iskandar, Nungki Anggorowati, Irene, Indaryati, Havid Kenny Tedja |
| EPI_ISL_707692 | Medical Research Center, Faculty of Medicine, Syarif Hidayatullah State Islamic University Jakarta | Medical Research Center, Faculty of Medicine, Syarif Hidayatullah State Islamic University Jakarta | Chris Adhiyanto, Laifa Hendarmin, Erike Suwarsono, Zeti Harriyati, Rini Puspitaningrum, Ferania Mela, Dennis Nurjadi |
| EPI_ISL_707696 | Medical Research Center, Faculty of Medicine, Syarif Hidayatullah State Islamic University Jakarta | Medical Research Center, Faculty of Medicine, Syarif Hidayatullah State Islamic University Jakarta | Laifa Hendarmin, Chris Adhiyanto, Erike Suwarsono, Zeti Harriyati, Rini Puspitaningrum, Ferania Mela, Dennis Nurjadi |
| EPI_ISL_707778 | Medical Research Center, Faculty of Medicine, Syarif Hidayatullah State Islamic University Jakarta | Medical Research Center, Faculty of Medicine, Syarif Hidayatullah State Islamic University Jakarta | Erike A Suwarsono, Chris Adhiyanto, Laifa Hendarmin, Zeti Harriyati, Rini Puspitaningrum, Ferania Mela, Dennis Nurjadi |
| EPI_ISL_707890 | Medical Research Center, Faculty of Medicine, Syarif Hidayatullah State Islamic University Jakarta | Medical Research Center, Faculty of Medicine, Syarif Hidayatullah State Islamic University Jakarta | Zeti Harriyati, Chris Adhiyanto, Laifa Hendarmin, Erike A Suwarsono, Dennis Nurjadi |
| EPI_ISL_708652 | Medical Research Center, Faculty of Medicine, Syarif Hidayatullah State Islamic University Jakarta | Medical Research Center, Faculty of Medicine, Syarif Hidayatullah State Islamic University Jakarta | Chris Adhiyanto, Laifa Hendarmin, Erike A Suwarsono, Zeti Harriyati, Endah Wulandari, Flori R Sari, Hari Hendarto |
| EPI_ISL_745033 | West Java Health Laboratory | West Java Health Laboratory | Azzania Fibriani, Ema Rahmawati, Ryan Bayusantika Ristandi, Rifky Waluyajati Rachman, Cut Nur Cinthia Alamanda, Isak Solihin, Rini Robiani, Miftahul Faridl, Karimatu Khoirunnisa |
| EPI_ISL_747194 | Clinical Microbiology Laboratory, Faculty of Medicine, Universitas Indonesia | Clinical Microbiology Laboratory, Faculty of Medicine, Universitas Indonesia | Fera Ibrahim, Augustine Natasha, Andi Yasmon, Pratiwi Sudarmono, Yulia Rosa Saharman, Anis Karuniawati, Fithriyah |
| EPI_ISL_747233, EPI_ISL_747234 | National Institute of Health Research and Development | National Institute of Health Research and Development | Subangkit; Indrasari,ND; Wulandari,D; Monika,M; Pawestri,HA; Puspa,KD; Nugraha,AA; Ikawati,HD; Pangesti,KNA; Soekarso,T; Paisal; Setiawaty,Vivi |
| EPI_ISL_747235 | Sukabumi Health Laboratory | West Java Health Laboratory; School of Life Sciences and Technology, Institut Teknologi Bandung | Azzania Fibriani, Ema Rahmawati, Ryan Bayusantika Ristandi, Rifky Waluyajati Rachman, Cut Nur Cinthia Alamanda, Isak Solihin, Rini Robiani, Miftahul Faridl, Karimatu Khoirunnisa |
| EPI_ISL_747236 | Sukabumi Public Health | West Java Health Laboratory; School of Life | Azzania Fibriani, Ema Rahmawati, Ryan Bayusantika Ristandi, Rifky Waluyajati Rachman, Cut Nur Cinthia Alamanda, Isak Solihin, Rini Robiani, Miftahul Faridl, Karimatu Khoirunnisa |

| | | | |
|---|---|---|---|
| | | Sciences and Technology, Institut Teknologi Bandung | |
| EPI_ISL_747237 | National Institute of Health Research and Development | National Institute of Health Research and Development | Subangkit,Nafiandi,Estaria,V;Pawestri,HA;Puspa,KD;Nugraha,AA;Ikawati,HD;Pangesti,KNA;Soekarso,T;Paisal;Puspandari,N;Setiawaty,V |
| EPI_ISL_747238 | Bogor Public Health | West Java Health Laboratory; School of Life Sciences and Technology, Institut Teknologi Bandung | Azzania Fibriani, Ema Rahmawati, Ryan Bayusantika Ristandi, Rifky Waluyajati Rachman, Cut Nur Cinthia Alamanda, Isak Solihin, Rini Robiani, Miftahul Faridl, Karimatu Khoirunnisa |
| EPI_ISL_747239 | Kuningan Publlic Health Office | West Java Health Laboratory; School of Life Sciences and Technology, Institut Teknologi Bandung | Azzania Fibriani, Ema Rahmawati, Ryan Bayusantika Ristandi, Rifky Waluyajati Rachman, Cut Nur Cinthia Alamanda, Isak Solihin, Rini Robiani, Miftahul Faridl, Karimatu Khoirunnisa |
| EPI_ISL_747240 | Al Islam Hospital | West Java Health Laboratory; School of Life Sciences and Technology, Institut Teknologi Bandung | Azzania Fibriani, Ema Rahmawati, Ryan Bayusantika Ristandi, Rifky Waluyajati Rachman, Cut Nur Cinthia Alamanda, Isak Solihin, Rini Robiani, Miftahul Faridl, Karimatu Khoirunnisa |
| EPI_ISL_747241 | Subang Public Health Office | West Java Health Laboratory; School of Life Sciences and Technology, Institut Teknologi Bandung | Azzania Fibriani, Ema Rahmawati, Ryan Bayusantika Ristandi, Rifky Waluyajati Rachman, Cut Nur Cinthia Alamanda, Isak Solihin, Rini Robiani, Miftahul Faridl, Karimatu Khoirunnisa |
| EPI_ISL_753698 | Balai Besar Veteriner Maros, Sulawesi Selatan | National Institute of Health Research and Development | Puspa,KD;Subangkit;Pawestri,HA;Nugraha,AA;Ikawati,HD;Pangesti,KNA;Soekarso,T;Puspandari,N;Muflihanah;Mangidi,R;Setiawaty,V |
| EPI_ISL_753699 | RSUD Ulin Banjarmasin Kalimantan Selatan | National Institute of Health Research and Development | Subangkit;Pawestri,HA;Puspa,KD;Nugraha,AA;Ikawati,HD;Pangesti,KNA;Soekarso,T;Puspandari,N;Pasaribu,M;Setiawaty,V |
| EPI_ISL_754239 | Santosa Hospital Bandung Central | "School of Life Sciences and Technology & School of Pharmacy-Institut Teknologi Bandung; Molecular Genetics Laboratory-Faculty of Medicine-Universitas Padjadjaran; Laboratorium Kesehatan Provinsi Jawa Barat" | Marselina Irasonia Tan, Yunia Sribudiani, Catur Riani, Azzania Fibriani, Husna Nugrahapraja, Tarwadi, Ema Rahmawati, Savira Ekawardhani, Hesti Lina Wiraswati, Ryan Bayusantika Ristandi, Rifky Waluyajati Rachman, Cut Nur Cinthia Alamanda, Lia Faridah, M<br>Solistia Wirawan, Agung Eru Wibowo, Irvan Faizal |
| EPI_ISL_754241 | Dinkes Tasikmalaya | "School of Life Sciences and Technology & School of Pharmacy-Institut Teknologi Bandung; Molecular Genetics Laboratory-Faculty of Medicine-Universitas Padjadjaran; Laboratorium Kesehatan Provinsi Jawa Barat" | Husna Nugrahapraja, Marselina Irasonia Tan, Yunia Sribudiani, Catur Riani, Azzania Fibriani, Tarwadi, Ema Rahmawati, Savira Ekawardhani, Hesti Lina Wiraswati, Ryan Bayusantika Ristandi, Rifky Waluyajati Rachman, Cut Nur Cinthia Alamanda, Lia Faridah, M<br>Solistia Wirawan, Agung Eru Wibowo, Irvan Faizal |
| EPI_ISL_756376 | National Institute of Health Research and Development | National Institute of Health Research and Development | Ramadhany,R; Rukminiati,Y; Agustiningsih; Adam,K; Wibowo,HA; Pawestri,HA; Subangkit;Puspa,KD; Nugraha,AA; Ikawati,HD; Pangesti,KNA; Soekarso,T; Susilarini,NK; Hariastuti,NI; Nikmah,UA;Mursinah; Febriyani,A; Herman,R; Susanti,N;Herna; Febriyanti,T<br>Rizki,A; Puspandari,N; Setiawaty,Vivi |
| EPI_ISL_756401 | National Institute of Health Research and Development | National Institute of Health Research and Development | Adam,K; Wibowo,HA; Ramadhany,R; Rukminiati,Y; Agustiningsih; Pawestri,HA; Subangkit; Puspa,KD; Nugraha,AA; Ikawati,HD; Pangesti,KNA; Soekarso,T; Susilarini,NK; Hariastuti,NI; Nikmah,UA; Mursinah; Febriyani,A; Herman,R; Susanti,N; Herna; Febriyanti,<br>Rizki,A; Puspandari,N; Setiawaty,Vivi |
| EPI_ISL_757287 | National Institute of Health Research and Development | National Institute of Health Research and Development | Rukminiati,Y;Agustiningsih;Adam,K;Wibowo,HA;Ramadhany,R;Pawestri,HA;Subangkit;Puspa,KD;Nugraha,AA;Ikawati,HD;Pangesti,KNA;Soekarso,T;Susilarini,NK;Hariastuti,NI;Nikmah,UA;Mursinah;Febriyani,A;Herman,R;Susan<br>Nurhadi,M;Kurniawati,J;Kipuw,NL;Muna,F;Indalao,IL;Rizki,A;Puspandari,N;Setiawaty,V |
| EPI_ISL_759701 | RS UNS, Surakarta | UNS/RS-UNS, Faculty of Medicine | Betty Suryawati, Yulia Sari, Hartono, Maryani, Afif Avicenna Gufron, HanaHana Apsari Pawestri, Kartika Dewi Puspa, Hartanti Dian Ika, Arie Ardiansyah Nugraha, Vivi Setiawaty. |
| EPI_ISL_759702 | RS UNS Surakarta | Universitas Sebelas Maret (UNS), Rumah sakit Universitas Sebelas Maret (RS-UNS) Surakarta, National Institute of Health Research and Development, Indonesian Ministry of Health | Betty Suryawati, Yulia sari, hartono, maryani, Afif Avicena Gufron, Hana Apsari Pawestri, Kartika, Dewi Puspa, Hartanti Dian Ika, Arie Ardiansyah Nugraha, Vivi Setiawaty |
| EPI_ISL_759703 | RSUD Bung Karno | Universitas Sebelas Maret (UNS), Rumah sakit Universitas Sebelas Maret (RS-UNS) Surakarta, National Institute of Health Research and Development, Indonesian Ministry of Health | Betty Suryawati, Yulia sari, hartono, maryani, Afif Avicena Gufron, Hana Apsari Pawestri, Kartika, Dewi Puspa, Hartanti Dian Ika, Arie Ardiansyah Nugraha, Vivi Setiawaty |
| EPI_ISL_759705 | RSUD Bung Karno, Surakarta | Universitas Sebelas Maret (UNS); Rumah Sakit Universitas Sebelas Maret (RS-UNS), Surakarta; National Institute of Health | Betty Suryawati, Yulia Sari, Hartono, Maryani, Afif Avicena Gufron, Hana Apsari Pawestri, Kartika, Dewi Puspa, Hartanti Dian Ika, Arie Ardiansyah Nugraha, Vivi Setiawaty. |

| | | | |
|---|---|---|---|
| | | Research and Development, Indonesian Ministry of Health , Jakarta). | |
| EPI_ISL_759842 | Rumah Sakit Universitas Sebelas Maret (RS-UNS) | Universitas Sebelas Maret (UNS); Rumah Sakit Universitas Sebelas Maret (RS-UNS), Surakarta; National Institute of Health Research and Development, Indonesian Ministry of Health, Jakarta. | Betty Suryawati, Yulia Sari, Hartono, Maryani, Afif Avicena Ghufron, Hana Apsari Pawestri, Kartika, Dewi Puspa, Hartanti Dian Ika, Arie Ardiansyah Nugraha, Vivi Setiawaty. |
| EPI_ISL_759955 | Primasatya Husada Citra Hospital | Institute of Tropical Disease, Universitas Airlangga | Aldise M Nastri, Jezzy R Dewantari, Rima R Prasetya, Krisnoadi Rahardjo, Pudji Djanuartono, Gatot Soegiarto, Laksmi Wulandari, Resti Yudhawati, Soetjipto, Yasuko Mori, Maria I Lusida, Kazufumi Shimizu |
| EPI_ISL_759956 | Husada Utama Hospital | Institute of Tropical Disease, Universitas Airlangga | Jezzy R Dewantari, Rima R Prasetya, Krisnoadi Rahardjo, Aldise M Nastri, Didi Dewanto, Gatot Soegiarto, Laksmi Wulandari, Resti Yudhawati, Soetjipto, Yasuko Mori, Maria I Lusida, Kazufumi Shimizu |
| EPI_ISL_759957 | Siti Khodijah Hospital | Institute of Tropical Disease, Universitas Airlangga | Rima R Prasetya, Krisnoadi Rahardjo, Aldise M Nastri, Jezzy R Dewantari, Muhammad Hamdan, Gatot Soegiarto, Laksmi Wulandari, Resti Yudhawati, Soetjipto, Yasuko Mori, Maria I Lusida, Kazufumi Shimi |
| EPI_ISL_759958 | Universitas Airlangga Hospital | Institute of Tropical Disease, Universitas Airlangga | Maria I Lusida, Krisnoadi Rahardjo, Aldise M Nastri, Jezzy R Dewantari, Rima R Prasetya, Nasronudin, Gatot Soegiarto, Laksmi Wulandari, Resti Yudhawati, Soetjipto, Yasuko Mori, Kazufumi Shimizu |
| EPI_ISL_759959 | Premier Hospital Surabaya | Institute of Tropical Disease, Universitas Airlangga | Krisnoadi Rahardjo, Aldise M Nastri, Jezzy R Dewantari, Rima R Prasetya, Hartono Tanto, Gatot Soegiarto, Laksmi Wulandari, Resti Yudhawati, Soetjipto, Yasuko Mori, Maria I Lusida, Kazufumi Shimizu |
| EPI_ISL_759960 | Surabaya Public Health Office | Institute of Tropical Disease, Universitas Airlangga | Aldise M Nastri, Jezzy R Dewantari, Rima R Prasetya, Krisnoadi Rahardjo, Febria Rachmanita, Gatot Soegiarto, Laksmi Wulandari, Resti Yudhawati, Soetjipto, Yasuko Mori, Maria I Lusida, Kazufumi Shimiz |
| EPI_ISL_759961 | Universitas Airlangga Hospital | Institute of Tropical Disease, Universitas Airlangga | Jezzy R Dewantari, Rima R Prasetya, Krisnoadi Rahardjo, Aldise M Nastri, Nasronudin, Gatot Soegiarto, Laksmi Wulandari, Resti Yudhawati, Soetjipto, Yasuko Mori, Maria I Lusida, Kazufumi Shimizu |
| EPI_ISL_759962 | RSUD Dr. Soetomo | Institute of Tropical Disease, Universitas Airlangga | Krisnoadi Rahardjo, Aldise M Nastri, Jezzy R Dewantari, Rima R Prasetya, Joni Wahyuhadi, Gatot Soegiarto, Laksmi Wulandari, Resti Yudhawati, Yasuko Mori, Soetjipto, Kazufumi Shimizu ,Maria I Lusida |
| EPI_ISL_759963 | RSUD Sidoarjo | Institute of Tropical Disease, Universitas Airlangga | Aldise M Nastri, Jezzy R Dewantari, Rima R Prasetya, Krisnoadi Rahardjo, Atok Irawan, Gatot Soegiarto, Laksmi Wulandari, Resti Yudhawati, Yasuko Mori, Soetjipto, Kazufumi Shimizu ,Maria I Lusida |
| EPI_ISL_759964 | Mitra Keluarga Hospital Waru | Institute of Tropical Disease, Universitas Airlangga | Jezzy R Dewantari, Rima R Prasetya, Krisnoadi Rahardjo, Aldise M Nastri, Christina Dian Anggraeni, Gatot Soegiarto, Laksmi Wulandari, Resti Yudhawati, Yasuko Mori, Soetjipto, Kazufumi Shimizu ,Maria I Lu |
| EPI_ISL_759965 | RSUD Dr. Soetomo | Institute of Tropical Disease, Universitas Airlangga | Rima R Prasetya, Krisnoadi Rahardjo, Aldise M Nastri, Jezzy R Dewantari, Joni Wahyuhadi, Gatot Soegiarto, Laksmi Wulandari, Resti Yudhawati, Yasuko Mori, Soetjipto, Kazufumi Shimizu ,Maria I Lusida |
| EPI_ISL_759966 | RSUD Sidoarjo | Institute of Tropical Disease, Universitas Airlangga | Krisnoadi Rahardjo, Aldise M Nastri, Jezzy R Dewantari, Rima R Prasetya, Atok Irawan, Gatot Soegiarto, Laksmi Wulandari, Resti Yudhawati, Yasuko Mori, Soetjipto, Kazufumi Shimizu ,Maria I Lusida |
| EPI_ISL_759967 | RSUD Dr. Iskak Tulungagung | Institute of Tropical Disease, Universitas Airlangga | Rima R Prasetya, Krisnoadi Rahardjo, Aldise M Nastri, Jezzy R Dewantari, Supriyanto Dharmoredjo, Gatot Soegiarto, Laksmi Wulandari, Resti Yudhawati, Soetjipto, Yasuko Mori, Maria I Lusida, Kazufumi Shir |
| EPI_ISL_759968 | RSI Darus Syifa' | Institute of Tropical Disease, Universitas Airlangga | Maria I Lusida, Krisnoadi Rahardjo, Aldise M Nastri, Jezzy R Dewantari, Rima R Prasetya, HM Faiz, Gatot Soegiarto, Laksmi Wulandari, Resti Yudhawati, Soetjipto, Yasuko Mori, Kazufumi Shimizu |
| EPI_ISL_763063 | RS Panti Waluyo, Surakarta | Universitas Sebelas Maret (UNS); Rumah Sakit UNS (RS-UNS); National Institute of Health Research and Development, Indonesian Ministry of Health. | Betty Suryawati, Yulia Sari, Hartono, Maryani, Afif A Ghufron, Revi G H Novika, Hana A Pawestri, Kartika, Dewi Puspa, Hartanti Dian Ika, Arie A Nugraha, Vivi Setiawaty. |
| EPI_ISL_763064 | RS Kasih Ibu, Surakarta | Universitas Sebelas Maret (UNS); Rumah Sakit UNS (RS-UNS); National Institute of Health Research and Development, Indonesian Ministry of Health. | Betty Suryawati, Yulia Sari, Hartono, Maryani, Afif A Ghufron, Revi G H Novika, Hana A Pawestri, Kartika, Dewi Puspa, Hartanti Dian Ika, Arie A Nugraha, Vivi Setiawaty. |
| EPI_ISL_763066 | Puskesmas Grogol, Sukoharjo | Universitas Sebelas Maret (UNS); Rumah Sakit UNS (RS-UNS); National Institute of Health Research and Development, Indonesian Ministry of Health. | Yulia Sari, Hartono, Maryani, Betty Suryawati, Afif A Ghufron, Revi G H Novika, Hana A Pawestri, Kartika, Dewi Puspa, Hartanti Dian Ika, Arie A Nugraha, Vivi Setiawaty. |
| EPI_ISL_763068 | RSUD Kota Surakarta | Universitas Sebelas Maret (UNS); Rumah Sakit UNS (RS-UNS); National Institute of Health Research | Maryani, Yulia Sari, Hartono, Afif A Ghufron, Revi G H Novika, Hana A Pawestri, Kartika, Dewi Puspa, Hartanti Dian Ika, Arie A Nugraha, Vivi Setiawaty, Betty Suryawati. |

| | | | |
|---|---|---|---|
| | | and Development, Indonesian Ministry of Health. | |
| EPI_ISL_763070 | Puskesmas Jambu Kulon, Klaten | Universitas Sebelas Maret (UNS); Rumah Sakit UNS (RS-UNS); National Institute of Health Research and Development, Indonesian Ministry of Health. | Yulia Sari, Hartono, Maryani, Afif A Ghufron, Hana A Pawestri, Kartika, Dewi Puspa, Hartanti Dian Ika, Arie A Nugraha, Vivi Setiawaty, Betty Suryawati. |
| EPI_ISL_766032 | RSUP Dr. Sardjito | Eijkman Institute for Molecular Biology, Ministry of Research and Technology/National Agency for Research and Innovation | Frilasita A Yudhaputri, Edison Johar, Hidayat Trimarsanto, David H Muljono, Safarina G Malik, Khin Saw Myint, Amin Soebandrio |
| EPI_ISL_766033 | RSUD Prof. DR. W. Z. Johannes Kupang | Eijkman Institute for Molecular Biology, Ministry of Research and Technology/National Agency for Research and Innovation | Frilasita A Yudhaputri, Edison Johar, Hidayat Trimarsanto, David H Muljono, Safarina G Malik, Khin Saw Myint, Amin Soebandrio |
| EPI_ISL_766034 | RS Mitra Keluarga Bintaro | Eijkman Institute for Molecular Biology, Ministry of Research and Technology/National Agency for Research and Innovation | Frilasita A Yudhaputri, Edison Johar, Hidayat Trimarsanto, David H Muljono, Safarina G Malik, Khin Saw Myint, Amin Soebandrio |
| EPI_ISL_766035 | RS Bhayangkara Brimob Kelapa Dua | Eijkman Institute for Molecular Biology, Ministry of Research and Technology/National Agency for Research and Innovation | Frilasita A Yudhaputri, Edison Johar, Hidayat Trimarsanto, David H Muljono, Safarina G Malik, Khin Saw Myint, Amin Soebandrio |
| EPI_ISL_766036 | RS Mitra Keluarga Bintaro | Eijkman Institute for Molecular Biology, Ministry of Research and Technology/National Agency for Research and Innovation | Frilasita A Yudhaputri, Edison Johar, Hidayat Trimarsanto, David H Muljono, Safarina G Malik, Khin Saw Myint, Amin Soebandrio |
| EPI_ISL_766037 | Puskesmas Palmerah | Eijkman Institute for Molecular Biology, Ministry of Research and Technology/National Agency for Research and Innovation | Frilasita A Yudhaputri, Edison Johar, Hidayat Trimarsanto, David H Muljono, Safarina G Malik, Khin Saw Myint, Amin Soebandrio |
| EPI_ISL_766038 | RSJPD Harapan Kita | Eijkman Institute for Molecular Biology, Ministry of Research and Technology/National Agency for Research and Innovation | Frilasita A Yudhaputri, Edison Johar, Hidayat Trimarsanto, David H Muljono, Safarina G Malik, Khin Saw Myint, Amin Soebandrio |
| EPI_ISL_766039 | RS Bhayangkara Tk. III | Eijkman Institute for Molecular Biology, Ministry of Research and Technology/National Agency for Research and Innovation | Frilasita A Yudhaputri, Edison Johar, Hidayat Trimarsanto, David H Muljono, Safarina G Malik, Khin Saw Myint, Amin Soebandrio |
| EPI_ISL_766040 | RS Tingkat II Moh. Ridwan Meuraksa | Eijkman Institute for Molecular Biology, Ministry of Research and Technology/National Agency for Research and Innovation | Frilasita A Yudhaputri, Edison Johar, Hidayat Trimarsanto, David H Muljono, Safarina G Malik, Khin Saw Myint, Amin Soebandrio |
| EPI_ISL_766041 | RS Polri | Eijkman Institute for Molecular Biology, Ministry of Research and Technology/National Agency for Research and Innovation | Frilasita A Yudhaputri, Edison Johar, Hidayat Trimarsanto, David H Muljono, Safarina G Malik, Khin Saw Myint, Amin Soebandrio |
| EPI_ISL_766042 | RS Sari Asih Ciledug | Eijkman Institute for Molecular Biology, Ministry of Research and Technology/National Agency for Research and Innovation | Frilasita A Yudhaputri, Edison Johar, Hidayat Trimarsanto, David H Muljono, Safarina G Malik, Khin Saw Myint, Amin Soebandrio |
| EPI_ISL_766043 | RS Pelni | Eijkman Institute for Molecular Biology, Ministry of Research and | Frilasita A Yudhaputri, Edison Johar, Hidayat Trimarsanto, David H Muljono, Safarina G Malik, Khin Saw Myint, Amin Soebandrio |

| | | | |
|---|---|---|---|
| | | Technology/National Agency for Research and Innovation | |
| EPI_ISL_766044 | RSUP Fatmawati | Eijkman Institute for Molecular Biology, Ministry of Research and Technology/National Agency for Research and Innovation | Frilasita A Yudhaputri, Edison Johar, Hidayat Trimarsanto, David H Muljono, Safarina G Malik, Khin Saw Myint, Amin Soebandrio |
| EPI_ISL_766045 | RS Pelni | Eijkman Institute for Molecular Biology, Ministry of Research and Technology/National Agency for Research and Innovation | Frilasita A Yudhaputri, Edison Johar, Hidayat Trimarsanto, David H Muljono, Safarina G Malik, Khin Saw Myint, Amin Soebandrio |
| EPI_ISL_766046 | RSUP Fatmawati | Eijkman Institute for Molecular Biology, Ministry of Research and Technology/National Agency for Research and Innovation | Frilasita A Yudhaputri, Edison Johar, Hidayat Trimarsanto, David H Muljono, Safarina G Malik, Khin Saw Myint, Amin Soebandrio |
| EPI_ISL_766047 | RS Kramat 128 | Eijkman Institute for Molecular Biology, Ministry of Research and Technology/National Agency for Research and Innovation | Frilasita A Yudhaputri, Edison Johar, Hidayat Trimarsanto, David H Muljono, Safarina G Malik, Khin Saw Myint, Amin Soebandrio |
| EPI_ISL_766048 | RS Bhayangkara Tk. III | Eijkman Institute for Molecular Biology, Ministry of Research and Technology/National Agency for Research and Innovation | Frilasita A Yudhaputri, Edison Johar, Hidayat Trimarsanto, David H Muljono, Safarina G Malik, Khin Saw Myint, Amin Soebandrio |
| EPI_ISL_775594 | Balitvet Lampung | National Institute of Health Research and Development | Nugraha,AA;Subangkit;Pawestri,HA;Ikawati,HD;Puspa,KD;Saswiyanti,E;Srihanto,EA;Pangesti,KNA;Soekarso,T;Puspandari,N;Setiawaty,V |
| EPI_ISL_775596 | Lab. Biologi Molekular FKIK Unwar, Bali | National Institute of Health Research and Development | Ikawati,HD;Subangkit;Pawestri,HA;Nugraha,AA;Puspa,KD;Dewi,SR;Pangesti,KNA;Soekarso,T;Puspandari,N;Setiawaty,V |
| EPI_ISL_775597 | RS Pertamina Sorong | National Institute of Health Research and Development | Puspa,KD;Subangkit;Pawestri,HA;Ikawati,HD;Nugraha,AA;Yosepin;Pangesti,KNA;Soekarso,T;Puspandari,N;Setiawaty,V |
| EPI_ISL_775598 | RSUD Kota Mataram | National Institute of Health Research and Development | Puspa,KD;Subangkit;Pawestri,HA;Ikawati,HD;Nugraha,AA;Sari,SK;Pangesti,KNA;Soekarso,T;Puspandari,N;Setiawaty,V |
| EPI_ISL_775599 | Balai Besar Veteriner Maros | National Institute of Health Research and Development | Subangkit;Pawestri,HA;Ikawati,HD;Nugraha,AA;Puspa,KD;Muflihanah;Mangidi,R;Pangesti,KNA;Soekarso,T;Puspandari,N;Setiawaty,V |
| EPI_ISL_779383, EPI_ISL_779384, EPI_ISL_779385, EPI_ISL_779386, EPI_ISL_779387, EPI_ISL_779388, EPI_ISL_779389, EPI_ISL_779390, EPI_ISL_779391, EPI_ISL_779392, EPI_ISL_779393, EPI_ISL_779394, EPI_ISL_779395 | | | |
| see above | Siloam Hospital Lippo Village | Mochtar Riady Institute for Nanotechnology-Universitas Pelita Harapan | Aksar C Lages, David Rustandi, Ivet M Suriapranata, Rinaldy Kusuma, Febi Andriani, Tri Shinta Kurniasih, Riska N Taufik, Young O Larasati, Rury M Wahyuni, Irawan Yusuf |
| EPI_ISL_791977 | Laboratorium Mikrobiologi Samarinda | National Institute of Health Research and Development | Nugraha,AA;Subangkit;Pawestri,HA;Ikawati,HD;Puspa,KD;Tandirogang,N;Yadi;Pangesti,KNA;Soekarso,T;Puspandari,N;Setiawaty,V |
| EPI_ISL_791978 | Balai Labkes Lampung | National Institute of Health Research and Development | Nugraha,AA;Subangkit;Pawestri,HA;Ikawati,HD;Puspa,KD;Yurina,L;Pangesti,KNA;Soekarso,T;Puspandari,N;Setiawaty,V |
| EPI_ISL_791979 | RSUD Curup Bengkulu | National Institute of Health Research and Development | Nugraha,AA;Subangkit;Pawestri,HA;Ikawati,HD;Puspa,KD;Mardiani;Pangesti,KNA;Soekarso,T;Puspandari,N;Setiawaty,V |
| EPI_ISL_791980 | RSUD Blambangan Banyuwangi | National Institute of Health Research and Development | Ikawati,HD;Subangkit;Pawestri,HA;Nugraha,AA;Puspa,KD;Noor,RI;Pangesti,KNA;Soekarso,T;Puspandari,N;Setiawaty,V |
| EPI_ISL_791981 | Balai Litbang Aceh | National Institute of Health Research and Development | Ikawati,HD;Subangkit;Pawestri,HA;Nugraha,AA;Puspa,KD;Ichwansyah,F;Fitria,E;Pangesti,KNA;Soekarso,T;Puspandari,N;Setiawaty,V |
| EPI_ISL_791982 | RSUP DR KARIADI Semarang | National Institute of Health Research and Development | Ikawati,HD;Subangkit;Pawestri,HA;Nugraha,AA;Puspa,KD;Puspitasari,I;Pangesti,KNA;Soekarso,T;Puspandari,N;Setiawaty,V |
| EPI_ISL_791983 | RSUD dr. Kanujoso Djatiwibowo Balikpapan | National Institute of Health Research and Development | Subangkit;Pawestri,HA;Ikawati,HD;Nugraha,AA;Puspa,KD;Ivanna;Pangesti,KNA;Soekarso,T;Puspandari,N;Setiawaty,V |
| EPI_ISL_791985 | RS Santa Maria Pekanbaru | National Institute of Health Research and Development | Puspa,KD;Subangkit;Pawestri,HA;Ikawati,HD;Nugraha,AA;Fridayenti;Pangesti,KNA;Soekarso,T;Puspandari,N;Setiawaty,V |
| EPI_ISL_791986 | RSUD Morotai Maluku Utara | National Institute of Health Research and Development | Subangkit;Pawestri,HA;Ikawati,HD;Nugraha,AA;Puspa,KD;Denggo,SD;Pangesti,KNA;Soekarso,T;Puspandari,N;Setiawaty,V |
| EPI_ISL_791987 | RSUD dr. Loekmono Hadi Kudus | National Institute of Health Research and Development | Puspa,KD;Subangkit;Pawestri,HA;Ikawati,HD;Nugraha,AA;Khoiriyah,S;Pangesti,KNA;Soekarso,T;Puspandari,N;Setiawaty,V |
| EPI_ISL_791988 | Balai Besar POM Semarang | National Institute of Health Research and Development | Subangkit;Pawestri,HA;Ikawati,HD;Nugraha,AA;Puspa,KD;Aryanti;Pangesti,KNA;Soekarso,T;Puspandari,N;Setiawaty,V |
| EPI_ISL_791989 | RSUD Waluyo Jati Kota Kraksaan, | National Institute of Health Research and Development | Subangkit;Pawestri,HA;Ikawati,HD;Nugraha,AA;Puspa,KD;Muhashonah,I;Pangesti,KNA;Soekarso,T;Puspandari,N;Setiawaty,V |

| | | | |
|---|---|---|---|
| | Jawa Timur | | |
| EPI_ISL_803876 | RSUD Tarakan Kalimantan Utara | National Institute of Health Research and Development | Subangkit;Pawestri,HA;Ikawati,HD;Nugraha,AA;Puspa,KD;Sriwedari,K;Sindunata,R;Pangesti,KNA;Soekarso,T;Puspandari,N;Setiawaty,V |
| EPI_ISL_833039 | RS TK II 02.05.01 dr. AK Gani Palembang | National Institute of Health Research and Development | Subangkit;Pawestri,HA;Ikawati,HD;Nugraha,AA;Puspa,KD;Pangesti,KNA;Soekarso,T;Tantula,A;Wiradharma,RP;Puspandari,N;Setiawaty,V |
| EPI_ISL_833040 | RSUD Dr. Abdul Aziz Singkawang | National Institute of Health Research and Development | Subangkit;Pawestri,HA;Ikawati,HD;Nugraha,AA;Puspa,KD;Pangesti,KNA;Soekarso,T;Junus,HN;Puspandari,N;Setiawaty,V |
| EPI_ISL_833130 | National Institute of Health Research and Development | National Institute of Health Research and Development | Agustiningsih;Adam,K;Wibowo,HA;Ramadhany,R;Rukminiati,Y;Pawestri,HA;Subangkit;Puspa,KD;Nugraha,AA;Ikawati,HD;Pangesti,KNA;Soekarso,T;Susilarini,NK;Hariastuti,NI;Nikmah,UA;Mursinah;Febriyani,A;Herman,R;Susanti,N;Herna;Febriyanti,T;Kurniawati,J;[illegible] |
| EPI_ISL_833388 | Klinik Apotek Dein, Jakarta, Indonesia | Biosafety Level-3 Laboratory, Indonesian Institute of Sciences (LIPI) | Syam Budi Iryanto, Andri Wardiana, Anggia Prasetyoputri, Isa Nuryana, Ade Andriani, Anik Budhi Dharmayanthi, Ahmad Fathoni, Dian Fitria Agustiyanti, Asrul M. Fuad, Ratih Asmana Ningrum |
| EPI_ISL_833389 | Klinik Apotek Dein, Jakarta, Indonesia | Biosafety Level-3 Laboratory, Indonesian Institute of Sciences (LIPI) | Syam Budi Iryanto, Andri Wardiana, Anggia Prasetyoputri, Isa Nuryana, Ade Andriani, Anik Budhi Dharmayanthi, Ahmad Fathoni, Popi Hadi Wisnuwardhani, Sri Swasthikawati, Ratih Asmana Ningrum |
| EPI_ISL_833390 | Klinik Apotek Dein, Jakarta, Indonesia | Biosafety Level-3 Laboratory, Indonesian Institute of Sciences (LIPI) | Andri Wardiana, Anggia Prasetyoputri, Isa Nuryana, Ade Andriani, Anik Budhi Dharmayanthi, Syam Budi Iryanto,Ahmad Fathoni, Fahrurrozy, Alfi Taufik Fathurahman, Ratih Asmana Ningrum |
| EPI_ISL_833391 | Klinik Apotek Dein, Jakarta, Indonesia | Biosafety Level-3 Laboratory, Indonesian Institute of Sciences (LIPI) | Andri Wardiana, Anggia Prasetyoputri, Isa Nuryana, Ade Andriani, Anik Budhi Dharmayanthi, Syam Budi Iryanto, Ahmad Fathoni, Herman Irawan, Dadang Supriatna, Ratih Asmana Ningrum |
| EPI_ISL_833392 | RSU Sayang Bunda, Makasar,Indonesia | Biosafety Level-3 Laboratory, Indonesian Institute of Sciences (LIPI) | Anggia Prasetyoputri, Isa Nuryana, Ade Andriani, Anik Budhi Dharmayanthi, Syam Budi Iryanto, Andri Wardiana, Ahmad Fathoni, Dini Nurdiani, Wahyuni, Ratih Asmana Ningrum |
| EPI_ISL_833393 | RS MMC Jakarta, Indonesia | Biosafety Level-3 Laboratory, Indonesian Institute of Sciences (LIPI) | Anggia Prasetyoputri, Isa Nuryana, Ade Andriani, Anik Budhi Dharmayanthi, Syam Budi Iryanto, Andri Wardiana, Ahmad Fathoni, Eva Erdayani, Linda Sukmarini, Ratih Asmana Ningrum |
| EPI_ISL_833492 | RS MMC Jakarta, Indonesia | Biosafety Level-3 Laboratory, Indonesian Institute of Sciences (LIPI) | Isa Nuryana, Ade Andriani, Anik Budhi Dharmayanthi, Syam Budi Iryanto, Andri Wardiana, Anggia Prasetyoputri, Ahmad Fathoni, Nova Dilla Yanthi, Yuliawati, Ratih Asmana Ningrum |
| EPI_ISL_833493 | RS PMI Bogor, Indonesia | Biosafety Level-3 Laboratory, Indonesian Institute of Sciences (LIPI) | Isa Nuryana, Ade Andriani, Anik Budhi Dharmayanthi, Syam Budi Iryanto, Andri Wardiana, Anggia Prasetyoputri, Ahmad Fathoni, Indriawati, Nurulliawaty, Ratih Asmana Ningrum |
| EPI_ISL_833494 | RS PMI Bogor, Indonesia | Biosafety Level-3 Laboratory, Indonesian Institute of Sciences (LIPI) | Ade Andriani, Anik Budhi Dharmayanthi, Syam Budi Iryanto, Andri Wardiana, Anggia Prasetyoputri, Isa Nuryana, Ahmad Fathoni, Maritsa Nurfatwa, Anky Zannati, Ratih Asmana Ningrum |
| EPI_ISL_833495 | Klinik Apotek Dein, Jakarta, Indonesia | Biosafety Level-3 Laboratory, Indonesian Institute of Sciences (LIPI) | Ade Andriani, Anik Budhi Dharmayanthi, Syam Budi Iryanto, Andri Wardiana, Anggia Prasetyoputri, Isa Nuryana, Ahmad Fathoni, Hariyatun, Akhirta Atikana, Ratih Asmana Ningrum |
| EPI_ISL_833496 | RS PMI Bogor, Indonesia | Biosafety Level-3 Laboratory, Indonesian Institute of Sciences (LIPI) | Anik Budhi Dharmayanthi, Syam Budi Iryanto, Andri Wardiana, Anggia Prasetyoputri, Isa Nuryana, Ade Andriani, Ahmad Fathoni, Arie Wulandari, Yana Rubiyana, Ratih Asmana Ningrum |
| EPI_ISL_833497 | Klinik Apotek Dein, Jakarta, Indonesia | Biosafety Level-3 Laboratory, Indonesian Institute of Sciences (LIPI) | Syam Budi Iryanto, Andri Wardiana, Anggia Prasetyoputri, Anik Budhi Dharmayanthi, Isa Nuryana, Ade Andriani, Ahmad Fathoni, Saiful anwar, Pamella Apriliana, Ratih Asmana Ningrum |
| EPI_ISL_833498 | Kartika Pulomas Hospital, Jakarta, Indonesia | Biosafety Level-3 Laboratory, Indonesian Institute of Sciences (LIPI) | Anggia Prasetyoputri, Isa Nuryana, Ade Andriani, Anik Budhi Dharmayanthi, Syam Budi Iryanto, Andri Wardiana, Ahmad Fathoni, Winda Tasia, Pinka Alisa Diena Shafa, Ratih Asmana Ningrum |
| EPI_ISL_833499 | RS Qadr, Tangerang, Indonesia | Biosafety Level-3 Laboratory, Indonesian Institute of Sciences (LIPI) | Isa Nuryana, Ade Andriani, Ahmad Fathoni, Anik Budhi Dharmayanthi, Syam Budi Iryanto, Andri Wardiana, Anggia Prasetyoputri, Ratna Dwi Ramadani, Sugiyono Saputra, Ratih Asmana Ningrum |
| EPI_ISL_833500 | Klinik Apotek Dein, Jakarta, Indonesia | Biosafety Level-3 Laboratory, Indonesian Institute of Sciences (LIPI) | Anik Budhi Dharmayanthi, Syam Budi Iryanto, Andri Wardiana, Anggia Prasetyoputri, Isa Nuryana, Ade Andriani, Ahmad Fathoni, Ratih Asmana Ningrum |
| EPI_ISL_833501 | RS Sayang Bunda, Makasar, Indonesia | Biosafety Level-3 Laboratory, Indonesian Institute of Sciences (LIPI) | Syam Budi Iryanto, Andri Wardiana, Anggia Prasetyoputri, Anik Budhi Dharmayanthi, Isa Nuryana, Ade Andriani, Ahmad Fathoni, Herjuno Ari Eko Nugroho, Masrukhin, Ratih Asmana Ningrum |
| EPI_ISL_833502 | RS Sayang Bunda, Makasar, Indonesia | Biosafety Level-3 Laboratory, Indonesian Institute of Sciences (LIPI) | Andri Wardiana, Anggia Prasetyoputri, Anik Budhi Dharmayanthi, Syam Budi Iryanto,I sa Nuryana, Ade Andriani, Ahmad Fathoni, Onny Nurrahman Marwayana, Fajarrudin Ahmad, Ratih Asmana Ningrum |
| EPI_ISL_833503 | RS MMC, Jakarta, Indonesia | Biosafety Level-3 Laboratory, Indonesian Institute of Sciences (LIPI) | Anggia Prasetyoputri, Isa Nuryana, Anik Budhi Dharmayanthi, Syam Budi Iryanto, Andri Wardiana, Ade Andriani, Ahmad Fathoni, Idris, Ruby Setiawan, Ratih Asmana Ningrum |
| EPI_ISL_833504 | RS MMC, Jakarta, Indonesia | Biosafety Level-3 Laboratory, Indonesian Institute of Sciences (LIPI) | Isa Nuryana, Anik Budhi Dharmayanthi, Syam Budi Iryanto, Andri Wardiana, Anggia Prasetyoputri, Ade Andriani, Ahmad Fathoni, Ahmad Randy, Listiana Oktavia, Ratih Asmana Ningrum |
| EPI_ISL_833505 | RS MMC, Jakarta, Indonesia | Biosafety Level-3 Laboratory, Indonesian Institute of Sciences (LIPI) | Ade Andriani, Anik Budhi Dharmayanthi, Syam Budi Iryanto, Andri Wardiana, Anggia Prasetyoputri, Isa Nuryana, Ahmad Fathoni, Muhammad Ridwan, Ratih Asmana Ningrum |
| EPI_ISL_833506 | RS PMI, Bogor, Indonesia | Biosafety Level-3 Laboratory, Indonesian Institute of Sciences (LIPI) | Isa Nuryana, Ade Andriani, Ahmad Fathoni, Anik Budhi Dharmayanthi, Syam Budi Iryanto, Andri Wardiana, Anggia Prasetyoputri, Satrio Haryo Benowo, Gilang Akbariani, Ratih Asmana Ningrum |

| | | | |
|---|---|---|---|
| EPI_ISL_833507 | RS MMC, Jakarta, Indonesia | Biosafety Level-3 Laboratory, Indonesian Institute of Sciences (LIPI) | Anggia Prasetyoputri, Anik Budhi Dharmayanthi, Syam Budi Iryanto, Andri Wardiana, Isa Nuryana, Ade Andriani, Ahmad Fathoni, Syaiful Rizal, Pangda Sopha, Ratih Asmana Ningrum |
| EPI_ISL_833508 | Klinik Apotek Dein, Jakarta, Indonesia | Biosafety Level-3 Laboratory, Indonesian Institute of Sciences (LIPI) | Andri Wardiana, Anggia Prasetyoputri, Anik Budhi Dharmayanthi, Syam Budi Iryanto, Isa Nuryana, Ade Andriani, Ahmad Fathoni, Ratih Asmana Ningrum |
| EPI_ISL_833509 | RS Melania, Bogor, Indonesia | Biosafety Level-3 Laboratory, Indonesian Institute of Sciences (LIPI) | Syam Budi Iryanto, Andri Wardiana, Anik Budhi Dharmayanthi, Anggia Prasetyoputri, Isa Nuryana, Ade Andriani, Ahmad Fathoni, Ratih Asmana Ningrum |
| EPI_ISL_833510 | RS Melania, Bogor, Indonesia | Biosafety Level-3 Laboratory, Indonesian Institute of Sciences (LIPI) | Anik Budhi Dharmayanthi, Syam Budi Iryanto, Andri Wardiana, Anggia Prasetyoputri, Isa Nuryana, Ade Andriani, Ahmad Fathoni, Ratih Asmana Ningrum |
| EPI_ISL_833571 | RS Qadr, Tangerang, Indonesia | Biosafety Level-3 Laboratory, Indonesian Institute of Sciences (LIPI) | Ade Andriani, Ahmad Fathoni, Anik Budhi Dharmayanthi, Syam Budi Iryanto, Andri Wardiana, Anggia Prasetyoputri, Isa Nuryana, Erwin Fajar Hasrianda, Ryan Haryo Setiawan, Ratih Asmana Ningrum |
| EPI_ISL_862039 | RSUP Dr. Sardjito | Genetics Working Group (Pokja Genetik) Faculty of Medicine, Public Health and Nursing Universitas Gadjah Mada (FK-KMK UGM); Disease Investigation Center Wates Ministry of Agriculture Indonesia; Department of Microbiology FK-KMK UGM; Laboratorium Diagnostik Yayasan Tahija World Mosquito Program (WMP) Yogyakarta Center for Tropical Medicine FK-KMK UGM; Integrated Research Center FK-KMK UGM; Department of Computer Science and Electronics FMIPA UGM; RSUP Dr. Sardjito | Gunadi, Hendra Wibawa, . Marcellus, Mohamad S. Hakim, Edwin W. Daniwijaya, Ludhang P. Rizki, Endah Supriyati, Eggi Arguni, Titik Nuryastuti, Tri Wibawa, Dwi AA Nugrahaningsih, Afiahayati, Siswanto, Kristy Iskandar, Nungki Anggorowati, Ika Trisnawati, Suma Munawar Gani, Satria Maulana, Nur Rahmi Ananda, Riat El Khair, Yunika Puspadewi, Osman Sianipar, Umi Solekhah Intansari, Elizabeth Henny Herningtiyas, Ira Puspitawati, Nur Imma Fatimah Harahap,William Widitjiarso, M |
| EPI_ISL_862040 | RSUP Dr. Sardjito | Genetics Working Group (Pokja Genetik) Faculty of Medicine, Public Health and Nursing Universitas Gadjah Mada (FK-KMK UGM); Disease Investigation Center Wates Ministry of Agriculture Indonesia; Department of Microbiology FK-KMK UGM; Laboratorium Diagnostik Yayasan Tahija World Mosquito Program (WMP) Yogyakarta Center for Tropical Medicine FK-KMK UGM; Integrated Research Center FK-KMK UGM; Department of Computer Science and Electronics FMIPA UGM; RSUP Dr. Sardjito | Gunadi, Hendra Wibawa, . Marcellus, Mohamad S. Hakim, Edwin W. Daniwijaya, Ludhang P. Rizki, Endah Supriyati, Eggi Arguni, Titik Nuryastuti, Tri Wibawa, Dwi AA Nugrahaningsih, Afiahayati, Siswanto, Kristy Iskandar, Nungki Anggorowati, Ika Trisnawati, Suma Munawar Gani, Satria Maulana, Nur Rahmi Ananda, Riat El Khair, Yunika Puspadewi, Osman Sianipar, Umi Solekhah Intansari, Elizabeth Henny Herningtiyas, Ira Puspitawati, Nur Imma Fatimah Harahap,Alvin Santoso Kali |
| EPI_ISL_862041 | RSUP Dr. Sardjito | Genetics Working Group (Pokja Genetik) Faculty of Medicine, Public Health and Nursing Universitas Gadjah Mada (FK-KMK UGM); Disease Investigation Center Wates Ministry of Agriculture Indonesia; Department of Microbiology FK-KMK UGM; Laboratorium Diagnostik Yayasan Tahija World Mosquito Program (WMP) Yogyakarta Center for Tropical Medicine FK-KMK UGM; Integrated Research Center FK-KMK UGM; Department of Computer Science and Electronics FMIPA UGM; RSUP Dr. Sardjito | Gunadi, Hendra Wibawa, . Marcellus, Mohamad S. Hakim, Edwin W. Daniwijaya, Ludhang P. Rizki, Endah Supriyati, Eggi Arguni, Titik Nuryastuti, Tri Wibawa, Dwi AA Nugrahaningsih, Afiahayati, Siswanto, Kristy Iskandar, Nungki Anggorowati, Ika Trisnawati, Suma Munawar Gani, Satria Maulana, Nur Rahmi Ananda, Riat El Khair, Yunika Puspadewi, Osman Sianipar, Umi Solekhah Intansari, Elizabeth Henny Herningtiyas, Ira Puspitawati, Nur Imma Fatimah Harahap, Dyah Ayu Puspitaran |
| EPI_ISL_872188 | RSUP Dr. Sardjito | Genetics Working Group (Pokja Genetik) Faculty of Medicine, Public Health and Nursing Universitas Gadjah Mada (FK-KMK UGM); | Gunadi, Hendra Wibawa, . Marcellus, Mohamad S. Hakim, Edwin W. Daniwijaya, Ludhang P. Rizki, Endah Supriyati, Eggi Arguni, Titik Nuryastuti, Tri Wibawa, Dwi AA Nugrahaningsih, Afiahayati, Siswanto, Kristy Iskandar, Nungki Anggorowati, Ika Trisnawati, Suma Munawar Gani, Satria Maulana, Nur Rahmi Ananda, Riat El Khair, Yunika Puspadewi, Osman Sianipar, Umi Solekhah Intansari, Elizabeth Henny Herningtiyas, Ira Puspitawati, Nur Imma Fatimah Harahap, Aditya Rifqi Fa |

| | | | |
|---|---|---|---|
| | | Disease Investigation Center Wates Ministry of Agriculture Indonesia; Department of Microbiology FK-KMK UGM; Laboratorium Diagnostik Yayasan Tahija World Mosquito Program (WMP) Yogyakarta Center for Tropical Medicine FK-KMK UGM; Integrated Research Center FK-KMK UGM; Department of Computer Science and Electronics FMIPA UGM; RSUP Dr. Sardjito | |
| EPI_ISL_872189 | RSUP Dr. Sardjito | Genetics Working Group (Pokja Genetik) Faculty of Medicine, Public Health and Nursing Universitas Gadjah Mada (FK-KMK UGM); Disease Investigation Center Wates Ministry of Agriculture Indonesia; Department of Microbiology FK-KMK UGM; Laboratorium Diagnostik Yayasan Tahija World Mosquito Program (WMP) Yogyakarta Center for Tropical Medicine FK-KMK UGM; Integrated Research Center FK-KMK UGM; Department of Computer Science and Electronics FMIPA UGM; RSUP Dr. Sardjito | Gunadi, Hendra Wibawa, . Marcellus, Mohamad S. Hakim, Edwin W. Daniwijaya, Ludhang P. Rizki, Endah Supriyati, Eggi Arguni, Titik Nuryastuti, Tri Wibawa, Dwi AA Nugrahaningsih, Afiahayati, Siswanto, Kristy Iskandar, Nungki Anggorowati, Ika Trisnawati, Suma<br>Munawar Gani, Satria Maulana, Nur Rahmi Ananda, Riat El Khair, Yunika Puspadewi, Osman Sianipar, Umi Solekhah Intansari, Elizabeth Henny Herningtiyas, Ira Puspitawati, Nur Imma Fatimah Harahap, Alvin S. Kalim |
| EPI_ISL_872190 | RSUP Dr. Sardjito | Genetics Working Group (Pokja Genetik) Faculty of Medicine, Public Health and Nursing Universitas Gadjah Mada (FK-KMK UGM); Disease Investigation Center Wates Ministry of Agriculture Indonesia; Department of Microbiology FK-KMK UGM; Laboratorium Diagnostik Yayasan Tahija World Mosquito Program (WMP) Yogyakarta Center for Tropical Medicine FK-KMK UGM; Integrated Research Center FK-KMK UGM; Department of Computer Science and Electronics FMIPA UGM; RSUP Dr. Sardjito | Gunadi, Hendra Wibawa, . Marcellus, Mohamad S. Hakim, Edwin W. Daniwijaya, Ludhang P. Rizki, Endah Supriyati, Eggi Arguni, Titik Nuryastuti, Tri Wibawa, Dwi AA Nugrahaningsih, Afiahayati, Siswanto, Kristy Iskandar, Nungki Anggorowati, Ika Trisnawati, Suma<br>Munawar Gani, Satria Maulana, Nur Rahmi Ananda, Riat El Khair, Yunika Puspadewi, Osman Sianipar, Umi Solekhah Intansari, Elizabeth Henny Herningtiyas, Ira Puspitawati, Nur Imma Fatimah Harahap, Dyah Ayu Puspitara |
| EPI_ISL_872897 | Stem Cell Research and Development Center | Stem Cell Research and Development Center | Igo Syaiful Ihsan, Eryk Hendrianto, Naimah Putri, Deya Karsari, Nora Ertanti, Aristika Dinaryanti, Purwati |
| EPI_ISL_877126 | Academic Hospital of Gajdah Mada University (RSA UGM) | Genetics Working Group (Pokja Genetik) Faculty of Medicine, Public Health and Nursing Universitas Gadjah Mada (FK-KMK UGM); Disease Investigation Center Wates Ministry of Agriculture Indonesia; Department of Microbiology FK-KMK UGM; Laboratorium Diagnostik Yayasan Tahija World Mosquito Program (WMP) Yogyakarta Center for Tropical Medicine FK-KMK UGM; Integrated Research center FK-KMK UGM; Department of Computer Science and Electronics FMIPA UGM | Gunadi, Hendra Wibawa, Marcellus, Mohamad S. Hakim, Edwin W. Daniwijaya, Ludhang P. Rizki, Endah Supriyati, Eggi Arguni, Titik Nuryastuti, Tri Wibawa, Dwi AA Nugrahaningsih, Afiahayati, Siswanto,Untung Riawan, W |
| EPI_ISL_877127 | Academic Hospital of Gajdah Mada University (RSA UGM) | Genetics Working Group (Pokja Genetik) Faculty of Medicine, Public Health and Nursing Universitas Gadjah | Gunadi, Hendra Wibawa, Marcellus, Mohamad S. Hakim, Edwin W. Daniwijaya, Ludhang P. Rizki, Endah Supriyati, Eggi Arguni, Titik Nuryastuti, Tri Wibawa, Dwi AA Nugrahaningsih, Afiahayati, Siswanto,Audric Kenny Tedja, M |

| | | | |
|---|---|---|---|
| | | Mada (FK-KMK UGM); Disease Investigation Center Wates Ministry of Agriculture Indonesia; Department of Microbiology FK-KMK UGM; Laboratorium Diagnostik Yayasan Tahija World Mosquito Program (WMP) Yogyakarta Center for Tropical Medicine FK-KMK UGM; Integrated Research center FK-KMK UGM; Department of Computer Science and Electronics FMIPA UGM | |
| EPI_ISL_877128 | Academic Hospital of Gajdah Mada University (RSA UGM) | Genetics Working Group (Pokja Genetik) Faculty of Medicine, Public Health and Nursing Universitas Gadjah Mada (FK-KMK UGM); Disease Investigation Center Wates Ministry of Agriculture Indonesia; Department of Microbiology FK-KMK UGM; Laboratorium Diagnostik Yayasan Tahija World Mosquito Program (WMP) Yogyakarta Center for Tropical Medicine FK-KMK UGM; Integrated Research center FK-KMK UGM; Department of Computer Science and Electronics FMIPA UGM | Gunadi, Hendra Wibawa, Marcellus, Mohamad S. Hakim, Edwin W. Daniwijaya, Ludhang P. Rizki, Endah Supriyati, Eggi Arguni, Titik Nuryastuti, Tri Wibawa, Dwi AA Nugrahaningsih, Afiahayati, Siswanto, Kristy Iskandar, Nungki Anggorow |
| EPI_ISL_877129 | Academic Hospital of Gajdah Mada University (RSA UGM) | Genetics Working Group (Pokja Genetik) Faculty of Medicine, Public Health and Nursing Universitas Gadjah Mada (FK-KMK UGM); Disease Investigation Center Wates Ministry of Agriculture Indonesia; Department of Microbiology FK-KMK UGM; Laboratorium Diagnostik Yayasan Tahija World Mosquito Program (WMP) Yogyakarta Center for Tropical Medicine FK-KMK UGM; Integrated Research Center FK-KMK UGM; Department of Computer Science and Electronics FMIPA UGM | Gunadi, Hendra Wibawa, Marcellus, Mohamad S. Hakim, Edwin W. Daniwijaya, Ludhang P. Rizki, Endah Supriyati, Eggi Arguni, Titik Nuryastuti, Tri Wibawa, Dwi AA Nugrahaningsih, Afiahayati, Siswanto, Kristy Iskandar, Nungki Anggorowati, |
| EPI_ISL_877130 | Academic Hospital of Gajdah Mada University (RSA UGM) | Genetics Working Group (Pokja Genetik) Faculty of Medicine, Public Health and Nursing Universitas Gadjah Mada (FK-KMK UGM); Disease Investigation Center Wates Ministry of Agriculture Indonesia; Department of Microbiology FK-KMK UGM; Laboratorium Diagnostik Yayasan Tahija World Mosquito Program (WMP) Yogyakarta Center for Tropical Medicine FK-KMK UGM; Integrated Research center FK-KMK UGM; Department of Computer Science and Electronics FMIPA UGM | Gunadi, Hendra Wibawa, Marcellus, Mohamad S. Hakim, Edwin W. Daniwijaya, Ludhang P. Rizki, Endah Supriyati, Eggi Arguni, Titik Nuryastuti, Tri Wibawa, Dwi AA Nugrahaningsih, Afiahayati, Siswanto, Kristy Iskandar, Nungki Anggorowat |
| EPI_ISL_877131 | Academic Hospital of Gajdah Mada University (RSA UGM) | Genetics Working Group (Pokja Genetik) Faculty of Medicine, Public Health and Nursing Universitas Gadjah Mada (FK-KMK UGM); Disease Investigation Center Wates Ministry of Agriculture Indonesia; Department of Microbiology | Gunadi, Hendra Wibawa, Marcellus, Mohamad S. Hakim, Edwin W. Daniwijaya, Ludhang P. Rizki, Endah Supriyati, Eggi Arguni, Titik Nuryastuti, Tri Wibawa, Dwi AA Nugrahaningsih, Afiahayati, Siswanto, Kristy Iskandar, Nungki Anggorowati, [illegible] |

| | | | |
|---|---|---|---|
| | | FK-KMK UGM; Laboratorium Diagnostik Yayasan Tahija World Mosquito Program (WMP) Yogyakarta Center for Tropical Medicine FK-KMK UGM; Integrated Research center FK-KMK UGM; Department of Computer Science and Electronics FMIPA UGM | |
| EPI_ISL_877419 | Siti Khodijah Hospital | Institute of Tropical Disease, Universitas Airlangga | Krisnoadi Rahardjo, Aldise M Nastri, Jezzy R Dewantari, Rima R Prasetya, Muhammad Hamdan, Gatot Soegiarto, Laksmi Wulandari, Resti Yudhawati, Yasuko Mori, Soetjipto, Kazufumi Shimizu, Maria I Lusi |
| EPI_ISL_877420 | Darus Syifa' Islamic Hospital | Institute of Tropical Disease, Universitas Airlangga | Aldise M Nastri, Jezzy R Dewantari, Rima R Prasetya, Krisnoadi Rahardjo, HM Faiz, Gatot Soegiarto, Laksmi Wulandari, Resti Yudhawati, Soetjipto, Yasuko Mori, Maria I Lusida, Kazufumi Shimizu |
| EPI_ISL_877421 | Institute of Tropical Disease | Institute of Tropical Disease, Universitas Airlangga | Jezzy R Dewantari, Rima R Prasetya, Krisnoadi Rahardjo, Aldise M Nastri, Gatot Soegiarto, Laksmi Wulandari, Resti Yudhawati, Soetjipto, Yasuko Mori, Maria I Lusida, Kazufumi Shimizu |
| EPI_ISL_877422 | Citra Medika Hospital | Institute of Tropical Disease, Universitas Airlangga | Rima R Prasetya, Krisnoadi Rahardjo, Aldise M Nastri, Jezzy R Dewantari, Wahidah Rachmaniyah, Gatot Soegiarto, Laksmi Wulandari, Resti Yudhawati, Soetjipto, Yasuko Mori, Maria I Lusida, Kazufumi Shin |
| EPI_ISL_877423 | Primasatya Husada Citra Hospital | Institute of Tropical Disease, Universitas Airlangga | Maria I Lusida, Krisnoadi Rahardjo, Aldise M Nastri, Jezzy R Dewantari, Rima R Prasetya, Pudji Djanuartono, Gatot Soegiarto, Laksmi Wulandari, Resti Yudhawati, Soetjipto, Yasuko Mori, Kazufumi Shimizu |
| EPI_ISL_877456 | Primasatya Husada Citra Hospital | Institute of Tropical Disease, Universitas Airlangga | Krisnoadi Rahardjo, Aldise M Nastri, Jezzy R Dewantari, Rima R Prasetya, Pudji Djanuartono, Gatot Soegiarto, Laksmi Wulandari, Resti Yudhawati, Soetjipto, Yasuko Mori, Maria I Lusida, Kazufumi Shimizu |
| EPI_ISL_877457 | Primasatya Husada Citra Hospital | Institute of Tropical Disease, Universitas Airlangga | Aldise M Nastri, Jezzy R Dewantari, Rima R Prasetya, Krisnoadi Rahardjo, Pudji Djanuartono, Gatot Soegiarto, Laksmi Wulandari, Resti Yudhawati, Soetjipto, Yasuko Mori, Maria I Lusida, Kazufumi Shimizu |
| EPI_ISL_877458 | Anwar Medika General Hospital | Institute of Tropical Disease, Universitas Airlangga | Jezzy R Dewantari, Rima R Prasetya, Krisnoadi Rahardjo, Aldise M Nastri, Nungky Taniasari, Gatot Soegiarto, Laksmi Wulandari, Resti Yudhawati, Soetjipto, Yasuko Mori, Maria I Lusida, Kazufumi Shimizu |
| EPI_ISL_877459 | Darmo Hospital | Institute of Tropical Disease, Universitas Airlangga | Rima R Prasetya, Krisnoadi Rahardjo, Aldise M Nastri, Jezzy R Dewantari, Sulung Budianto, Gatot Soegiarto, Laksmi Wulandari, Resti Yudhawati, Soetjipto, Yasuko Mori, Maria I Lusida, Kazufumi Shimizu |
| EPI_ISL_877460 | Siti Khodijah Hospital | Institute of Tropical Disease, Universitas Airlangga | Maria I Lusida, Krisnoadi Rahardjo, Aldise M Nastri, Jezzy R Dewantari, Rima R Prasetya, Muhammad Hamdan, Gatot Soegiarto, Laksmi Wulandari, Resti Yudhawati, Soetjipto, Yasuko Mori, Kazufumi Shimi |
| EPI_ISL_877538 | Siti Khodijah Hospital | Institute of Tropical Disease, Universitas Airlangga | Krisnoadi Rahardjo, Aldise M Nastri, Jezzy R Dewantari, Rima R Prasetya, Muhammad Hamdan, Gatot Soegiarto, Laksmi Wulandari, Resti Yudhawati, Soetjipto, Yasuko Mori, Maria I Lusida, Kazufumi Shimi |
| EPI_ISL_882645 | Medical Research Center, Faculty of Medicine, Syarif Hidayatullah State Islamic University Jakarta | Medical Research Center, Faculty of Medicine, Syarif Hidayatullah State Islamic University Jakarta | Chris Adhiyanto, Erike A Suwarsono, Laifa Hendarmin, Zeti Harriyati, Endah Wulandari, Flori Ratna Sari, Fika Ekayanti, Hari Hendarto |
| EPI_ISL_882646 | Medical Research Center, Faculty of Medicine, Syarif Hidayatullah State Islamic University Jakarta | Medical Research Center, Faculty of Medicine, Syarif Hidayatullah State Islamic University Jakarta | Chris Adhiyanto, Laifa Hendarmin, Zeti Harriyati, Endah Wulandari, Flori Ratna Sari, Fika Ekayanti, Hari Hendarto |
| EPI_ISL_882656 | Medical Research Center, Faculty of Medicine, Syarif Hidayatullah State Islamic University Jakarta | Medical Research Center, Faculty of Medicine, Syarif Hidayatullah State Islamic University Jakarta | Erike A Suwarsono, Chris Adhiyanto, Laifa Hendarmin, Zeti Harriyati, Flori Ratna Sari, Endah Wulandari, Fika Ekayanti |
| EPI_ISL_882769 | Institute of Tropical Disease | Institute of Tropical Disease, Universitas Airlangga | Kazufumi Shimizu, Krisnoadi Rahardjo, Aldise M Nastri, Jezzy R Dewantari, Rima R Prasetya, Gatot Soegiarto, Laksmi Wulandari, Resti Yudhawati, Yasuko Mori, Soetjipto, Maria I Lusida |
| EPI_ISL_882770 | Darus Syifa' Islamic Hospital | Institute of Tropical Disease, Universitas Airlangga | Aldise M Nastri, Jezzy R Dewantari, Rima R Prasetya, Krisnoadi Rahardjo, HM Faiz, Gatot Soegiarto, Laksmi Wulandari, Resti Yudhawati, Soetjipto, Yasuko Mori, Kazufumi Shimizu, Maria I Lusida |
| EPI_ISL_882771 | Siti Khodijah Hospital | Institute of Tropical Disease, Universitas Airlangga | Kazufumi Shimizu, Krisnoadi Rahardjo, Aldise M Nastri, Jezzy R Dewantari, Rima R Prasetya, Muhammad Hamdan, Gatot Soegiarto, Laksmi Wulandari, Resti Yudhawati, Yasuko Mori, Soetjipto, Maria I Lusi |
| EPI_ISL_883957 | Medical Research Center, Faculty of Medicine, Syarif Hidayatullah State Islamic University Jakarta | Medical Research Center, Faculty of Medicine, Syarif Hidayatullah State Islamic University Jakarta | Erike A Suwarsono, Chris Adhiyanto, Laifa Hendarmin, Zeti Harriyati, Endah Wulandari, Flori Ratna Sari, Fika Ekayanti, Hari Hendarto |
| EPI_ISL_885142 | RSUP Dr. Sardjito | Genetics Working Group | Gunadi, Hendra Wibawa, . Marcellus, Mohamad S. Hakim, Edwin W. Daniwijaya, Ludhang P. Rizki, Endah Supriyati, Eggi Arguni, Titik Nuryastuti, Tri Wibawa, Dwi AA Nugrahaningsih, Afiahayati, Siswanto, Kristy Iskandar, Nungki Anggorowati, Ika Trisnawati, Suma |

| | | | |
|---|---|---|---|
| | | (Pokja Genetik) Faculty of Medicine, Public Health and Nursing Universitas Gadjah Mada (FK-KMK UGM); Disease Investigation Center Wates Ministry of Agriculture Indonesia; Department of Microbiology FK-KMK UGM; Laboratorium Diagnostik Yayasan Tahija World Mosquito Program (WMP) Yogyakarta Center for Tropical Medicine FK-KMK UGM; Integrated Research Center FK-KMK UGM; Department of Computer Science and Electronics FMIPA UGM; RSUP Dr. Sardjito | Munawar Gani, Satria Maulana, Nur Rahmi Ananda, Riat El Khair, Yunika Puspadewi, Osman Sianipar, Umi Solekhah Intansari, Elizabeth Henny Herningtiyas, Ira Puspitawati, Nur Imma Fatimah Harahap, Alvin Santoso Kalin |
| EPI_ISL_888591 | Siti Khodijah Hospital | Institute of Tropical Disease, Universitas Airlangga | Krisnoadi Rahardjo, Aldise M Nastri, Jezzy R Dewantari, Rima R Prasetya, Muhammad Hamdan, Gatot Soegiarto, Laksmi Wulandari, Resti Yudhawati, Yasuko Mori, Soetjipto, Kazufumi Shimizu, Maria I Lusi |
| EPI_ISL_888593 | Siti Khodijah Hospital | Institute of Tropical Disease, Universitas Airlangga | Rima R Prasetya, Krisnoadi Rahardjo, Aldise M Nastri, Jezzy R Dewantari, Muhammad Hamdan, Gatot Soegiarto, Laksmi Wulandari, Resti Yudhawati, Soetjipto, Yasuko Mori, Kazufumi Shimizu, Maria I Lusi |
| EPI_ISL_888979 | RS Bhakti Mulia | Eijkman Institute for Molecular Biology, Ministry of Research and Technology/National Agency for Research and Innovation | Willy Agustine, Edison Johar, Hidayat Trimarsanto, Iskandar Adnan, Lydia V. Panggalo, Sukma Oktavianthi, Frilasita A Yudhaputri, Safarina G Malik, Khin Saw Myint, Amin Soebandrio |
| EPI_ISL_888980 | RS Anna | Eijkman Institute for Molecular Biology, Ministry of Research and Technology/National Agency for Research and Innovation | Hidayat Trimarsanto, Iskandar Adnan, Lydia V. Panggalo, Sukma Oktavianthi, Willy Agustine, Edison Johar, Frilasita A Yudhaputri, Safarina G Malik, Khin Saw Myint, Amin Soebandrio |
| EPI_ISL_888981 | RS Omni Pulomas | Eijkman Institute for Molecular Biology, Ministry of Research and Technology/National Agency for Research and Innovation | Iskandar Adnan, Lydia V. Panggalo, Sukma Oktavianthi, Willy Agustine, Edison Johar, Hidayat Trimarsanto, Frilasita A Yudhaputri, Safarina G Malik, Khin Saw Myint, Amin Soebandrio |
| EPI_ISL_888982 | RS Pondok Indah Hospital - Pondok Indah | Eijkman Institute for Molecular Biology, Ministry of Research and Technology/National Agency for Research and Innovation | Edison Johar, Frilasita A Yudhaputri, Hidayat Trimarsanto, Iskandar Adnan, Lydia V. Panggalo, Sukma Oktavianthi, Willy Agustine, Safarina G Malik, Khin Saw Myint, Amin Soebandrio |
| EPI_ISL_888983 | RS Hermina Daan Mogot | Eijkman Institute for Molecular Biology, Ministry of Research and Technology/National Agency for Research and Innovation | Edison Johar, Frilasita A Yudhaputri, Hidayat Trimarsanto, Iskandar Adnan, Lydia V. Panggalo, Sukma Oktavianthi, Willy Agustine, Safarina G Malik, Khin Saw Myint, Amin Soebandrio |
| EPI_ISL_888984 | RS Hermina Depok | Eijkman Institute for Molecular Biology, Ministry of Research and Technology/National Agency for Research and Innovation | Edison Johar, Frilasita A Yudhaputri, Hidayat Trimarsanto, Iskandar Adnan, Lydia V. Panggalo, Sukma Oktavianthi, Willy Agustine, Safarina G Malik, Khin Saw Myint, Amin Soebandrio |
| EPI_ISL_888985 | RS Hermina Serpong | Eijkman Institute for Molecular Biology, Ministry of Research and Technology/National Agency for Research and Innovation | Edison Johar, Frilasita A Yudhaputri, Hidayat Trimarsanto, Iskandar Adnan, Lydia V. Panggalo, Sukma Oktavianthi, Willy Agustine, Safarina G Malik, Khin Saw Myint, Amin Soebandrio |
| EPI_ISL_888986 | RSU Bhakti Kartini | Eijkman Institute for Molecular Biology, Ministry of Research and Technology/National Agency for Research and Innovation | Edison Johar, Frilasita A Yudhaputri, Hidayat Trimarsanto, Iskandar Adnan, Lydia V. Panggalo, Sukma Oktavianthi, Willy Agustine, Safarina G Malik, Khin Saw Myint, Amin Soebandrio |
| EPI_ISL_888987 | RS Citra Medika Depok | Eijkman Institute for Molecular Biology, Ministry of Research and Technology/National Agency for Research and Innovation | Frilasita A Yudhaputri, Hidayat Trimarsanto, Iskandar Adnan, Lydia V. Panggalo, Sukma Oktavianthi, Willy Agustine, Edison Johar, Safarina G Malik, Khin Saw Myint, Amin Soebandrio |
| EPI_ISL_888988 | RS Qadr | Eijkman Institute for | Frilasita A Yudhaputri, Hidayat Trimarsanto, Iskandar Adnan, Lydia V. Panggalo, Sukma Oktavianthi, Willy Agustine, Edison Johar, Safarina G Malik, Khin Saw Myint, Amin Soebandrio |

| | | | |
|---|---|---|---|
| | | Molecular Biology, Ministry of Research and Technology/National Agency for Research and Innovation | |
| EPI_ISL_888989, EPI_ISL_888990 | RS Pondok Indah Bintaro Jaya | Eijkman Institute for Molecular Biology, Ministry of Research and Technology/National Agency for Research and Innovation | Frilasita A Yudhaputri, Hidayat Trimarsanto, Iskandar Adnan, Lydia V. Panggalo, Sukma Oktavianthi, Willy Agustine, Edison Johar, Safarina G Malik, Khin Saw Myint, Amin Soebandrio |
| EPI_ISL_888991 | RS Mitra Keluarga Gading Serpong | Eijkman Institute for Molecular Biology, Ministry of Research and Technology/National Agency for Research and Innovation | Frilasita A Yudhaputri, Hidayat Trimarsanto, Iskandar Adnan, Lydia V. Panggalo, Sukma Oktavianthi, Willy Agustine, Edison Johar, Safarina G Malik, Khin Saw Myint, Amin Soebandrio |
| EPI_ISL_888992 | RS Mitra Keluarga Gading Serpong | Eijkman Institute for Molecular Biology, Ministry of Research and Technology/National Agency for Research and Innovation | Hidayat Trimarsanto, Iskandar Adnan, Lydia V. Panggalo, Sukma Oktavianthi, Willy Agustine, Edison Johar, Frilasita A Yudhaputri, Safarina G Malik, Khin Saw Myint, Amin Soebandrio |
| EPI_ISL_888993 | RSU Medirossa | Eijkman Institute for Molecular Biology, Ministry of Research and Technology/National Agency for Research and Innovation | Iskandar Adnan, Lydia V. Panggalo, Sukma Oktavianthi, Willy Agustine, Edison Johar, Hidayat Trimarsanto, Frilasita A Yudhaputri, Safarina G Malik, Khin Saw Myint, Amin Soebandrio |
| EPI_ISL_888994 | RSU MMC | Eijkman Institute for Molecular Biology, Ministry of Research and Technology/National Agency for Research and Innovation | Lydia V. Panggalo, Sukma Oktavianthi, Willy Agustine, Edison Johar, Hidayat Trimarsanto, Iskandar Adnan, Frilasita A Yudhaputri, Safarina G Malik, Khin Saw Myint, Amin Soebandrio |
| EPI_ISL_888995 | RSU Sumber Waras | Eijkman Institute for Molecular Biology, Ministry of Research and Technology/National Agency for Research and Innovation | Lydia V. Panggalo, Sukma Oktavianthi, Willy Agustine, Edison Johar, Hidayat Trimarsanto, Iskandar Adnan, Frilasita A Yudhaputri, Safarina G Malik, Khin Saw Myint, Amin Soebandrio |
| EPI_ISL_888996 | RSU Graha Juanda | Eijkman Institute for Molecular Biology, Ministry of Research and Technology/National Agency for Research and Innovation | Lydia V. Panggalo, Sukma Oktavianthi, Willy Agustine, Edison Johar, Hidayat Trimarsanto, Iskandar Adnan, Frilasita A Yudhaputri, Safarina G Malik, Khin Saw Myint, Amin Soebandrio |
| EPI_ISL_888997 | RSU Asysyifaa | Eijkman Institute for Molecular Biology, Ministry of Research and Technology/National Agency for Research and Innovation | Sukma Oktavianthi, Willy Agustine, Edison Johar, Hidayat Trimarsanto, Iskandar Adnan, Lydia V. Panggalo, Frilasita A Yudhaputri, Safarina G Malik, Khin Saw Myint, Amin Soebandrio |
| EPI_ISL_888998 | RSUD Cileungsi | Eijkman Institute for Molecular Biology, Ministry of Research and Technology/National Agency for Research and Innovation | Sukma Oktavianthi, Willy Agustine, Edison Johar, Hidayat Trimarsanto, Iskandar Adnan, Lydia V. Panggalo, Frilasita A Yudhaputri, Safarina G Malik, Khin Saw Myint, Amin Soebandrio |
| EPI_ISL_888999 | RS Mitra Keluarga Cibubur | Eijkman Institute for Molecular Biology, Ministry of Research and Technology/National Agency for Research and Innovation | Sukma Oktavianthi, Willy Agustine, Edison Johar, Hidayat Trimarsanto, Iskandar Adnan, Lydia V. Panggalo, Frilasita A Yudhaputri, Safarina G Malik, Khin Saw Myint, Amin Soebandrio |
| EPI_ISL_889000 | RSUD Cileungsi | Eijkman Institute for Molecular Biology, Ministry of Research and Technology/National Agency for Research and Innovation | Sukma Oktavianthi, Willy Agustine, Edison Johar, Hidayat Trimarsanto, Iskandar Adnan, Lydia V. Panggalo, Frilasita A Yudhaputri, Safarina G Malik, Khin Saw Myint, Amin Soebandrio |
| EPI_ISL_889001 | RSU Azra | Eijkman Institute for Molecular Biology, Ministry of Research and Technology/National Agency for Research and Innovation | Sukma Oktavianthi, Willy Agustine, Edison Johar, Hidayat Trimarsanto, Iskandar Adnan, Lydia V. Panggalo, Frilasita A Yudhaputri, Safarina G Malik, Khin Saw Myint, Amin Soebandrio |
| EPI_ISL_889002 | RSU Sumber Waras | Eijkman Institute for Molecular Biology, Ministry of Research and Technology/National | Sukma Oktavianthi, Willy Agustine, Edison Johar, Hidayat Trimarsanto, Iskandar Adnan, Lydia V. Panggalo, Frilasita A Yudhaputri, Safarina G Malik, Khin Saw Myint, Amin Soebandrio |

| | | | |
|---|---|---|---|
| | | Agency for Research and Innovation | |
| EPI_ISL_889003, EPI_ISL_889004 | RS Mitra Keluarga Gading Serpong | Eijkman Institute for Molecular Biology, Ministry of Research and Technology/National Agency for Research and Innovation | Willy Agustine, Edison Johar, Hidayat Trimarsanto, Iskandar Adnan, Lydia V. Panggalo, Sukma Oktavianthi, Frilasita A Yudhaputri, Safarina G Malik, Khin Saw Myint, Amin Soebandrio |
| EPI_ISL_889005 | Eijkman Institute for Molecular Biology, Ministry of Research and Technology/National Agency for Research and Innovation | Eijkman Institute for Molecular Biology, Ministry of Research and Technology/National Agency for Research and Innovation | Willy Agustine, Edison Johar, Hidayat Trimarsanto, Iskandar Adnan, Lydia V. Panggalo, Sukma Oktavianthi, Frilasita A Yudhaputri, Safarina G Malik, Khin Saw Myint, Amin Soebandrio |
| EPI_ISL_889006 | RSU Harapan Bunda | Eijkman Institute for Molecular Biology, Ministry of Research and Technology/National Agency for Research and Innovation | Willy Agustine, Edison Johar, Hidayat Trimarsanto, Iskandar Adnan, Lydia V. Panggalo, Sukma Oktavianthi, Frilasita A Yudhaputri, Safarina G Malik, Khin Saw Myint, Amin Soebandrio |
| EPI_ISL_889007 | RSUD Cileungsi | Eijkman Institute for Molecular Biology, Ministry of Research and Technology/National Agency for Research and Innovation | Willy Agustine, Edison Johar, Hidayat Trimarsanto, Iskandar Adnan, Lydia V. Panggalo, Sukma Oktavianthi, Frilasita A Yudhaputri, Safarina G Malik, Khin Saw Myint, Amin Soebandrio |
| EPI_ISL_889008 | RS Sentra Medika Cibinong | Eijkman Institute for Molecular Biology, Ministry of Research and Technology/National Agency for Research and Innovation | Edison Johar, Frilasita A Yudhaputri, Hidayat Trimarsanto, Iskandar Adnan, Lydia V. Panggalo, Sukma Oktavianthi, Willy Agustine, Safarina G Malik, Khin Saw Myint, Amin Soebandrio |
| EPI_ISL_889009 | RS Mitra Keluarga Bintaro | Eijkman Institute for Molecular Biology, Ministry of Research and Technology/National Agency for Research and Innovation | Edison Johar, Frilasita A Yudhaputri, Hidayat Trimarsanto, Iskandar Adnan, Lydia V. Panggalo, Sukma Oktavianthi, Willy Agustine, Safarina G Malik, Khin Saw Myint, Amin Soebandrio |
| EPI_ISL_889010 | RSU Medika Dramaga | Eijkman Institute for Molecular Biology, Ministry of Research and Technology/National Agency for Research and Innovation | Frilasita A Yudhaputri, Hidayat Trimarsanto, Iskandar Adnan, Lydia V. Panggalo, Sukma Oktavianthi, Willy Agustine, Edison Johar, Safarina G Malik, Khin Saw Myint, Amin Soebandrio |
| EPI_ISL_889011 | RSU Permata Cibubur | Eijkman Institute for Molecular Biology, Ministry of Research and Technology/National Agency for Research and Innovation | Hidayat Trimarsanto, Iskandar Adnan, Lydia V. Panggalo, Sukma Oktavianthi, Willy Agustine, Edison Johar, Frilasita A Yudhaputri, Safarina G Malik, Khin Saw Myint, Amin Soebandrio |
| EPI_ISL_889012 | RS Mitra Keluarga Pratama Jatiasih | Eijkman Institute for Molecular Biology, Ministry of Research and Technology/National Agency for Research and Innovation | Hidayat Trimarsanto, Iskandar Adnan, Lydia V. Panggalo, Sukma Oktavianthi, Willy Agustine, Edison Johar, Frilasita A Yudhaputri, Safarina G Malik, Khin Saw Myint, Amin Soebandrio |
| EPI_ISL_889013 | RS Rawa Lumbu | Eijkman Institute for Molecular Biology, Ministry of Research and Technology/National Agency for Research and Innovation | Iskandar Adnan, Lydia V. Panggalo, Sukma Oktavianthi, Willy Agustine, Edison Johar, Hidayat Trimarsanto, Frilasita A Yudhaputri, Safarina G Malik, Khin Saw Myint, Amin Soebandrio |
| EPI_ISL_889014 | RSU Medika Dramaga | Eijkman Institute for Molecular Biology, Ministry of Research and Technology/National Agency for Research and Innovation | Iskandar Adnan, Lydia V. Panggalo, Sukma Oktavianthi, Willy Agustine, Edison Johar, Hidayat Trimarsanto, Frilasita A Yudhaputri, Safarina G Malik, Khin Saw Myint, Amin Soebandrio |
| EPI_ISL_889015 | RS Anna | Eijkman Institute for Molecular Biology, Ministry of Research and Technology/National Agency for Research and Innovation | Iskandar Adnan, Lydia V. Panggalo, Sukma Oktavianthi, Willy Agustine, Edison Johar, Hidayat Trimarsanto, Frilasita A Yudhaputri, Safarina G Malik, Khin Saw Myint, Amin Soebandrio |
| EPI_ISL_889016 | RS Mitra Husada | Eijkman Institute for Molecular Biology, Ministry of Research and | Iskandar Adnan, Lydia V. Panggalo, Sukma Oktavianthi, Willy Agustine, Edison Johar, Hidayat Trimarsanto, Frilasita A Yudhaputri, Safarina G Malik, Khin Saw Myint, Amin Soebandrio |

| | | | |
|---|---|---|---|
| | | Technology/National Agency for Research and Innovation | |
| EPI_ISL_889017 | RS Hermina Tangerang | Eijkman Institute for Molecular Biology, Ministry of Research and Technology/National Agency for Research and Innovation | Lydia V. Panggalo, Sukma Oktavianthi, Willy Agustine, Edison Johar, Hidayat Trimarsanto, Iskandar Adnan, Frilasita A Yudhaputri, Safarina G Malik, Khin Saw Myint, Amin Soebandrio |
| EPI_ISL_889018 | RSU Medika Dramaga | Eijkman Institute for Molecular Biology, Ministry of Research and Technology/National Agency for Research and Innovation | Lydia V. Panggalo, Sukma Oktavianthi, Willy Agustine, Edison Johar, Hidayat Trimarsanto, Iskandar Adnan, Frilasita A Yudhaputri, Safarina G Malik, Khin Saw Myint, Amin Soebandrio |
| EPI_ISL_889019 | RSIA Sayyidah | Eijkman Institute for Molecular Biology, Ministry of Research and Technology/National Agency for Research and Innovation | Lydia V. Panggalo, Sukma Oktavianthi, Willy Agustine, Edison Johar, Hidayat Trimarsanto, Iskandar Adnan, Frilasita A Yudhaputri, Safarina G Malik, Khin Saw Myint, Amin Soebandrio |
| EPI_ISL_889020 | RS Omni Cikarang | Eijkman Institute for Molecular Biology, Ministry of Research and Technology/National Agency for Research and Innovation | Edison Johar, Frilasita A Yudhaputri, Hidayat Trimarsanto, Iskandar Adnan, Lydia V. Panggalo, Sukma Oktavianthi, Willy Agustine, Safarina G Malik, Khin Saw Myint, Amin Soebandrio |
| EPI_ISL_889021 | RSIA Kenari Graha Medika | Eijkman Institute for Molecular Biology, Ministry of Research and Technology/National Agency for Research and Innovation | Hidayat Trimarsanto, Iskandar Adnan, Lydia V. Panggalo, Sukma Oktavianthi, Willy Agustine, Edison Johar, Frilasita A Yudhaputri, Safarina G Malik, Khin Saw Myint, Amin Soebandrio |
| EPI_ISL_889022 | RS Anna | Eijkman Institute for Molecular Biology, Ministry of Research and Technology/National Agency for Research and Innovation | Hidayat Trimarsanto, Iskandar Adnan, Lydia V. Panggalo, Sukma Oktavianthi, Willy Agustine, Edison Johar, Frilasita A Yudhaputri, Safarina G Malik, Khin Saw Myint, Amin Soebandrio |
| EPI_ISL_890185 | Academic Hospital of Gadjah Mada University (RSA UGM) | Genetics Working Group (Pokja Genetik) Faculty of Medicine, Public Health and Nursing Universitas Gadjah Mada (FK-KMK UGM); Disease Investigation Center Wates Ministry of Agriculture Indonesia; Department of Microbiology FK-KMK UGM; Laboratorium Diagnostik Yayasan Tahija World Mosquito Program (WMP) Yogyakarta Center for Tropical Medicine FK-KMK UGM; Integrated Research center FK-KMK UGM; Department of Computer Science and Electronics FMIPA UGM | Gunadi, Hendra Wibawa, Marcellus, Mohamad S. Hakim, Edwin W. Daniwijaya, Ludhang P. Rizki, Endah Supriyati, Eggi Arguni, Titik Nuryastuti, Tri Wibawa, Dwi AA Nugrahaningsih, Afiahayati, Siswanto, Kristy Iskandar, Nungki Anggorow |
| EPI_ISL_890186 | Academic Hospital of Gadjah Mada University (RSA UGM) | Genetics Working Group (Pokja Genetik) Faculty of Medicine, Public Health and Nursing Universitas Gadjah Mada (FK-KMK UGM); Disease Investigation Center Wates Ministry of Agriculture Indonesia; Department of Microbiology FK-KMK UGM; Laboratorium Diagnostik Yayasan Tahija World Mosquito Program (WMP) Yogyakarta Center for Tropical Medicine FK-KMK UGM; Integrated Research center FK-KMK UGM; Department of Computer Science and Electronics FMIPA UGM | Gunadi, Hendra Wibawa, Marcellus, Mohamad S. Hakim, Edwin W. Daniwijaya, Ludhang P. Rizki, Endah Supriyati, Eggi Arguni, Titik Nuryastuti, Tri Wibawa, Dwi AA Nugrahaningsih, Afiahayati, Siswanto, Kristy Iskandar, Nungki Anggorowati, Au |
| EPI_ISL_890187 | Academic Hospital of Gadjah Mada University (RSA UGM) | Genetics Working Group (Pokja Genetik) Faculty of Medicine, Public Health and Nursing Universitas Gadjah | Gunadi, Hendra Wibawa, Marcellus, Mohamad S. Hakim, Edwin W. Daniwijaya, Ludhang P. Rizki, Endah Supriyati, Eggi Arguni, Titik Nuryastuti, Tri Wibawa, Dwi AA Nugrahaningsih, Afiahayati, Siswanto, Kristy Iskandar, Nungki Anggorowati |

| | | | |
|---|---|---|---|
| | | Mada (FK-KMK UGM); Disease Investigation Center Wates Ministry of Agriculture Indonesia; Department of Microbiology FK-KMK UGM; Laboratorium Diagnostik Yayasan Tahija World Mosquito Program (WMP) Yogyakarta Center for Tropical Medicine FK-KMK UGM; Integrated Research center FK-KMK UGM; Department of Computer Science and Electronics FMIPA UGM | |
| EPI_ISL_891151 | Academic Hospital of Gadjah Mada University (RSA UGM) | Genetics Working Group (Pokja Genetik) Faculty of Medicine, Public Health and Nursing Universitas Gadjah Mada (FK-KMK UGM); Disease Investigation Center Wates Ministry of Agriculture Indonesia; Department of Microbiology FK-KMK UGM; Laboratorium Diagnostik Yayasan Tahija World Mosquito Program (WMP) Yogyakarta Center for Tropical Medicine FK-KMK UGM; Integrated Research center FK-KMK UGM; Department of Computer Science and Electronics FMIPA UGM | Gunadi, Hendra Wibawa, Marcellus, Mohamad S. Hakim, Edwin W. Daniwijaya, Ludhang P. Rizki, Endah Supriyati, Eggi Arguni, Titik Nuryastuti, Tri Wibawa, Dwi AA Nugrahaningsih, Afiahayati, Siswanto, Kristy Iskandar, Nungki Anggorowati, [illegible] |
| EPI_ISL_902737 | RSA UGM | Genetics Working Group (Pokja Genetik) Faculty of Medicine, Public Health and Nursing Universitas Gadjah Mada (FK-KMK UGM); Disease Investigation Center Wates Ministry of Agriculture Indonesia; Department of Microbiology FK-KMK UGM; Laboratorium Diagnostik Yayasan Tahija World Mosquito Program (WMP) Yogyakarta Center for Tropical Medicine FK-KMK UGM; Integrated Research center FK-KMK UGM; Department of Computer Science and Electronics FMIPA UGM | Gunadi, Hendra Wibawa, Marcellus, Mohamad S. Hakim, Edwin W. Daniwijaya, Ludhang P. Rizki, Endah Supriyati, Eggi Arguni, Titik Nuryastuti, Tri Wibawa, Dwi AA Nugrahaningsih, Afiahayati, Siswanto, Kristy Iskandar, Nungki Anggorow[illegible] |
| EPI_ISL_902749 | RSA UGM | Genetics Working Group (Pokja Genetik) Faculty of Medicine, Public Health and Nursing Universitas Gadjah Mada (FK-KMK UGM); Disease Investigation Center Wates Ministry of Agriculture Indonesia; Department of Microbiology FK-KMK UGM; Laboratorium Diagnostik Yayasan Tahija World Mosquito Program (WMP) Yogyakarta Center for Tropical Medicine FK-KMK UGM; Integrated Research center FK-KMK UGM; Department of Computer Science and Electronics FMIPA UGM | Gunadi, Hendra Wibawa, Marcellus, Mohamad S. Hakim, Edwin W. Daniwijaya, Ludhang P. Rizki, Endah Supriyati, Eggi Arguni, Titik Nuryastuti, Tri Wibawa, Dwi AA Nugrahaningsih, Afiahayati, Siswanto, Kristy Iskandar, Nungki Anggorowati, [illegible] |
| EPI_ISL_902917 | Tanjungpura University Hospital | Tanjungpura University Hospital | Andriani; Mahyarudin; Novianry V; Liana DF; Shofiyah SS Astuti P; Kahtan MI; Rialita A; Putri EA; Windarti W; Sastriawan W |
| EPI_ISL_902918 | Tanjungpura University Hospital | Tanjungpura University Hospital | Andriani; Mahyarudin; Virhan Novianry ; Delima Fajar Liana; Sofi Siti Shofiyah; Puji Astuti P; Muhammad Ibnu Kahtan; Ambar Rialita; Eka Ardiani Putri EA; Wiwik Windarti ; Helmi Sastriawan; Willy Handok[illegible] |
| EPI_ISL_902919 | Tanjungpura University Hospital | Tanjungpura University Hospital | Andriani; Mahyarudin; Virhan Novianry ; Delima Fajar Liana; Sofi Siti Shofiyah; Puji Astuti P; Muhammad Ibnu Kahtan; Ambar Rialita; Eka Ardiani Putri EA; Wiwik Windarti ; Helmi Sastriawan; Arif Wicakson[illegible] |
| EPI_ISL_902921 | Tanjungpura University Hospital | Tanjungpura University Hospital | Puji Astuti; Andriani; Mahyarudin; Delima Fajar Liana; Virhan Novianry ; Sofi Siti Shofiyah |

| | | | |
|---|---|---|---|
| EPI_ISL_902922 | Tanjungpura University Hospital | Tanjungpura University Hospital | Andriani; Mahyarudin; Virhan Novianry ; Delima Fajar Liana; Sofi Siti Shofiyah; Puji Astuti; Muhammad Ibnu Kahtan; Ambar Rialita; Eka Ardiani Putri; Wiwik Windarti ; Helmi Sastriawan |
| EPI_ISL_902924 | Tanjungpura University Hospital | Tanjungpura University Hospital | Andriani; Mahyarudin; Virhan Novianry; Delima Fajar Liana; Sofi Siti Shofiyah; Puji Astuti ; Muhammad Ibnu Kahtan; Ambar Rialita; Eka Ardiani Putri; Wiwik Windarti; Helmi Sastriawan |
| EPI_ISL_905731 | Academic Hospital of Gadjah Mada University (RSA UGM) | Genetics Working Group (Pokja Genetik) Faculty of Medicine, Public Health and Nursing Universitas Gadjah Mada (FK-KMK UGM); Disease Investigation Center Wates Ministry of Agriculture Indonesia; Department of Microbiology FK-KMK UGM; Laboratorium Diagnostik Yayasan Tahija World Mosquito Program (WMP) Yogyakarta Center for Tropical Medicine FK-KMK UGM; Integrated Research center FK-KMK UGM; Department of Computer Science and Electronics FMIPA UGM | Gunadi, Hendra Wibawa, Marcellus, Mohamad S. Hakim, Edwin W. Daniwijaya, Ludhang P. Rizki, Endah Supriyati, Eggi Arguni, Titik Nuryastuti, Tri Wibawa, Dwi AA Nugrahaningsih, Afiahayati, Siswanto, Kristy Iskandar,Nungki Anggorowati, |
| EPI_ISL_906050 | RSA UGM | Genetics Working Group (Pokja Genetik) Faculty of Medicine, Public Health and Nursing Universitas Gadjah Mada (FK-KMK UGM); Disease Investigation Center Wates Ministry of Agriculture Indonesia; Department of Microbiology FK-KMK UGM; Laboratorium Diagnostik Yayasan Tahija World Mosquito Program (WMP) Yogyakarta Center for Tropical Medicine FK-KMK UGM; Integrated Research center FK-KMK UGM; Department of Computer Science and Electronics FMIPA UGM | Gunadi, Hendra Wibawa, Marcellus, Mohamad S. Hakim, Edwin W. Daniwijaya, Ludhang P. Rizki, Endah Supriyati, Eggi Arguni, Titik Nuryastuti, Tri Wibawa, Dwi AA Nugrahaningsih, Afiahayati, Siswanto,Kristy Iskandar,Nungki Anggorowati, |
| EPI_ISL_906051 | RSA UGM | Genetics Working Group (Pokja Genetik) Faculty of Medicine, Public Health and Nursing Universitas Gadjah Mada (FK-KMK UGM); Disease Investigation Center Wates Ministry of Agriculture Indonesia; Department of Microbiology FK-KMK UGM; Laboratorium Diagnostik Yayasan Tahija World Mosquito Program (WMP) Yogyakarta Center for Tropical Medicine FK-KMK UGM; Integrated Research center FK-KMK UGM; Department of Computer Science and Electronics FMIPA UGM | Gunadi, Hendra Wibawa, Marcellus, Mohamad S. Hakim, Edwin W. Daniwijaya, Ludhang P. Rizki, Endah Supriyati, Eggi Arguni, Titik Nuryastuti, Tri Wibawa, Dwi AA Nugrahaningsih, Afiahayati, Siswanto, Kristy Iskandar, Nungki Anggorowat |
| EPI_ISL_906052 | RSA UGM | Genetics Working Group (Pokja Genetik) Faculty of Medicine, Public Health and Nursing Universitas Gadjah Mada (FK-KMK UGM); Disease Investigation Center Wates Ministry of Agriculture Indonesia; Department of Microbiology FK-KMK UGM; Laboratorium Diagnostik Yayasan Tahija World Mosquito Program (WMP) Yogyakarta Center for Tropical Medicine FK-KMK UGM; Integrated Research center FK-KMK UGM; Department of Computer Science and Electronics FMIPA UGM | Gunadi, Hendra Wibawa, Marcellus, Mohamad S. Hakim, Edwin W. Daniwijaya, Ludhang P. Rizki, Endah Supriyati, Eggi Arguni, Titik Nuryastuti, Tri Wibawa, Dwi AA Nugrahaningsih, Afiahayati, Siswanto, Kristy Iskandar, Nungki Anggorowati |

| | | | |
|---|---|---|---|
| EPI_ISL_909772 | National Institute of Health Research and Development | National Institute of Health Research and Development | Holy Arif Wibowo; Ririn Ramadhany; Yuni Rukminiati; Agustiningsih; Kindi Adam; Hana Apsari Pawestri; Subangkit; Kartika Dewi Puspa; Arie Ardiansyah Nugraha; Hartanti Dian Ikawati; Krisna Nur Andriana Pangesti; Triyani Soekarso; Ni Ketut Susilarini; Nur Ika Herman; Nike Susanti; Herna; Tati Febriyanti; Juwita Kurniawati; Natalie Laurencia Kipuw; Fauzul Muna; Irene Lorinda Indalao; Aulia Rizki; Nelly Puspandari; vivi Setiawaty |
| EPI_ISL_910014 | National Institute of Health Research and Development | National Institute of Health Research and Development | Agustiningsih; Kindi Adam; Holy Arif Wibowo; Ririn Ramadhany; Yuni Rukminiati; Hana Apsari Pawestri; Subangkit; Kartika Dewi Puspa; Arie Ardiansyah Nugraha; Hartanti Dian Ikawati; Krisna Nur Andriana Pangesti; Triyani Soekarso; Ni Ketut Susilarini; Nur Ika Herman; Nike Susanti; Herna; Tati Febriyanti; Muhammad Nurhadi; Juwita Kurniawati; Natalie Laurencia Kipuw; Fauzul Muna; Irene Lorinda Indalao; Aulia Rizki; Nelly Puspandari; Vivi Setiawaty |
| EPI_ISL_911633, EPI_ISL_911675 | Tanjungpura University Hospital | Tanjungpura University Hospital | Mahyarudin; Andriani; Virhan Novianry; Delima Fajar Liana ; Sofi Siti Shofiyah; Puji Astuti ; Muhammad Ibnu Kahtan ; Ambar Rialita; Eka Ardiani Putri; Wiwik Windarti; Helmi Sastriawan |
| EPI_ISL_911707 | Academic Hospital of Gadjah Mada University (RSA UGM) | Genetics Working Group (Pokja Genetik) Faculty of Medicine, Public Health and Nursing Universitas Gadjah Mada (FK-KMK UGM); Disease Investigation Center Wates Ministry of Agriculture Indonesia; Department of Microbiology FK-KMK UGM; Laboratorium Diagnostik Yayasan Tahija World Mosquito Program (WMP) Yogyakarta Center for Tropical Medicine FK-KMK UGM; Integrated Research center FK-KMK UGM; Department of Computer Science and Electronics FMIPA UGM | Gunadi, Hendra Wibawa, Marcellus, Mohamad S. Hakim, Edwin W. Daniwijaya, Ludhang P. Rizki, Endah Supriyati, Eggi Arguni, Titik Nuryastuti, Tri Wibawa, Dwi AA Nugrahaningsih, Afiahayati, Siswanto, Kristy Iskandar, Nungki Anggorowa |
| EPI_ISL_911708 | Tanjungpura University Hospital | Tanjungpura University Hospital | Puji Astuti; Andriani; Mahyarudin; Delima Fajar Liana; Virhan Novianry ; Sofi Siti Shofiyah |
| EPI_ISL_911709 | Academic Hospital of Gadjah Mada University (RSA UGM) | Genetics Working Group (Pokja Genetik) Faculty of Medicine, Public Health and Nursing Universitas Gadjah Mada (FK-KMK UGM); Disease Investigation Center Wates Ministry of Agriculture Indonesia; Department of Microbiology FK-KMK UGM; Laboratorium Diagnostik Yayasan Tahija World Mosquito Program (WMP) Yogyakarta Center for Tropical Medicine FK-KMK UGM; Integrated Research center FK-KMK UGM; Department of Computer Science and Electronics FMIPA UGM | Gunadi, Hendra Wibawa, Marcellus, Mohamad S. Hakim, Edwin W. Daniwijaya, Ludhang P. Rizki, Endah Supriyati, Eggi Arguni, Titik Nuryastuti, Tri Wibawa, Dwi AA Nugrahaningsih, Afiahayati, Siswanto, Kristy Iskandar, Nungki Anggorowati, |
| EPI_ISL_911732 | Tanjungpura University Hospital | Tanjungpura University Hospital | Delima Fajar Liana; Mahyarudin; Andriani; Virhan Novianry; Sofi Siti Shofiyah; Puji Astuti; Muhammad Ibnu Kahtan; Ambar Rialita; Eka Ardiani Putri; Wiwik Windarti; Helmi Sastriawan |
| EPI_ISL_911750 | Tanjungpura University Hospital | Tanjungpura University Hospital | Virhan Novianry; Delima Fajar Liana; Mahyarudin; Andriani; Sofi Siti Shofiyah; Puji Astuti; Muhammad Kahtan; Ambar Rialita; Eka Ardiani Putri; Wiwik Windarti; Helmi Sastriawan, Rini Andriani |
| EPI_ISL_933575, EPI_ISL_933576 | Medical Research Center, Faculty of Medicine, Syarif Hidayatullah State Islamic University Jakarta | Medical Research Center, Faculty of Medicine, Syarif Hidayatullah State Islamic University Jakarta | Erike A Suwarsono, Chris Adhiyanto, Laifa Hendarmin, Zeti Harriyati, Endah Wulandari, Flori Ratna Sari, Fika Ekayanti, Hari Hendarto |
| EPI_ISL_942960, EPI_ISL_942971 | National Institute of Health Research and Development | National Institute of Health Research and Development | Subangkit, Hana Apsari Pawestri, Kartika Dewi Puspa, Arie Ardiansyah Nugraha, Hartanti Dian Ikawati, Krisna Nur Andriana Pangesti, Yuni Rukminiati, Ririn Ramadhany, Agustiningsih, Kindi Adam, Holy Arif Wibowo, Triyani Soekarso, Ni Ketut Susilarini, Nurika Har Tati Febriyanti, Natalie Laurencia Kipuw, Fauzul Muna, Irene Lorinda Indalao, Nelly Puspandari, Vivi Setiawaty. |
| EPI_ISL_943548 | National Institute of Health Research and Development | National Institute of Health Research and Development | Subangkit, Hana Apsari Pawestri, Kartika Dewi Puspa, Arie Ardiansyah Nugraha, Hartanti Dian Ikawati, Krisna Nur Andriana Pangesti, Yuni Rukminiati, Ririn Ramadhany, Agustiningsih, Kindi Adam, Holy Arif Wibowo, Triyani Soekarso, Ni Ketut Susilarini, Nurika Har Tati Febriyanti, Natalie Laurencia Kipuw, Fauzul Muna, Irene Lorinda Indalao, Nelly Puspandari, Vivi Setiawaty |
| EPI_ISL_947247 | RSU Bunda Mulia | Eijkman Institute for Molecular Biology, Ministry of Research and Technology/National Agency for Research and Innovation | Edison Johar, Frilasita A Yudhaputri, Hidayat Trimarsanto, Iskandar Adnan, Lydia V. Panggalo, Sukma Oktavianthi, Willy Agustine, Safarina G Malik, Khin Saw Myint, Amin Soebandrio |
| EPI_ISL_947249, EPI_ISL_947252 | RS Bina Husada Cibinong | Eijkman Institute for Molecular Biology, Ministry of Research and Technology/National Agency for Research and Innovation | Edison Johar, Frilasita A Yudhaputri, Hidayat Trimarsanto, Iskandar Adnan, Lydia V. Panggalo, Sukma Oktavianthi, Willy Agustine, Safarina G Malik, Khin Saw Myint, Amin Soebandrio |
| EPI_ISL_947255 | RSU Meilia | Eijkman Institute for Molecular Biology, Ministry of Research and | Edison Johar, Frilasita A Yudhaputri, Hidayat Trimarsanto, Iskandar Adnan, Lydia V. Panggalo, Sukma Oktavianthi, Willy Agustine, Safarina G Malik, Khin Saw Myint, Amin Soebandrio |

| | | | |
|---|---|---|---|
| | | Technology/National Agency for Research and Innovation | |
| EPI_ISL_947257 | RS Bina Husada Cibinong | Eijkman Institute for Molecular Biology, Ministry of Research and Technology/National Agency for Research and Innovation | Edison Johar, Frilasita A Yudhaputri, Hidayat Trimarsanto, Iskandar Adnan, Lydia V. Panggalo, Sukma Oktavianthi, Willy Agustine, Safarina G Malik, Khin Saw Myint, Amin Soebandrio |
| EPI_ISL_947260 | RSU Bunda Mulia | Eijkman Institute for Molecular Biology, Ministry of Research and Technology/National Agency for Research and Innovation | Iskandar Adnan, Lydia V. Panggalo, Sukma Oktavianthi, Willy Agustine, Edison Johar, Hidayat Trimarsanto, Frilasita A Yudhaputri, Safarina G Malik, Khin Saw Myint, Amin Soebandrio |
| EPI_ISL_947262 | RS Qadr | Eijkman Institute for Molecular Biology, Ministry of Research and Technology/National Agency for Research and Innovation | Frilasita A Yudhaputri, Hidayat Trimarsanto, Iskandar Adnan, Lydia V. Panggalo, Sukma Oktavianthi, Willy Agustine, Edison Johar, Safarina G Malik, Khin Saw Myint, Amin Soebandrio |
| EPI_ISL_947265 | RS Sari Asih Ciledug | Eijkman Institute for Molecular Biology, Ministry of Research and Technology/National Agency for Research and Innovation | Frilasita A Yudhaputri, Hidayat Trimarsanto, Iskandar Adnan, Lydia V. Panggalo, Sukma Oktavianthi, Willy Agustine, Edison Johar, Safarina G Malik, Khin Saw Myint, Amin Soebandrio |
| EPI_ISL_947268 | RSU Anggrek Mas | Eijkman Institute for Molecular Biology, Ministry of Research and Technology/National Agency for Research and Innovation | Frilasita A Yudhaputri, Hidayat Trimarsanto, Iskandar Adnan, Lydia V. Panggalo, Sukma Oktavianthi, Willy Agustine, Edison Johar, Safarina G Malik, Khin Saw Myint, Amin Soebandrio |
| EPI_ISL_947270 | RS Mitra Keluarga Gading Serpong | Eijkman Institute for Molecular Biology, Ministry of Research and Technology/National Agency for Research and Innovation | Frilasita A Yudhaputri, Hidayat Trimarsanto, Iskandar Adnan, Lydia V. Panggalo, Sukma Oktavianthi, Willy Agustine, Edison Johar, Safarina G Malik, Khin Saw Myint, Amin Soebandrio |
| EPI_ISL_947273 | RSIA Sayyidah | Eijkman Institute for Molecular Biology, Ministry of Research and Technology/National Agency for Research and Innovation | Frilasita A Yudhaputri, Hidayat Trimarsanto, Iskandar Adnan, Lydia V. Panggalo, Sukma Oktavianthi, Willy Agustine, Edison Johar, Safarina G Malik, Khin Saw Myint, Amin Soebandrio |
| EPI_ISL_947275 | RS Umum Mulia Pajajaran | Eijkman Institute for Molecular Biology, Ministry of Research and Technology/National Agency for Research and Innovation | Frilasita A Yudhaputri, Hidayat Trimarsanto, Iskandar Adnan, Lydia V. Panggalo, Sukma Oktavianthi, Willy Agustine, Edison Johar, Safarina G Malik, Khin Saw Myint, Amin Soebandrio |
| EPI_ISL_947278 | RS Umum Ciputra Hospital Citragarden City | Eijkman Institute for Molecular Biology, Ministry of Research and Technology/National Agency for Research and Innovation | Hidayat Trimarsanto, Iskandar Adnan, Lydia V. Panggalo, Sukma Oktavianthi, Willy Agustine, Edison Johar, Frilasita A Yudhaputri, Safarina G Malik, Khin Saw Myint, Amin Soebandrio |
| EPI_ISL_947281 | RSU Medika Dramaga | Eijkman Institute for Molecular Biology, Ministry of Research and Technology/National Agency for Research and Innovation | Hidayat Trimarsanto, Iskandar Adnan, Lydia V. Panggalo, Sukma Oktavianthi, Willy Agustine, Edison Johar, Frilasita A Yudhaputri, Safarina G Malik, Khin Saw Myint, Amin Soebandrio |
| EPI_ISL_947284 | RSIA Bunda Sejati | Eijkman Institute for Molecular Biology, Ministry of Research and Technology/National Agency for Research and Innovation | Hidayat Trimarsanto, Iskandar Adnan, Lydia V. Panggalo, Sukma Oktavianthi, Willy Agustine, Edison Johar, Frilasita A Yudhaputri, Safarina G Malik, Khin Saw Myint, Amin Soebandrio |
| EPI_ISL_947287 | RSIA PKU Muhammadiyah | Eijkman Institute for Molecular Biology, Ministry of Research and Technology/National Agency for Research and Innovation | Hidayat Trimarsanto, Iskandar Adnan, Lydia V. Panggalo, Sukma Oktavianthi, Willy Agustine, Edison Johar, Frilasita A Yudhaputri, Safarina G Malik, Khin Saw Myint, Amin Soebandrio |
| EPI_ISL_947290 | RS Hermina Bitung | Eijkman Institute for Molecular Biology, Ministry of Research and Technology/National Agency for Research and Innovation | Hidayat Trimarsanto, Iskandar Adnan, Lydia V. Panggalo, Sukma Oktavianthi, Willy Agustine, Edison Johar, Frilasita A Yudhaputri, Safarina G Malik, Khin Saw Myint, Amin Soebandrio |

| | | | |
|---|---|---|---|
| EPI_ISL_947293 | Eijkman Institute for Molecular Biology, Ministry of Research and Technology/National Agency for Research and Innovation | Eijkman Institute for Molecular Biology, Ministry of Research and Technology/National Agency for Research and Innovation | Edison Johar, Frilasita A Yudhaputri, Hidayat Trimarsanto, Iskandar Adnan, Lydia V. Panggalo, Sukma Oktavianthi, Willy Agustine, Safarina G Malik, Khin Saw Myint, Amin Soebandrio |
| EPI_ISL_947296 | RSU Kartika Husada | Eijkman Institute for Molecular Biology, Ministry of Research and Technology/National Agency for Research and Innovation | Iskandar Adnan, Lydia V. Panggalo, Sukma Oktavianthi, Willy Agustine, Edison Johar, Hidayat Trimarsanto, Frilasita A Yudhaputri, Safarina G Malik, Khin Saw Myint, Amin Soebandrio |
| EPI_ISL_947298 | RSU Medirossa | Eijkman Institute for Molecular Biology, Ministry of Research and Technology/National Agency for Research and Innovation | Iskandar Adnan, Lydia V. Panggalo, Sukma Oktavianthi, Willy Agustine, Edison Johar, Hidayat Trimarsanto, Frilasita A Yudhaputri, Safarina G Malik, Khin Saw Myint, Amin Soebandrio |
| EPI_ISL_947301 | RS Mitra Keluarga Cibubur | Eijkman Institute for Molecular Biology, Ministry of Research and Technology/National Agency for Research and Innovation | Iskandar Adnan, Lydia V. Panggalo, Sukma Oktavianthi, Willy Agustine, Edison Johar, Hidayat Trimarsanto, Frilasita A Yudhaputri, Safarina G Malik, Khin Saw Myint, Amin Soebandrio |
| EPI_ISL_947304 | RSU Harapan Bunda | Eijkman Institute for Molecular Biology, Ministry of Research and Technology/National Agency for Research and Innovation | Iskandar Adnan, Lydia V. Panggalo, Sukma Oktavianthi, Willy Agustine, Edison Johar, Hidayat Trimarsanto, Frilasita A Yudhaputri, Safarina G Malik, Khin Saw Myint, Amin Soebandrio |
| EPI_ISL_947307, EPI_ISL_947309 | RS Sentra Medika Cikarang | Eijkman Institute for Molecular Biology, Ministry of Research and Technology/National Agency for Research and Innovation | Lydia V. Panggalo, Sukma Oktavianthi, Willy Agustine, Edison Johar, Hidayat Trimarsanto, Iskandar Adnan, Frilasita A Yudhaputri, Safarina G Malik, Khin Saw Myint, Amin Soebandrio |
| EPI_ISL_947312 | RSU Bunda Mulia | Eijkman Institute for Molecular Biology, Ministry of Research and Technology/National Agency for Research and Innovation | Lydia V. Panggalo, Sukma Oktavianthi, Willy Agustine, Edison Johar, Hidayat Trimarsanto, Iskandar Adnan, Frilasita A Yudhaputri, Safarina G Malik, Khin Saw Myint, Amin Soebandrio |
| EPI_ISL_947315 | RSU Harapan Bunda | Eijkman Institute for Molecular Biology, Ministry of Research and Technology/National Agency for Research and Innovation | Lydia V. Panggalo, Sukma Oktavianthi, Willy Agustine, Edison Johar, Hidayat Trimarsanto, Iskandar Adnan, Frilasita A Yudhaputri, Safarina G Malik, Khin Saw Myint, Amin Soebandrio |
| EPI_ISL_947317 | RSU Medirossa | Eijkman Institute for Molecular Biology, Ministry of Research and Technology/National Agency for Research and Innovation | Lydia V. Panggalo, Sukma Oktavianthi, Willy Agustine, Edison Johar, Hidayat Trimarsanto, Iskandar Adnan, Frilasita A Yudhaputri, Safarina G Malik, Khin Saw Myint, Amin Soebandrio |
| EPI_ISL_947320 | RS Sari Asih Ciledug | Eijkman Institute for Molecular Biology, Ministry of Research and Technology/National Agency for Research and Innovation | Sukma Oktavianthi, Willy Agustine, Edison Johar, Hidayat Trimarsanto, Iskandar Adnan, Lydia V. Panggalo, Frilasita A Yudhaputri, Safarina G Malik, Khin Saw Myint, Amin Soebandrio |
| EPI_ISL_947322 | RS Mitra Keluarga Kelapa Gading | Eijkman Institute for Molecular Biology, Ministry of Research and Technology/National Agency for Research and Innovation | Sukma Oktavianthi, Willy Agustine, Edison Johar, Hidayat Trimarsanto, Iskandar Adnan, Lydia V. Panggalo, Frilasita A Yudhaputri, Safarina G Malik, Khin Saw Myint, Amin Soebandrio |
| EPI_ISL_947323 | RS Siloam Kebon Jeruk | Eijkman Institute for Molecular Biology, Ministry of Research and Technology/National Agency for Research and Innovation | Sukma Oktavianthi, Willy Agustine, Edison Johar, Hidayat Trimarsanto, Iskandar Adnan, Lydia V. Panggalo, Frilasita A Yudhaputri, Safarina G Malik, Khin Saw Myint, Amin Soebandrio |
| EPI_ISL_947324 | RSU Medirossa | Eijkman Institute for Molecular Biology, Ministry of Research and Technology/National Agency for Research and Innovation | Sukma Oktavianthi, Willy Agustine, Edison Johar, Hidayat Trimarsanto, Iskandar Adnan, Lydia V. Panggalo, Frilasita A Yudhaputri, Safarina G Malik, Khin Saw Myint, Amin Soebandrio |
| EPI_ISL_947325 | RSIA PKU | Eijkman Institute for | Sukma Oktavianthi, Willy Agustine, Edison Johar, Hidayat Trimarsanto, Iskandar Adnan, Lydia V. Panggalo, Frilasita A Yudhaputri, Safarina G Malik, Khin Saw Myint, Amin Soebandrio |

| | | | |
|---|---|---|---|
| | Muhammadiyah | Molecular Biology, Ministry of Research and Technology/National Agency for Research and Innovation | |
| EPI_ISL_947326, EPI_ISL_947327 | RS Mitra Keluarga Gading Serpong | Eijkman Institute for Molecular Biology, Ministry of Research and Technology/National Agency for Research and Innovation | Willy Agustine, Edison Johar, Hidayat Trimarsanto, Iskandar Adnan, Lydia V. Panggalo, Sukma Oktavianthi, Frilasita A Yudhaputri, Safarina G Malik, Khin Saw Myint, Amin Soebandrio |
| EPI_ISL_947328 | RS Mitra Keluarga Depok | Eijkman Institute for Molecular Biology, Ministry of Research and Technology/National Agency for Research and Innovation | Willy Agustine, Edison Johar, Hidayat Trimarsanto, Iskandar Adnan, Lydia V. Panggalo, Sukma Oktavianthi, Frilasita A Yudhaputri, Safarina G Malik, Khin Saw Myint, Amin Soebandrio |
| EPI_ISL_947329, EPI_ISL_947330 | RSU Bunda Mulia | Eijkman Institute for Molecular Biology, Ministry of Research and Technology/National Agency for Research and Innovation | Willy Agustine, Edison Johar, Hidayat Trimarsanto, Iskandar Adnan, Lydia V. Panggalo, Sukma Oktavianthi, Frilasita A Yudhaputri, Safarina G Malik, Khin Saw Myint, Amin Soebandrio |
| EPI_ISL_949183 | National Institute of Health Research and Development | National Institute of Health Research and Development | Kindi Adam; Holy Arif Wibowo; Ririn Ramadhany; Yuni Rukminiati; Agustiningsih; Hana Apsari Pawestri; Subangkit; Kartika Dewi Puspa; Arie Ardiansyah Nugraha; Hartanti Dian Ikawati; Krisna Nur Andriana Pangesti; Triyani Soekarso; Ni Ketut Susilarini; Nur Ika [illegible] Herman; Nike Susanti; Herna; Tati Febriyanti; Natalie Laurencia Kipuw; Fauzul Muna; Irene Lorinda Indalao; Aulia Rizki; Nelly Puspandari; Vivi Setiawaty |
| EPI_ISL_949192 | National Institute of Health Research and Development | National Institute of Health Research and Development | Ririn Ramadhany; Yuni Rukminiati; Agustiningsih; Kindi Adam; Holy Arif Wibowo; Hana Apsari Pawestri; Subangkit; Kartika Dewi Puspa; Arie Ardiansyah Nugraha; Hartanti Dian Ikawati; Krisna Nur Andriana Pangesti; Ni Ketut Susilarini; Nur Ika Hariastuti; Uly A[illegible] Susanti; Herna; Tati Febriyanti; Natalie Laurentia Kipuw; Fauzul Muna; Irene Lorinda Indalao; Aulia Rizki; Nelly Puspandari; Vivi Setiawaty |
| EPI_ISL_953427 | National Institute of Health Research and Development | National Institute of Health Research and Development | Subangkit, Hana Apsari Pawestri, Kartika Dewi Puspa, Arie Ardiansyah Nugraha, Hartanti Dian Ikawati, Krisna Nur Andriana Pangesti, Yuni Rukminiati, Ririn Ramadhany, Agustiningsih, Kindi Adam, Holy Arif Wibowo, Triyani Soekarso, Ni Ketut Susilarini, Nurika Ha[illegible] Tati Febriyanti, Natalie Laurencia Kipuw, Fauzul Muna, Irene Lorinda Indalao, Nelly Puspandari, Vivi Setiawaty |
| EPI_ISL_956270 | Universitas Airlangga Hospital | Institute of Tropical Disease, Universitas Airlangga | Aldise M Nastri, Jezzy R Dewantari, Rima R Prasetya, Krisnoadi Rahardjo, Nasronudin, Gatot Soegiarto, Laksmi Wulandari, Resti Yudhawati, Soetjipto, Yasuko Mori, Maria I Lusida, Kazufumi Shimizu |
| EPI_ISL_956271 | Adi Husada Undaan Hospital | Institute of Tropical Disease, Universitas Airlangga | Aldise M Nastri, Jezzy R Dewantari, Rima R Prasetya, Krisnoadi Rahardjo, Irawati Marga, Gatot Soegiarto, Laksmi Wulandari, Retno A Setyoningrum, Resti Yudhawati, Yohko K Shimizu, Mitsuhiro Nishimura, Soetjipto, Yasuko Mori, Ma[illegible] |
| EPI_ISL_956273 | Universitas Airlangga Hospital | Institute of Tropical Disease, Universitas Airlangga | Jezzy R Dewantari, Rima R Prasetya, Krisnoadi Rahardjo, Aldise M Nastri, Nasronudin, Gatot Soegiarto, Laksmi Wulandari, Resti Yudhawati, Soetjipto, Yasuko Mori, Maria I Lusida, Kazufumi Shimizu |
| EPI_ISL_956274 | RSUD Sidoarjo | Institute of Tropical Disease, Universitas Airlangga | Rima R Prasetya, Krisnoadi Rahardjo, Aldise M Nastri, Jezzy R Dewantari, Atok Irawan, Gatot Soegiarto, Laksmi Wulandari, Resti Yudhawati, Soetjipto, Yasuko Mori, Maria I Lusida, Kazufumi Shimizu |
| EPI_ISL_956276 | Mitra Keluarga Hospital Waru | Institute of Tropical Disease, Universitas Airlangga | Maria I Lusida, Krisnoadi Rahardjo, Aldise M Nastri, Jezzy R Dewantari, Rima R Prasetya, Christina Dian Anggraeni, Gatot Soegiarto, Laksmi Wulandari, Resti Yudhawati, Soetjipto, Yasuko Mori, Kazufumi Shi[illegible] |
| EPI_ISL_956277 | Paru Hospital | Institute of Tropical Disease, Universitas Airlangga | Krisnoadi Rahardjo, Aldise M Nastri, Jezzy R Dewantari, Rima R Prasetya, Dyah Retno, Gatot Soegiarto, Laksmi Wulandari, Resti Yudhawati, Soetjipto, Yasuko Mori, Maria I Lusida, Kazufumi Shimizu |
| EPI_ISL_956278 | Siti Khodijah Hospital | Institute of Tropical Disease, Universitas Airlangga | Aldise M Nastri, Jezzy R Dewantari, Rima R Prasetya, Krisnoadi Rahardjo, Muhammad Hamdan, Gatot Soegiarto, Laksmi Wulandari, Resti Yudhawati, Soetjipto, Yasuko Mori, Maria I Lusida, Kazufumi Shimi[illegible] |
| EPI_ISL_956279 | Siti Khodijah Hospital | Institute of Tropical Disease, Universitas Airlangga | Jezzy R Dewantari, Rima R Prasetya, Krisnoadi Rahardjo, Aldise M Nastri, Muhammad Hamdan, Gatot Soegiarto, Laksmi Wulandari, Resti Yudhawati, Soetjipto, Yasuko Mori, Maria I Lusida, Kazufumi Shimi[illegible] |
| EPI_ISL_956280 | Institute of Tropical Disease | Institute of Tropical Disease, Universitas Airlangga | Rima R Prasetya, Krisnoadi Rahardjo, Aldise M Nastri, Jezzy R Dewantari, Gatot Soegiarto, Laksmi Wulandari, Resti Yudhawati, Soetjipto, Yasuko Mori, Maria I Lusida, Kazufumi Shimizu |
| EPI_ISL_956281 | Siti Khodijah Hospital | Institute of Tropical Disease, Universitas Airlangga | Maria I Lusida, Krisnoadi Rahardjo, Aldise M Nastri, Jezzy R Dewantari, Rima R Prasetya, Muhammad Hamdan, Gatot Soegiarto, Laksmi Wulandari, Resti Yudhawati, Soetjipto, Yasuko Mori, Kazufumi Shimi[illegible] |
| EPI_ISL_956307 | Primasatya Husada Citra Hospital | Institute of Tropical Disease, Universitas Airlangga | Krisnoadi Rahardjo, Aldise M Nastri, Jezzy R Dewantari, Rima R Prasetya, Pudji Djanuartono, Gatot Soegiarto, Laksmi Wulandari, Resti Yudhawati, Yasuko Mori, Soetjipto, Kazufumi Shimizu, Maria I Lusid[illegible] |
| EPI_ISL_956308 | RSUD Dr. M. Soewandhie | Institute of Tropical Disease, Universitas Airlangga | Jezzy R Dewantari, Rima R Prasetya, Krisnoadi Rahardjo, Aldise M Nastri, Febria Rachmanita, Gatot Soegiarto, Laksmi Wulandari, Resti Yudhawati, Yasuko Mori, Soetjipto, Kazufumi Shimizu, Maria I Lusid[illegible] |
| EPI_ISL_956309 | RSUD Sidoarjo | Institute of Tropical Disease, Universitas Airlangga | Rima R Prasetya, Krisnoadi Rahardjo, Aldise M Nastri, Jezzy R Dewantari, Atok Irawan, Gatot Soegiarto, Laksmi Wulandari, Resti Yudhawati, Yasuko Mori, Soetjipto, Kazufumi Shimizu, Maria I Lusida |
| EPI_ISL_956310 | RSUD Dr. M. Soewandhie | Institute of Tropical Disease, Universitas Airlangga | Aldise M Nastri, Jezzy R Dewantari, Rima R Prasetya, Krisnoadi Rahardjo, Febria Rachmanita, Gatot Soegiarto, Laksmi Wulandari, Resti Yudhawati, Yasuko Mori, Soetjipto, Kazufumi Shimizu, Maria I Lusid[illegible] |
| EPI_ISL_956311 | Islamic Hospital | Institute of Tropical Disease, Universitas Airlangga | Kazufumi Shimizu, Krisnoadi Rahardjo, Aldise M Nastri, Jezzy R Dewantari, Rima R Prasetya, Dodo Anondo, Gatot Soegiarto, Laksmi Wulandari, Resti Yudhawati, Yasuko Mori, Soetjipto, Maria I Lusida |

| | | | |
|---|---|---|---|
| EPI_ISL_956312 | Institute of Tropical Disease | Institute of Tropical Disease, Universitas Airlangga | Krisnoadi Rahardjo, Aldise M Nastri, Jezzy R Dewantari, Rima R Prasetya, Gatot Soegiarto, Laksmi Wulandari, Resti Yudhawati, Yasuko Mori, Soetjipto, Kazufumi Shimizu, Maria I Lusida |
| EPI_ISL_956313 | Siti Khodijah Hospital | Institute of Tropical Disease, Universitas Airlangga | Jezzy R Dewantari, Rima R Prasetya, Krisnoadi Rahardjo, Aldise M Nastri, Muhammad Hamdan, Gatot Soegiarto, Laksmi Wulandari, Resti Yudhawati, Yasuko Mori, Soetjipto, Kazufumi Shimizu, Maria I Lusi |
| EPI_ISL_956314 | Paru Hospital | Institute of Tropical Disease, Universitas Airlangga | Rima R Prasetya, Krisnoadi Rahardjo, Aldise M Nastri, Jezzy R Dewantari, Dyah Retno, Gatot Soegiarto, Laksmi Wulandari, Resti Yudhawati, Yasuko Mori, Soetjipto, Kazufumi Shimizu, Maria I Lusida |
| EPI_ISL_956315 | Siti Khodijah Hospital | Institute of Tropical Disease, Universitas Airlangga | Kazufumi Shimizu, Krisnoadi Rahardjo, Aldise M Nastri, Jezzy R Dewantari, Rima R Prasetya, Muhammad Hamdan, Gatot Soegiarto, Laksmi Wulandari, Resti Yudhawati, Yasuko Mori, Soetjipto, Maria I Lusi |
| EPI_ISL_956316 | Institute of Tropical Disease | Institute of Tropical Disease, Universitas Airlangga | Krisnoadi Rahardjo, Aldise M Nastri, Jezzy R Dewantari, Rima R Prasetya, Gatot Soegiarto, Laksmi Wulandari, Resti Yudhawati, Yasuko Mori, Soetjipto, Kazufumi Shimizu, Maria I Lusida |
| EPI_ISL_984264 | RSUPN Dr. Cipto Mangunkusumo | National Institute of Health Research and Development | Subangkit, Hana Apsari Pawestri, Kartika Dewi Puspa, Arie Ardiansyah Nugraha, Hartanti Dian Ikawati, Krisna Nur Andriana Pangesti, Nuri Dyah Indrasari, Dewi Wulandari, Nelly Puspandari, Vivi Setiawaty |
| EPI_ISL_984276 | National Institute of Health Research and Development | National Institute of Health Research and Development | Holy Arif Wibowo, Ririn Ramadhany, Yuni Rukminiati, Agustiningsih, Kindi Adam, Hana Apsari Pawestri, Subangkit, Kartika Dewi Puspa, Arie Ardiansyah Nugraha, Hartanti Dian Ikawati, Krisna Nur Andriana Pangesti, Ni Ketut Susilarini, Nur Ika Hariastuti, Uly Al Susanti, Herna, Tati Febriyanti, Natalie Laurencia Kipuw, Fauzul Muna, Irene Lorinda Indalao, Aulia Rizki, Nelly Puspandari, Vivi Setiawaty |
| EPI_ISL_984277 | National Institute of Health Research and Development | National Institute of Health Research and Development | Kindi Adam, Holy Arif Wibowo, Ririn Ramadhany, Yuni Rukminiati, Agustiningsih, Hana Apsari Pawestri, Subangkit, Kartika Dewi Puspa, Arie Ardiansyah Nugraha, Hartanti Dian Ikawati, Krisna Nur Andriana Pangesti, Ni Ketut Susilarini, Nur Ika Hariastuti, Uly Al Susanti, Herna, Tati Febriyanti, Natalie Laurencia Kipuw, Fauzul Muna, Irene Lorinda Indalao, Aulia Rizki, Nelly Puspandari, Vivi Setiawaty |
| EPI_ISL_984309 | National Institute of Health Research and Development | National Institute of Health Research and Development | Agustiningsih, Kindi Adam, Holy Arif Wibowo, Ririn Ramadhany, Yuni Rukminiati, Hana Apsari Pawestri, Subangkit, Kartika Dewi Puspa, Arie Ardiansyah Nugraha, Hartanti Dian Ikawati, Krisna Nur Andriana Pangesti, Ni Ketut Susilarini, Nur Ika Hariastuti, Uly Al Susanti, Herna, Tati Febriyanti, Natalie Laurencia Kipuw, Fauzul Muna, Irene Lorinda Indalao, Aulia Rizki, Nelly Puspandari, Vivi Setiawaty |
| EPI_ISL_984313 | National Institute of Health Research and Development | National Institute of Health Research and Development | Yuni Rukminiati, Agustiningsih, Kindi Adam, Holy Arif Wibowo, Ririn Ramadhany, Hana Apsari Pawestri, Subangkit, Kartika Dewi Puspa, Arie Ardiansyah Nugraha, Hartanti Dian Ikawati, Krisna Nur Andriana Pangesti, Ni Ketut Susilarini, Nur Ika Hariastuti, Uly Al Susanti, Herna, Tati Febriyanti, Natalie Laurencia Kipuw, Fauzul Muna, Irene Lorinda Indalao, Aulia Rizki, Nelly Puspandari, Vivi Setiawaty |
| EPI_ISL_985396 | RSUD Dr. Tjitrowardojo | Genetics Working Group (Pokja Genetik) Faculty of Medicine, Public Health and Nursing Universitas Gadjah Mada (FK-KMK UGM); Disease Investigation Center Wates Ministry of Agriculture Indonesia; Department of Microbiology FK-KMK UGM; Laboratorium Diagnostik Yayasan Tahija World Mosquito Program (WMP) Yogyakarta Center for Tropical Medicine FK-KMK UGM; Integrated Research Center FK-KMK UGM; Department of Computer Science and Electronics FMIPA UGM; Balai Besar Teknik Kesehatan Lingkungan dan Pengendalian Penyakit (BBTKLPP) Yogyakarta | Gunadi, Hendra Wibawa, Marcellus, Mohamad S. Hakim, Edwin W. Daniwijaya, Ludhang P. Rizki, Endah Supriyati, Eggi Arguni, Titik Nuryastuti, Tri Wibawa, Dwi AA Nugrahaningsih, Afiahayati, Siswanto, Kristy Iskandar, Nungki Anggorowati, Irene, Indaryati, Havic Sari, Dwiki afandy, Dyah Ayu Puspitarani |
| EPI_ISL_985397 | RSUD Dr. Tjitrowardojo | Genetics Working Group (Pokja Genetik) Faculty of Medicine, Public Health and Nursing Universitas Gadjah Mada (FK-KMK UGM); Disease Investigation Center Wates Ministry of Agriculture Indonesia; Department of Microbiology FK-KMK UGM; Laboratorium Diagnostik Yayasan Tahija World Mosquito Program (WMP) Yogyakarta Center for Tropical Medicine FK-KMK UGM; Integrated Research Center FK-KMK UGM; Department of Computer Science and Electronics FMIPA UGM; Balai Besar Teknik Kesehatan Lingkungan dan Pengendalian Penyakit (BBTKLPP) Yogyakarta | Gunadi, Hendra Wibawa, Marcellus, Mohamad S. Hakim, Edwin W. Daniwijaya, Ludhang P. Rizki, Endah Supriyati, Eggi Arguni, Titik Nuryastuti, Tri Wibawa, Dwi AA Nugrahaningsih, Afiahayati, Siswanto, Kristy Iskandar, Nungki Anggorowati, Irene, Indaryati, Havic Sari, Untung Riawan, William Widitjiarso |
| EPI_ISL_985398 | RSA UGM | Genetics Working Group (Pokja Genetik) Faculty of Medicine, Public Health and Nursing Universitas Gadjah | Gunadi, Hendra Wibawa, Marcellus, Mohamad S. Hakim, Edwin W. Daniwijaya, Ludhang P. Rizki, Endah Supriyati, Eggi Arguni, Titik Nuryastuti, Tri Wibawa, Dwi AA Nugrahaningsih, Afiahayati, Siswanto, Kristy Iskandar, Nungki Anggorowati, Su |

Mada (FK-KMK UGM);
Disease Investigation
Center Wates Ministry of
Agriculture Indonesia;
Department of Microbiology
FK-KMK UGM;
Laboratorium Diagnostik
Yayasan Tahija World
Mosquito Program (WMP)
Yogyakarta Center for
Tropical Medicine FK-KMK
UGM; Integrated Research
center FK-KMK UGM;
Department of Computer
Science and Electronics
FMIPA UGM
